# Gene Conversion Directed Successive Engineering of Modular Polyketide Synthases

**DOI:** 10.1101/2024.12.29.630704

**Authors:** Wenzheng Jin, Jiaming Tu, Bei Zhang, Xuri Wu, Yijun Chen

## Abstract

Modular polyketide synthases (PKSs) can produce various secondary metabolites in a collinearity fashion. Although rational engineering of modular PKS can ultimately create a diverse array of novel compounds, *de novo* generation of defined structures usually results in the loss or remarkable decline of productivity due primarily to the incompatibility of different elements. Here, we present a modular PKS engineering strategy driven by an evolutionary event of gene conversion to accomplish successive engineering of the modular PKS in cinnamomycin biosynthetic gene cluster (*cmm* BGC). By simulating the gene conversion process, *cmm* BGC is consecutively reprogrammed to generate a novel macrolide with predicted structural features. Moreover, in contrast to previous notion, the intra-module KS domain is demonstrated to associate with the selectivity of extender units. Collectively, the coordination between evolutionary consequence and functional manipulation of assembly line may shed a new light on modular PKS engineering.

**Graphical Abstract:** Gene conversion process, typically occurred in KS (purple squares) and AT (blue squares), plays an integral role of structural diversity of polyketides. In this study, an unusual gene conversion was observed in *cmm* BGC. Subsequently, a homologous BGC was obtained through genome mining by a gene conversion-associated KS domain. Under the direction of gene conversion, the modular PKS in *cmm* BGC was successively reprogrammed, resulting in *de novo* biosynthesis of a new-to-nature polyketide.

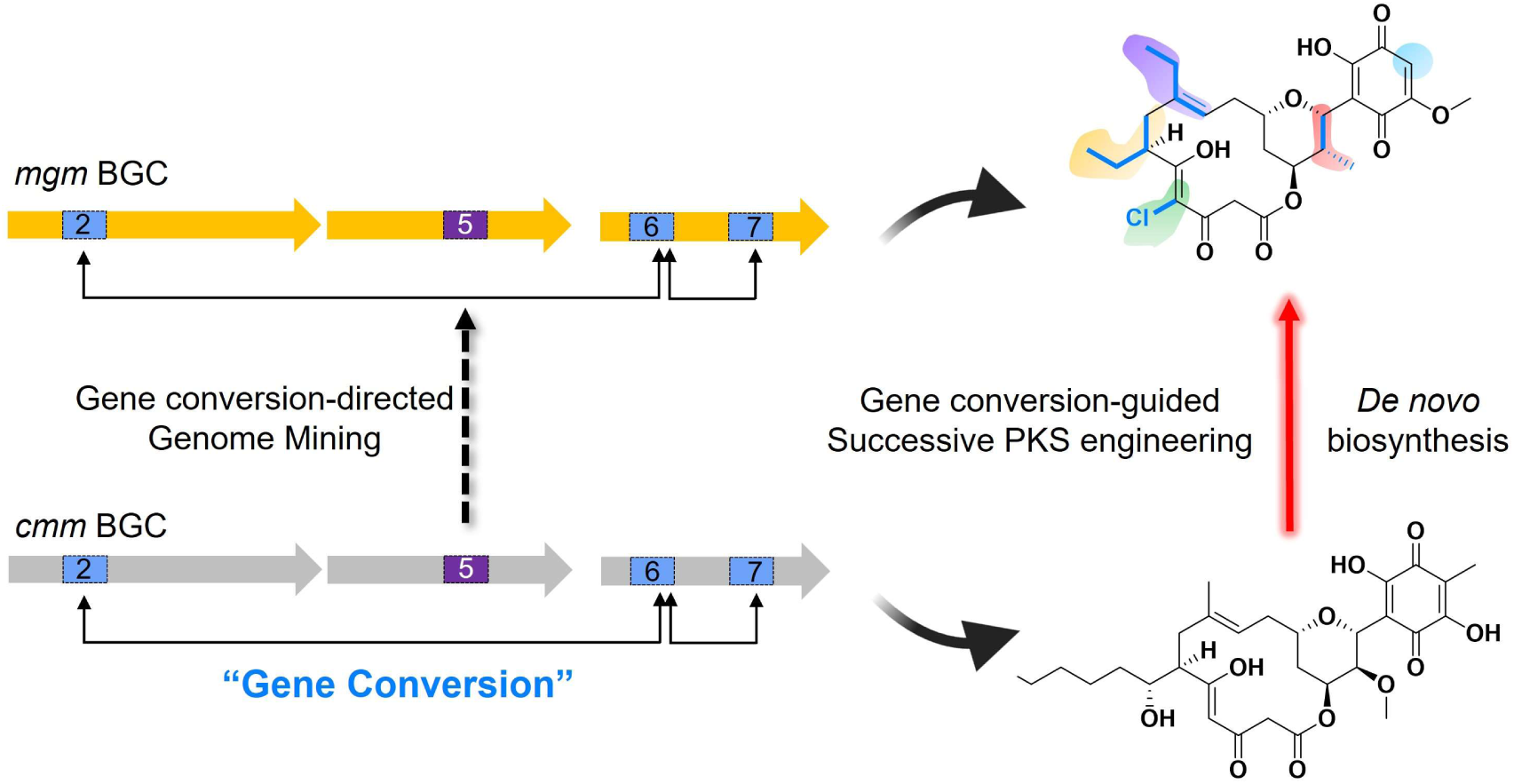

## Introduction

Polyketides are biosynthesized by polyketide synthases (PKSs) to possess remarkable structural diversities and valuable medicinal utilities^[1]^. Among different types of PKS, type I PKS functions in a modular fashion^[2,3]^, in which the modules are linearly arranged to guide the unidirectional chain extension.

Since the early investigations on modular PKSs^[4,5]^, the concept of "Lego-ization" in polyketide biosynthesis has continuously been evolved^[6]^, allowing systematic reprogramming of PKSs to produce designed natural products with high stereo- and regio-specificity^[7–9]^. At the same time, various approaches, including site-directed mutagenesis, fragment swaps and subunit insertions/deletions, have been implemented to the engineering of PKSs, aiming to side chain alteration or scaffold reconstruction. More recently, the strategies for PKS reprogramming have been moved forward from sequence-based design to structure-guided engineering^[10,11]^. However, the complex and dynamic conformations of PKSs^[12–14]^, along with sophisticated interactions and functional interdependencies of different PKS domains^[15,16]^, have posed considerable challenges for efficient engineering. Consequently, PKS assembly lines tend to become fragile following a single round of engineering, preventing successive reprogramming and applications^[17]^.

The findings on the evolutionary events in modular PKSs^[18]^, such as point mutation, gene duplication, gene loss, gene conversion, gene recombination, and horizontal gene transfer, have provided a new opportunity for PKS engineering. In 2017, Abe lab proposed an evolutionary mechanism for structural diversification of polyketides and defining unique module organizations ^[19]^. Keatinge-Clay lab demonstrated improvement on success rate by using the “Abe boundaries" for domain-fusion ^[20]^. Based on the statistics of the massive *trans*-AT PKS sequences, Piel lab applied the concept of evolutionary-guided engineering for *trans*-AT PKSs^[21]^. Collectively, the evolutionary information not only guides optimal recombination boundary, but also facilitates the engineering efforts. For example, by simulating the evolutionary recombination of homologous modules, a ring-contracted mini-azalomycin was generated ^[22]^. Regarding gene loss, the interconversions between polyene-pyrone structures, differing in the skeleton sizes, were accomplished ^[23,24]^. Although engineering of PKS remains as a try-and-correct, recent studies mutually demonstrate that evolutionary-guided engineering is broadly applicable, extending the scenarios beyond isolated and special cases.

Gene conversion is a prevalent evolutionary phenomenon observed in PKSs, whereby genetic material is exchanged between adjacent and homologous modules particularly between regions with high sequence identity^[25–27]^,. Thus, the event of gene conversion can alter specific regions for fine-tuning the chemical diversity of polyketides. This evolutionary event is widely distributed in *Streptomyces*, frequently occurring in KS and AT domains of modular PKS assembly lines according to the nucleotide homolog and phylogenetic tree analysis^[26–28]^. Consequently, the gene conversion process is regarded as an important feature for the alteration of alkyl group of the side chain of polyketide backbone. Meanwhile, the intra-module KS-AT didomain is found to often engage in gene conversion as a complete entity^[23]^. Although current knowledge on gene conversion is very limited, the preservation of exquisite compatibility of catalytic elements to steadily produce polyketides through consecutive gene conversion events may offer a promising avenue to achieve successive PKS engineering.

Previously, we unveiled an unusual type I/type III PKS hybrid *cmm* BGC and its products of cinnamomycin A-F (**1,1b, 1c** and **2**-**4**), a series of 14-membered macrolides with selective and significant anti-proliferative activity^[29]^. When we performed further analysis of *cmm* BGC, we found the presence of gene conversion with high homolog or even identical DNA fragments in module 2, 6 and 7 (Fig. 1a, Supplementary Table 1). These regions are specifically located in malonyl-CoA-specific AT domains, spanning from C-terminus of KS domain to post-AT linker. Hence, we named this evolutionary region as "AT_conversion_" because of the lack of obvious presence in other domains (Fig. 1). The 100% nucleotide sequence identity of AT_conversion_ regions between modules 2 and 6 is exceptionally rare, strongly supporting the connection of gene conversion with the biosynthesis of cinnamomycin skeleton. Therefore, previous successes in utilizing evolutionary approach for PKS engineering promoted us to test whether gene conversion process could be applied to empowering PKS engineering.

**Fig 1.**
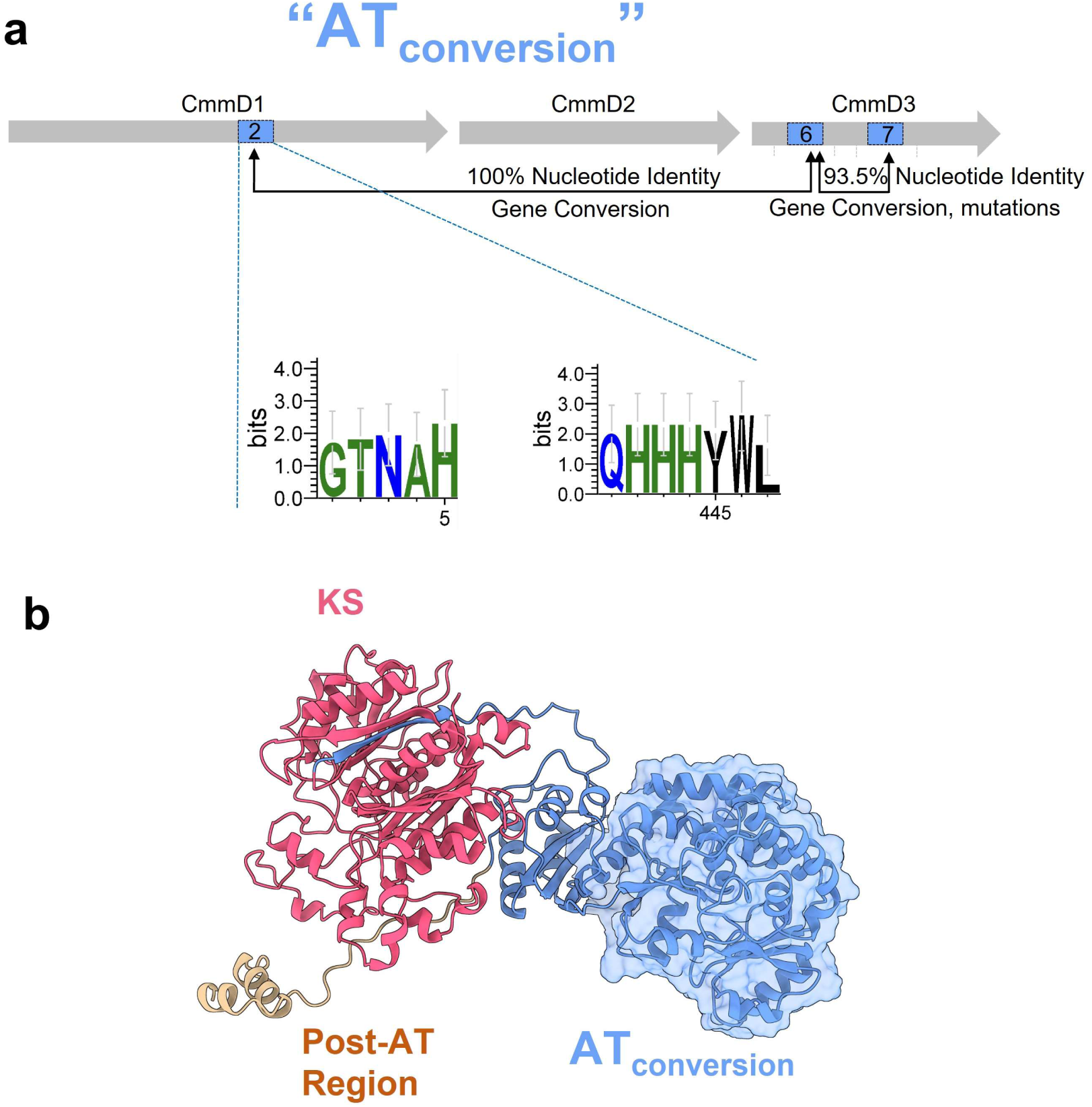
Identification of AT_conversion_ regions in *cmm* BGC and their structural features. a) The locations of AT_conversion_ regions in *cmm* BGC. Rectangles colored in blue represent AT_conversion_ regions in CmmD1 and CmmD3. Numbers in rectangles are the abbreviation of module number. The boundaries of AT_conversion_ in module 2 are shown by amino acid residues of His5 and Tyr445. b) Structural models of AT_conversion_ region created by AlphaFold 2.0. Structures colored in red represent the location neighboring KS domain; structures colored in blue represent AT_conversion_ region; structures colored in brown represent post-AT region.

In this study, a homologous *mgm* BGC for possibly producing novel macrolides in *S. mangrovisoli*^[30]^ was discovered by gene conversion-oriented genome mining. This homologous *mgm* BGC could be combined with *cmm* BGC to serve as a pair of templates for precisely exchanging genes from each other. By taking the advantages of gene conversion in both *mgm* and *cmm* BGCs, successive engineering of the modular PKSs was accomplished, resulting in the rejuvenation of *mgm* BGC for *de novo* production of mangromycin. Moreover, the intra-module KS domain was revealed to act as a proofreading element for ensuring the fidelity of extender unit incorporation.

## Results

### Discovery of a homologous *mgm* BGC by gene conversion-guided genome mining

Bacterial natural products frequently exist as a large family of structural analogues, encoded by evolutionarily related homologous biosynthetic gene clusters. Similarly, the observation of gene conversion in *cmm* BGC could lead to the discovery of cinnamomycin-type compounds and their corresponding BGCs.

Since multiple ancestral cluster-specific fragments among homologous BGCs could be preserved by the process of gene conversion, these regions should be useful probes for genome mining. First, we used an AT_conversion_ fragment (Fig. 1a) as the probe to mine BGC(s) with gene conversion characteristics in the NCBI genomic database. Unfortunately, the initial attempts did not yield any meaningful results, possibly due to a significant difference in sequence identity of AT domains caused by a large number of independent recombination events of the AT domain in homologous BGCs. Meanwhile, KS domains are usually accompanied with AT domains to possess gene conversion^[26,28]^, which might be suitable as a potential marker for genome mining. Next, a BLAST search using KS fragments in *cmm* BGC was conducted, and an uncharacterized homologous *mgm* BGC in *S. mangrovisoli* (GCF_000974985.2) was revealed (Supplementary Fig. 1). The products from this newly identified BGC were designated as mangromycins.

While *mgm* BGC shows significant similarity to *cmm* BGC (Fig. 2a, Supplementary Table 2 and Supplementary Figs. 2-5), notable differences do present, particularly the genes involved in the biosynthesis of extender units and tailoring steps (Fig. 2a). Specifically, a unique *mgmO* gene, encoding a phenol-type FAD-dependent halogenase^[31]^, is located in the upstream of *mgm* BGC. After integration of *mgmO* gene in *cmm* BGC, chlorinated derivatives (**5**-**8**) were isolated and structurally elucidated. Unexpectedly, the chlorination reaction took place at C4 position of the olefin instead of C19 position of the aromatic ring system, confirming that MgmO catalyzes an unconventional olefin chlorination on cinnamomycin scaffold (Additional descriptions are listed in Supplementary Text, supplementary Figs. 6-34, and Supplementary Tables 3, 6-9). By contrast, the homologous genes of a P450 monooxygenase encoding *cmmA* and a methyltransferase encoding *cmmB* are not present in *mgm* BGC, suggesting distinct tailoring processes. Since the substrate specificity of AT domains determines the structural diversity of side chains, the differences of AT domains in the modules 1, 4, and 5 of *mgm* BGC were compared based on the signature motifs^[2]^ (Supplementary Fig. 2). While MgmD1 is likely to incorporate methylmalonyl-CoA in module 1, MgmD2 might utilize ethylmalonyl-CoA in both modules 4 and 5, indicating the acyl group variations between the skeletons of mangromycin and cinnamomycin (Fig. 2b). According to the biosynthetic logic and experimental evidence on the biosynthesis of cinnamomycin (Fig. 2c), we translated the genetic information of *mgm* BGC into possible structures of mangromycin **A**-**C** (Fig. 2d) along with a proposed biosynthetic pathway (Supplementary Fig. 35).

**Fig 2.**
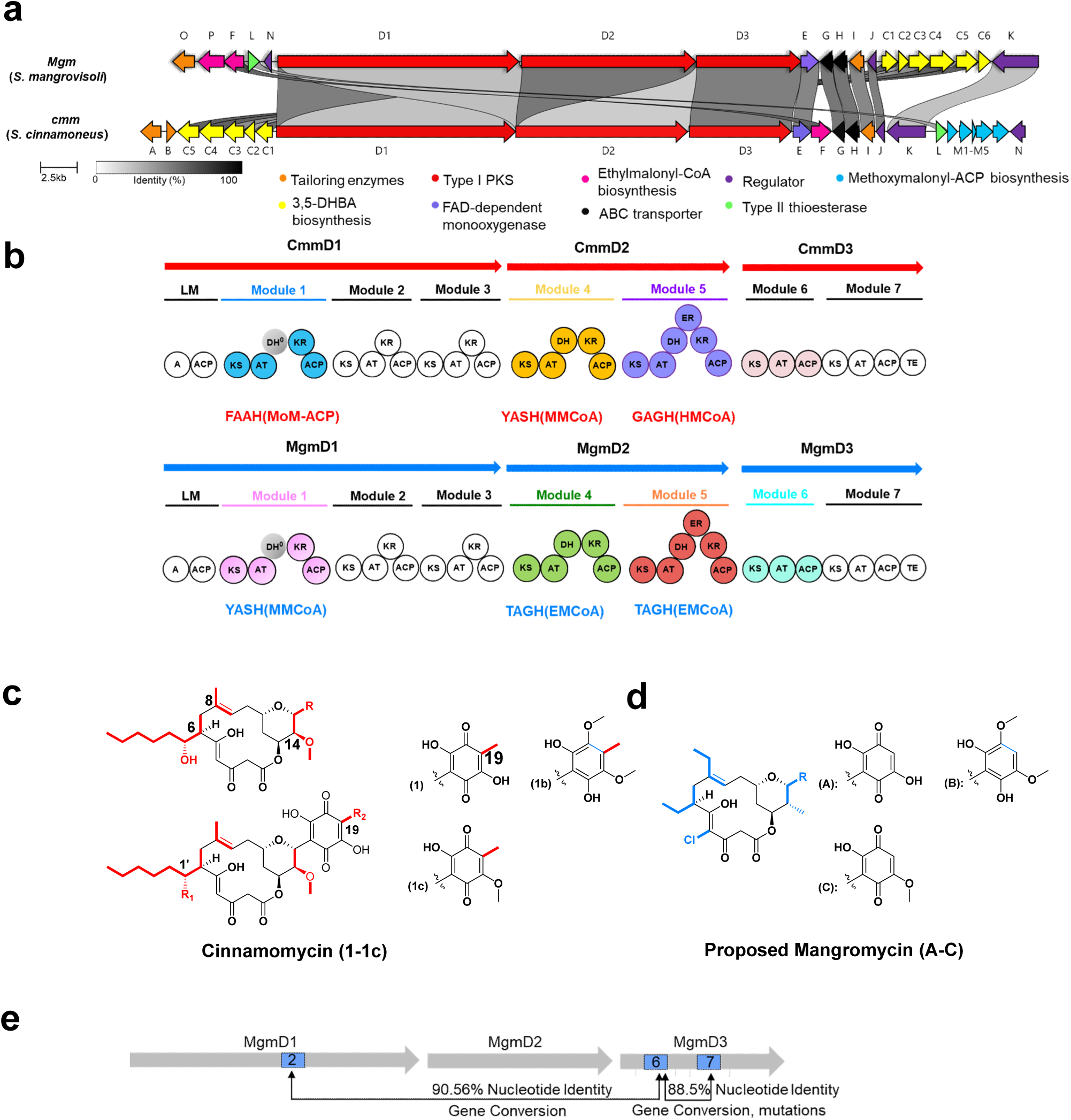
Comparisons of cinnamomycin and its BGC with *mgm* BGC and prediction of mangromycin structure. a) Clinker analysis of *mgm* BGC and *cmm* BGC. Each ORF is labeled by a capital letter. b) Domain organizations of *cmm* and *mgm* modular PKSs. Each circle represents a catalytic domain with its characteristic amino acid motif. The circles with the same color are originated from the same module. LM: 3,5-DHBA-specific loading module; DH^0^: non-functional dehydratase domain verified in previous work^26^. The signature motif “LAF” of CmmD1-DH^0^ is present in MgmD1-DH^0^ (Supplementary Fig. 4). Signature motifs for acyltransferase specificity with the corresponding extender unit are labeled under each module. OMM-ACP: methoxymalonyl-ACP; MMCoA: methylmalonyl-CoA; HMCoA: hexylmalonyl-CoA; EMCoA: ethylmalonyl-CoA. c) Structures of cinnamomycin (**1, 1b, 1c** and **2**-**4**) elucidated in previous work^[26]^. d) Proposed structures of mangromycin **A**-**C** based on the biosynthetic logic and bioinformatics analysis. e) The locations of gene conversion regions in *mgm* BGC. Rectangles colored in blue represent gene conversion fragments. Numbers in rectangles are the abbreviation of module number.

Next, we comparatively analyzed the AT_conversion_ regions of modules 2, 6, and 7 within *mgm* BGC. A profound level of nucleotide identity (≥88%) was found in these regions, confirming the occurrence of gene conversion in *mgm* BGC. Differently, there are no identical gene fragments, which could be attributed to other later evolutionary events (Fig. 2e). Meanwhile, the phylogenetic tree analysis of KS domains indicated that most KS domains in *cmm* and *mgm* BGCs are clustered at the same clade (Supplementary Fig. 36). In particular, CmmD2-KS_5_ and MgmD2-KS_5_ are in a single node, confirming that they might be derived from the same ancestral sequence.

Because AT domains control the specificity of extender units, we therefore focused on AT_conversion_ for structural diversification through the evolutionary event-based PKS engineering in this study, aiming to address the compatibility issues by the exchanges of catalytic elements between *cmm* and *mgm* BGCs.

### Gene conversion-guided successive engineering to create mangromycin skeleton

To reprogram *cmm* BGC for producing predicted mangromycin, various extender units incorporated by modules 1, 4, and 5 in *cmm* BGC were required to be individually altered for matching those in the corresponding regions of *mgm* BGC (Fig. 2b). To mimic the AT_conversion_ processes in nature, we proposed following guidelines for AT engineering: i) The DNA fragments corresponding to AT_conversion_, spanning the amino acid motifs from “GTNAH” to “LPTY” in each module, are designated as AT_c_ region to locate the boundaries; ii) Prioritize the use of catalytic elements from the same BGC; iii) Exogenous elements with higher sequence homology to host BGCs are beneficial for functional compatibility.

Following the guidelines i) and ii), we utilized the AT_c_ region from CmmD2-module 4, specific to methylmalonyl-CoA, to replace the corresponding region in CmmD1-module1, creating mutant S1 (Supplementary Table 5 and Supplementary Fig. 37), aiming to substitute the methoxy group by a methyl group at C14 (Fig. 3a). In accordance with guideline iii), the MgmD2-AT_5-conversion_ region showed higher homology to *cmm* BGC than MgmD2-AT_4-conversion_ region. Thus, this region was chosen for the exchange with the corresponding region in module 4 of CmmD2 to generate mutant S2 (Supplementary Table 5 and Supplementary Fig. 38) or module 5 to produce mutant S3 (Supplementary Table 5 and Supplementary Fig. 39).

**Fig. 3.**
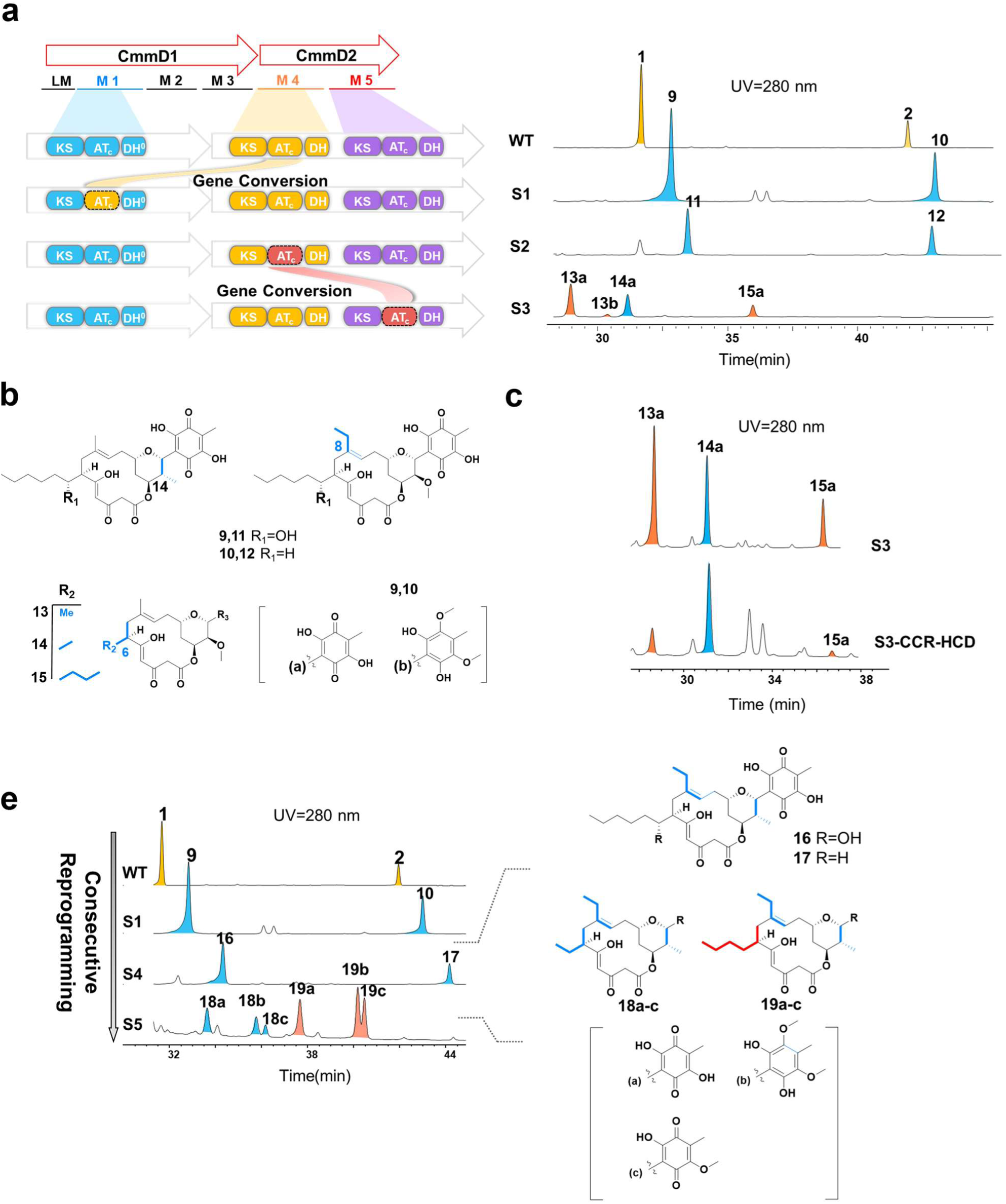
The generation of mangromycin scaffold by successive AT_conversion_ replacements. a) Diagram for AT_conversion_ replacements in wild-type *S. cinnamoneus* (left) and HPLC analysis of different strains (right). Abbreviations represent different catalytic domains. AT_c_ colored in yellow represent CmmD2-AT_4-conversion_; AT_c_ colored in red represent MgmD2-AT_5-conversion_. b) Structures of compounds **9**-**15a**. c) HPLC analysis of strains of S3 and S3-CCR-HCD. d) HPLC analysis of strains of S4 and S5 (left) and the structures of the analogues (right).

After the fermentation of mutant strains S1, S2, and S3, HPLC analyses (Fig. 3a) indicated the production of a series of new peaks (**9**-**15a**) to display identical UV-Vis spectra to cinnamomycin **1** (Supplementary Fig. 40), along with the disappearance of cinnamomycin **1** and **2** in all mutants. Then, fermentation was scaled up for isolation and structural characterization. LC/MS and 1D and 2D NMR analyses (Supplementary Figs. 41-96 and Supplementary Tables 10-17) confirmed the structures of compounds **9**-**15a** (Fig. 3b). Compounds **9**-**12** from mutants S1 and S2 were in line with the alkyl group substitutions at C14 and C8 positions. Compared to **9** and **11**, compounds **10** and **12** lack a C1’ hydroxyl group, due to lower catalytic activity of CmmA^[29]^. Notably, the yields of compounds **9**-**12** were not sacrificed, which was close to that of cinnamomycin **1** and **2** in wild-type strain (∼50 mg/L).

Despite the utilization of Mgm AT_5-conversion_ region in both S2 and S3 strains, they surprisingly behaved significantly different (Fig. 4a). In S2 strain, Mgm AT_5-conversion_ region in module 4 resulted in ethyl substitution at C8 with high specificity. By contrast, S3 strain generated a group of structurally diverse cinnamomycin analogues **13a-15a**, varying in acyl groups at C6 position, whereas the desired product of **14a** was much less than expected. To enhance the biosynthesis of ethylmalonyl-CoA for improving the titer of **14a**, the genes of crotonyl-CoA carboxylase (CCR) and 3-hydroxybutyryl-CoA dehydrogenase (HCD) from *Streptomyces coelicolor* A3(2) (GCA_008931305.1), critical in the ethylmalonyl-CoA biosynthesis pathway^[32]^, were inserted into pSET152, resulting in a construct of pSET152-CCR-HCD^[33]^ (Supplementary Fig. 97). Subsequently, CCR and HCD was co-expressed in mutant S3, yielding S3-CCR-HCD mutant strain. Unexpectedly, although **14a** became the major product in the fermentation broth of the S3-CCR-HCD strain (Fig. 3c), its titer remained comparable to that of S3 strain (∼3 mg/L).

**Fig. 4.**
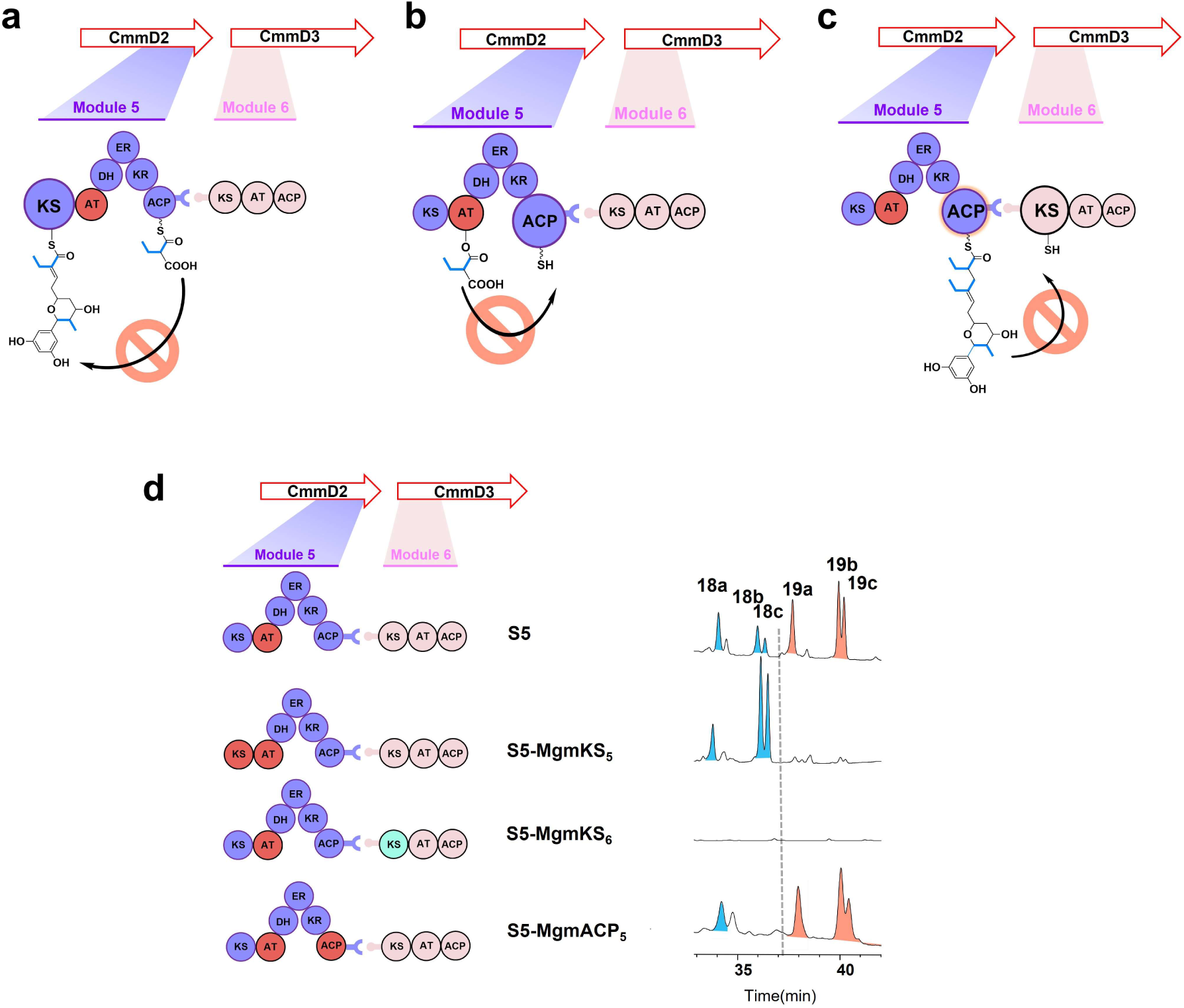
The creation of domain swap mutants for altering the specificity for extender units. a-c) Potential routes and domain organizations for different extender units by domain swapping in S5 strain. Red circles represent exogenous MgmD2-_AT5-conversion_, and purple and pink circles are natural domains in *cmm* BGC. ACP with red circle and KS with cyan circle indicate the counterpart domains from *mgm* BGC. d) HPLC chromatograms of engineered strains at λ = 280 nm. The structures of **18a-c** and **19a-c** are shown in Fig. 3d along with their positions in HPLC analyses.

To further examine the feasibility of using AT_conversion_ for the creation of mangromycin skeleton, CmmD2-AT_4-conversion_ in S1 strain was replaced by MgmD2-AT_5-conversion_ to construct S4 strain. As expected, the S4 strain produced two new compounds **16** and **17** at satisfactory titers (∼15 mg/L). The structures of **16** and **17** were fully elucidated (Supplementary Figs. 98-111 and Supplementary Tables 18-10). Indeed, the substitutions at C8 and C14 took place to generate anticipated alkyl chains (Fig. 3d).

When CmmD2-AT_5_ in S4 strain was substituted by MgmD2-AT_5_-_conversion,_ S5 strain was generated. However, different from S4 strain, HPLC-HRMS profiling of the fermentation broth of S5 strain showed six new peaks (Fig. 3d). Following the scale-up fermentation of S5 strain and isolation, the structures of compounds **18b-c** and **19a-c** were characterized (Supplementary Figs. 112-143 and Supplementary Tables 20-24). Compounds **19a-c** contain a butyl chain attached at C6 position, whereas compounds **18b-c** feature desired ethyl substitution. Such a diversification of extender units in module 5 was similar to that in strain S3, indicating that AT domains are not the only elements for selective incorporation of extender units.

### Identification of the proofreading role of the intra-module KS domains for extender units

To increase the production of **18a-c**, we sought to elucidate the underlying mechanism that discriminates different acyl groups. Base on previous observation on the substrate specificity of the AT domain in module 5, we speculated that the situation in S5 strain could be caused by unnatural domain-domain interactions from exogenous MgmD2-AT_5-conversion_.

Theoretically, there could be three routes to decide extender unit specificity during chain extension (Fig. 4). First, MgmD2-KS_5_, as an intra-module KS, might function as a proof-reading element during the Claisen-condensation step^[34]^. Second, CmmD3-KS_6_ might serve as a proofreader in the transfer of growing intermediates^[35–37]^. Third, the replacement of MgmD2-AT_5-conversion_ might result in unnatural AT_5_-ACP_5_ interactions^[38]^, affecting the fidelity of extender unit incorporation.

To examine the actual route, we employed PKS domain counterparts from *mgm* BGC to replace internal elements for eliminating functional interferences from non-native domain-domain interactions. Additionally, through structural modeling by AlphaFold 2.0, the boundaries for the replacement of KS or ACP domains were precisely defined to ensure their compatibilities (Supplementary Figs. 144-146). Specifically, CmmD2-KS_5_ was replace by its counterpart MgmD2-KS_5_ from *mgm* BGC in S5 strain, resulting in S5-MgmKS_5_ strain (Supplementary Figs. 147). Furthermore, S5-MgmKS_6_ (Supplementary Fig. 148) and S5-MgmACP_5_ (Supplementary Fig. 149) strains were similarly generated (Fig. 5). After fermentation of these newly created mutant strains, HPLC analyses revealed that S5-MgmKS_6_ strain only produced trace amounts of **18a-c** and **19a-c** (Fig. 4), suggesting low efficiency of the assembly line. On the other hand, S5-MgmACP_5_ strain could produce products equivalent to S5 strain with an unchanged ratio of **18a-c** and **19a-c**, implying that the interaction between non-cognate MgmD2-AT_5_ and CmmD2-ACP_5_ did not affect extender unit incorporation (Fig. 4). Intriguingly, unlike the cases in S5-MgmKS_6_ and S5-MgmACP_5_, the production of desired macrolides **18a-c** was remarkably increased in the extracts of S5-MgmKS_5_ strain, along with complete abolishment of **19a-c** (Fig. 4), suggesting the involvement of MgmKS_5_ in the selection of extender units.

**Fig. 5.**
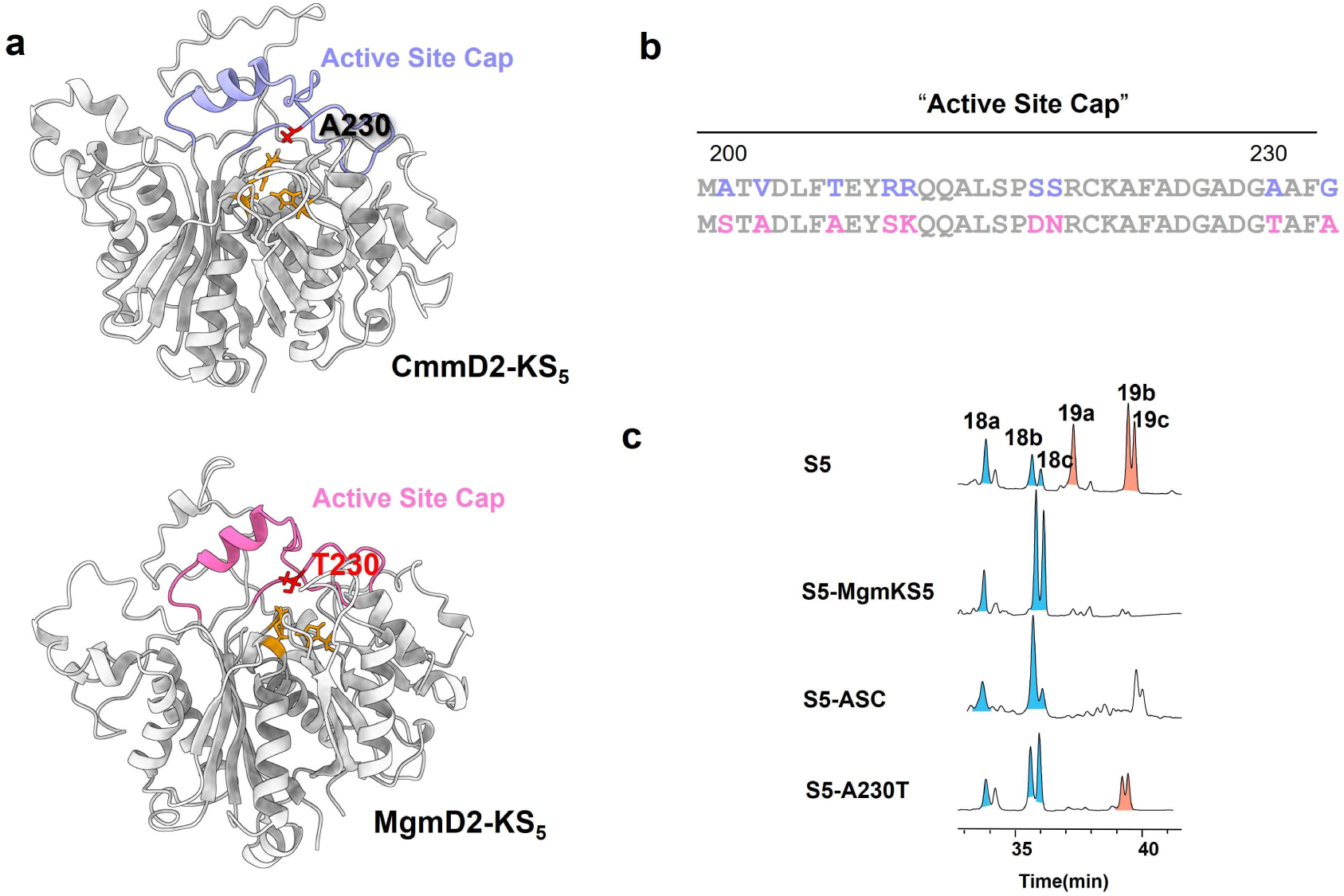
“Active Site Cap” region of KS domain contribute to the extender unit specificity a) Structural models of CmmD2-KS_5_ and MgmD2-KS_5_ created by using AlphaFold 2.0. The catalytic essential residues are colored in yellow with the indication of residues 230. b) Sequence comparison of “Active Site Cap” region of CmmD2-KS_5_ and MgmD2-KS_5_. c) HPLC chromatograms of engineered strains with the same treatments and amounts of fermentation broth at λ = 280 nm with three independent repeats.

CmmD2-KS_5_ and MgmD2-KS_5_ shares 78.64% amino acid sequence identity (Supplementary Fig. 150), but they possessed different specificity on extender units. This difference provided us an opportunity to identify the molecular basis in KS domains to determine the specificity. Multiple sequence alignments and structural modeling of these proteins indicated the presence of a specific region located near catalytically essential residues, which was previously named as "Active-Site Cap" ^[39,40]^ (Fig. 5). Based on the structure and location, we then speculated that the "Active-Site Cap" might be closely related to the specificity of KS domain towards the extender unit. To test this hypothesis, a S5-ASC mutant strain was constructed (Supplementary Fig. 151), in which the "Active-Site Cap" of CmmD2-KS_5_ was modified to match that of MgmD2-KS_5_. Similar to the findings in S3-mgmKS_5_ strain, the production of **19a-c** was completely abolished along with the accumulation of **18a-c** in S5-ASC strain (Fig. 5d), confirming that the specificity of KS domain on extender units is associated with “Active Site Cap”.

To pinpoint the possible determinates for the specificity, amino acid residue 230 in KS domain, the position closest to catalytically essential residues within “Active Site Cap”, was identified to show difference between CmmD2-KS_5_ and MgmD2-KS_5_ (Fig. 5b). To demonstrate functional role of residue 230, site-directed mutagenesis was undertaken to generate a mutant strain S3-A230T, where Ala was substituted by Thr in CmmD2-KS_5_ domain (Supplementary Fig. 152). Consequently, a significant decrease of **19a-c** in the fermentation broth of S5-A230T strain (Fig. 5d) provided additional evidence on the contribution of “Active Site Cap” of KS domain to the process of selecting a proper extender unit.

### Rejuvenation of mangromycin C by morphing *cmm* BGC

With the establishment of macrolide skeleton (**18a-c**), complete biosynthesis of mangromycin required further engineering of tailoring steps. Since CmmA was unable to directly hydroxylate **18a-c**, the first step was to disrupt *cmmB* in S5-MgmKS_5_ strain to yield S6 strain (Supplementary Fig. 153), thereby preventing the methylation at C19 (Fig. 6). Upon examining the fermentation broth of S6 strain, compound **20** was produced, but its yield was drastically decreased compared to **18a-c**. Consequently, sufficient amount of compound **20** was unable to obtain from 50 L fermentation, and its structure therefore was proposed by detailed HRMS/MS analysis (Supplementary Figs. 154 and 155).

**Fig. 6.**
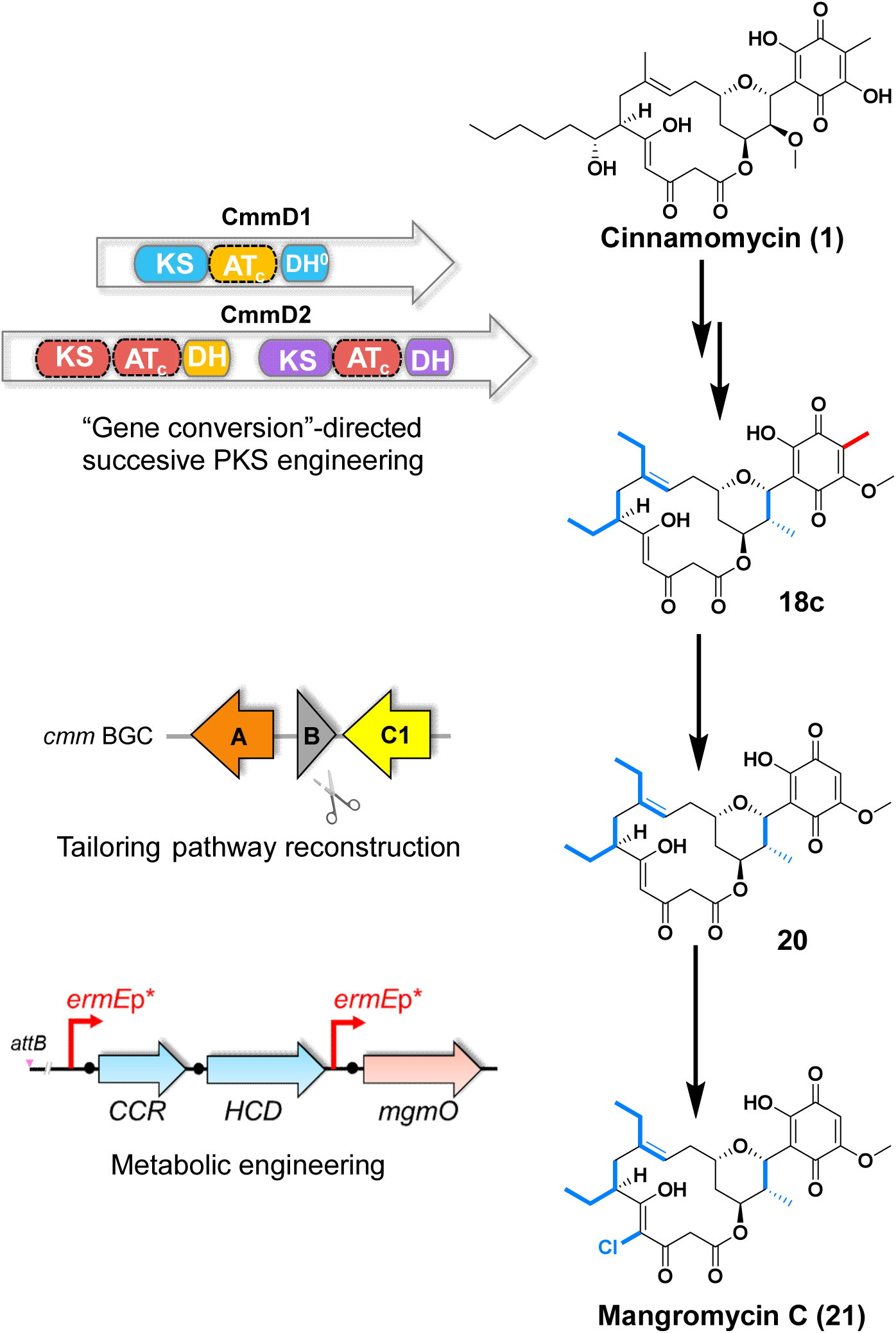
Establishing the biosynthetic route of mangromycin C (**21**) by *de novo* engineering in *Streptomyces cinnamoneus*. Proposed biosynthetic route of mangromycin **C** generated using *S. cinnamoneus* as a template. The grey triangle labeled **B** represents the in-frame deletion of *cmmB* gene in S5-mgmKS_5_ strain. The arrows labeled in **A** and **C1** represent the open reading frames of *cmmA* and *cmmC1*.

According to the characteristics of flavin-dependent halogenase MgmO, C4-chlorination would be the final step to complete the biosynthesis of mangromycin. Given the extremely low abundance of compound **20** and chemical inversion upon the absence of methyl group, CCR-HCD dual gene fragments, was then inserted, under the control of independent *ermE*p* promoter, into pSET152-mgmO plasmid to yield S7 strain (Supplementary Fig. 156). The fermentation of S7 strain led to the production of compound **21** (Supplementary Figs. 154). Following a 50 L fermentation, 1.5 mg of compound **21** was isolated for structural elucidation (Supplementary Figs. 157-162 and Supplementary Table 25), confirming its structure as predicted mangromycin **C** (Fig. 2d and Fig. 6). Thus, through successive engineering efforts, a novel compound of mangromycin **C** was obtained by *de novo* construction of the biosynthetic pathway.

## Discussion

Polyketides biosynthesized by PKSs are a class important compounds with therapeutic values^[41]^. Currently, the emergence of drug-resistance and the growing demand for improved draggability have led to the need of structural diversity. The unique biosynthetic logics of modular PKSs provide an opportunity to utilize synthetic biology approach to reprogram PKSs. However, the production of structural analogs by PKS reprogramming is still in the stage of trial-and-correct, due primarily to insufficient knowledge on dynamic conformations of different modules, domain-domain interactions, and speed-limiting bottlenecks in the complex biosynthetic processes.

Inspired by nature’s creation of diverse PKSs and their biosynthetic pathways, evolution-oriented strategies have shown the usefulness and a promising potential in PKS engineering. For instance, mimicking and accelerating natural evolution in laboratory settings for modular PKSs were practiced to yield new families of polyketides with high yields, such as ring-contracted or ring-expanded rapamycins^[25]^. Meanwhile, evolution-related PKS engineering approaches possess the capability of refining the assembly lines in well-organized and coordinated modular PKSs. Additionally, natural evolution contributes to the guidance of optimal selection of splice points during the designs for PKS engineering. Recent reports have extensively demonstrated that the modular unit for insertion or deletion can adhere to a "redefined module"^[19,20]^, spanning from upstream AT to downstream KS, rather than following canonical order from KS to ACP. Therefore, emulating evolutionary processes observed in nature and elucidating their underlying logics could ultimately facilitate the advancements in PKS engineering.

In this study, we found the presence of an evolutionary event of gene conversion in the AT domains of *cmm* BGC. The finding of three identical nucleotide sequences in different AT domains is quite rare, but may reflect how nature creates the diversity for acyl groups. Thus, we named such a unique phenomenon as AT_conversion_, which may offer us a new way for seamless engineering of PKS assembly line. Subsequent application of gene conversion guided genome mining resulted in the identification of a homologous *mgm* BGC, which enabled us to test the usefulness of gene conversion guided strategy for PKS engineering. Through successive engineering of modular PKS in *cmm* BGC, this gene conversion-directed approach was paid off to introduce three desired acyl groups, specifically achieved by replacing exogenous AT_conversion_ regions to reconstruct the scaffold of cinnamomycins in an efficient manner. Notably, the yields of analogues **9** and **10** resulted from S1 mutant strain even surpassed those of cinnamomycin **1** and **2** in wild-type strain. This surprising finding implies that CmmD2-AT_4_ domain might be transferred from ancestral CmmD1-module 1 to CmmD2-module 4 through the process of gene conversion.

Commonly, the quality control system of modular PKSs consists of a set of precise events to ensure irreversible assembly of substrate and current product output. The system includes the dictation of the flow of inter-modular substrate transfer by docking domain pairing^[42]^, the gatekeeping roles of downstream KS domains and the hydrolysis of inaccurate intermediates by type II thioesterases^[43]^. On the other hand, it is also observed that some PKSs, such as the ones producing antimycin or epothilone, display a tolerance to incorporate different extender units, often attributing to the substrate promiscuity of AT domain. Regardless of various outcomes, the precise mechanism on ensuring the fidelity of extender units during Claisen-condensation step is not fully elucidated. In this study, we identified the proofreading role of intra-module KS domains to control the fidelity of extender units, providing a new possibility for the governance of polyketide synthase assembly line and the involvement of KS-AT didomain. Recently, the condensation domain in NRPS biosynthesis was also revealed to be proofreading for the control of substrate- and stereo-specificity^[44]^, suggesting wide occurrence of proofreading for well-coordinated stepwise biosynthesis of natural products. Therefore, because of the pivotal roles in determining substrate specificity and stereochemistry, organizational and functional characteristics of intra-module KS domains may largely decide the outcomes of PKS reprogramming.

With the development of combinational biosynthesis, successive PKS engineering has emerged as a new fashion to achieve double or even triple substitutions to produce "unnatural" natural products^[45]^. However, unpredictable functionality of chimeric assembly line and historically low yield in PKS engineering have continuously hindered efficient PKS engineering. In this study, we take the advantages of high compatibility and product yield exhibited by AT_conversion_ replacements and intra-module KS engineering, a quadruple-substitution chimeric assembly line is created for the generation of a series of cinnamomycin derivatives This also demonstrates the feasibility and versatility of gene conversion-directed successive PKS engineering to expand the scope of structural diversity and chemical space.

In the past decades, how to efficiently activate cryptic BGCs to obtain their products has been one of the major challenges for natural product discovery^[46]^. Despite the availability of vast genomic information and the advancement of genetic manipulation and heterologous expression, a significant degree of unpredictability still leaves us away from "rational discovery". In addition, although *streptomyces* are well-recognized as a rich source for secondary metabolites with exceptional structural diversity, natural product discovery in *Streptomyces* has become opportunistic due to high rate of redundancy and rediscovery of strains and metabolites. Consequently, engineering of known biosynthetic pathways and reconstruction of BGCs by synthetic biology approach have gained notable attention on the production of novel compounds. Regardless of the significant investments in various engineering technologies, the pace on discovering novel natural products remain relatively slow, particularly working with megaenzyme systems such as modular PKSs because of the process complexity and highly orchestrated assembly line. As one of the approaches, the present successive PKS engineering for *de novo* creation of mangromycin offers a new prospect on the access to natural products biosynthesized by modular megaenzymes from uncultured strains or metagenomic data. Moreover, as exemplified in the consecutive generation of S1-S7 strains guided by gene conversion, the present approach may pave a new way to deal with the issues of compatibility and efficiency.

## Conclusions

In the present study, we have accomplished successive engineering of PKSs for targeted generation of polyketides based on the evolutionary event of gene conversion typically occurred in KS and AT domains. This new approach has led to the creation of an artificial PKS gene cluster to produce a new-to-nature compound of mangromycin C. In addition, as summarized in Supplementary Table 26, a series of novel macrolides with diverse side chains have been generated during the engineering efforts. Moreover, we have demonstrated that the selectivity of extender units within the module is governed by intra-module KS domain, rather than previously proposed by AT domain alone. The present PKS engineering strategy enabled by gene conversion may facilitate the discovery of novel bacterial natural products and provide new insights into the correlation between PKS biosynthetic rationale and evolutionary feature.

## Acknowledgements

This work was supported by the grants of the Project Program of the State Key Laboratory of Natural Medicines, China Pharmaceutical University (SKLNMZZ202201), National Key Research and Development Program of China (No.2018YFA0902000), Key Research and Development Project of Guangdong Province (2022B1111070004), and the “Double First-Class” University Project of China Pharmaceutical University (CPU2022QZ08).

## Competing interests

A Chinese patent application was filed with the number of 2024112777429. YC and WJ are the inventors of the patent, and China Pharmaceutical University owns the patent rights.

## Author contributions

WJ: design, data curation, formal analysis, methodology, and writing–original draft. JT: data curation, formal analysis. BZ: data curation, methodology. XW: resources, supervision. YC: conceptualization, funding acquisition, supervision, and writing–review and editing.

