## Supplemental tables and figures for "Gene Conversion Directed Successive Engineering of Modular Polyketide Synthases"

Supplementary Fig. 36. Phylogenetic tree analysis of KS domains. 63
Supplementary Fig. 37. Construction and confirmation of mutant S1 strain. 64
Supplementary Fig. 38. Construction and confirmation of mutant S2 strain. 65
Supplementary Fig. 39. Construction and confirmation of mutant S3 strain. 66
Supplementary Fig. 40. UV-Vis spectra of compounds **1**-**21**. 67
Supplementary Fig. 41. HR-ESI-MS of derivates **9**. 68
Supplementary Fig. 42. $^{1}$H NMR spectrum (500 MHz, $CDCl_3$) of compound **9**. 68
Supplementary Fig. 43. $^{13}$C NMR spectrum (125 MHz, $CDCl_3$) of compound **9**. 69
Supplementary Fig. 44. HSQC NMR spectrum (500 MHz, $CDCl_3$) of compound **9**. 69
Supplementary Fig. 45. HMBC NMR spectrum (500 MHz, $CDCl_3$) of compound **9**. 70
Supplementary Fig. 46. $^{1}$H-$^{1}$H COSY NMR spectrum (500 MHz, $CDCl_3$) of compound **9**. 70
Supplementary Fig. 47. NOESY NMR spectrum (500 MHz, $CDCl_3$) of compound **9**. 71
Supplementary Fig. 48. HR-ESI-MS spectrum of compound **10**. 71
Supplementary Fig. 49. $^{1}$H NMR spectrum (500 MHz, $CDCl_3$) of compound **10**. 72
Supplementary Fig. 50. $^{13}$C NMR spectrum (125 MHz, $CDCl_3$) of compound **10**. 72
Supplementary Fig. 51. HSQC NMR spectrum (500 MHz, $CDCl_3$) of compound **10**. 73
Supplementary Fig. 52. HMBC NMR spectrum (500 MHz, CDCl3) of compound **10**. 73
Supplementary Fig. 53. $^{1}$H-$^{1}$H COSY NMR spectrum (500 MHz, $CDCl_3$) of compound **10**. 74
Supplementary Fig. 54. NOESY NMR spectrum (500 MHz, $CDCl_3$) of compound **10**. 74
Supplementary Fig. 55. HR-ESI-MS spectrum of compound **11**. 75
Supplementary Fig. 56. $^{1}$H NMR spectrum (600 MHz, $CDCl_3$) of compound **11**. 75
Supplementary Fig. 57. $^{13}$C NMR spectrum (150 MHz, $CDCl_3$) of compound **11**. 76
Supplementary Fig. 58. HSQC NMR spectrum (600 MHz, $CDCl_3$) of compound **11**. 76
Supplementary Fig. 59. HMBC NMR spectrum (600 MHz, $CDCl_3$) of compound **11**. 77
Supplementary Fig. 60. $^{1}$H-$^{1}$H COSY NMR spectrum (600 MHz, $CDCl_3$) of compound **11**. 77
Supplementary Fig. 61. NOESY NMR spectrum (600 MHz, $CDCl_3$) of compound **11**. 78
Supplementary Fig. 62. HR-ESI-MS spectrum of compound **12**. 78
Supplementary Fig. 63. $^{1}$H NMR spectrum (500 MHz, $CDCl_3$) of compound **12**. 79
Supplementary Fig. 64. $^{13}$C NMR spectrum (125 MHz, $CDCl_3$) of compound **12**. 79
Supplementary Fig. 65. HSQC NMR spectrum (500 MHz, $CDCl_3$) of compound **12**. 80
Supplementary Fig. 66. HMBC NMR spectrum (500 MHz, $CDCl_3$) of compound **12**. 80
Supplementary Fig. 67. $^{1}$H-$^{1}$H COSY NMR spectrum (500 MHz, $CDCl_3$) of compound **12**. 81
Supplementary Fig. 68. NOESY NMR spectrum (500 MHz, $CDCl_3$) of compound **12**. 81
Supplementary Fig. 69. HR-ESI-MS spectrum of compound **13a**. 82
Supplementary Fig. 70. $^{1}$H NMR spectrum (600 MHz, $CDCl_3$) of compound **13a**. 82
Supplementary Fig. 71. $^{13}$C NMR spectrum (150 MHz, $CDCl_3$) of compound **13a**. 83
Supplementary Fig. 72. HSQC NMR spectrum (600 MHz, $CDCl_3$) of compound **13a**. 83

### Supplementary Text

#### Characterization of FAD-dependent halogenase *mgmO* to catalyze an unexpected olefin chlorination

To characterize the flavin-dependent halogenases (FDHs) are recognized for their capability to incorporate halogens into natural products, which are further categorized into subclasses based on their substrate-specificity and sequence homology. A phylogenetic comparison of MgmO in *mgm* BGC with known FDHs indicated that MgmO falls in the phenolic-type FDHs, known to chlorinate phenolic moieties (Supplementary Fig. 6). Accordingly, it was reasonable to speculate that MgmO catalyzes the chlorination at C19 position of 2,5-dihydroxy-*p*-benzoquinone moiety in cinnamomycins (Supplementary Fig. 8a), the site also to be methylated by CmmB. To examine the function of MgmO, the gene of *mgmO* was synthesized (Supplementary Table 3) and incorporated into pSET152 under the control of constitutive *ermE**p promoter (Supplementary Table 3 and Supplementary Fig. 7). To eliminate potential interference from CmmB, pSET152-mgmO construct was introduced into wild-type *S. cinnamoneus* and*ΔcmmB* mutant, resulting in the strains of WT-mgmO and ΔcmmB-mgmO for heterologous expression of *mgmO*. After fermentation of these strains, HPLC analyses showed the appearance of four new peaks (Supplementary Fig. 8b), which were confirmed as chlorinated derivatives of cinnamomycin **1**-**4** based on their MS fragmentations (Supplementary Figs. 9 and 10). Surprisingly, the chlorination occurred at C4 position of the double bond in cinnamomycins after structural elucidation of compounds **5-8** from large-scale fermentation, purification and NMR spectroscopy (Supplementary Figs. 11-34 and Supplementary Tables 5-8). The chlorination at the same position by the strains of WT-mgmO and ΔcmmB-mgmO further ruled out the possibility of competing for C19 position. Thus, the function of MgmO was characterized as a FAD-dependent halogenase to chlorinate the olefin group of cinnamomycin-type macrolides (Supplementary Fig. 8a).

### Isolation and purification of cinnamomycin analogues **5**-**21**

(1) For isolation of compounds **5** and **6** from strain WT+mgmO (Supplementary Table 25), a total of 5 L of fermentation media were extracted with an equal volume of ethyl acetate. The extracts were evaporated and dissolved in ethyl acetate. Then, the crude extracts were subjected to C18 silica gel column chromatography, and eluted stepwise using an acetonitrile/water gradient from 10% acetonitrile to 100% acetonitrile. The fractions containing the target compounds were confirmed by HPLC analyses, and the same fractions were combined. Finally, 80 mg of **5** and 60 mg of **6** was obtained.

compound **5**: yellow powder. NMR data, see Supplementary Table 5.

compound **6**: yellow powder. NMR data, see Supplementary Table 6.

(2) For isolation of compounds **7** and **8** from strain ΔcmmB+mgmO (Supplementary Table 25), a total of 5 L of fermentation media were extracted with an equal volume of ethyl acetate. The extracts were evaporated and dissolved in ethyl acetate. Then, the crude extracts were subjected to C18 silica gel column chromatography, and eluted stepwise using an acetonitrile/water gradient from 10% acetonitrile to 100% acetonitrile. The fractions containing the target compounds were confirmed by HPLC analyses, and the same fractions were combined. Finally, 70 mg of **7** and 50 mg of **8** was obtained.

compound **7**: yellow powder. NMR data, see Supplementary Table 7.

compound **8**: yellow powder. NMR data, see Supplementary Table 8.

(3) For isolation of compounds **9** and **10** from strain S1 (Supplementary Table 25), a total of 3 L of fermentation media were extracted with an equal volume of ethyl acetate. The extracts were evaporated and dissolved in ethyl acetate. Then, the crude extracts were subjected to C18 silica gel column chromatography, and eluted stepwise using an acetonitrile/water gradient from 10% acetonitrile to 100% acetonitrile. The fractions containing the target compounds were confirmed by HPLC analyses, and the same fractions were combined. Finally, 180 mg of **9** and 60 mg of **10** was obtained.

compound **9**: yellow powder. NMR data, see Supplementary Table 9.

compound **10**: yellow powder. NMR data, see Supplementary Table 10.

(4) For isolation of compounds **11** and **12** from strain S2 (Supplementary Table 25), a total of 3 L of fermentation media were extracted with an equal volume of ethyl acetate. The extracts were evaporated and dissolved in ethyl acetate. Then, the crude extracts were subjected to C18 silica gel column chromatography, and eluted stepwise using an acetonitrile/water gradient from 10% acetonitrile to 100% acetonitrile. The fractions containing the target compounds were confirmed by HPLC analyses, and the same fractions were combined. Finally, 75 mg of **11** and 35 mg of **12** was obtained.

compound **11**: yellow powder. NMR data, see Supplementary Table 11.

compound **12**: yellow powder. NMR data, see Supplementary Table 12.

(5) For isolation of compounds **13a**, **13b**, **14a** and **15a** from strain S3 (Supplementary Table 25), a total of 15 L of fermentation media were extracted with an equal volume of ethyl acetate. The extracts were evaporated and dissolved in ethyl acetate. Then, the crude extracts were subjected to C18 silica gel column chromatography, and eluted stepwise using an acetonitrile/water gradient from 10%

acetonitrile to 100% acetonitrile. The fractions containing the target compounds were confirmed by HPLC analyses, and the same fractions were combined. Fractions containing the desired compounds were further purified using semi-preparative HPLC on a YMC-Pack ODS-A column with a water/acetonitrile gradient (35:65) over 25 minutes at a flow rate of 1.0 mL/min monitored at 280 nm. Finally, 75 mg of **13a**, 7.5 mg of **13b,** 30mg of **14a** and 20mg of **15a** was obtained.

compound **13a**: yellow powder. NMR data, see Supplementary Table 13.

compound **13b**: white powder. NMR data, see Supplementary Table 14.

compound **14a**: yellow powder. NMR data, see Supplementary Table 15.

compound **15a**: yellow powder. NMR data, see Supplementary Table 16.

(6) For isolation of compounds **16** and **17** from strain S4 (Supplementary Table 25), a total of 3 L of fermentation media were extracted with an equal volume of ethyl acetate. The extracts were evaporated and dissolved in ethyl acetate. Then, the crude extracts were subjected to C18 silica gel column chromatography, and eluted stepwise using an acetonitrile/water gradient from 10% acetonitrile to 100% acetonitrile. The fractions containing the target compounds were confirmed by HPLC analyses, and the same fractions were combined. Finally, 75 mg of **16** and 30 mg of **17** was obtained.

compound **16**: yellow powder. NMR data, see Supplementary Table 17.

compound **17**: yellow powder. NMR data, see Supplementary Table 18.

(7) For isolation of compounds **18b and 18c** from strain S5-mgmKS$_5$ (Supplementary Table 25), a total of 10 L of fermentation media were extracted with an equal volume of ethyl acetate. The extracts were evaporated and dissolved in ethyl acetate. Then, the crude extracts were subjected to C18 silica gel column chromatography, and eluted stepwise using an acetonitrile/water gradient from 10% acetonitrile to 100% acetonitrile. The fractions containing the target compounds were confirmed by HPLC analyses, and the same fractions were combined. Fractions containing the desired compounds were further purified using semi-preparative HPLC on a YMC-Pack ODS-A column with a water/acetonitrile gradient (35:65) over 25 minutes at a flow rate of 1.0 mL/min monitored at 280 nm. Compound **18a** degrades during purification process. Finally, 35mg of **18b** and 30mg of **15a** was obtained.

compound **18b**: white powder. NMR data, see Supplementary Table 19.

compound **18c**: yellow powder. NMR data, see Supplementary Table 20.

(8) For isolation of compounds **19a, 19b** and **19c** from strain S5 (Supplementary Table 25), a total of 10 L of fermentation media were extracted with an equal volume of ethyl acetate. The extracts were evaporated and dissolved in ethyl acetate. Then, the crude extracts were subjected to C18 silica gel column chromatography, and eluted stepwise using an acetonitrile/water gradient from 10% acetonitrile to 100% acetonitrile. The fractions containing the target compounds were confirmed by HPLC analyses, and the same fractions were combined. Fractions containing the desired compounds were further purified using semi-preparative HPLC on a YMC-Pack ODS-A column with a water/acetonitrile gradient (15:85) over 25 minutes at a flow rate of 1.0 mL/min monitored at 280 nm. Finally, 25mg of **19a**, 20mg of **19b** and 25mg of **19c** was obtained.

compound **19a**: yellow powder. NMR data, see Supplementary Table 21.

compound **19b**: white powder. NMR data, see Supplementary Table 22.

compound **19c**: yellow powder. NMR data, see Supplementary Table 23.

(9) For isolation of compound **21** from strain S7 (Supplementary Table 25), a total of 20 L of fermentation media were extracted with an equal volume of ethyl acetate. The extracts were evaporated and dissolved in ethyl acetate. Then, the crude extracts were subjected to C18 silica gel column chromatography, and eluted stepwise using an acetonitrile/water gradient from 10% acetonitrile to 100% acetonitrile. Compound **21** easily disperses on the column, so that we increased the column pressure during the purification process. The fractions containing the **21** were confirmed by HPLC analyses, and the same fractions were combined. Fractions containing the desired compounds were further purified using semi-preparative HPLC on a YMC-Pack ODS-A column with a water/acetonitrile gradient (35:65) over 25 minutes at a flow rate of 1.0 mL/min monitored at 280 nm. Finally, 3.5 mg of **21** was obtained.

compound **21**: yellow powder. NMR data, see Supplementary Table 24.

### Supplementary Table

Supplementary Table 1. Nucleotide sequence of $AT_{conversion}$ region in *cmm* BGC

| |
|---|
| $AT_2$ , $AT_6$ region： |
| GGAACCAACGCCCACGTCATCCTGGAACAGGCACCACCAGAGACAGTGGAGGAGCAGGCGCC<br>CGCCACTGGGCAGAGCGACGTGGTCGTGCCCTGGGTCGTCTCGGGCAAGACCGAAGCAGCCG<br>TCGACGAACAGCTCGCACGGCTACGCCAATGGGCCCAGCAACGGCCGGACGCCCGGCCAGTC<br>GACGTCGCCCACGCCTTGGCCACGAGTCGCACGCACTTCGAATACCGGGCCGCGGTCGTAGG<br>CCGAACCCACGAAGAACTTGTCACCGCGCTGGCCTCGCCCGCATCGGTCTTACGAGGCCGGCG<br>ACAAGGCCGCGAACTGGCGGTCCTCTTCGCGGGACAGGGCTCACAACGCCCCGGCATGGGAC<br>GCGAACTCCACGCCACCTACCCAGCCTTCGCCGACGCCTTCGACGCCATCCGCACCGAACTCG<br>ACCAACACCTAGACCTGCCACTCACCCACATCATGTGGCACGACACCACCGGCCTACTCCACCA<br>AACCGCCTATACCCAGGCCGCGCTCTTCGCCCTCGAAACAGCCCAATACAGGCTCATAGAAAGC<br>TGGGGACTACGGCCAGAGGCACTGCTCGGGCACTCGATCGGGGAACTCACCGCAGCCCACGT<br>CGCCGGCATCTGGTCTCTCGAAGACGCCTGCACCCTGGTCGCCGCACGCGGACGCCTCATGCA<br>AACCCTCCCCACCAGCGGCACCATGACCGCCCTCCAAGCCACAGAAAACGAAATCGCTCCTCTT<br>CTCAACGAGCGAGTGAGCCTGGCAGCCATCAACGGCCCCTCATCCGTCGTCATCTCCGGCGAC<br>AAAGACGCCGTCGACACCATCGCAACAACCGTCACCAACTGGGGCCGCAAAACCAAGAAACTC<br>CACGTCAGCCACGCCTTCCACTCCCCCCACATGGACCCCATACTCGACGAATTCCAAACCATCG<br>CCGAGTCCCTCACCTACCACCCACCCCAGCTGACCATCATCTCCAACCTCACCGGCCAACCCAC<br>CACCACCGACACCCTCACCCCCACCTACTGGACCCACCACATCCGCCAACCCGTCCGCTTCAA<br>CGACGGCCTCACCCACCTCACCACCCACACCCTCCTCGAACTCGGCCCCGACAGCACCCTCAC<br>CGCCCTCACCCAACAAACCCACCCCCACACCACCGCCACCCCCCTCCTCCGCAAAAACCACCC<br>CGAACCCCACACCACCATCACCGCCCTCACCCACCTCCACACCACCGGCACCACCCCCAACTG<br>GAACACCCTCCTCCCCACCACCCACCCCACCCACGACCTCCCCACCTACCCCTTCCAACACCAC<br>CACTACTGGCTG |
| AT7 region： |
| GCACGAACGCGCACCTCATCCTTGAGGAATCACCCCAGGAGACGGCCGACACGGCGGATCCGA<br>CCGCGCCCGTCGAGGCCAAGGACGTCGTCGTGCCGTGGGTCGTCTCGGCGAAGTCACCGGAG<br>GCGCTGGACGAGCAGCTGAGCCGGCTGCGGCGCTGGGCCGAGGACCGTCCCGACGTGCACC<br>CCGCCGACACGGCGTACGCCCTCGCGACCACCCGCACCCACTTCGATCACCGGGCGGCCGTG<br>GTGGGCAGCACCCGCGAAGAACTGATCGAGGCCCTGGGCTCGGTCTCGGTGATCCGCGGCAG<br>GCGGCACGACGGCAGGCTGGCGATCCTCTTCGCGGGACAGGGCTCACAACGGCCCGGCATGG<br>GGCGCGAACTCCACGCCACCTACCCAGCCTTCGCCGACGCCTTCGACGCCATCCGCACCGAAC<br>TCGACCAACACCTAGACCAGCCACTCACCCACATCATGTGGCACGACACCACCGGCCTGCTCCA<br>CCAAACCGCCTACACCCAGGCCGCGCTCTTCGCCCTCGAAACAGCCCAATACAGGCTCATAGAA<br>AGCTGGGGACTACGGCCAGAGGCACTGCTCGGGCACTCGATCGGGGAACTCACCGCAGCCCA<br>CGTCGCCGGCATCTGGTCTCTCGAAGACGCCTGCACCCTGGTCGCCGCACGCGGACGCCTCAT<br>GCAAACCCTCCCCACCAGCGGCACCATGACCGCCCTCCAAGCCACAGAAAACGAAATCGCTCC<br>TCTTCTCAACGAGCGAGTGAGCCTGGCAGCCATCAACGGCCCCTCATCCGTCGTCATCTCCGG<br>CGACAAAGACGCCGTCGACACCATCGCAACAACCGTCACCAACTGGGGCCGCAAAACCAAGAA<br>ACTCCACGTCAGCCACGCCTTCCACTCCCCCCACATGGACCCCATACTCGACGAATTCCAAACC<br>ATCGCCGAGTCCCTCACCTACCACCCACCCCAGCTGACCATCATCTCCAACCTCACCGGCCAAC<br>CCACCACCACCGACACCCTCACCCCCACCTACTGGACCCACCACATCCGCCAACCCGTCCGCT<br>TCAACGACGGCCTCACCCACCTCACCACCCACACCCTCCTCGAACTCGGCCCCGACAGCACCC<br>TCACCGCCCTCACCCAACAAACCCACCCCCACACCACCGCCACCCCCCTCCTCCGCAAAAACC<br>ACCCCGAACCCCACACCACCATCACCGCCCTCACCCACCTCCACACCACCGGCACCACCCCCA<br>ACTGGAACACCCTCCTCCCCACCACCCACCCCACCCACGACCTCCCCACCTACCCCTTCCAACA<br>CCACCACTACTGGCTCGAC |

Supplementary Table 2 Gene annotations of *mgm* BGC.

| Gene | Amino Acids | Proposed Function | Protein Homology | Identity (%) |
|---|---|---|---|---|
| *mgmO* | *493* | Tryptophan-halogenase | TiaM (GenBank: ADU85999.1) | 63.95% |
| *mgmP* | 600 | 3-hydroxybutyryl-coa dehydrogenase | WP_011030948.1 | 47.88% |
| *mgmF* | 450 | Crotonyl-coa carboxylase/reductase | WP_030863150.1 | 87.64% |
| *mgmL* | 258 | Type II thioesterase | CmmL | 50.42% |
| *mgmN* | 251 | Transcriptional regulator | WP_104630222.1 | 93.15% |
| *mgmD1* | 5523 | Type I polyketide synthase | CmmD1 | |
| *mgmD2* | 3992 | Type I polyketide synthase | CmmD2 | |
| *mgmD3* | 2397 | Type I polyketide synthase | CmmD3 | |
| *mgmE* | 402 | FAD-dependent monooxygenase | CmmE | 55.50% |
| *mgmG* | 282 | ABC transporter | CmmG | 62.01% |
| *mgmH* | 315 | ABC transporter | CmmH | 62.03% |
| *mgmI* | 330 | Zinc-binding alcohol dehydrogenase | CmmI | 57.23% |
| *mgmJ* | 192 | Transcriptional regulator | CmmJ | 58.76% |
| *mgmC1* | 356 | Type III polyketide synthase | CmmC1 | 70.11% |
| *mgmC2* | 238 | enoyl-CoA hydratase/isomerase family protein | CmmC2 | 48.68% |
| *mgmC3* | 485 | Enoyl-CoA hydratase/isomerase family protein | CmmC3 | 58.55% |
| *mgmC4* | 546 | Thiamine pyrophosphate-binding protein | CmmC4 | 71.38% |
| *mgmC5* | 486 | Benzaldehyde dehydrogenase | CmmC5 | 69.20% |
| *mgmC6* | 268 | Enoyl-CoA-hydratase | VemD | 79.85% |
| *mgmK* | 1058 | Transcriptional regulator | CmmK | 36.20% |

Supplementary Table 3. Bacterial strains used in this work.

| Strains | Characteristics | Source or References |
|---|---|---|
| ***E. coil* strains** | | |
| *E. coil* DH5α | Host for general cloning | Tiangen biotech |
| *E. coil* ET12567/pUZ8002 | Donor strain used for *E. coli-Streptomyces* conjugation | Ref. 1 |
| ***Streptomyces* strains** | | |
| *S. cinnamoneus* ATCC 21532 (WT) | Wild-type strain for cinnamonycin production | Ref. 2 |
| *S. coelicolor A3(2)* | CCR and HCD gene containing strain | GCA_008931305.1 |
| ΔcmmB mutant | *cmmB* in-frame deletion mutant | Ref. 2 |
| WT+mgmO | *mgmO* gene integrated strain of WT | This work |
| *Δ*cmmB+mgmO | *mgmO* gene integrated strain of ΔcmmB mutant | This work |
| S1 | Cmm-AT1 replacement mutant with Cmm-AT4 in wild-type s*treptomyces cinnamoneus* | This work |
| S2 | Cmm-AT4 replacement mutant with Mgm-AT5 in wild-type s*treptomyces cinnamoneus* | This work |
| S3 | Cmm-AT5 replacement mutant with Mgm-AT5 in wild-type s*treptomyces cinnamoneus* | This work |
| S3-CCR-HCD | Ethylmalonyl-CoA pathway key gene overexpression strain of mutant S3 | This work |
| S4 | Cmm-AT4 replacement mutant with Mgm-AT5 in S3 | This work |
| S5 | Cmm-AT5 replacement mutant with Mgm-AT5 in S4 | This work |
| S5-mgmKS$_5$ | Cmm-KS5 replacement mutant with Mgm-KS5 in S5 | This work |
| S5-mgmKS$_6$ | Cmm-KS6 replacement mutant with Mgm-KS6 in S5 | This work |
| S5-mgmACP$_5$ | Cmm-ACP5 replacement mutant with Mgm-ACP in S5 | This work |
| S6 | *cmmB* in-frame deletion mutant in S5-mgmKS5 | This work |
| S7 | Mangromycin **C** produced strain | This work |
| S5-ASC | Mutants with “active site cap” region substitutions in Cmm-KS5 | This work |
| S5-A230T | Cmm-KS5 A230T substituted mutant | This work |

Supplementary Table 4. Plasmids and primers used to construct of exogenous gene integration mutants in this study.

| Mutant Strain | Plasmid | Cloning Method | Fragment | Primer Name | Primer Sequence (5'→3') | Template |
|---|---|---|---|---|---|---|
| WT+mgmO<br>△cmmB+mgmO | pSET152-mgmO | DNA ligation | mgmO-2 | mgmO-P1 | GGAATTCCATATGACGCGGAAAGTCCTAG (*Nde*I) | Gene fragment synthesized by GenScript Biotechnology Co., Ltd |
| | | | | mgmO-P2 | CCGGATATCTCAGGCGACGACGGCGTCGGC (*Eco*RV) | |
| | | | linearized pSET152- *ermE**p digested with *Nde*I/*Eco*RV | | | |
| S3-CCR-HCD | pSET152-CCR-HCD | Gibson assembly | CCR | CCR-HCD-P1 | ACTCCACAGGAGGACCCATATGACCGTGAAGGACATCCTGG | genomic DNA of *S. coelicolor* A3(2) |
| | | | | CCR-HCD-P2 | GGACTCCAGGAATGAGGTCAGATGTTCCGGAAGCGGTTGATG | |
| | | | HCD | CCR-HCD-P3 | CCTCATTCCTGGAGTCCCGCGATGGCCACTCCCCTGTCCGAC | |
| | | | | CCR-HCD-P4 | TTGGGCTGCAGGTCGACTCTAGATTATCAGCGGCGGGCATGCTCG | |
| | | | linearized pSET152-*ermE**p digested with *Nde*I/*Xba*I | | | |
| S7 | pSET152-CCR-HCD-mgmO | Gibson assembly | mgmO-1 | CCR-HCD-mgmO-P1 | CATGCCCGCCGCTGATAATCGAGTGTCCGTTCGAGTGGC | pSET152-mgmO |
| | | | | CCR-HCD-mgmO-P2 | CTGCAGGTCGACTCTAGGCGCTGCAGGTCCGCGGATC | |
| | | | linearized pSET152-CCR-HCD digested with *Xba*I | | | |

Supplementary Table 5. Plasmids and primers used to construct genetic manipulating mutants in this study.

| Mutant Strain | Plasmid | Cloning Method | Fragment | Primer Name | Primer Sequence (5'→3') | Template |
|---|---|---|---|---|---|---|
| S1 | pKC1139-S1 | Gibson assembly | S1-1 | S1-P1 | **AGCTATGACATGATTAC**GAATTCGAGTTCGGCTACCTGAGCATCAC | genomic DNA of *S. cinnamoneus* ATCC 21532 |
| | | | | S1-P2 | **TCTTCGAGGATGACGT**GGGCGTTGGTGCCGCTGAC | |
| | | | S1-2 | S1-P3 | **ACGTCATCCTCGA**AGAGGCAC | |
| | | | | S1-P4 | **TGCTCGAACGGGTACG**TCGGCAGGTCCACCC | |
| | | | S1-3 | S1-P5 | **ACGTACCCGTTCGAGCA**CCGGCACTAC | |
| | | | | S1-P6 | **TAAAACGACGGCCAGTGCC**AAGCTTGACGAGAACATCAAGAACGCGGTG | |
| | | | linearized pKC1139 digested with *EcoR*I/*Hind*III | | | |
| S2<br>S4 | pKC1139-S2 | Gibson assembly | SAT4-1 | S2-P1 | **CAGCTATGACATGATTAC**GAATTCATGGCCACCAGTGAGGCGGTG | genomic DNA of *S. cinnamoneus* ATCC 21532 |
| | | | | S2-P2 | **TTCGAGGATGACGTGG**GCGTTG | |
| | | | SAT4-2 | S2-P3 | **CCACGTCATCCTCGAA**CAGGCCCCCGC | Gene fragment synthesized by GenScript Biotechnology Co., Ltd |
| | | | | S2-P4 | **CTCGAAGGCGTACGT**CGGCAGCTCCACCGTCCGACCCGAGC | |
| | | | SAT4-3 | S2-P5 | **ACGTACGCCTTCGAG**CGGC | genomic DNA of *S. cinnamoneus* ATCC 21532 |
| | | | | S2-P6 | **TAAAACGACGGCCAGTGCC**AAGCTTTCCTCCCTGCCGAGCGTCAG | |
| | | | linearized pKC1139 digested with *EcoR*I/*Hind*III | | | |
| S3<br>S5 | pKC1139-S3 | Gibson assembly | SAT5-1 | S3-P1 | **CAGCTATGACATGATTAC**GAATTCTGCTCGACGACGGCGTCCTC | genomic DNA of *S. cinnamoneus* ATCC 21532 |
| | | | | S3-P2 | **CCTGTTCGAGGATCAC**GTGGGCGTTGGTGCCGCTG | |
| | | | SAT5-2 | S3-P3 | **GTGATCCTCGAACAGG**CCCC | Gene fragment synthesized |
| | | | | S3-P4 | **TACGTCGGCAGCT** | |

| | | | | | | |
|---|---|---|---|---|---|---|
| | | | | | **CCA**CCGTCCGACCCGAGC | by GenScript Biotechnology Co., Ltd |
| | | | SAT5-3 | S3-P5 | **TGGAGCTGCCGACGTA**CGCGTTCGAG | genomic DNA of *S. cinnamoneus* ATCC 21532 |
| | | | | S3-P6 | **TAAAACGACGGCCAGTGCC**AAGCTTAGTTGAGGACCACGTCCACTC | |
| | | | linearized pKC1139 digested with *Eco*RI/*Hind*III | | | |
| S5-mgmKS$_5$ | pKC1139-KS5 | Gibson assembly | SKS5-1 | KS5-P1 | **CAGCTATGACATGATTAC**GAATTCGACGAGGAGGAGTGGACCTGTC | genomic DNA of *S. cinnamoneus* ATCC 21532 |
| | | | | KS5-P2 | **ATGGACACGATGAC**GACCGGGTCCCCGTCCGCCGCGCCCGACC | |
| | | | SKS5-2 | KS5-P3 | GACCCG**GTCGTCATCGTGTCC** | Gene fragment synthesized by GenScript Biotechnology Co., Ltd |
| | | | | KS5-P4 | **CTGTTCGAGGATCACGTG**CGCGTTGGTGCCGGAG | |
| | | | SKS5-3 | KS5-P5 | **CACGTGATCCTCGAACAG**GC | pKC1139-S3 |
| | | | | KS5-P6 | **TAAAACGACGGCC**AGTGCCAAGCTTCAGGTGAGCCGGAACAGCGAG | |
| | | | linearized pKC1139 digested with *Eco*RI/*Hind*III | | | |
| S5-mgmKS$_6$ | pKC1139-KS6 | Gibson assembly | SKS6-1 | KS6-P1 | **CAGCTATGACATGATTAC**GAATTCGTCTTCCTGACCGCCTACTACG | genomic DNA of *S. cinnamoneus* ATCC 21532 |
| | | | | KS6-P2 | **ATCGCGATGGGCTCGTG**ACGCCGGCTCTCGACCTCCTGCAGC | |
| | | | SKS6-2 | KS6-P3 | **CACGAGCCCATCGCGAT**CG | Gene fragment synthesized by GenScript Biotechnology Co., Ltd |
| | | | | KS6-P4 | **AGGATGACGTGGGCGTT**GGTGCCGGAGATGCCGAAG | |
| | | | SKS6-3 | KS6-P5 | **AACGCCCACGTCATCCT**GGAAC | genomic DNA of *S. cinnamoneus* ATCC 21532 |
| | | | | KS6-P6 | **TAAAACGACGGCCAGTGCC**AAGCTTTGCATGGCCACCAAGGAGGAC | |
| | | | linearized pKC1139 digested with *Eco*RI/*Hind*III | | | |
| S5-mgmACP$_5$ | pKC1139-ACP5 | Gibson assembly | ACP5-1 | ACP5-P1 | CAGCTATGACATGATTACGAATTCTGGCACTGGCCCAGGAG | genomic DNA of *S. cinnamoneus* |

| | | | | | | |
|---|---|---|---|---|---|---|
| | | | | | TG | ATCC 21532 |
| | | | | ACP5-P2 | AGCCGTTCGGCCAGGGTGCCGCCGCCCTCGG | |
| | | | ACP5-2 | ACP5-P3 | ACCCTGGCCGAACGGCTC | Gene fragment synthesized by GenScript Biotechnology Co., Ltd |
| | | | | ACP5-P4 | TCCTCCGGCACCAGCAGC | |
| | | | ACP5-3 | ACP5-P5 | AGCTGCTGGTGCCGGAGGAGCCGTC | genomic DNA of *S. cinnamoneus* ATCC 21532 |
| | | | | ACP5-P6 | TAAAACGACGGCCAGTGCCAAGCTTGCTGATCCCGAACGACGACAC | |
| | | | linearized pKC1139 digested with *Eco*RI/*Hind*III | | | |
| S6 | pKC1139-cmmB (Ref. 3) | DNA ligation | cmmB-1 | cmmB-P1 | CG**GAATTC**CGATCACCACCGGACTGAAG (*Eco*RI) | genomic DNA of *S. cinnamoneus* ATCC 21532 |
| | | | | cmmB-P2 | GC**TCTAGA**GGATGGCTGCCGCTCATCCAG (*Xba*I) | |
| | | | cmmB-2 | cmmB-P3 | GC**TCTAGA**GATCACGGAGCCTCCAGCG (*Xba*I) | |
| | | | | cmmB-P4 | CCC**AAGCTT**CGGCTCGGCACCATGAAGACGTC (*Hind*III) | |
| | | | linearized pKC1139 digested with *Eco*RI/*Hind*III | | | |
| S5-ASC | pKC1139-ASC | Gibson assembly | ASC-1 | ASC-P1 | **CAGCTATGACATGATTAC**GAATTCGACGAGGAGGAGTGGACCTGTC | genomic DNA of *S. cinnamoneus* ATCC 21532 |
| | | | | ASC-P2 | **GTGGACATGACGGCCA**CCCCGC | |
| | | | ASC-2 | ASC-P3 | **TGGCCGTCATGTCCAC**GGCCGATCTGTTCG | Gene fragment synthesized by GenScript Biotechnology Co., Ltd |
| | | | | ASC-P4 | **GGCGAACGCGGTGCCGTC** | |
| | | | ASC-3 | A230T-P5 | **GACGGCACCGCGTTCGCC**GAGGGGGTGGGCATGGTGC | genomic DNA of *S. cinnamoneus* ATCC 21532 |
| | | | | A230T-P6 | **CTGTTCGAGGATCAC**GTGGGCGTTGGTGCCGCTGAC | |
| | | | ASC-4 | ASC-P7 | CAC**GTGATCCTCGAACAG**GC | pKC1139-S3 |
| | | | | ASC-P8 | **TAAAACGACGGCCAGTGCC**AAGCTTCAGGTGAGCCGGAACAGCGAG | |
| | | | linearized pKC1139 digested with *Eco*RI/*Hind*III | | | |

| | | | | | | |
|---|---|---|---|---|---|---|
| S5-A230T | pKC1139-A230T | Gibson assembly | A230T1 | A230T-P1 | **CAGCTATGACATGATTAC**GAATTCGACGAGGAGGAGTGGACCTGTC | genomic DNA of *S. cinnamoneus* ATCC 21532 |
| | | | | A230T-P2 | **TCCGCAAAGGCCTTGCA**GCG | |
| | | | A230T-2 | A230T-P3 | **TGCAAGGCCTTTGCGGACGGCGCGGA**CGGCACCGCCTTCGGTGAGGG | |
| | | | | A230T-P4 | **CTGTTCGAGGATCAC**GTGGGCGTTGGTGCCGCTGAC | |
| | | | A230T-3 | A230T-P5 | CAC**GTGATCCTCGAACAG**GC | pKC1139-S3 |
| | | | | A230T-P6 | **TAAAACGACGGCCAGTGCC**AAGCTTCAGGTGAGCCGGAACAGCGAG | |
| | | | linearized pKC1139 digested with *EcoR*I/*Hind*III | | | |

Supplementary Table 6. NMR data of compound **5**.

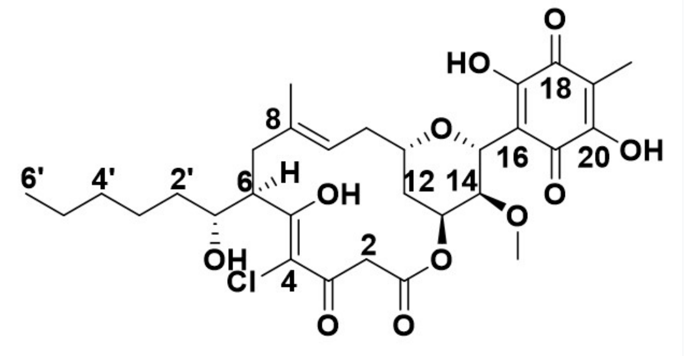

(5)

| NO. | $\delta_H$ (mult., *J* in Hz) | $\delta_C$ | C-type |
|---|---|---|---|
| 1 | | 165.1 | C |
| 2a | 3.55 (d, 1H, *J*=15.7Hz) | 46.5 | $CH_2$ |
| 2b | 3.82 (d, 1H, *J*=15.7Hz) | | |
| 3 | | 188.3 | C |
| 4 | | 110.4 | C |
| 5 | | 188.7 | C |
| 6 | 3.41 (m, 1H) | 47.6 | CH |
| 7a | 2.11 (dd, 1H, *J*=12.9, 3.2Hz) | 39.7 | $CH_2$ |
| 7b | 2.69 (t, 1H, *J*=12.7Hz) | | |
| 8 | | 134.6 | C |
| 9 | 5.19 (t, 1H, *J*=8.4Hz) | 122.6 | CH |
| 10a | 2.16 (dd, 1H, *J*=12.8Hz, 8.0Hz) | 33.7 | $CH_2$ |
| 10b | 2.23 (dt, 1H, *J*=12.7Hz, 10.0Hz) | | |
| 11 | 3.26 (overlapped, 1H) | 75.5 | CH |
| 12a | 1.66 (m, 1H) | 31.2 | $CH_2$ |
| 12b | 1.85 (ddd, 1H, J=16.8, 19.9, 4.2 Hz) | | |
| 13 | 5.39 (q, 1H, *J*=3.0Hz) | 67.1 | CH |
| 14 | 3.25 (overlapped, 1H) | 75.5 | CH |
| 15 | 4.96 (d, 1H, *J*=1.8Hz) | 75.1 | CH |
| 16 | | 111.1 | C |
| 17 | | * | C |
| 18 | | * | C |
| 19 | | 114.3 | C |
| 20 | | * | C |
| 21 | | * | C |
| 8-$CH_3$ | 1.68 (s, 3H) | 16.7 | $CH_3$ |
| 14-$OCH_3$ | 3.40 (s, 3H) | 59.3 | $CH_3$ |
| 19-$CH_3$ | 1.94 (s, 3H) | 7.8 | $CH_3$ |
| 1` | 3.74 (d, 1H, *J*=5.0Hz) | 74.3 | CH |
| 2`a | 1.40 (m, 1H) | 35.8 | $CH_2$ |
| 2`b | 1.58 (m, 1H) | | |
| 3`a | 1.39 (m, 1H) | 25.2 | $CH_2$ |
| 3`b | 1.53 (m, 1H) | | |
| 4` | 1.30 (m, 2H) | 31.7 | $CH_2$ |
| 5` | 1.32 (m, 2H) | 22.6 | $CH_2$ |
| 6’ | 0.90 (t, 3H, *J*=6.9Hz) | 14.0 | $CH_3$ |

$^1$H NMR: 600 MHz; $^{13}$C NMR: 150 MHz (in $CDCl_3$).

* The interconversion of the benzoquinone moiety led to the missing of these carbon signals.

Supplementary Table 7. NMR data of compound **6**.

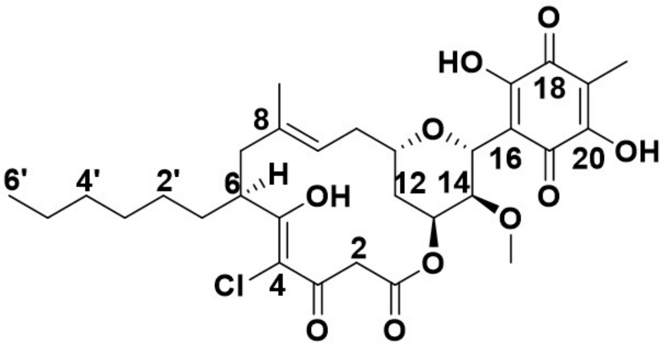

**(6)**

| NO. | δH (mult., J in Hz) | δC | C-type |
|---|---|---|---|
| 1 | | 165.3 | C |
| 2a | 3.54 (d, 1H, J=15.7Hz) | 46.9 | CH2 |
| 2b | 3.82 (d, 1H, J=15.7Hz) | | |
| 3 | | 189.7 | C |
| 4 | | 109.1 | C |
| 5 | | 188.5 | C |
| 6 | 3.29 (m, 1H) | 42.0 | CH |
| 7a | 2.07 (dd, 1H, J=13.0, 3.0Hz) | 42.1 | CH2 |
| 7b | 2.47 (dd, 1H, J=13.5, 6.2Hz) | | |
| 8 | | 135.4 | C |
| 9 | 5.16 (t, 1H, J=8.3Hz) | 122.0 | CH |
| 10a | 2.17 (m, 1H) | 33.7 | CH2 |
| 10b | 2.21 (m, 1H) | | |
| 11 | 3.25 (overlapped, 1H) | 75.5 | CH |
| 12a | 1.67 (m, 1H) | 31.1 | CH2 |
| 12b | 1.83 (ddd, 1H, J=14.0, 11.3, 2.5 Hz) | | |
| 13 | 5.39 (q, 1H, J=3.0Hz) | 67.1 | CH |
| 14 | 3.25 (overlapped, 1H) | 75.5 | CH |
| 15 | 4.95 (d, 1H, J =1.8Hz) | 75.2 | CH |
| 16 | | 111.2 | C |
| 17 | | * | C |
| 18 | | * | C |
| 19 | | 114.1 | C |
| 20 | | * | C |
| 21 | | * | C |
| 8-CH3 | 1.66 (s, 3H) | 16.8 | CH3 |
| 14-OCH3 | 3.40 (s, 3H) | 59.2 | CH3 |
| 19-CH3 | 1.94 (s, 3H) | 7.8 | CH3 |
| 1`a | 1.47 (m, 1H) | 33.9 | CH2 |
| 1`b | 1.67 (m, 1H) | | |
| 2`a | 1.27 (m, 1H) | 27.2 | CH2 |
| 2’b | 1.34 (m, 1H) | | |
| 3` | 1.27 (m, 2H) | 29.3 | CH2 |
| 4` | 1.26 (m, 2H) | 31.6 | CH2 |
| 5` | 1.28 (m, 2H) | 22.6 | CH2 |
| 6` | 0.88 (t, 3H, J=7.0 Hz) | 14.1 | CH3 |

$^{1}H$ NMR: 600 MHz; $^{13}C$ NMR: 150 MHz (in DMSO-d6).

* The interconversion of the benzoquinone moiety led to the missing of these carbon signals.

Supplementary Table 8. NMR data of compound **7**.

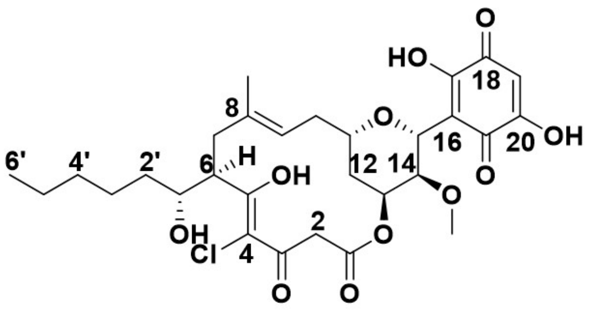

(7)

| NO. | $\delta_H$ (mult., *J* in Hz) | $\delta_C$ | C-type |
|---|---|---|---|
| 1 | | 165.0 | C |
| 2a | 3.55 (d, 1H, *J*=15.7Hz) | 46.7 | $CH_2$ |
| 2b | 3.82 (d, 1H, *J*=15.7Hz) | | |
| 3 | | 188.2 | C |
| 4 | | 110.4 | C |
| 5 | | 188.8 | C |
| 6 | 3.41 (m, 1H) | 47.7 | CH |
| 7a | 2.11 (dd, 1H, *J*=12.9, 3.2Hz) | 39.6 | $CH_2$ |
| 7b | 2.69 (t, 1H, *J*=12.7Hz) | | |
| 8 | | 134.7 | C |
| 9 | 5.19 (t, 1H, *J*=8.4Hz) | 122.6 | CH |
| 10a | 2.18 (dd, 1H, *J*=12.8, 6.9Hz) | 33.7 | $CH_2$ |
| 10b | 2.24 (dt, 1H, *J*=12.7, 10.0Hz) | | |
| 11 | 3.26 (overlapped, 1H) | 75.5 | CH |
| 12a | 1.67 (m, 1H) | 31.2 | $CH_2$ |
| 12b | 1.86 (ddd, 1H, *J*=14.0, 11.3, 2.5 Hz) | | |
| 13 | 5.39 (q, 1H, *J*=2.9Hz) | 67.1 | CH |
| 14 | 3.26 (overlapped, 1H) | 75.5 | CH |
| 15 | 4.98 (d, 1H, *J* =1.8Hz) | 75.1 | CH |
| 16 | | 112.0 | C |
| 17 | | * | C |
| 18 | | * | C |
| 19 | 6.00 (s, 1H) | 105.0 | CH |
| 20 | | * | C |
| 21 | | * | C |
| 8-$CH_3$ | 1.68 (s, 3H) | 16.8 | $CH_3$ |
| 14-$OCH_3$ | 3.41 (s, 3H) | 59.3 | $CH_3$ |
| 1` | 1.47 (m, 1H) | 33.9 | CH |
| 2`a | 1.40 (m, 1H) | 35.8 | $CH_2$ |
| 2’b | 1.58 (m, 1H) | | |
| 3`a | 1.39 (m, 1H) | 25.2 | $CH_2$ |
| 3’b | 1.53 (m, 1H) | | |
| 4` | 1.31 (m, 2H) | 31.9 | $CH_2$ |
| 5` | 1.32 (m, 2H) | 22.7 | $CH_2$ |
| 6` | 0.90 (t, 3H, *J*=6.9 Hz) | 13.9 | $CH_3$ |

$^1$H NMR: 600 MHz; $^{13}$C NMR: 150 MHz (in DMSO-$d_6$).

* The interconversion of the benzoquinone moiety led to the missing of these carbon signals.

Supplementary Table 9. NMR data of compound **8**.

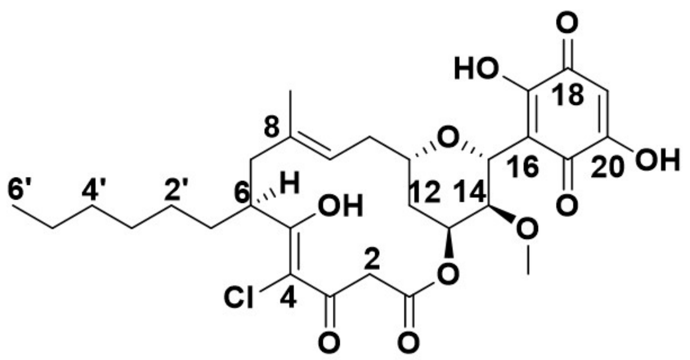

(8)

| NO. | $\delta_H$ (mult., *J* in Hz) | $\delta_C$ | C-type |
|---|---|---|---|
| 1 | | 165.2 | C |
| 2a | 3.54 (d, 1H, *J*=15.7Hz) | 47.0 | $CH_2$ |
| 2b | 3.83 (d, 1H, *J*=15.7Hz) | | |
| 3 | | 189.7 | C |
| 4 | | 109.1 | C |
| 5 | | 188.5 | C |
| 6 | 3.29 (m, 1H) | 42.2 | CH |
| 7a | 2.07 (dd, 1H, *J*=12.9, 2.4Hz) | 42.4 | $CH_2$ |
| 7b | 2.48 (t, 1H, *J*=12.6) | | |
| 8 | | 135.5 | C |
| 9 | 5.16 (t, 1H, *J*=8.4Hz) | 122.0 | CH |
| 10a | 2.17 (m, 1H) | 33.6 | $CH_2$ |
| 10b | 2.21 (m, 1H) | | |
| 11 | 3.25 (overlapped, 1H) | 75.5 | CH |
| 12a | 1.68 (m, 1H) | 31.1 | $CH_2$ |
| 12b | 1.84 (ddd, 1H, *J*=14.1, 11.2, 2.5 Hz) | | |
| 13 | 5.39 (q, 1H, *J*=3.0Hz) | 67.1 | CH |
| 14 | 3.25 (overlapped, 1H) | 75.5 | CH |
| 15 | 4.95 (d, 1H, *J* =1.7Hz) | 75.2 | CH |
| 16 | | 111.2 | C |
| 17 | | * | C |
| 18 | | * | C |
| 19 | 5.99 (s, 1H) | 105.0 | CH |
| 20 | | * | C |
| 21 | | * | C |
| 8-$CH_3$ | 1.67 (s, 3H) | 16.7 | $CH_3$ |
| 14-$OCH_3$ | 3.39 (s, 3H) | 59.2 | $CH_3$ |
| 1`a | 1.47 (ddt, 1H, *J* =14.3, 10.1, 5.2 Hz) | 33.9 | $CH_2$ |
| 1`b | 1.67 (m, 1H) | | |
| 2`a | 1.27 (m, 1H) | 27.2 | $CH_2$ |
| 2'b | 1.34 (m, 1H) | | |
| 3` | 1.26 (m, 2H) | 29.5 | $CH_2$ |
| 4` | 1.26 (m, 2H) | 31.6 | $CH_2$ |
| 5` | 1.28 (m, 2H) | 22.5 | $CH_2$ |
| 6` | 0.88 (t, 3H, *J*=6.9 Hz) | 14.0 | $CH_3$ |

$^1$H NMR: 600 MHz; $^{13}$C NMR: 150 MHz (in DMSO-$d_6$).

* The interconversion of the benzoquinone moiety led to the missing of these carbon signals.

Supplementary Table 10. NMR data of compound **9**.

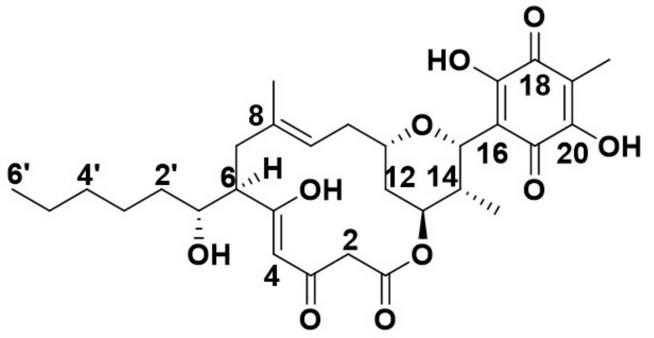

(9)

| NO. | $\delta_H$ (mult., *J* in Hz) | $\delta_C$ | C-type |
|---|---|---|---|
| 1 | | 167.0 | C |
| 2 | 3.43 (td, 2H, *J*=21.4, 7.4Hz) | 47.5 | $CH_2$ |
| 3 | | 190.0 | C |
| 4 | 5.48 (s, 1H) | 101.2 | CH |
| 5 | | 191.2 | C |
| 6 | 2.30 (ddd, *J*=12.4, 5.7, 3.2Hz) | 52.6 | CH |
| 7a | 2.08 (dd, 1H, *J*=13.1, 3.2Hz) | 39.7 | $CH_2$ |
| 7b | 2.59 (t, 1H, *J*=12.7 Hz) | | |
| 8 | | 135.6 | C |
| 9 | 5.12 (overlapped, 1H) | 121.6 | CH |
| 10a | 2.16 (dt, 1H, *J*=7.4, 2.7Hz) | 34.5 | $CH_2$ |
| 10b | 2.24 (m, 1H) | | |
| 11 | 3.36 (ddd, 1H, *J*=12.9, 7.1, 4.3Hz) | 76.0 | CH |
| 12 | 1.68 (m, 2H) | 30.7 | $CH_2$ |
| 13 | 5.14 (overlapped, 1H) | 72.1 | CH |
| 14 | 2.16 (dtd, 1H, *J*=10.0, 7.2, 2.6Hz) | 34.5 | CH |
| 15 | 5.21 (d, 1H, *J*=2.7Hz) | 76.0 | CH |
| 16 | | 112.4 | C |
| 17 | | * | C |
| 18 | | * | C |
| 19 | | 114.2 | C |
| 20 | | * | C |
| 21 | | * | C |
| 8-$CH_3$ | 1.61 (s, 3H) | 17.1 | $CH_3$ |
| 14-$CH_3$ | 1.01 (d, 3H, *J*=7.4Hz) | 11.5 | $CH_3$ |
| 19-$CH_3$ | 1.93 (s, 3H) | 7.9 | $CH_3$ |
| 1` | 3.69 (m, 1H) | 73.7 | CH |
| 2`a | 1.40 (m, 1H) | 36.0 | $CH_2$ |
| 2`b | 1.50 (m, 1H) | | |
| 3`a | 1.35 (m, 1H) | 25.4 | $CH_2$ |
| 3`b | 1.49 (m, 1H) | | |
| 4` | 1.31 (m, 2H) | 31.7 | $CH_2$ |
| 5` | 1.29 (m, 2H) | 22.6 | $CH_2$ |
| 6` | 0.89 (t, 3H, *J*=6.9Hz) | 14.1 | |

$^1H$ NMR: 600 MHz; $^{13}C$ NMR: 150 MHz (in DMSO-$d_6$).

* The interconversion of the benzoquinone moiety led to the missing of these carbon signals.

Supplementary Table 11. NMR data of compound **10**.

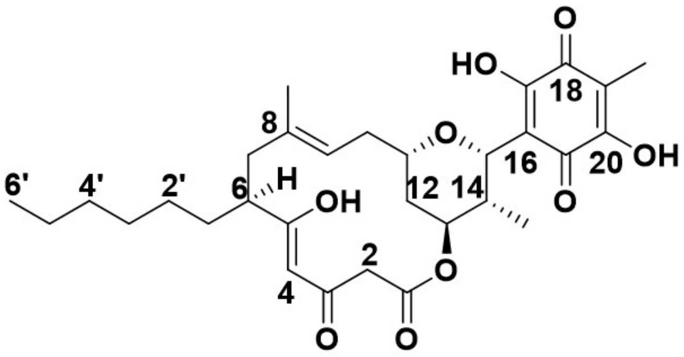

**(10)**

| NO. | $\delta_H$ (mult., $J$ in Hz) | $\delta_C$ | C-type |
|---|---|---|---|
| 1 | | 167.2 | C |
| 2 | 3.42 (m, 2H) | 48.5 | $CH_2$ |
| 3 | | 191.4 | C |
| 4 | 5.44 (s, 1H) | 99.8 | CH |
| 5 | | 190.5 | C |
| 6 | 2.17 (overlapped, 1H) | 47.2 | CH |
| 7a | 2.02 (dd, 1H, $J$=12.8, 3.0Hz) | 42.5 | $CH_2$ |
| 7b | 2.36 (t, 1H, $J$=12.5 Hz) | | |
| 8 | | 136.2 | C |
| 9 | 5.09 (t, 1H, $J$=8.0Hz) | 121.3 | CH |
| 10 | 2.23 (m, 2H) | 34.4 | $CH_2$ |
| 11 | 3.35 (m, 1H) | 76.5 | CH |
| 12a | 1.65 (ddd, 1H, $J$=14.4, 11.1, 2.5Hz) | 30.7 | $CH_2$ |
| 12b | 1.69 (overlapped, 1H) | | |
| 13 | 5.14 (q, 1H, $J$=2.8 Hz) | 72.1 | CH |
| 14 | 2.16 (overlapped, 1H) | 35.0 | CH |
| 15 | 5.21 (d, 1H, $J$=2.7Hz) | 76.5 | CH |
| 16 | | 112.4 | C |
| 17 | | * | C |
| 18 | | * | C |
| 19 | | 114.1 | C |
| 20 | | * | C |
| 21 | | * | C |
| 8-$CH_3$ | 1.59 (d, 3H, $J$=1.4Hz) | 17.0 | $CH_3$ |
| 14-$CH_3$ | 1.01 (d, 3H, $J$=7.4Hz) | 11.6 | $CH_3$ |
| 19-$CH_3$ | 1.93 (s, 3H) | 7.8 | $CH_3$ |
| 1`a | 1.38 (m, 1H) | 33.2 | $CH_2$ |
| 1`b | 1.68 (overlapped, 1H) | | |
| 2` | 1.26 (overlapped, 2H) | 29.3 | $CH_2$ |
| 3` | 1.26 (overlapped, 2H) | 27.7 | $CH_2$ |
| 4` | 1.26 (overlapped, 2H) | 31.6 | $CH_2$ |
| 5` | 1.26 (overlapped, 2H) | 22.6 | $CH_2$ |
| 6` | 0.88 (t, 3H, $J$=6.9Hz) | 14.1 | $CH_3$ |

$^1$H NMR: 600 MHz; $^{13}$C NMR: 150 MHz (in DMSO-$d_6$).

* The interconversion of the benzoquinone moiety led to the missing of these carbon signals.

Supplementary Table 12. NMR data of compound **11**.

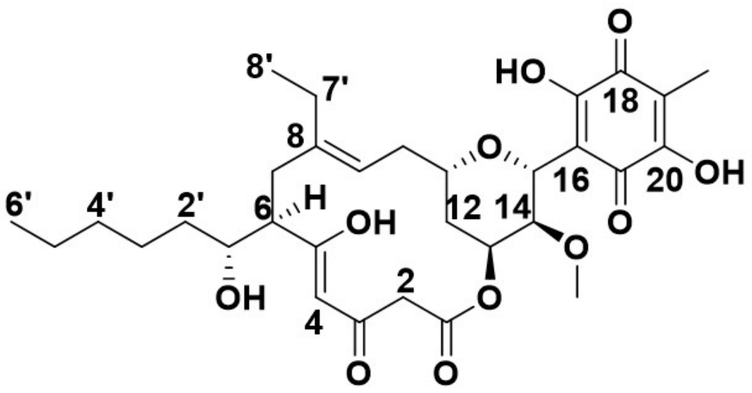

**(11)**

| NO. | $\delta_H$ (mult., *J* in Hz) | $\delta_C$ | C-type |
|---|---|---|---|
| 1 | | 166.7 | C |
| 2a | 3.43 (d, 1H, *J*=14.1Hz) | 47.3 | $CH_2$ |
| 2b | 3.46 (d, 1H, *J*=14.1Hz) | | |
| 3 | | 189.3 | C |
| 4 | 5.46 (s, 1H) | 101.3 | CH |
| 5 | | 191.3 | C |
| 6 | 2.27 (ddd, 1H, *J*=12.3, 5.7, 3.1Hz) | 52.8 | CH |
| 7a | 2.20 (overlapped, 1H) | 35.5 | $CH_2$ |
| 7b | 2.50 (t, 1H, *J*=12.8Hz) | | |
| 8 | | 141.1 | C |
| 9 | 5.04 (overlapped, 1H) | 121.0 | CH |
| 10 | 2.22 (overlapped, 1H) | 33.6 | $CH_2$ |
| 11 | 3.29 (t, 1H, *J*=10.3Hz) | 74.9 | CH |
| 12a | 1.74 (d, 1H, *J*=14.3Hz) | 30.1 | $CH_2$ |
| 12b | 1.80 (d, 1H, *J*=14.3Hz) | | |
| 13 | 5.34 (q, 1H, *J*=3.0Hz) | 66.3 | CH |
| 14 | 3.28 (m, 1H) | 75.8 | CH |
| 15 | 5.03 (overlapped, 1H) | 74.6 | CH |
| 16 | | 111.2 | C |
| 17 | | * | C |
| 18 | | * | C |
| 19 | | 114.2 | C |
| 20 | | * | C |
| 21 | | * | C |
| 14-$OCH_3$ | 3.37 (s, 3H) | 59.2 | $CH_3$ |
| 19-$CH_3$ | 1.94 (s, 3H) | 7.7 | $CH_3$ |
| 1` | 3.69 (ddd, 1H, *J*=9.0, 5.8, 2.9Hz) | 73.9 | CH |
| 2`a | 1.41 (m, 1H) | 35.9 | $CH_2$ |
| 2`b | 1.50 (m, 1H) | | |
| 3`a | 1.35 (m, 2H) | 25.3 | $CH_2$ |
| 3`b | 1.50 (m, 2H) | | |
| 4` | 1.31 (m, 2H) | 31.6 | $CH_2$ |
| 5` | 1.31 (m, 2H) | 23.2 | $CH_2$ |
| 6` | 0.90 (t, 3H, *J*=6.9Hz) | 13.9 | $CH_3$ |
| 7`a | 1.82 (m, 1H) | 23.21 | $CH_2$ |
| 7`b | 2.10 (dq, 1H, *J*=14.8, 7.5Hz) | | |
| 8` | 0.95 (t, 3H, *J*=7.2Hz) | 12.8 | $CH_3$ |

$^1$H NMR: 600 MHz; $^{13}$C NMR: 150 MHz (in DMSO-$d_6$).

* The interconversion of the benzoquinone moiety led to the missing of these carbon signals.

Supplementary Table 13. NMR data of compound **12**.

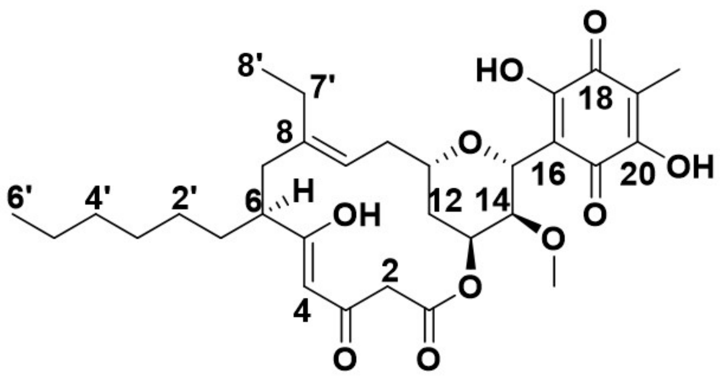

**(12)**

| NO. | $\delta_H$ (mult., *J* in Hz) | $\delta_C$ | C-type |
|---|---|---|---|
| 1 | | 167.0 | C |
| 2a | 3.43 (d, 1H, *J*=13.9Hz) | 48.1 | $CH_2$ |
| 2b | 3.48 (d, 1H, *J*=13.9Hz) | | |
| 3 | | 190.8 | C |
| 4 | 5.44 (s, 1H) | 99.9 | CH |
| 5 | | 190.8 | C |
| 6 | 2.17 (overlapped, 1H) | 48.1 | CH |
| 7a | 2.16 (overlapped, 1H) | 38.3 | $CH_2$ |
| 7b | 2.31 (m, 1H) | | |
| 8 | | 141.6 | C |
| 9 | 5.04 (t, 1H, *J*=8.3Hz) | 120.6 | CH |
| 10 | 2.24 (overlapped, 2H) | 33.6 | $CH_2$ |
| 11 | 3.30 (overlapped, 1H) | 75.6 | CH |
| 12 | 1.79 (q, 2H, *J*=2.5Hz) | 30.2 | $CH_2$ |
| 13 | 5.37 (q, 1H, *J*=2.9Hz) | 66.4 | CH |
| 14 | 3.30 (overlapped, 1H) | 75.6 | CH |
| 15 | 5.07 (d, 1H, *J*=1.7Hz) | 74.7 | CH |
| 16 | | 111.2 | C |
| 17 | | * | C |
| 18 | | * | C |
| 19 | | 114.2 | C |
| 20 | | * | C |
| 21 | | * | C |
| 14-$OCH_3$ | 3.40 (s, 3H) | 59.2 | $CH_3$ |
| 19-$CH_3$ | 1.97 (s, 3H) | 7.8 | $CH_3$ |
| 1`a | 1.43 (ddd, 1H, *J*=13.8, 8.7, 5.0Hz) | 33.4 | CH |
| 1`b | 1.71 (m, 1H) | | |
| 2` | 1.29 (overlapped, 2H) | 29.2 | $CH_2$ |
| 3` | 1.29 (overlapped, 2H) | 27.5 | $CH_2$ |
| 4` | 1.29 (overlapped, 2H) | 31.6 | $CH_2$ |
| 5` | 1.30 (overlapped, 2H) | 22.5 | $CH_2$ |
| 6` | 0.91 (q, 3H, *J*=6.2Hz) | 14.1 | $CH_3$ |
| 7`a | 1.86 (m, 1H) | 23.4 | $CH_2$ |
| 7`b | 2.11 (m, 1H) | | |
| 8` | 0.98 (t, 3H, *J*=7.5Hz) | 13.0 | $CH_3$ |

$^{1}$H NMR: 600 MHz; $^{13}$C NMR: 150 MHz (in DMSO-$d_6$).

* The interconversion of the benzoquinone moiety led to the missing of these carbon signals.

Supplementary Table 14. NMR data of compound **13a**.

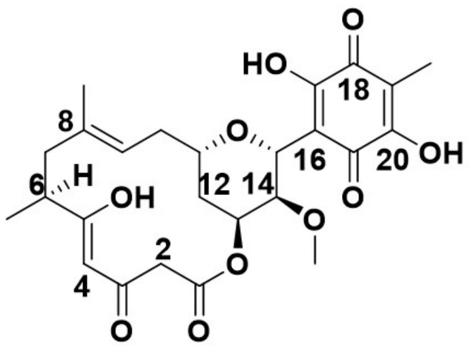

**(13a)**

| NO. | $\delta_H$ (mult., *J* in Hz) | $\delta_C$ | C-type |
|---|---|---|---|
| 1 | | 167.1 | C |
| 2a | 3.39 (d, 1H, *J*=13.7Hz) | 48.2 | $CH_2$ |
| 2b | 3.45 (d, 1H, *J*=13.7Hz) | | |
| 3 | | 190.9 | C |
| 4 | 5.42 (s, 1H) | 98.3 | CH |
| 5 | | 190.9 | C |
| 6 | 2.33 (m, 1H) | 41.1 | CH |
| 7a | 1.98 (m, 1H) | 43.3 | $CH_2$ |
| 7b | 2.38 (overlapped, 1H) | | |
| 8 | | 135.9 | C |
| 9 | 5.08 (t, 1H, *J*=8.4Hz) | 121.4 | CH |
| 10a | 2.21 (m, 1H) | 34.2 | $CH_2$ |
| 10b | 2.35 (overlapped, 1H) | | |
| 11 | 3.29 (m, 1H) | 74.7 | CH |
| 12a | 1.60 (overlapped, 1H) | 30.4 | $CH_2$ |
| 12b | 1.78 overlapped, 1H) | | |
| 13 | 5.34 (q, 1H, *J*=3.0Hz) | 66.2 | CH |
| 14 | 3.27 (m, 1H) | 75.7 | CH |
| 15 | 5.03 (d, 1H, *J* =1Hz) | 74.7 | CH |
| 16 | | 111.1 | C |
| 17 | | * | C |
| 18 | | * | C |
| 19 | | 114.0 | C |
| 20 | | * | C |
| 21 | | * | C |
| 6-$CH_3$ | 1.18 (d, 3H, *J* = 6.3Hz) | 18.4 | $CH_3$ |
| 8-$CH_3$ | 1.58 (d, 3H, J=1.6Hz) | 18.2 | $CH_3$ |
| 14-$OCH_3$ | 3.36 (s, 3H) | 59.1 | $CH_3$ |
| 19-$CH_3$ | 1.94 (s, 3H) | 7.5 | $CH_3$ |

$^1$H NMR: 600 MHz; $^{13}$C NMR: 150 MHz (in CDCl3).

* The interconversion of the benzoquinone moiety led to the missing of these carbon signals.

Supplementary Table 15. NMR data of compound **13b**.

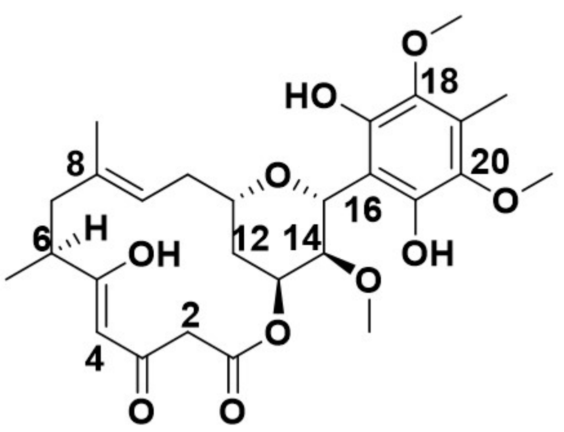

(13b)

| NO. | $\delta_H$ (mult., *J* in Hz) | $\delta_C$ | C-type |
|---|---|---|---|
| 1 | | 167.1 | C |
| 2a | 3.39 (d, 1H, *J*=14.0Hz) | 48.3 | $CH_2$ |
| 2b | 3.44 (d, 1H, *J*=14.0Hz) | | |
| 3 | | 191.5 | C |
| 4 | 5.46 (s, 1H) | 98.5 | CH |
| 5 | | 191.2 | C |
| 6 | 2.35 (m, 1H) | 41.4 | CH |
| 7a | 1.99 (d, 1H, *J*=10.66 Hz) | 44.0 | $CH_2$ |
| 7b | 2.36 (m, 1H) | | |
| 8 | | 135.5 | C |
| 9 | 5.11 (t, 1H, *J*=8.4Hz) | 121.8 | CH |
| 10a | 2.21 (overlapped, 1H) | 34.3 | $CH_2$ |
| 10b | 2.29 (m, 1H) | | |
| 11 | 3.31 (m, 1H) | 74.8 | CH |
| 12a | 1.60 (overlapped, 1H) | 30.7 | $CH_2$ |
| 12b | 1.82 (ddd, 1H, *J*=14.0, 11.3, 2.5 Hz) | | |
| 13 | 5.32 (q, 1H, *J*=2.9Hz) | 67.9 | CH |
| 14 | 3.32 (m, 1H) | 76.6 | CH |
| 15 | 5.26 (d, 1H, *J* =1.6Hz) | 75.4 | CH |
| 16 | | 107.38 | C |
| 17 | | 144.3 | C |
| 18 | | 138.4 | C |
| 19 | | 123.8 | C |
| 20 | | 138.4 | C |
| 21 | | 144.3 | C |
| 6-$CH_3$ | 1.18 (d, 3H, *J* =6.1Hz) | 18.5 | $CH_3$ |
| 8-$CH_3$ | 1.60 (s, 3H) | 17.0 | $CH_3$ |
| 14-$OCH_3$ | 3.21 (s, 3H) | 59.6 | $CH_3$ |
| 18-OCH3 | 3.74 (s, 3H) | 60.6 | $CH_3$ |
| 19-$CH_3$ | 2.21 (s, 3H) | 9.5 | $CH_3$ |
| 20-OCH3 | 3.74 (s, 3H) | 60.6 | $CH_3$ |

$^1$H NMR: 600 MHz; $^{13}$C NMR: 150 MHz (in DMSO-$d_6$).

Supplementary Table 16. NMR data of compound **14a**.

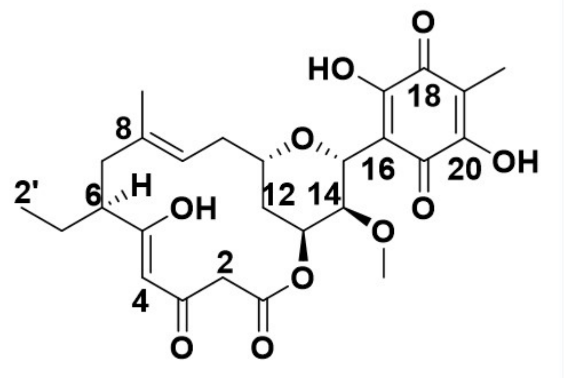

**(14a)**

| NO. | $\delta_H$ (mult., *J* in Hz) | $\delta_C$ | C-type |
|---|---|---|---|
| 1 | | 166.9 | C |
| 2a | 3.40 (d, 1H) | 48.2 | $CH_2$ |
| 2b | 3.46 (m, 1H) | | |
| 3 | | 191.1 | C |
| 4 | 5.44 (s, 1H) | 100.0 | CH |
| 5 | | 190.1 | C |
| 6 | 2.09 (m, 1H) | 48.9 | CH |
| 7a | 2.04 (dd, 1H, *J*=12.8, 3.2Hz) | 42.1 | $CH_2$ |
| 7b | 2.36 (t, 1H, *J*=12.5Hz) | | |
| 8 | | 135.9 | C |
| 9 | 5.09 (t, 1H, *J*=8.4Hz) | 121.5 | CH |
| 10 | 2.21 (m, 1H) | 34.1 | $CH_2$ |
| 11 | 3.28 (overlapped, 1H) | 75.3 | CH |
| 12 | 1.75 (ddd, 1H, *J* = 14.0, 11.2, 2.6 Hz) | 30.4 | $CH_2$ |
| 13 | 5.35 (q, 1H, *J*=3.0Hz) | 66.3 | CH |
| 14 | 3.28 (overlapped, 1H) | 75.3 | CH |
| 15 | 5.04 (d, 1H, *J* =1.7Hz) | 74.8 | CH |
| 16 | | 111.2 | C |
| 17 | | * | C |
| 18 | | * | C |
| 19 | | 114.2 | C |
| 20 | | * | C |
| 21 | | * | C |
| 8-$CH_3$ | 1.59 (s, 3H) | 17.0 | $CH_3$ |
| 14-$OCH_3$ | 3.37 (s, 3H) | 59.2 | $CH_3$ |
| 19-$CH_3$ | 1.95 (s, 3H) | 7.7 | $CH_3$ |
| 1\`a | 1.47 (m, 1H) | 26.2 | $CH_2$ |
| 1\`b | 1.71 (m, 1H) | | |
| 2\` | 0.89 (t, 3H, *J* =7.4Hz) | 12.1 | $CH_3$ |

$^1$H NMR: 600 MHz; $^{13}$C NMR: 150 MHz (in DMSO-$d_6$).

* The interconversion of the benzoquinone moiety led to the missing of these carbon signals.

Supplementary Table 17. NMR data of compound **15a**.

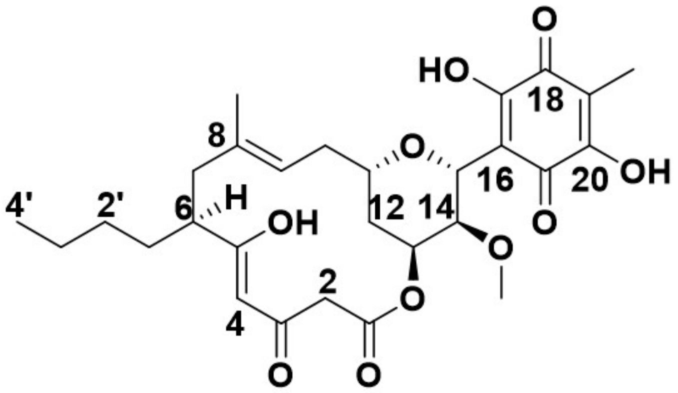

**(15a)**

| NO. | δH (mult., *J* in Hz) | δC | C-type |
|---|---|---|---|
| 1 | | 167.1 | C |
| 2a | 3.40 (d, 1H, *J*=13.8Hz) | 48.2 | $CH_2$ |
| 2b | 3.46 (d, 1H, *J*=13.8Hz) | | |
| 3 | | 191.0 | C |
| 4 | 5.43 (s, 1H) | 99.7 | CH |
| 5 | | 190.5 | C |
| 6 | 2.16 (m, 1H) | 46.9 | CH |
| 7a | 2.02 (m, 1H) | 42.5 | $CH_2$ |
| 7b | 2.36 (t, 1H, *J*=12.5) | | |
| 8 | | 136.0 | C |
| 9 | 5.08 (t, 1H, *J*=8.4Hz) | 121.4 | CH |
| 10 | 2.20 (t, 1H, *J*=7.9Hz) | 34.1 | $CH_2$ |
| 11 | 3.28 (overlapped, 1H) | 74.7 | CH |
| 12a | 1.70 (m, 1H) | 30.4 | $CH_2$ |
| 12b | 1.78 (ddd, 1H, *J*=14.1, 11.3, 2.5 Hz) | | |
| 13 | 5.35 (q, 1H, *J*=2.9Hz) | 66.3 | CH |
| 14 | 3.27 (overlapped, 1H) | 75.7 | CH |
| 15 | 5.03 (d, 1H, *J* =1.7Hz) | 74.7 | CH |
| 16 | | 111.2 | C |
| 17 | | * | C |
| 18 | | * | C |
| 19 | | 114.1 | CH |
| 20 | | * | C |
| 21 | | * | C |
| 8-$CH_3$ | 1.59 (s, 3H) | 17.00 | $CH_3$ |
| 14-$OCH_3$ | 3.37 (s, 3H) | 59.3 | $CH_3$ |
| 1\`a | 1.40 (m, 1H) | 33.0 | $CH_2$ |
| 1\`b | 1.69 (m, 1H) | | |
| 2\` | 1.23 (m, 1H) | 30.0 | $CH_2$ |
| 3\` | 1.30 (m, 2H) | 22.7 | $CH_2$ |
| 4\` | 0.88 (t, 3H, *J*=7.2 Hz) | 13.9 | $CH_3$ |

$^{1}$H NMR: 600 MHz; $^{13}$C NMR: 150 MHz (in DMSO-$d_6$).

* The interconversion of the benzoquinone moiety led to the missing of these carbon signals.

Supplementary Table 18. NMR data of compound **16**.

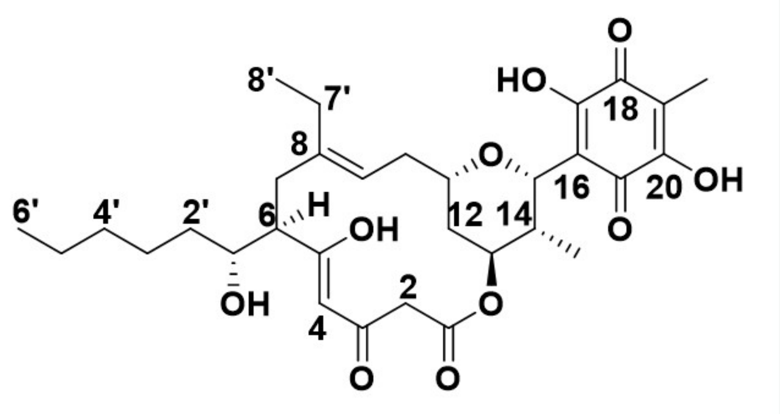

**(16)**

| NO. | δ_H (mult., *J* in Hz) | δ_C | C-type |
|---|---|---|---|
| 1 | | 166.9 | C |
| 2a | 3.42 (d, H, *J*=14.1Hz) | 47.7 | $CH_2$ |
| 2b | 3.46 (d, 1H, J=14.1Hz) | | |
| 3 | | 189.4 | C |
| 4 | 5.47 (s, 1H) | 101.4 | CH |
| 5 | | 191.5 | C |
| 6 | 2.26 (overlapped, 1H) | 52.5 | CH |
| 7a | 2.17 (m, 1H) | 35.4 | $CH_2$ |
| 7b | 2.50 (t, 1H, *J*=12.7 Hz) | | |
| 8 | | 141.0 | C |
| 9 | 5.04 (t, 1H, *J*=8.3Hz) | 120.9 | CH |
| 10 | 2.26 (overlapped, 2H) | 52.5 | $CH_2$ |
| 11 | 3.36 (m, 1H) | 76.5 | CH |
| 12a | 1.66 (m, 1H) | 30.2 | $CH_2$ |
| 12b | 1.72 (m, 1H) | | |
| 13 | 5.14 (t, 1H, J=2.9Hz) | 72.1 | CH |
| 14 | 2.16 (m, 1H) | 35.0 | CH |
| 15 | 5.21 (d, 1H, *J*=2.7Hz) | 76.5 | CH |
| 16 | | 112.2 | C |
| 17 | | * | C |
| 18 | | * | C |
| 19 | | 114.1 | C |
| 20 | | * | C |
| 21 | | * | C |
| 14-$CH_3$ | 1.01 (d, 3H, *J*=7.4Hz) | 11.3 | $CH_3$ |
| 19-$CH_3$ | 1.93 (s, 3H) | 7.6 | $CH_3$ |
| 1` | 3.68 (m, 1H) | 72.1 | CH |
| 2`a | 1.40 (m, 1H) | 35.9 | $CH_2$ |
| 2`b | 1.49 (m, 1H) | | |
| 3`a | 1.35 (m, 1H) | 25.1 | $CH_2$ |
| 3`b | 1.48 (m, 1H) | | |
| 4` | 1.29 (m, 2H) | 31.6 | $CH_2$ |
| 5` | 1.31 (m, 2H) | 22.8 | $CH_2$ |
| 6` | 0.89 (d, 3H, *J*=7.1Hz) | 13.9 | $CH_3$ |
| 7`a | 1.83 (m, 1H) | 23.2 | $CH_2$ |
| 7`b | 2.10 (m, 1H) | | |
| 8` | 0.96 (t, 3H, *J*=7.5Hz) | 12.8 | $CH_3$ |

$^{1}$H NMR: 600 MHz; $^{13}$C NMR: 150 MHz (in DMSO-$d_6$).

* The interconversion of the benzoquinone moiety led to the missing of these carbon signals.

Supplementary Table 19. NMR data of compound **17**.

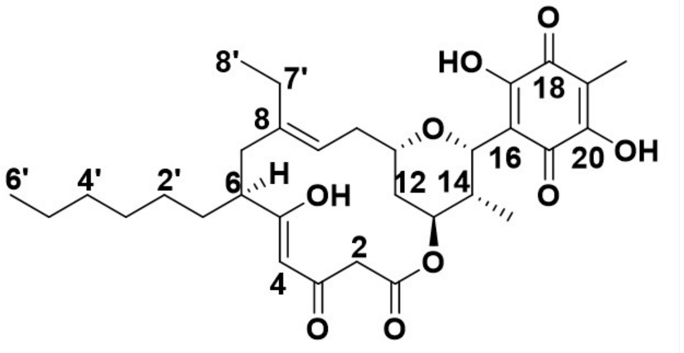

**(17)**

| NO. | $\delta_H$ (mult., *J* in Hz) | $\delta_C$ | C-type |
|---|---|---|---|
| 1 | | 167.1 | C |
| 2a | 3.41 (d, 1H, *J*=13.8Hz) | 48.3 | $CH_2$ |
| 2b | 3.44 (d, 1H, *J*=13.8Hz) | | |
| 3 | | 191.2 | C |
| 4 | 5.42 (s, 1H) | 100.0 | CH |
| 5 | | 190.5 | C |
| 6 | 2.13 (overlapped, 1H) | 47.4 | CH |
| 7a | 2.13 (overlapped, 1H) | 38.2 | $CH_2$ |
| 7b | 2.27 (t, 1H, *J*=13.3 Hz) | | |
| 8 | | 141.6 | C |
| 9 | 5.01 (t, 1H, *J*=8.3Hz) | 120.1 | CH |
| 10 | 2.23 (m, 2H) | 33.8 | $CH_2$ |
| 11 | 3.34 (m, 1H) | 75.9 | CH |
| 12a | 1.64 (overlapped, 1H) | 30.4 | $CH_2$ |
| 12b | 1.73 (overlapped, 1H) | | |
| 13 | 5.14 (q, 1H, *J*=2.8 Hz) | 71.8 | CH |
| 14 | 2.16 (m, 1H) | 35.0 | CH |
| 15 | 5.22 (d, 1H, *J*=2.7Hz) | 76.2 | CH |
| 16 | | 112.2 | C |
| 17 | | * | C |
| 18 | | * | C |
| 19 | | 114.1 | C |
| 20 | | * | C |
| 21 | | * | C |
| 14-$CH_3$ | 1.01 (d, 3H, *J*=7.4Hz) | 11.5 | $CH_3$ |
| 19-$CH_3$ | 1.93 (s, 3H) | 7.6 | $CH_3$ |
| 1\`a | 1.40 (m, 1H) | 33.8 | $CH_2$ |
| 1\`b | 1.67 (overlapped, 1H) | | |
| 2\` | 1.26 (overlapped, 2H) | 29.3 | $CH_2$ |
| 3\` | 1.25 (overlapped, 2H) | 27.1 | $CH_2$ |
| 4\` | 1.29 (overlapped, 2H) | 31.6 | $CH_2$ |
| 5\` | 1.28 (overlapped, 2H) | 22.4 | $CH_2$ |
| 6\` | 0.88 (t, 3H, *J*=7.0Hz) | 14.0 | $CH_3$ |
| 7\`a | 1.81 (m, 1H) | 23.1 | $CH_2$ |
| 7\`b | 2.07 (m, 1H) | | |
| 8\` | 0.95 (t, 3H, *J*=7.5Hz) | 13.1 | $CH_3$ |

$^1$H NMR: 600 MHz; $^{13}$C NMR: 150 MHz (in DMSO-$d_6$).

* The interconversion of the benzoquinone moiety led to the missing of these carbon signals.

Supplementary Table 20. NMR data of compound **18b**.

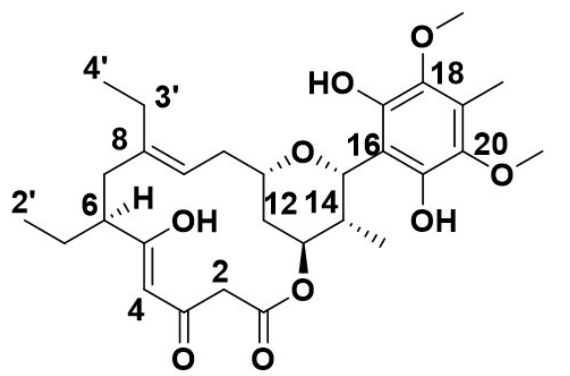

(18b)

| NO. | δH (mult., *J* in Hz) | δC | C-type |
|---|---|---|---|
| 1 | | 167.3 | C |
| 2 | 3.42(d, 2H, *J*=13.8Hz) | 48.4 | $CH_2$ |
| 3 | | 191.3 | C |
| 4 | 5.46 (overlapped, 1H) | 100.2 | CH |
| 5 | | 190.5 | C |
| 6 | 2.07 (m, 1H) | 49.2 | CH |
| 7a | 2.14 (m, 1H) | 38.1 | $CH_2$ |
| 7b | 2.26 (d, 1H, *J*=12.8Hz) | | |
| 8 | | 141.3 | C |
| 9 | 5.04 (t, 1H, *J*=8.1Hz) | 121.1 | CH |
| 10a | 2.22 (m, 1H) | 34.1 | $CH_2$ |
| 10b | 2.31 (m, 1H) | | |
| 11 | 3.38 (m, 1H) | 75.8 | CH |
| 12a | 1.68 (m, 1H) | 30.5 | $CH_2$ |
| 12b | 1.74 (m, 1H) | | |
| 13 | 5.17 (td, 1H, *J*=22.2, 1.7Hz) | 72.7 | CH |
| 14 | 2.18 (overlapped, 1H) | 35.4 | CH |
| 15 | 5.46 (overlapped, 1H) | 75.8 | CH |
| 16 | | 108.5 | C |
| 17 | | 141.7 | C |
| 18 | | 138.7 | C |
| 19 | | 123.4 | C |
| 20 | | 138.7 | C |
| 21 | | 141.7 | C |
| 14-$CH_3$ | 0.98 (m, 3H) | 11.6 | $CH_3$ |
| 18-$OCH_3$ | 3.73 (s, 3H) | 60.7 | |
| 19-$CH_3$ | 2.20 (s, 3H) | 9.4 | $CH_3$ |
| 20-$OCH_3$ | 3.73 (s, 3H) | 60.7 | |
| 1\`a | 1.46 (m, 1H) | 26.4 | CH |
| 1\`b | 1.70 (m, 1H) | | |
| 2\` | 0.89 (t, 3H, *J*=7.4 Hz) | 12.2 | $CH_2$ |
| 3\`a | 1.83 (dq, 1H, *J*=14.8, 7.6 Hz) | 23.4 | $CH_2$ |
| 3\`b | 2.09 (overlapped, 1H) | | |
| 4\` | 0.95 (t, 3H, *J*=7.6Hz) | 12.4 | $CH_2$ |

$^1H$ NMR: 600 MHz; $^{13}C$ NMR: 150 MHz (in DMSO-$d_6$).

Supplementary Table 21. NMR data of compound **18c**.

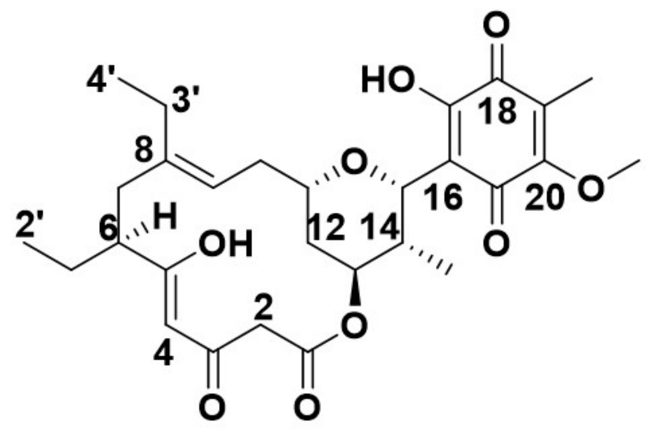

(18c)

| NO. | $\delta_H$ (mult., *J* in Hz) | $\delta_C$ | C-type |
|---|---|---|---|
| 1 | | 167.1 | C |
| 2 | 3.43 (s, 1H) | 48.4 | $CH_2$ |
| 3 | | 191.5 | C |
| 4 | 5.44 (s, 1H) | 100.0 | CH |
| 5 | | 190.0 | C |
| 6 | 2.06 (m, 1H) | 48.4 | CH |
| 7a | 2.14 (m, 1H) | 38.1 | $CH_2$ |
| 7b | 2.27 (t, 1H, *J*=12.5Hz) | | |
| 8 | | 141.8 | C |
| 9 | 5.02 (t, *J*=8.3Hz, 1H) | 120.4 | CH |
| 10a | 2.22 (m, 1H) | 33.2 | $CH_2$ |
| 10b | 2.36 (q, 1H, *J*=7.7Hz) | | |
| 11 | 3.36 (ddt, 1H, *J*=12.0, 9.7, 2.4Hz) | 76.6 | CH |
| 12a | 1.63 (m, 1H) | 30.3 | $CH_2$ |
| 12b | 1.73 (m, 1H) | | |
| 13 | 5.14 (q, 1H, *J*=2.8Hz) | 71.8 | CH |
| 14 | 2.18 (m, 1H) | 35.1 | CH |
| 15 | 5.21 (d, 1H, *J*=2.7 Hz) | 76.6 | CH |
| 16 | | 114.6 | C |
| 17 | | 155.3 | C |
| 18 | | 183.2 | C |
| 19 | | 125.8 | C |
| 20 | | 155.5 | C |
| 21 | | 182.00 | C |
| 14-$CH_3$ | 1.01 (d, 3H, *J*=7.4Hz) | 11.4 | $CH_3$ |
| 19-$CH_3$ | 1.94 (s, 3H) | 8.0 | $CH_3$ |
| 20-$OCH_3$ | 4.01 (s, 3H) | 61.5 | $CH_3$ |
| 1`a | 1.46 (m, 1H) | 25.9 | $CH_2$ |
| 1`b | 1.70 (m, 1H) | | |
| 2` | 0.89 (t, 3H, *J*=7.4Hz) | 12.2 | $CH_3$ |
| 3`a | 1.82 (m, 1H) | 23.4 | $CH_2$ |
| 3`b | 2.08 (m, 1H) | | |
| 4` | 0.95 (t, 3H, *J*=7.5Hz) | 13.0 | $CH_3$ |

$^1$H NMR: 600 MHz; $^{13}$C NMR: 150 MHz (in DMSO-$d_6$).

Supplementary Table 22. NMR data of compound **19a**.

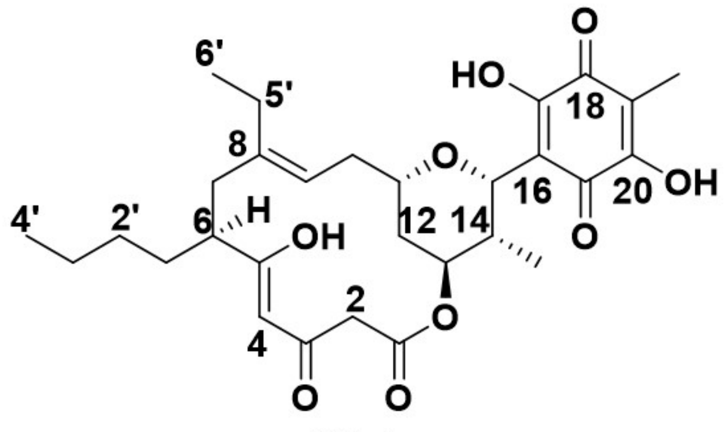

**(19a)**

| NO. | $\delta_H$ (mult., *J* in Hz) | $\delta_C$ | C-type |
|---|---|---|---|
| 1 | | 167.1 | C |
| 2a | 3.41(d, 1H, *J*=13.8Hz) | 48.4 | $CH_2$ |
| 2b | 3.45(d, 1H, *J*=13.8Hz) | | |
| 3 | | 191.2 | C |
| 4 | 5.44 (s, 1H) | 99.9 | CH |
| 5 | | 190.5 | C |
| 6 | 2.13 (overlapped, 1H) | 47.4 | CH |
| 7a | 2.13 (overlapped, 1H) | 38.6 | $CH_2$ |
| 7b | 2.28 (t, 1H, *J*=13.2Hz) | | |
| 8 | | 141.7 | C |
| 9 | 5.01 (t, 1H, *J*=8.3Hz) | 121.4 | CH |
| 10 | 2.23 (m, 2H) | 33.8 | $CH_2$ |
| 11 | 3.35 (m, 1H) | 76.6 | CH |
| 12a | 1.65 (m, 1H) | 30.4 | $CH_2$ |
| 12b | 1.72 (m, 1H) | | |
| 13 | 5.14 (q, 1H, *J*=2.8Hz) | 71.7 | CH |
| 14 | 2.16 (m, 1H) | 34.7 | CH |
| 15 | 5.22 (d, 1H, *J*=2.7Hz) | 76.6 | CH |
| 16 | | 112.3 | C |
| 17 | | * | C |
| 18 | | * | C |
| 19 | | 114.0 | C |
| 20 | | * | C |
| 21 | | * | C |
| 14-$CH_3$ | 1.01 (d, 3H, *J*=7.4Hz) | 11.9 | $CH_3$ |
| 19-$CH_3$ | 1.93 (s, 3H) | 7.5 | $CH_3$ |
| 1\`a | 1.40 (ddt, 1H, *J*=13.4, 10.6, 5.8Hz) | 33.3 | CH |
| 1\`b | 1.67 (ddd, 1H, *J*=8.9, 4.4, 3.0 Hz) | | |
| 2\` | 1.23 (m, 2H) | 29.9 | $CH_2$ |
| 3\` | 1.29 (m, 2H) | 22.6 | $CH_2$ |
| 4\` | 0.88 (t, 3H, *J*=7.2Hz) | 13.8 | $CH_2$ |
| 5\`a | 1.81 (dq, 1H, *J*=14.7, 7.6Hz) | 23.5 | $CH_2$ |
| 5\`b | 2.08 (tt, 1H, *J*=14.2, 7.4Hz) | | |
| 6\` | 0.94 (t, 3H, *J*=7.2Hz) | 11.9 | $CH_3$ |

$^1$H NMR: 600 MHz; $^{13}$C NMR: 150 MHz (in DMSO-$d_6$).

* The interconversion of the benzoquinone moiety led to the missing of these carbon signals.

Supplementary Table 23. NMR data of compound **19b**.

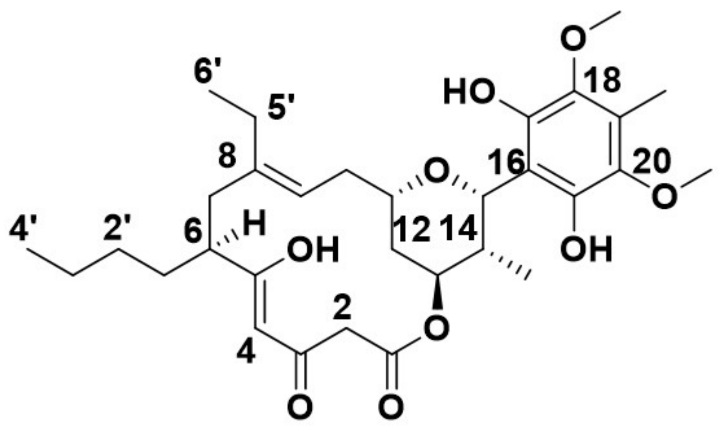

**(19b)**

| NO. | $\delta_H$ (mult., *J* in Hz) | $\delta_C$ | C-type |
|---|---|---|---|
| 1 | | 167.3 | C |
| 2a | 3.40 (d, 1H, *J*=14.0Hz) | 38.4 | $CH_2$ |
| 2b | 3.44 (d, 1H, *J*=14.0Hz) | | |
| 3 | | 191.4 | C |
| 4 | 5.45 (s, 1H) | 100.1 | CH |
| 5 | | 190.6 | C |
| 6 | 2.15 (overlapped, 1H) | 47.4 | CH |
| 7a | 2.14 (overlapped, 1H) | 38.4 | $CH_2$ |
| 7b | 2.27 (m, 1H) | | |
| 8 | | 141.6 | C |
| 9 | 5.04 (t, 1H, *J*=8.3Hz) | 121.2 | CH |
| 10a | 2.22 (m, 1H) | 34.2 | $CH_2$ |
| 10b | 2.31(m, 1H) | | |
| 11 | 3.38 (m, 1H) | 75.9 | CH |
| 12a | 1.68 (m, 1H) | 30.5 | $CH_2$ |
| 12b | 1.74 (m, 1H) | | |
| 13 | 5.17 (q, 1H, *J*=2.7Hz) | 72.7 | CH |
| 14 | 2.19 (m, 1H) | 35.4 | CH |
| 15 | 5.46 (d, 1H, *J*=2.8Hz) | 77.1 | CH |
| 16 | | 108.5 | C |
| 17 | | * | C |
| 18 | | 138.6 | C |
| 19 | | 123.6 | C |
| 20 | | 138.6 | C |
| 21 | | * | C |
| 14-$CH_3$ | 0.97 (d, 3H, *J*=7.4Hz) | 11.9 | $CH_3$ |
| 18-$OCH_3$ | 3.73 (s, 3H) | 60.6 | $CH_3$ |
| 19-$CH_3$ | 2.20 (s, 3H) | 9.4 | $CH_3$ |
| 20-$OCH_3$ | 3.73 (s, 3H) | 60.6 | $CH_3$ |
| 1`a | 1.41 (m, 1H) | 33.04 | $CH_2$ |
| 1`b | 1.69 (m, 1H) | | |
| 2` | 1.25 (overlapped, 2H) | 27.5 | $CH_2$ |
| 3` | 1.30 (overlapped, 2H) | 22.8 | $CH_2$ |
| 4` | 0.88 (t, 3H, *J*=6.8Hz) | 13.8 | $CH_3$ |
| 5`a | 1.83 (m, 1H) | 23.5 | $CH_2$ |
| 5`b | 2.09 (m, 1H) | | |
| 6` | 0.94 (t, 3H, *J*=7.6Hz) | 11.9 | $CH_3$ |

$^1$H NMR: 600 MHz; $^{13}$C NMR: 150 MHz (in DMSO-$d_6$).

* The interconversion of the benzoquinone moiety led to the missing of these carbon signals.

Supplementary Table 24. NMR data of compound **19c**.

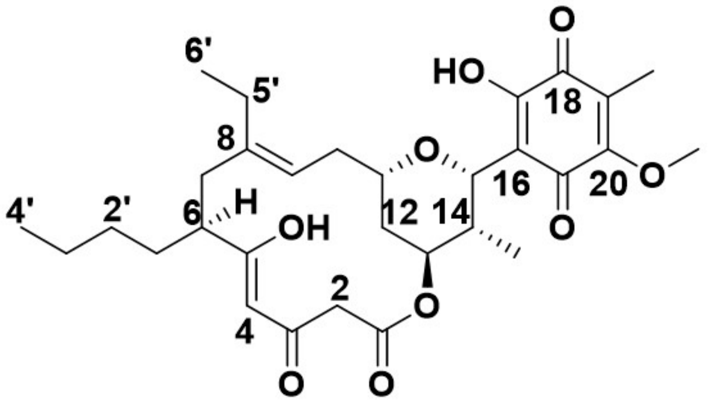

**(19c)**

| NO. | $\delta_H$ (mult., *J* in Hz) | $\delta_C$ | C-type |
|---|---|---|---|
| 1 | | 166.5 | C |
| 2 | 3.45 (s, 2H) | 48.4 | $CH_2$ |
| 3 | | 191.00 | C |
| 4 | 5.45 (s, 1H) | 99.9 | CH |
| 5 | | 190.5 | C |
| 6 | 2.15 (overlapped, 1H) | 47.4 | CH |
| 7a | 2.16 (overlapped, 1H) | 38.4 | $CH_2$ |
| 7b | 2.29 (t, 1H, *J*=13.2Hz) | | |
| 8 | | 141.8 | C |
| 9 | 5.03 (t, 1H, *J*=8.3Hz) | 120.6 | CH |
| 10 | 2.25 (m, 2H) | 33.9 | $CH_2$ |
| 11 | 3.37 (m, 1H) | 76.1 | CH |
| 12a | 1.64 (m, 1H) | 30.4 | $CH_2$ |
| 12b | 1.76 (m, 1H) | | |
| 13 | 5.16 (q, 1H, *J*=2.7Hz) | 72.0 | CH |
| 14 | 2.20 (m, 1H) | 35.3 | CH |
| 15 | 5.23 (d, 1H, *J*=2.7Hz) | 76.9 | CH |
| 16 | | 114.3 | C |
| 17 | | 155.3 | C |
| 18 | | 183.2 | C |
| 19 | | 125.5 | C |
| 20 | | 155.4 | C |
| 21 | | 181.7 | C |
| 14-$CH_3$ | 1.03 (d, 3H, *J*=7.2Hz) | 11.4 | $CH_3$ |
| 19-$CH_3$ | 1.96 (s, 3H) | 8.3 | $CH_3$ |
| 20-OCH3 | 4.03 (s, 3H) | 61.2 | $CH_3$ |
| 1\`a | 1.43 (m, 1H) | 33.1 | $CH_2$ |
| 1\`b | 1.70 (m, 1H) | | |
| 2\` | 1.27 (m, 2H) | 29.8 | $CH_2$ |
| 3\` | 1.32 (m, 2H) | 22.6 | $CH_2$ |
| 4\` | 0.90 (t, 3H, *J*=7.2Hz) | 14.0 | $CH_3$ |
| 5\`a | 1.84 (m, 1H) | 23.5 | $CH_2$ |
| 5\`b | 2.10 (m, 1H) | | |
| 6\` | 0.96 (t, 3H, *J*=7.5Hz) | 13.0 | $CH_3$ |

$^{1}$H NMR: 600 MHz; $^{13}$C NMR: 150 MHz (in DMSO-$d_6$).

Supplementary Table 25. NMR data of compound **21**.

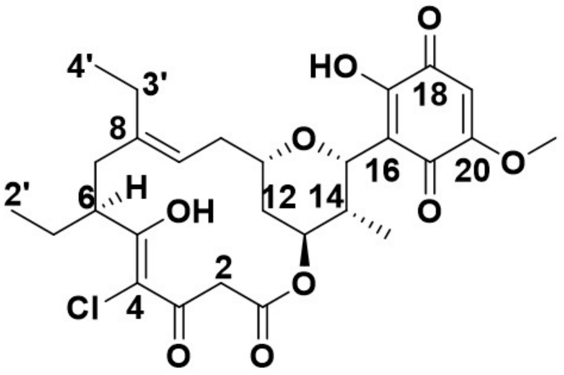

(21)

| NO. | $\delta_H$ (mult., *J* in Hz) | $\delta_C$ | C-type |
|---|---|---|---|
| 1 | | 165.1 | C |
| 2a | 3.53(d, 1H, *J*=15.6Hz) | 47.0 | $CH_2$ |
| 2b | 3.53(d, 1H, *J*=15.6Hz) | | |
| 3 | | 189.6 | C |
| 4 | | 109.2 | C |
| 5 | | 188.8 | C |
| 6 | 3.19 (m, 1H) | 43.8 | CH |
| 7a | 2.23 (overlapped, 1H) | 37.8 | $CH_2$ |
| 7b | 2.36 (t, 1H, *J*=12.7Hz) | | |
| 8 | | 141.3 | C |
| 9 | 5.10 (t, 1H, *J*=8.3Hz) | 120.9 | CH |
| 10 | 2.20 (overlapped, 1H) | 33.4 | $CH_2$ |
| 11 | 3.35 (m, 1H) | 76.6 | CH |
| 12 | 1.67 (m, 1H) | 30.8 | $CH_2$ |
| 13 | 5.17 (brs, 1H) | 72.5 | CH |
| 14 | 2.17 (m, 1H) | 34.9 | CH |
| 15 | 5.16 (brs, 1H) | 77.4 | CH |
| 16 | | 114.9 | C |
| 17 | | 156.0 | C |
| 18 | | * | C |
| 19 | 5.80 (s, 1H) | 104.5 | CH |
| 20 | | 158.7 | C |
| 21 | | 180.0 | C |
| 14-$CH_3$ | 0.99 (overlapped, 3H) | 11.4 | $CH_3$ |
| 20-$OCH_3$ | 3.80 (s, 3H) | 56.3 | |
| 1`a | 1.53 (m, 1H) | 27.0 | $CH_2$ |
| 1`b | 1.69 (m, 1H) | | |
| 2` | 0.94 (t, 3H, *J*=7.4 Hz) | 11.6 | $CH_3$ |
| 3` | 1.88 (m, 1H) | 23.0 | $CH_2$ |
| 4` | 0.98 (overlapped, 3H) | 12.8 | $CH_3$ |

$^1$H NMR: 600 MHz; $^{13}$C NMR: 150 MHz (in DMSO-$d_6$).

* The interconversion of the benzoquinone moiety led to the missing of these carbon signals.

Supplementary Table 26. Information on the compounds obtained in this study.

| Strain | Compound | Production Yield | Construction method |
|---|---|---|---|
| Wild-type (*S.cinnamoneus*) | **1** | ~50mg/L | |
| | **1b** | ~2mg/L | |
| | **1c** | ~1mg/L | |
| | **2** | ~10mg/L | |
| ΔcmmB | **3** | ~50mg/L | *cmmB* in-frame deletion mutant[2] |
| | **4** | ~10mg/L | |
| WT+mgmO | **5** | ~10mg/L | *mgmO* gene integrated strain of WT |
| | **6** | ~2mg/L | |
| ΔcmmB+mgmO | **7** | ~10mg/L | *mgmO* gene integrated strain of ΔcmmB mutant |
| | **8** | ~2mg/L | |
| S1 | **9** | ~50mg/L | Cmm-$AT_1$ replacement mutant with Cmm-$AT_4$ in WT |
| | **10** | ~15mg/L | |
| S2 | **11** | ~30mg/L | Cmm-$AT_4$ replacement mutant with Mgm-$AT_5$ in WT |
| | **12** | ~7mg/L | |
| S3 | **13a** | ~5mg/L | Cmm-AT5 replacement mutant with Mgm-$AT_5$ in WT |
| | **13b** | ~0.5mg/L | |
| | **14a** | ~3mg/L | |
| | **15a** | ~2mg/L | |
| S4 | **16** | ~15mg/L | Cmm-$AT_4$ replacement mutant with Mgm-$AT_5$ in S3 |
| | **17** | ~3mg/L | |
| S5 | **18a** | ~2mg/L | Cmm-AT5 replacement mutant with Mgm-$AT_5$ in S4 |
| | **18b** | ~1.5mg/L | |
| | **18c** | ~0.5mg/L | |
| | **19a** | ~4mg/L | |
| | **19b** | ~5mg/L | |
| | **19c** | ~5mg/L | |
| S5-mgm$KS_5$ | **18a** | ~1.5mg/L | Cmm-$KS_5$ replacement mutant with Mgm-$KS_5$ in S5 |
| | **18b** | ~4mg/L | |
| | **18c** | ~4mg/L | |
| S6 | **20** | Trace | *cmmB* in-frame deletion mutant in S5-mgm$KS_5$ |
| S7 | **21** | ~0.5mg/L | CCR-HCD-mgmO co-overexpression in mutant S6 |

### Supplementary Figure

Compare these results against the new Clustered nr database BLAST

Descriptions | Graphic Summary | Alignments | Taxonomy

**Sequences producing significant alignments** Download Select columns Show 100

select all *100 sequences selected* GenPept Graphics Distance tree of results Multiple alignment MSA Viewer

| Description | Scientific Name | Max Score | Total Score | Query Cover | E value | Per. Ident | Acc. Len | Accession |
|---|---|---|---|---|---|---|---|---|
| type I polyketide synthase [Streptomyces cinnamoneus] | Streptomyces ci... | 848 | 1411 | 100% | 0.0 | 100.00% | 3925 | WP_099197735.1 |
| type I polyketide synthase [Streptomyces mangrovisoli] | Streptomyces m... | 680 | 1241 | 100% | 0.0 | 78.64% | 3992 | WP_046583961.1 |
| type I polyketide synthase [Streptomyces cinnamoneus] | Streptomyces ci... | 672 | 1893 | 100% | 0.0 | 80.14% | 5475 | WP_240003137.1 |
| type I polyketide synthase [Streptomyces cinnamoneus] | Streptomyces ci... | 671 | 1892 | 100% | 0.0 | 80.14% | 5475 | WP_243469264.1 |
| SDR family NAD(P)-dependent oxidoreductase [Streptomyces sp. NBC_00659] | Streptomyces s... | 657 | 1200 | 100% | 0.0 | 76.06% | 3919 | WP_329297130.1 |
| SDR family NAD(P)-dependent oxidoreductase [Streptomyces sp. NBC_01393] | Streptomyces s... | 653 | 1200 | 100% | 0.0 | 76.48% | 3918 | WTZ06591.1 |
| type I polyketide synthase [Streptomyces hygroscopicus] | Streptomyces h... | 649 | 3043 | 100% | 0.0 | 74.30% | 8595 | WP_236256936.1 |
| type I polyketide synthase [Streptomyces hygroscopicus] | Streptomyces h... | 649 | 3051 | 100% | 0.0 | 74.30% | 8606 | MEU5273606.1 |
| type I polyketide synthase [Streptomyces hygroscopicus] | Streptomyces h... | 649 | 3037 | 100% | 0.0 | 74.30% | 8605 | MEU1908929.1 |
| type I polyketide synthase [Saccharothrix saharensis] | Saccharothrix s... | 648 | 1271 | 100% | 0.0 | 75.23% | 4484 | WP_141974657.1 |
| type I polyketide synthase [Streptomyces sp. 5-10] | Streptomyces s... | 647 | 647 | 100% | 0.0 | 74.07% | 1620 | WP_191070427.1 |
| type I polyketide synthase [Streptomyces hygroscopicus] | Streptomyces h... | 647 | 3038 | 100% | 0.0 | 74.07% | 8582 | MEU7368327.1 |

Supplementary Fig. 1. BLAST search results using CmmD2-$KS_5$ as the probe.

MgmD2-$KS_5$ are circled in red square.

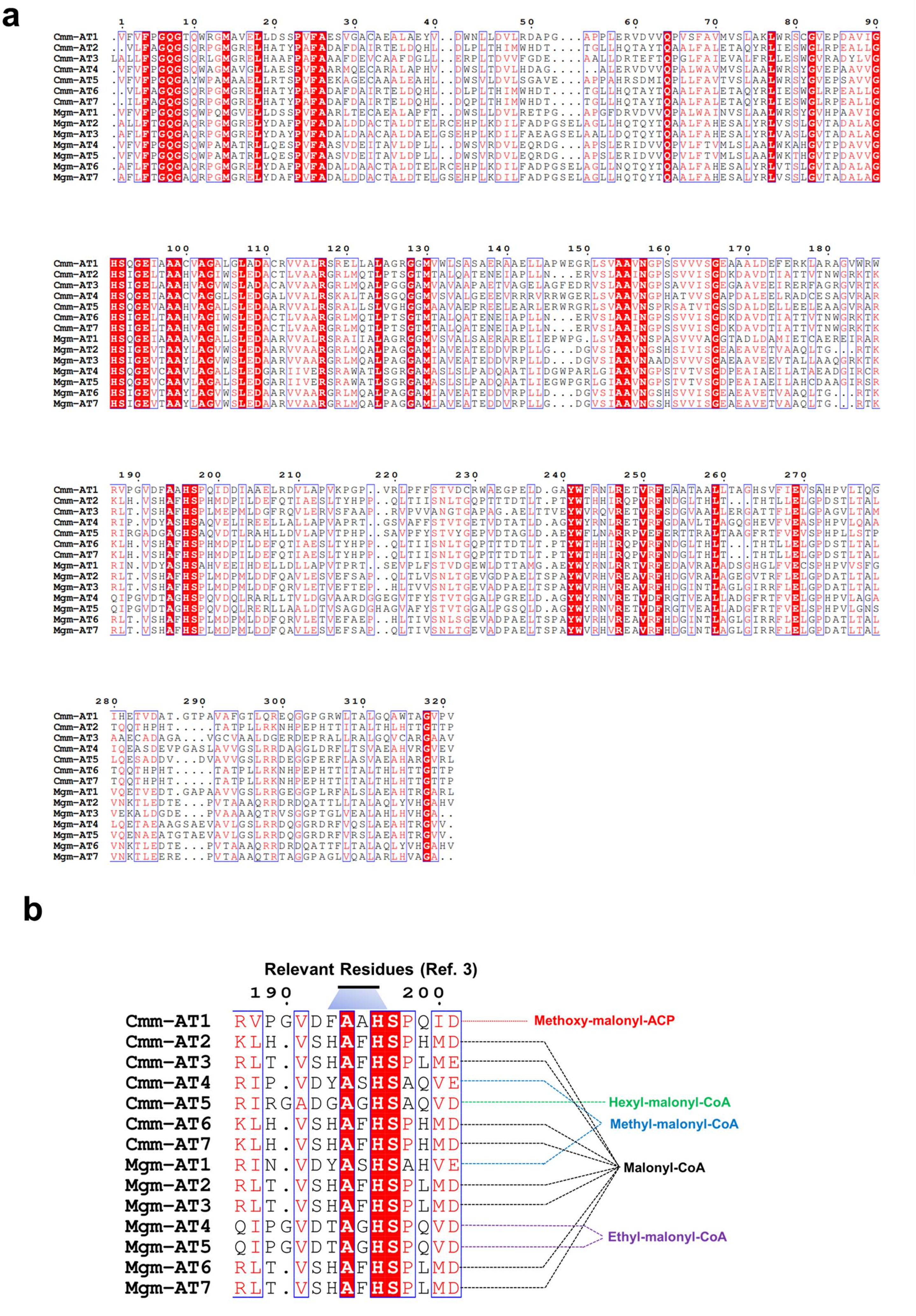

Supplementary Fig. 2. Amino acid sequence alignments of AT domains between *cmm* and *mgm* BGCs. (**a**) AT domains. (**b**) The substrate-specific motif[3], covering residues 193-196, in AT domains. The corresponding extender unit of each AT is labeled.

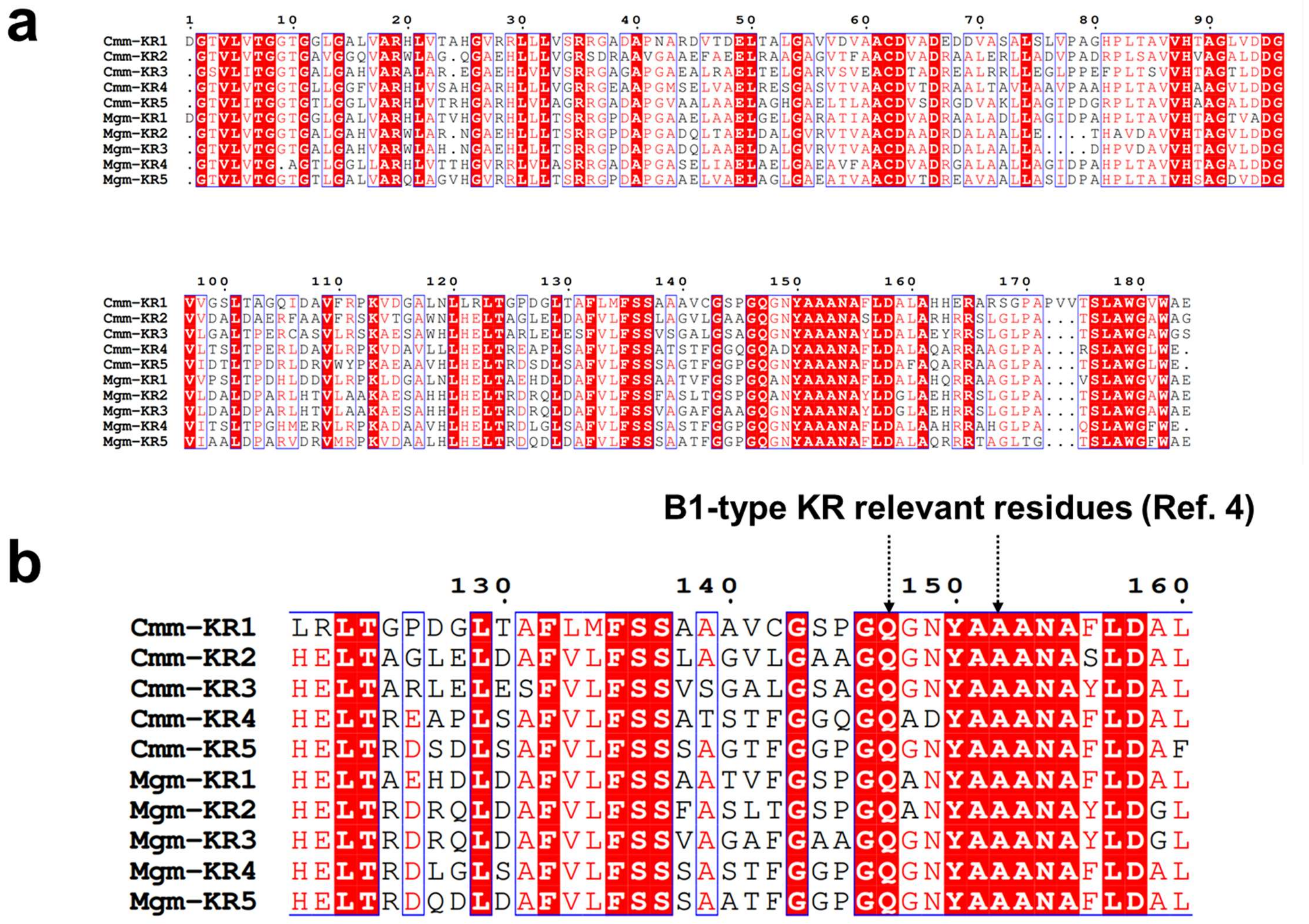

Supplementary Fig. 3. Amino acid sequence alignments of KR domains between *cmm* and *mgm* BGCs. (**a**) KR domains. (**b**) Comparison of the signature amino acids of B1-type KRs[4].

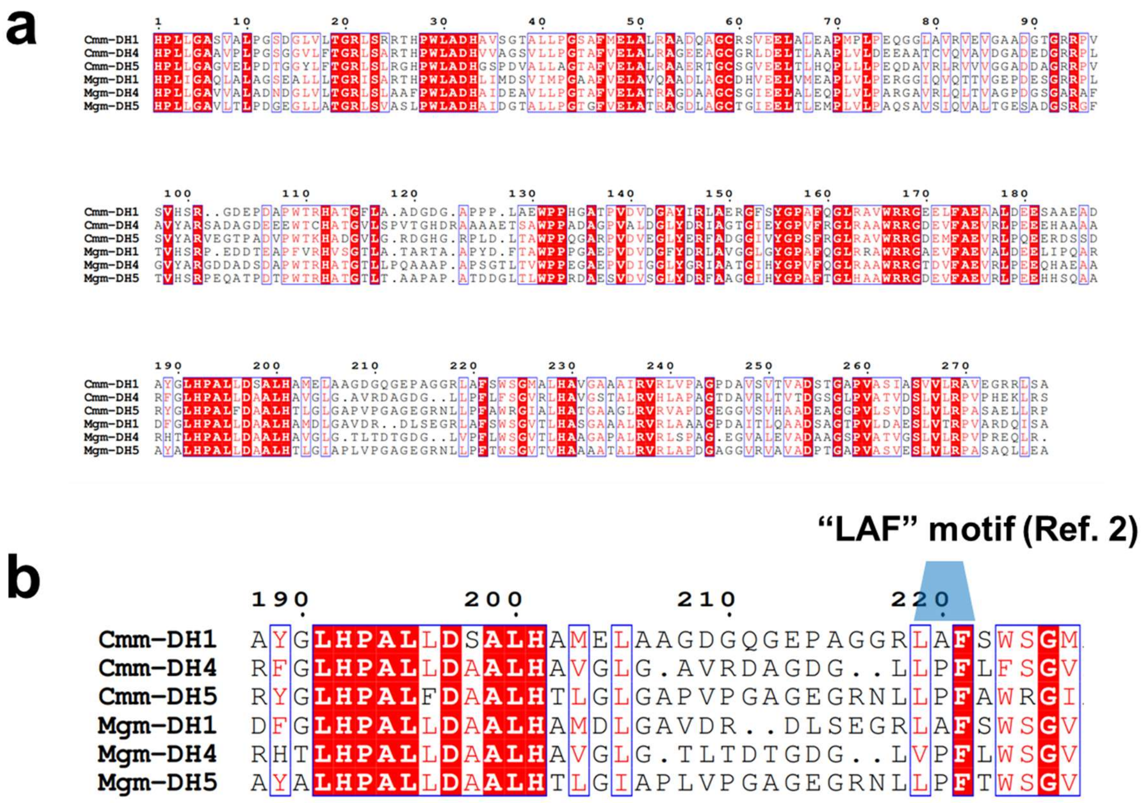

Supplementary Fig. 4. Amino acid sequence alignments of DH domains between *cmm* and *mgm* BGCs. (**a**) DH domains. (**b**) Comparison of the catalytically inactive signature motif “LAF” of Cmm-DH1 with Mgm-DH1.

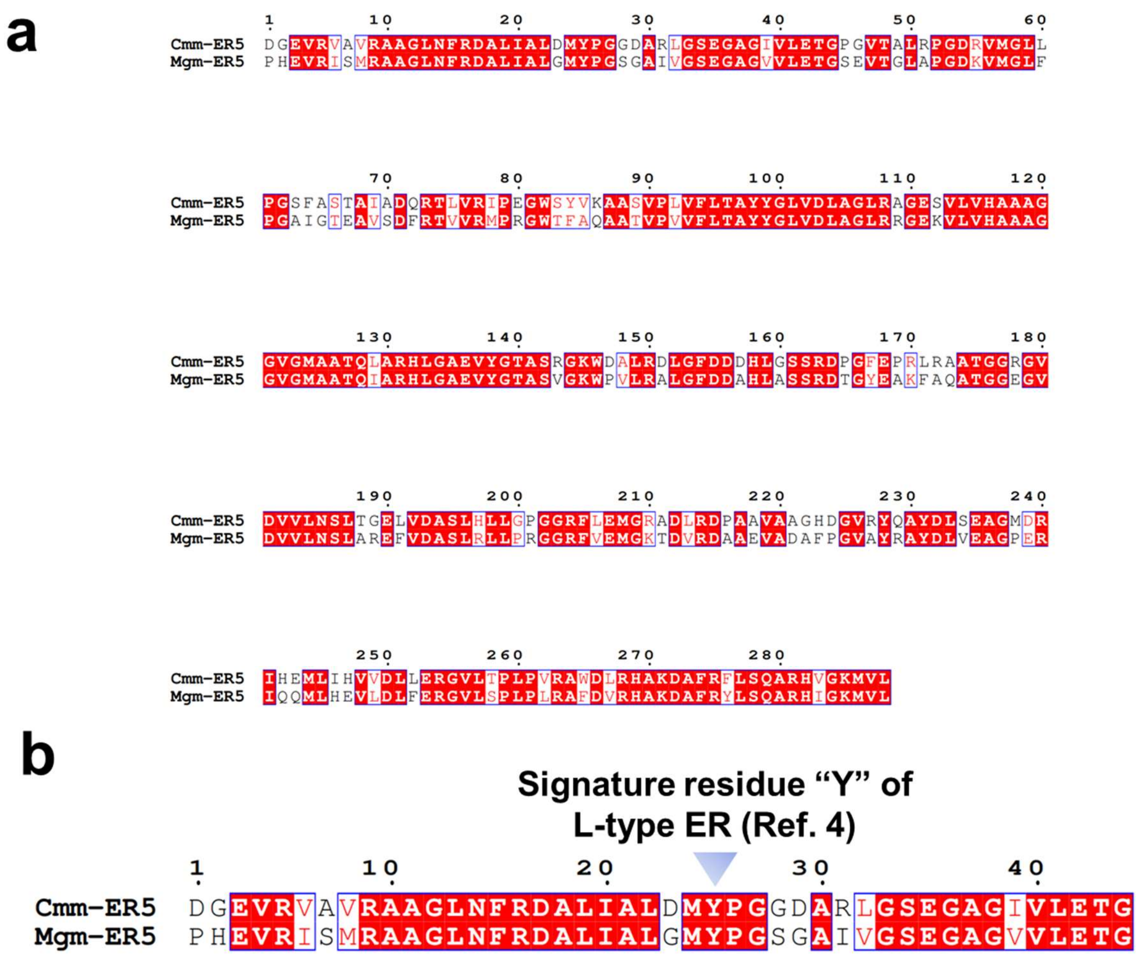

Supplementary Fig. 5. Amino acid sequence alignments of ER domains between *cmm* and *mgm* BGCs. (**a**) ER domains. (**b**) Comparison of the signature amino acid “Y” of L-type ER[4].

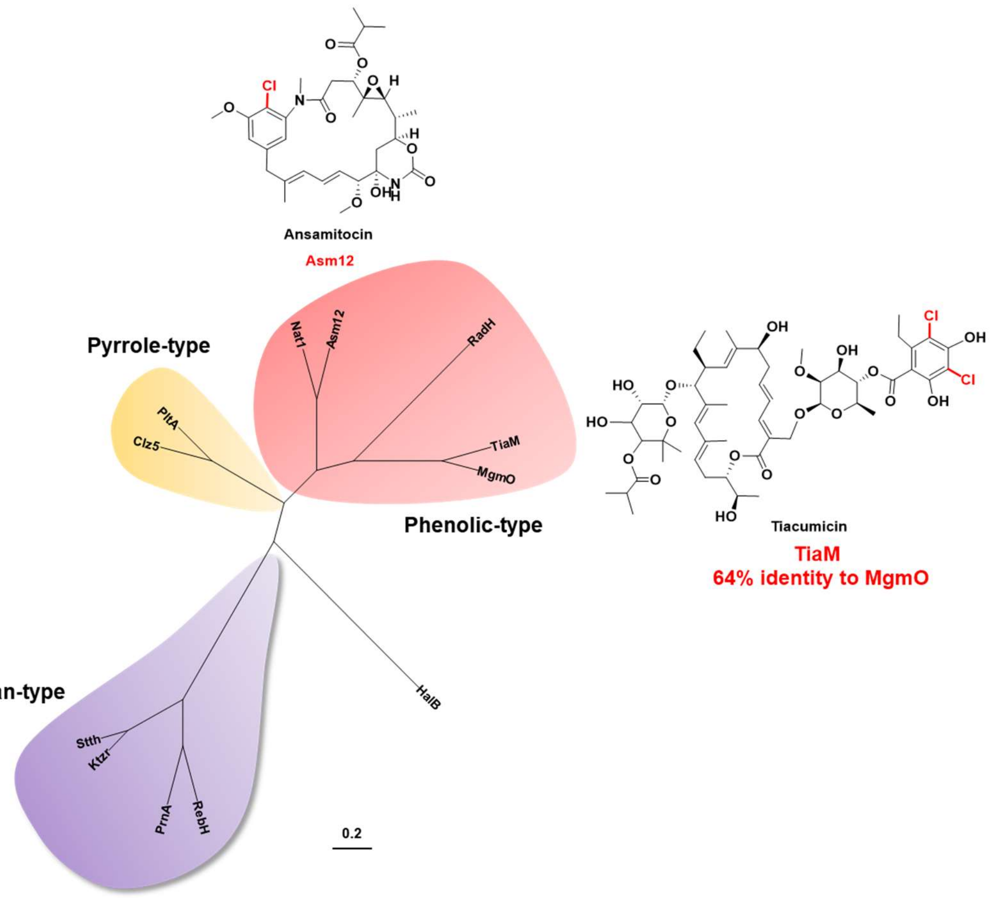

Supplementary Fig. 6. Phylogenetic tree analysis of FAD-dependent halogenases[5].

MgmO (WP_046591495.1); TiaM (ADU85999.1); RadH (C5H881.1); Asm12 (AAM54090.1); Nat1 (ADM46362.1); PltA (Q4KCZ0.1); Clz5(A0A345BJN5.1); SttH (E9P162.1); Ktzr (A8CF74.2); PrnA (P95480.1); RebH (Q8KHZ8.1); HalB (AAQ04685.1).

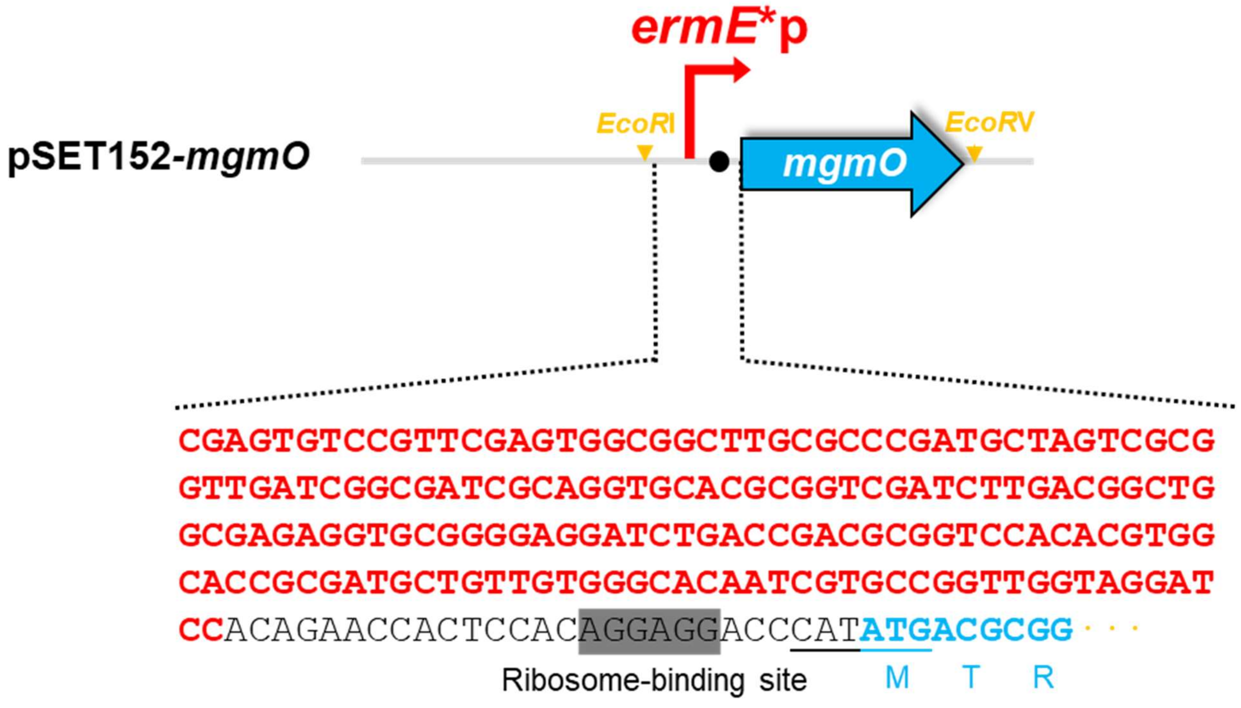

Supplementary Fig. 7. Nucleotide sequence of plasmid pSET1252-mgmO.

The "*ermE**p" nucleotide sequence is displayed in red bold font.

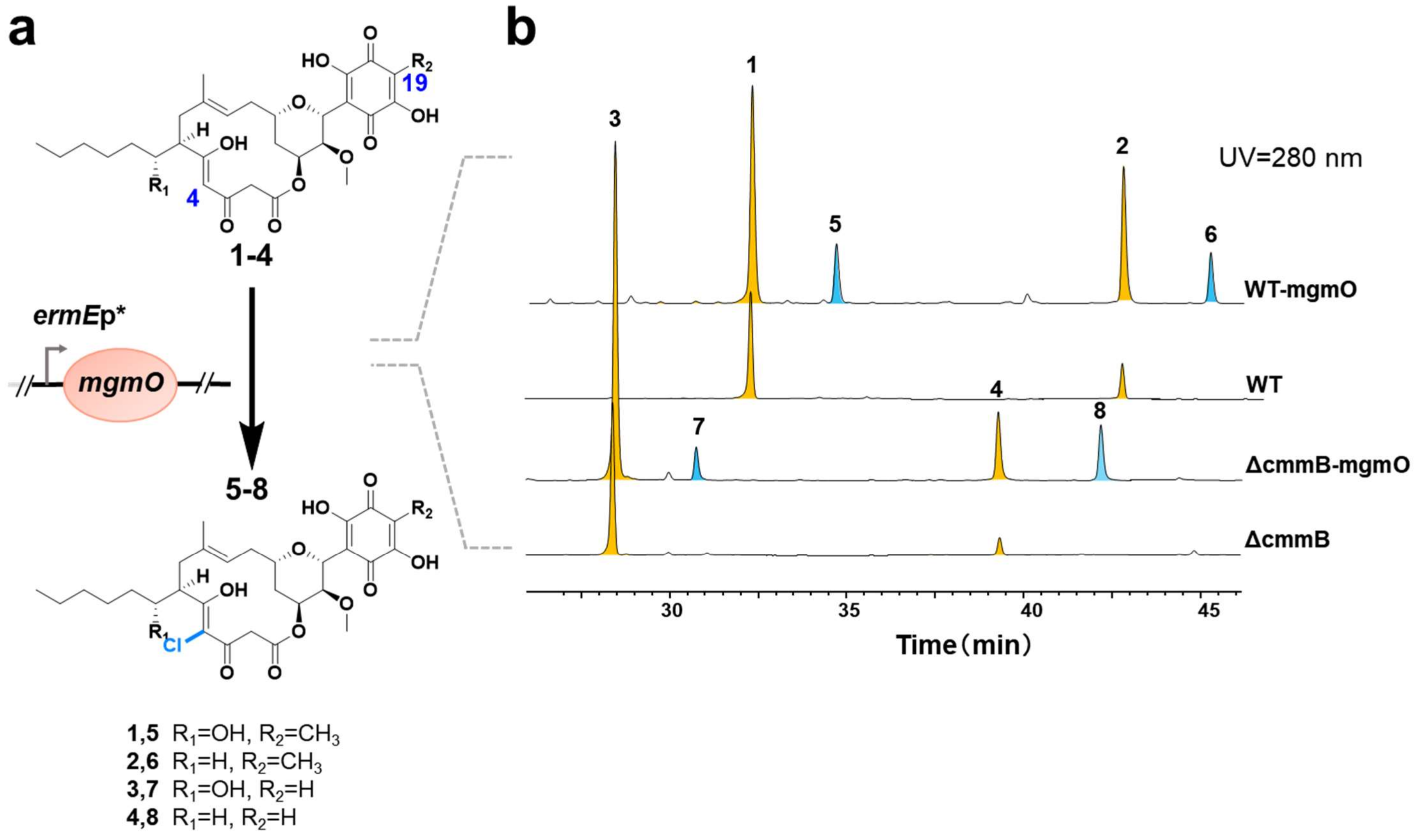

Supplementary Fig. 8. Olefin chlorination catalyzed by MgmO.

**a.** Cinnamomycin **1-4** converted to chlorinated-cinnamomycin **5-8** by MgmO. *ermE**p: the promoter used.

**b.** HPLC analyses of mutant strains after fermentation. Cinnamomycin **1-4** are colored as yellow. Chlorinated cinnamomycin **5-8** are colored as blue.

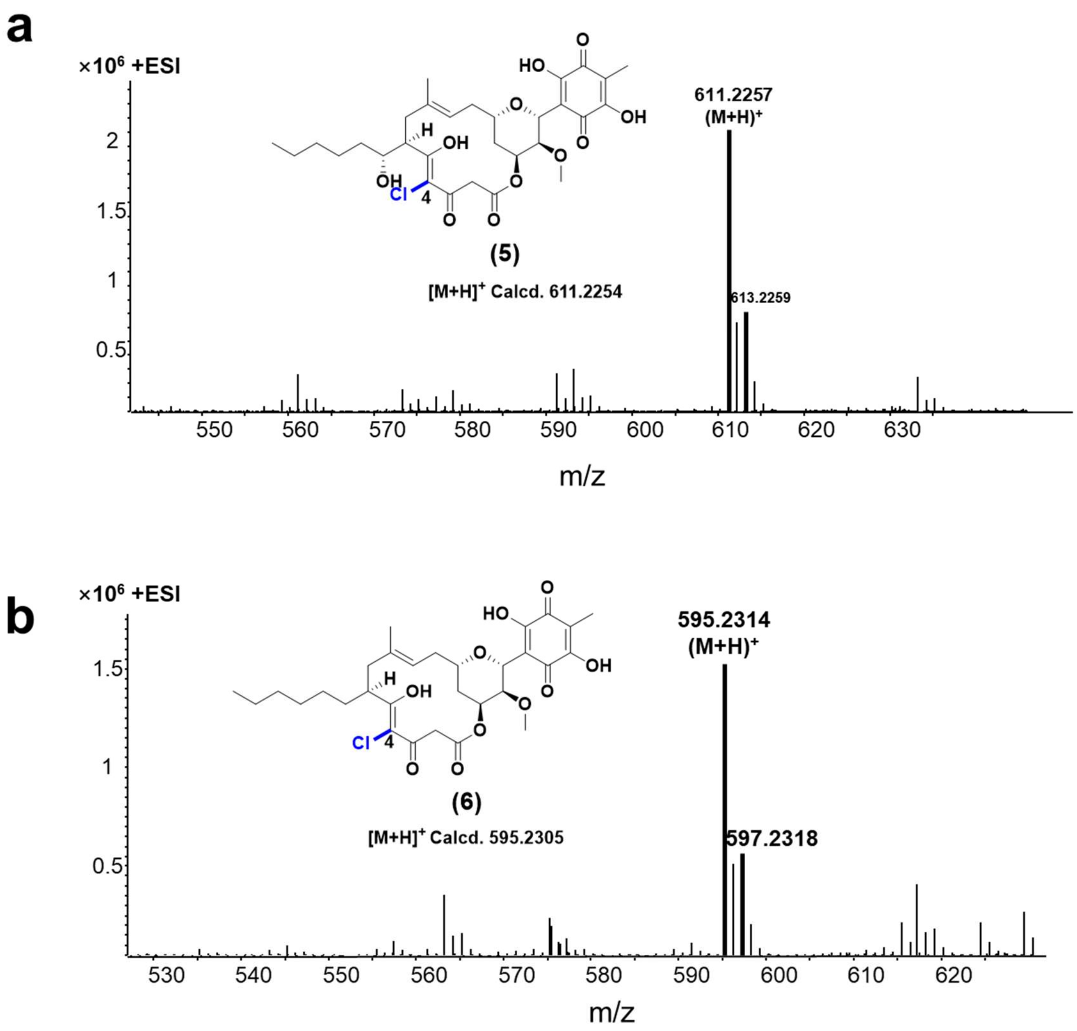

Supplementary Fig. 9. HR-ESI-MS spectra of chlorinated derivates **5** (a) and **6** (b).

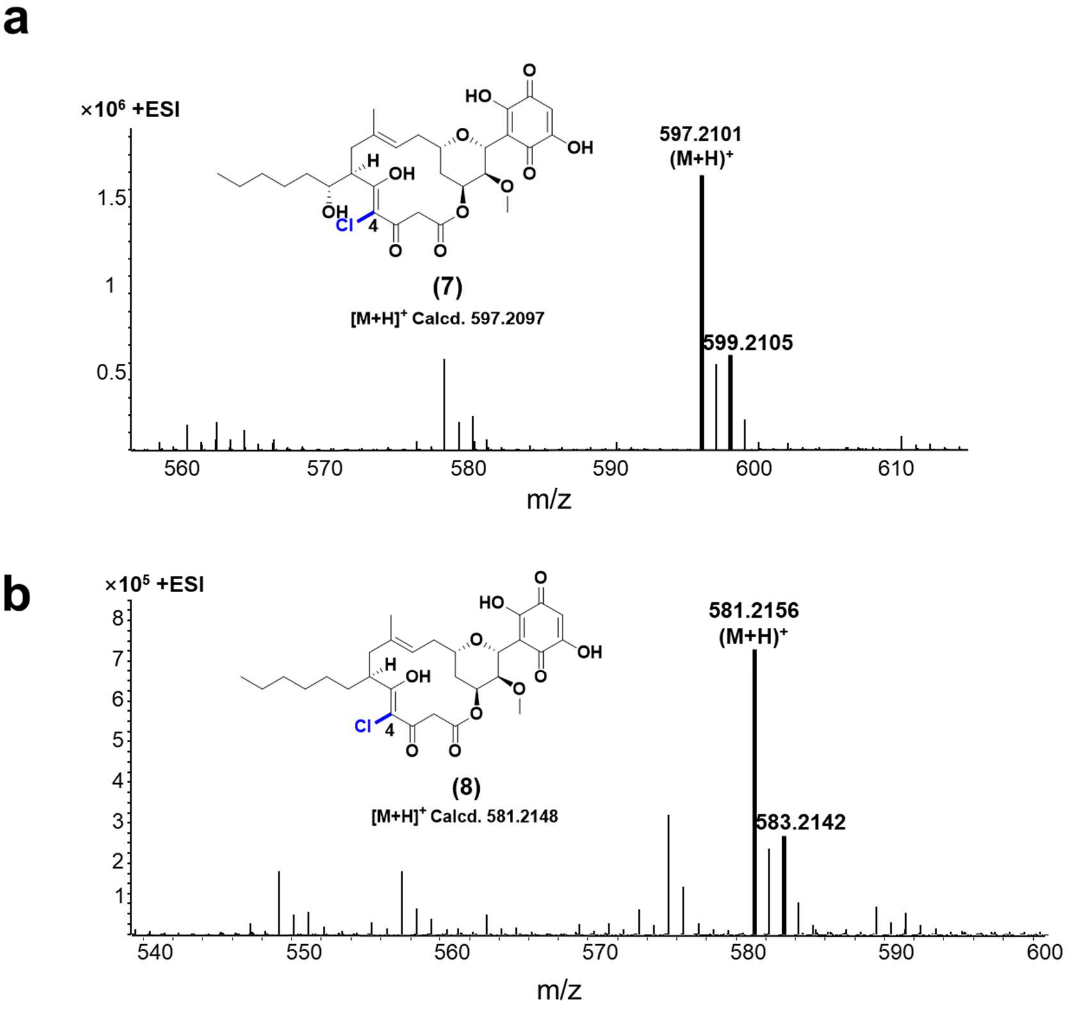

Supplementary Fig. 10. HR-ESI-MS spectra of chlorinated derivates **7** (a) and **8** (b).

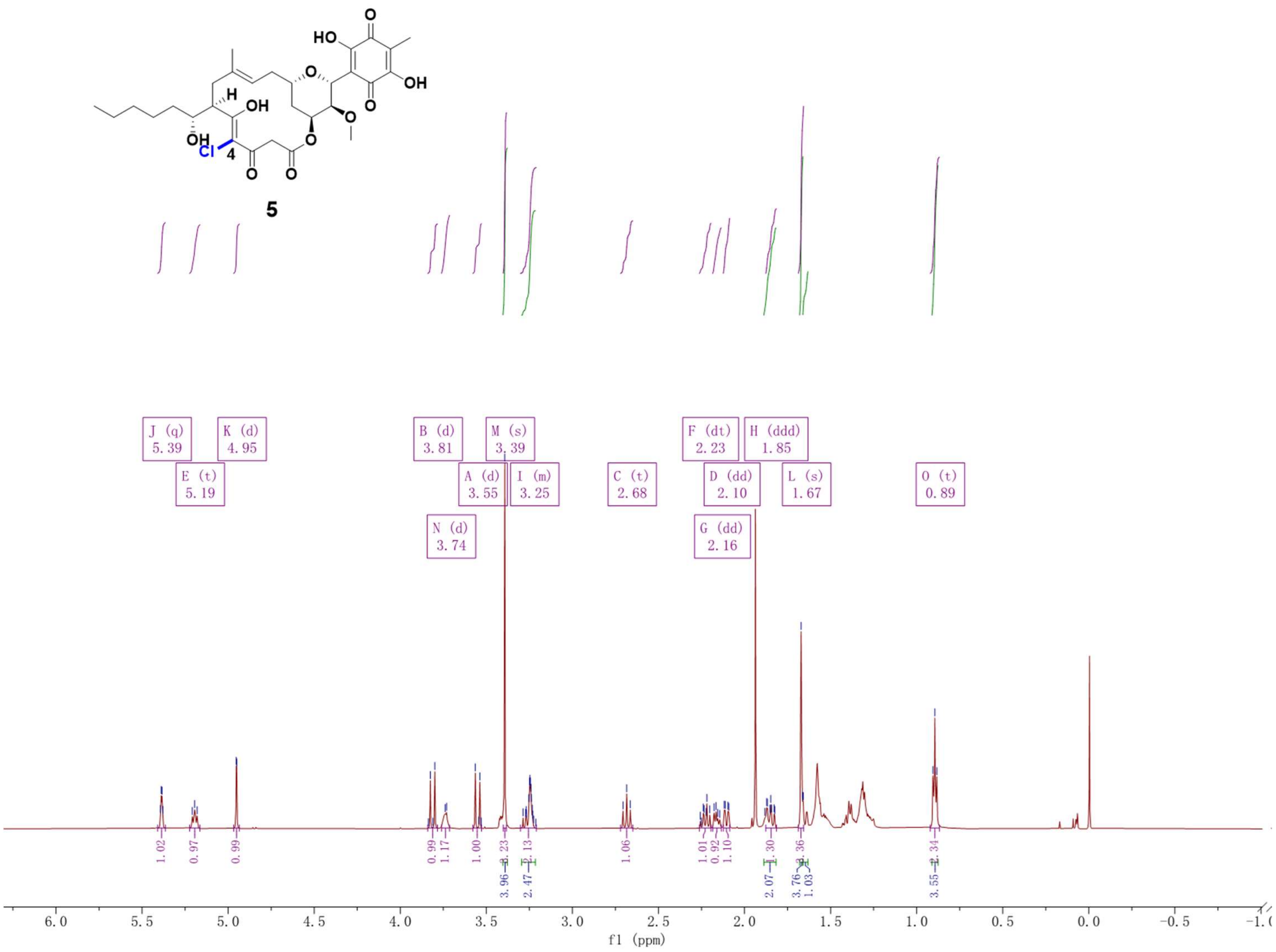

Supplementary Fig. 11. $^{1}$H NMR spectrum (600 MHz, $CDCl_3$) of compound **5**.

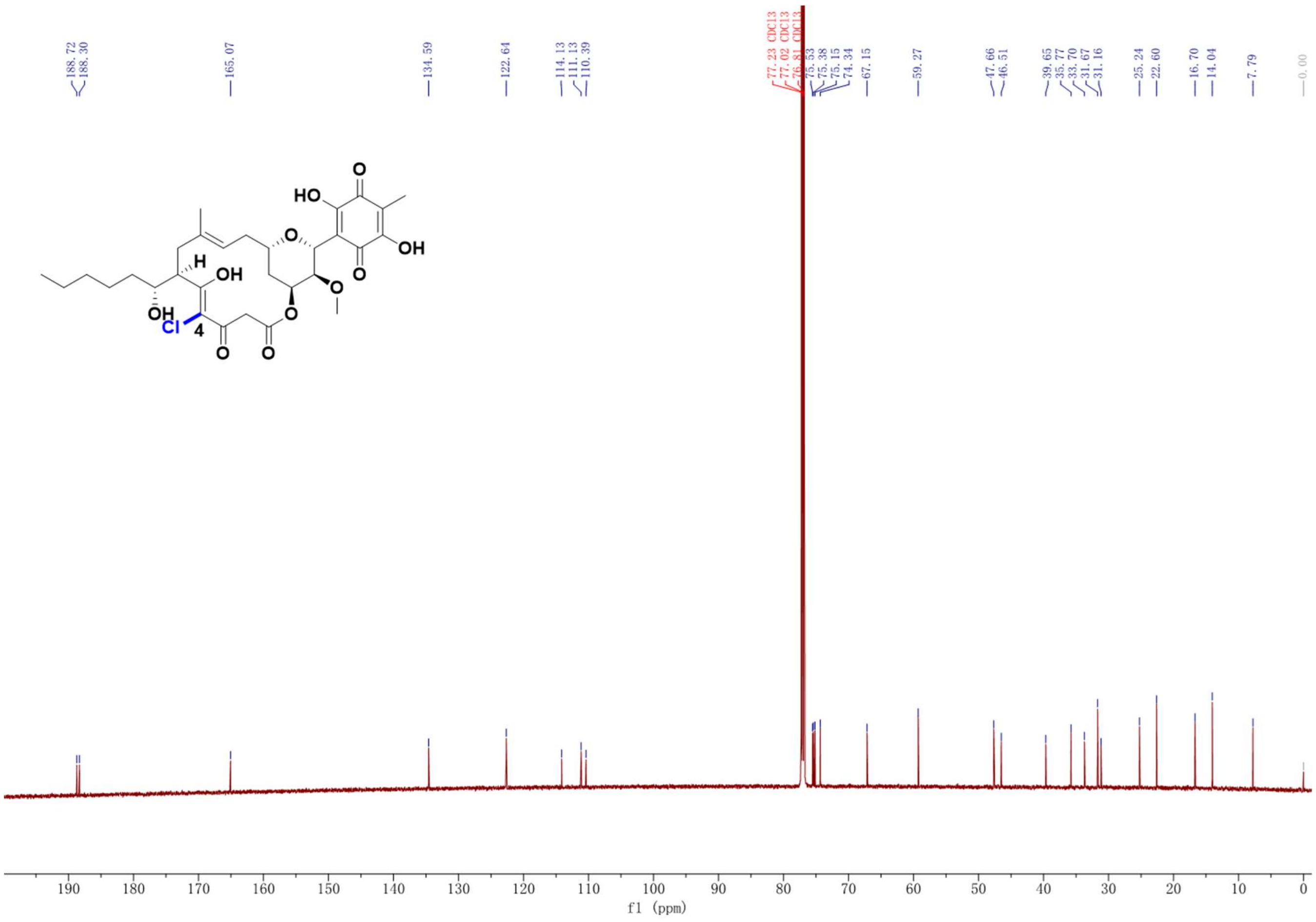

Supplementary Fig. 12. $^{13}$C NMR spectrum (150 MHz, $CDCl_3$) of compound **5**.

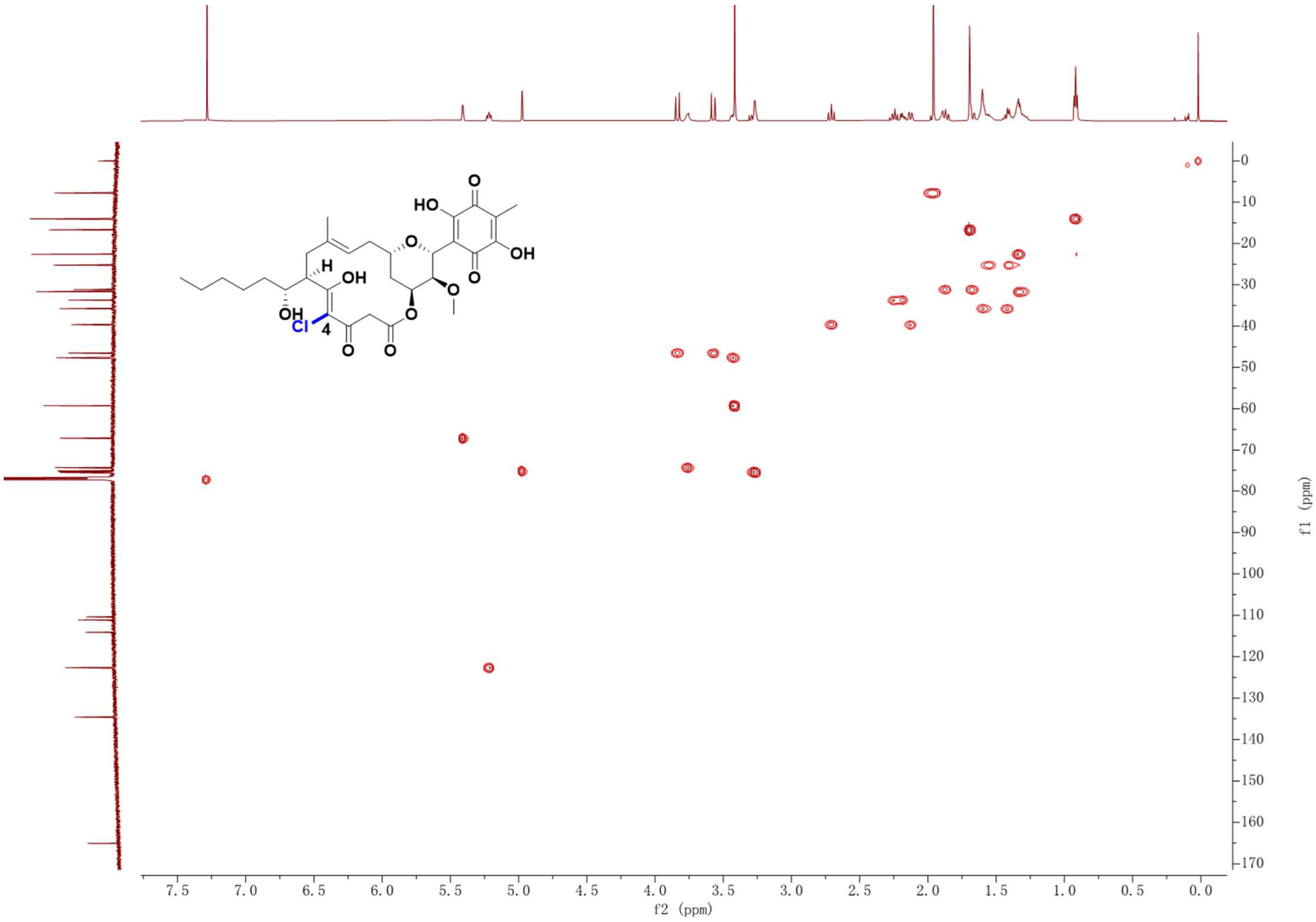

Supplementary Fig. 13. HSQC NMR spectrum (600 MHz, $CDCl_3$) of compound **5**.

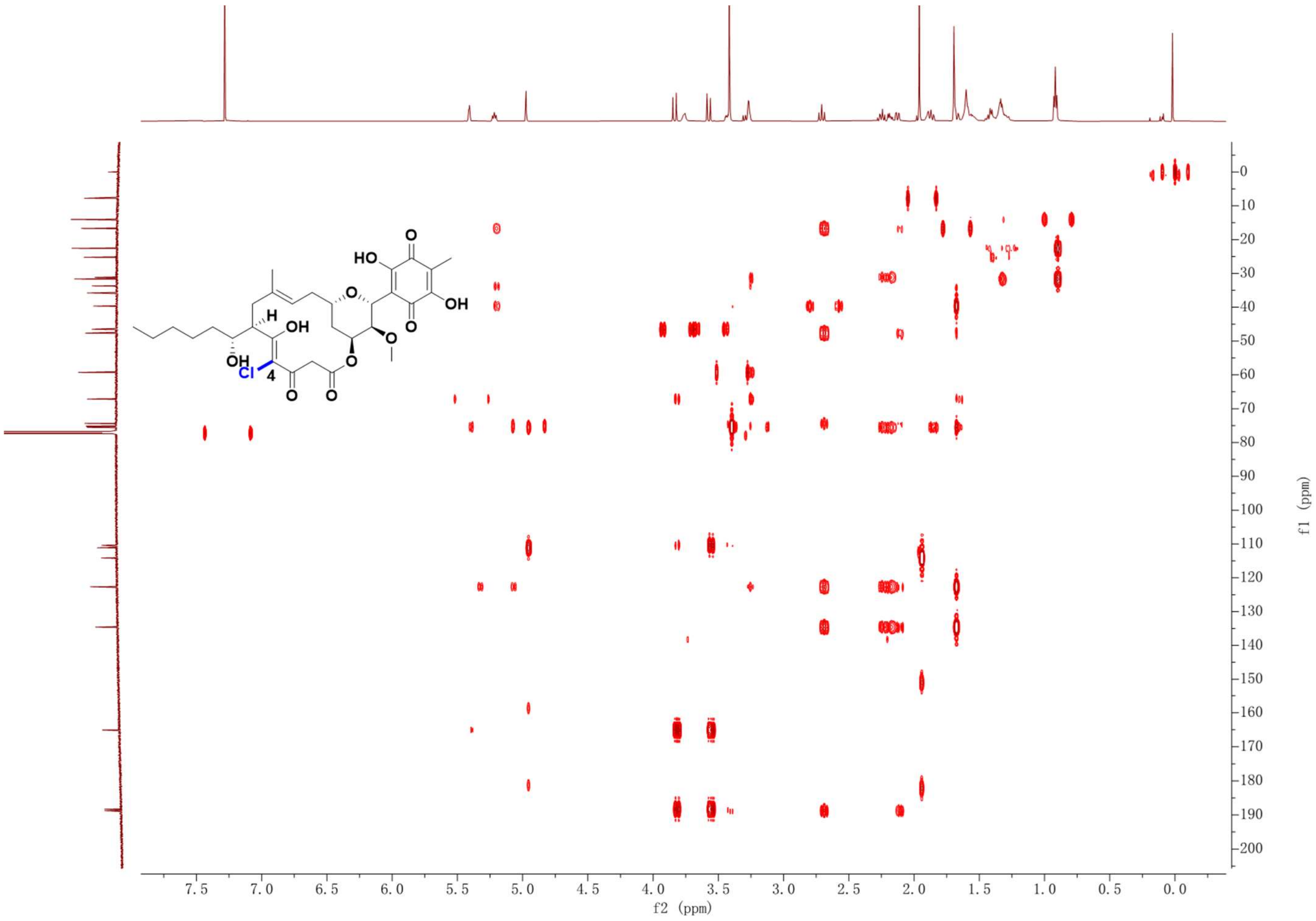

Supplementary Fig. 14. HMBC NMR spectrum (600 MHz, $CDCl_3$) of compound **5**.

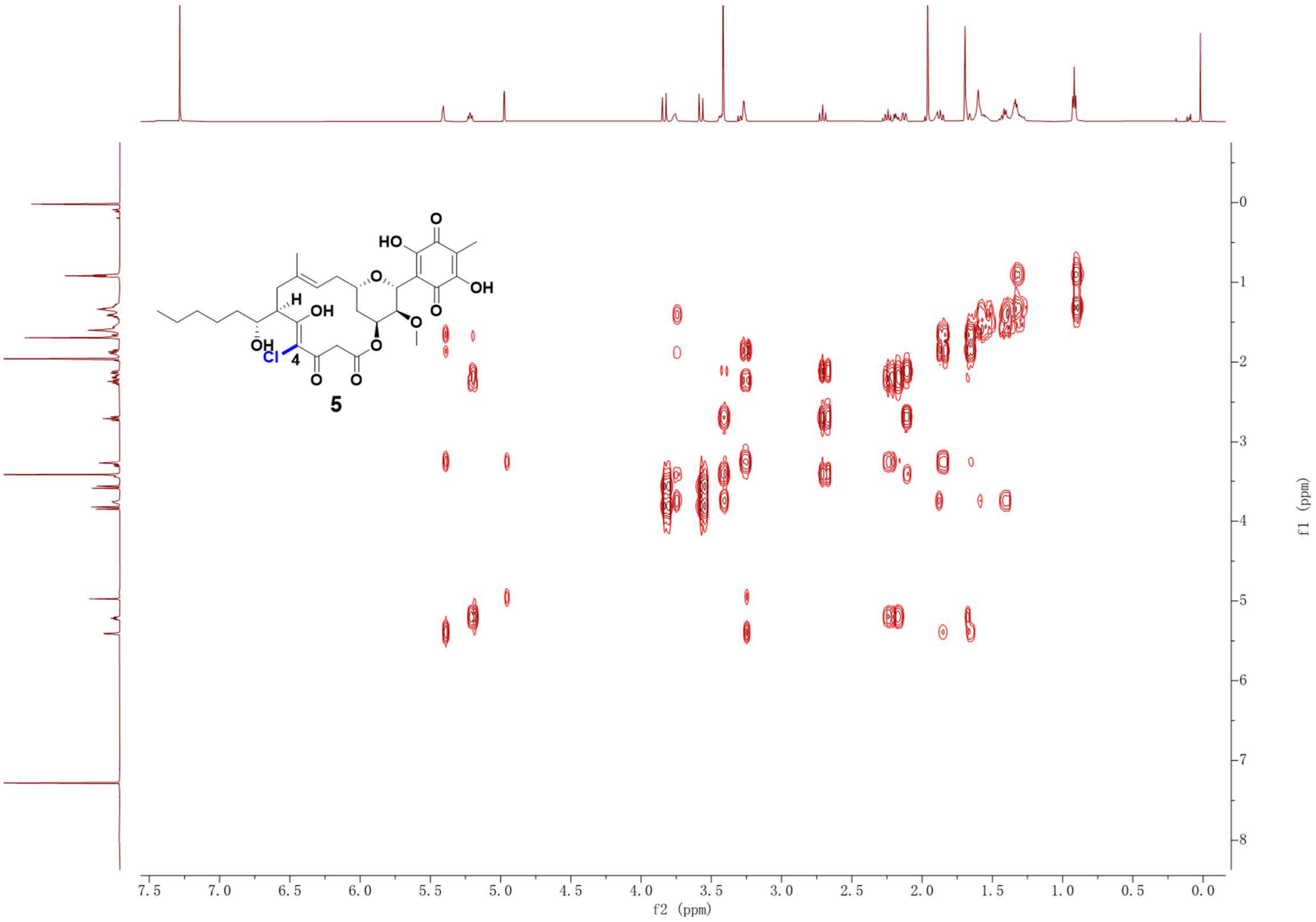

Supplementary Fig. 15. $^{1}H$-$^{1}H$ COSY NMR spectrum (600 MHz, $CDCl_3$) of compound **5**.

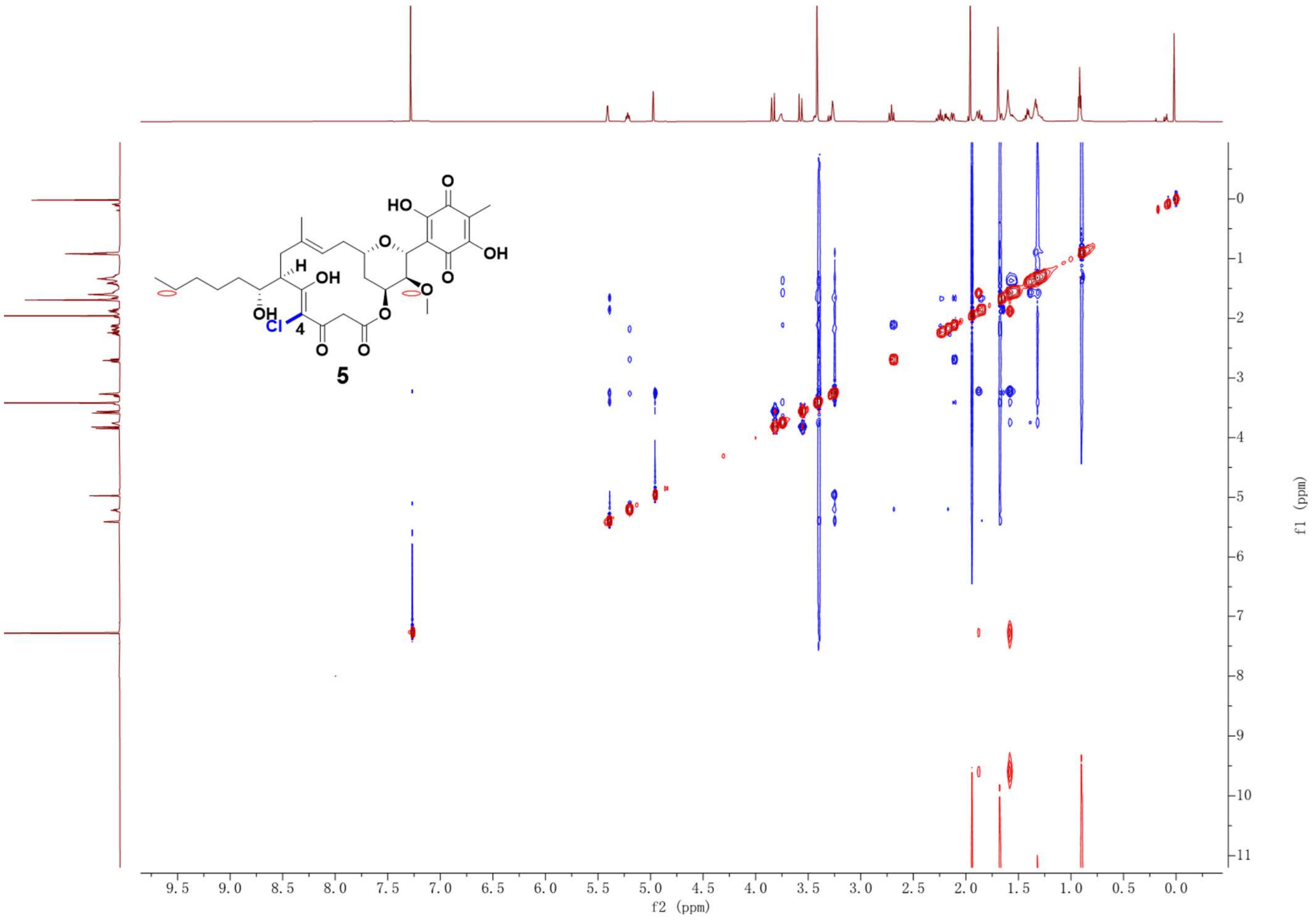

Supplementary Fig. 16. NOESY NMR spectrum (600 MHz, $CDCl_3$) of compound **5**.

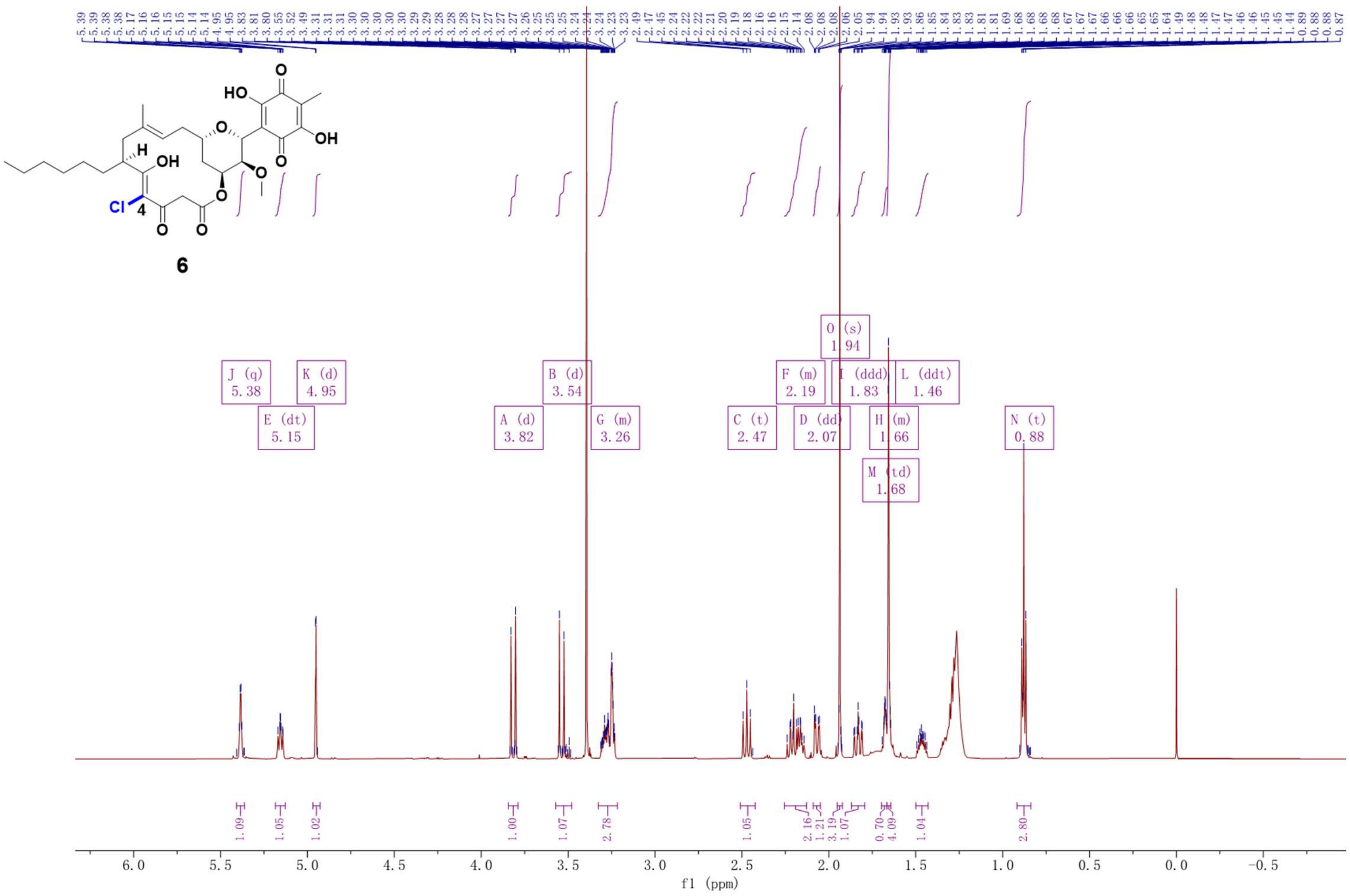

Supplementary Fig. 17. $^{1}$H NMR spectrum (600 MHz, $CDCl_3$) of compound **6**.

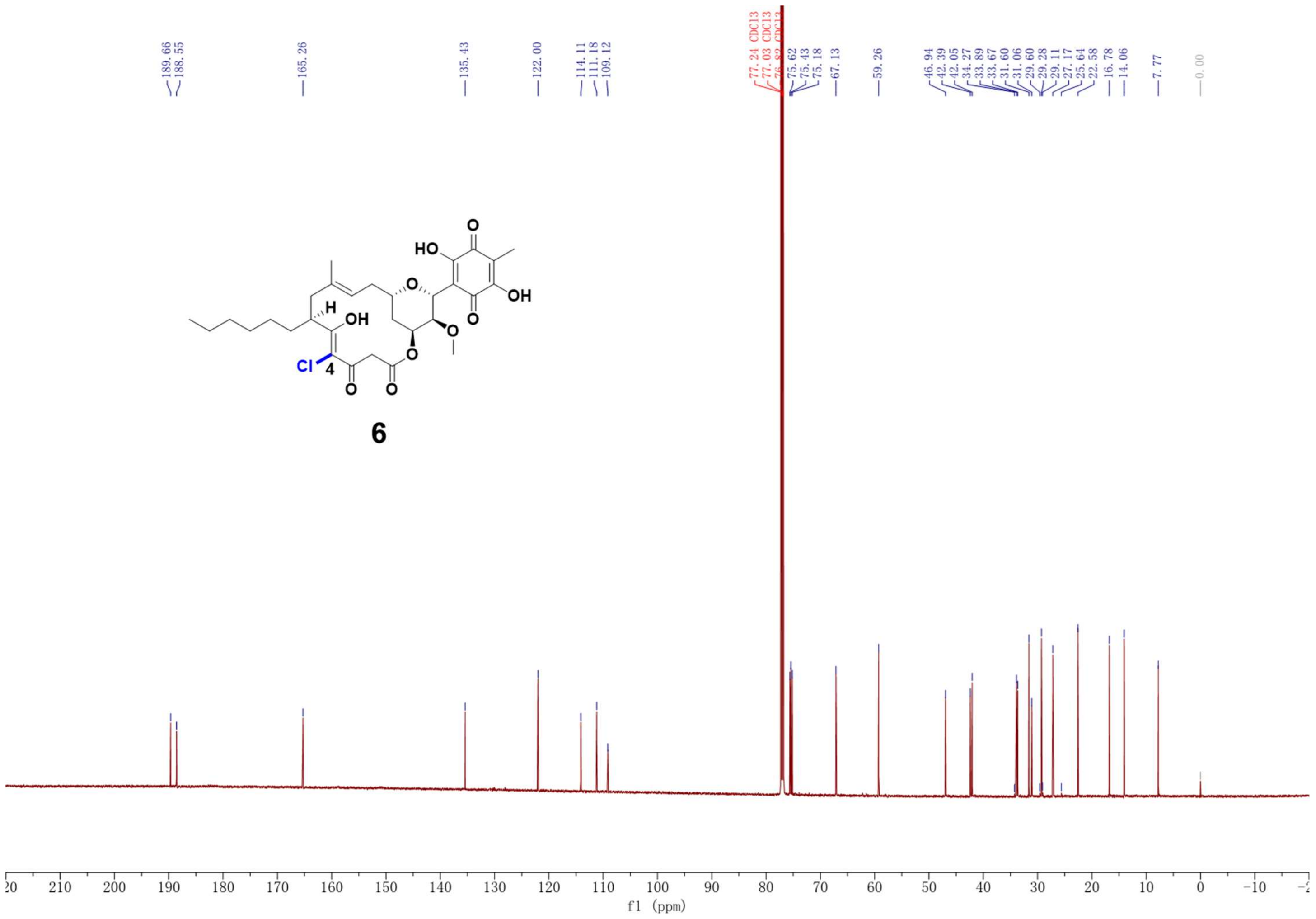

Supplementary Fig. 18. $^{13}$C NMR spectrum (150 MHz, $CDCl_3$) of compound **6**.

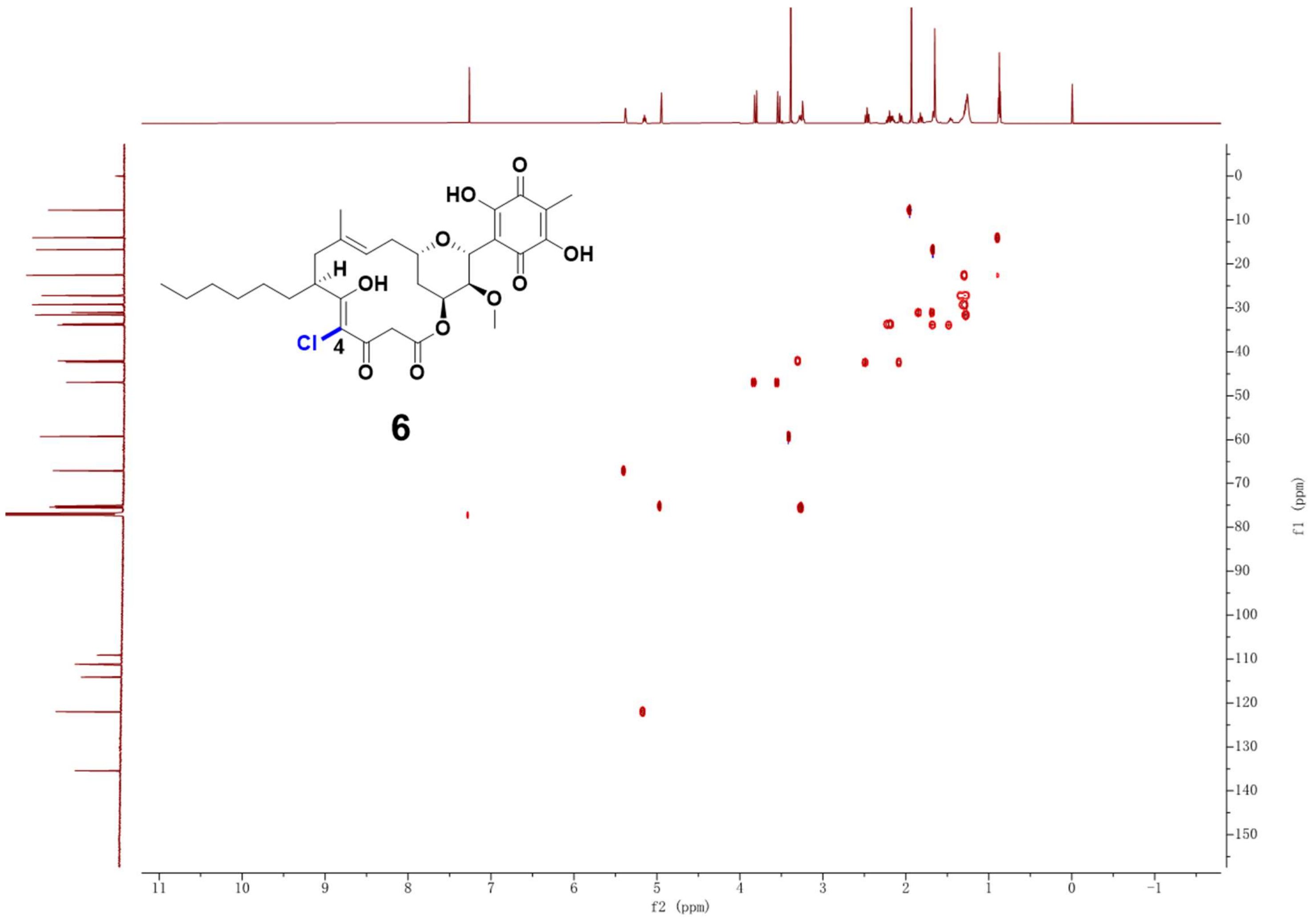

Supplementary Fig. 19. HSQC NMR spectrum (600 MHz, $CDCl_3$) of compound **6**.

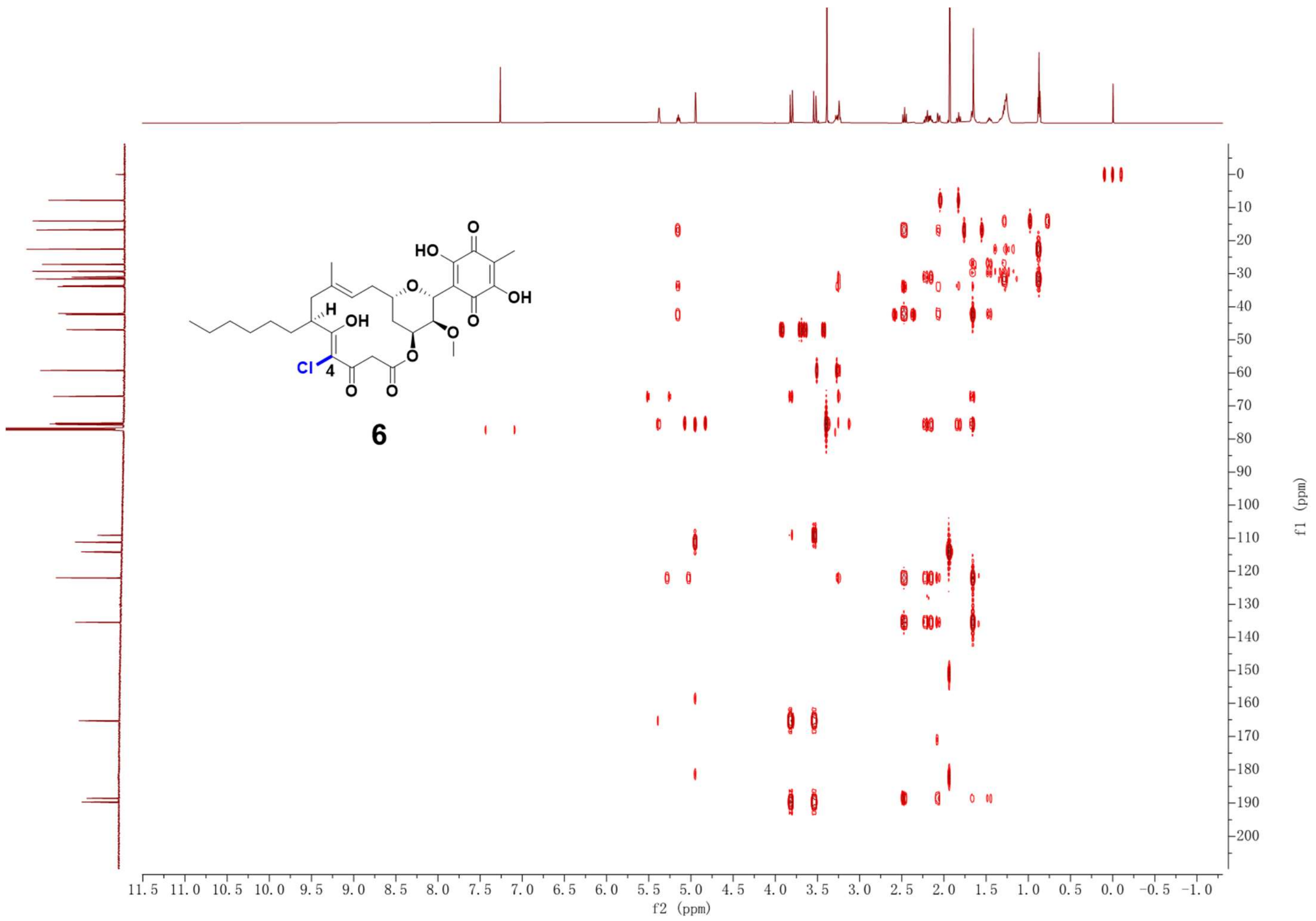

Supplementary Fig. 20. HMBC NMR spectrum (600 MHz, $CDCl_3$) of compound **6**.

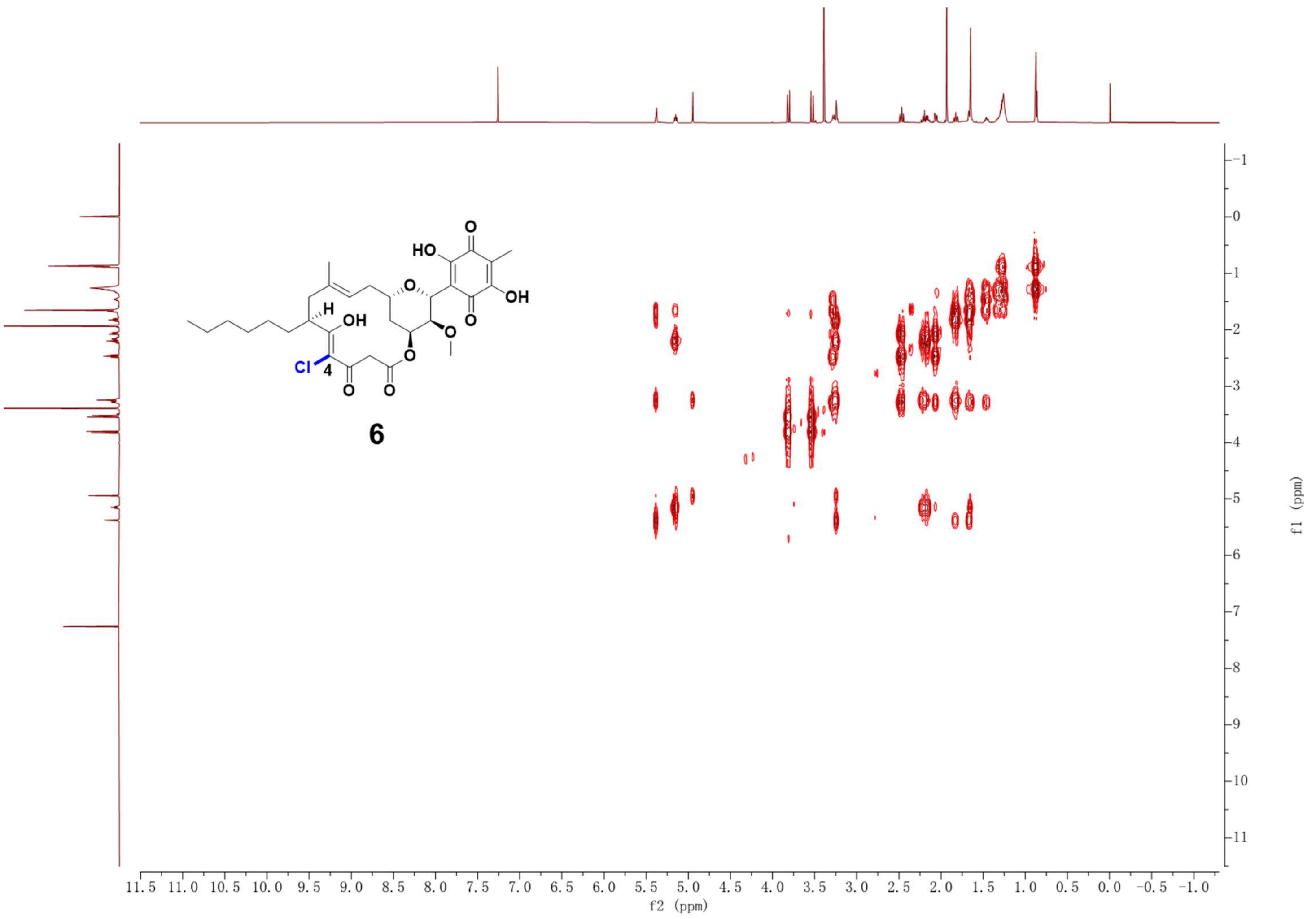

Supplementary Fig. 21. $^{1}H$-$^{1}H$ COSY NMR spectrum (600 MHz, $CDCl_3$) of compound **6**.

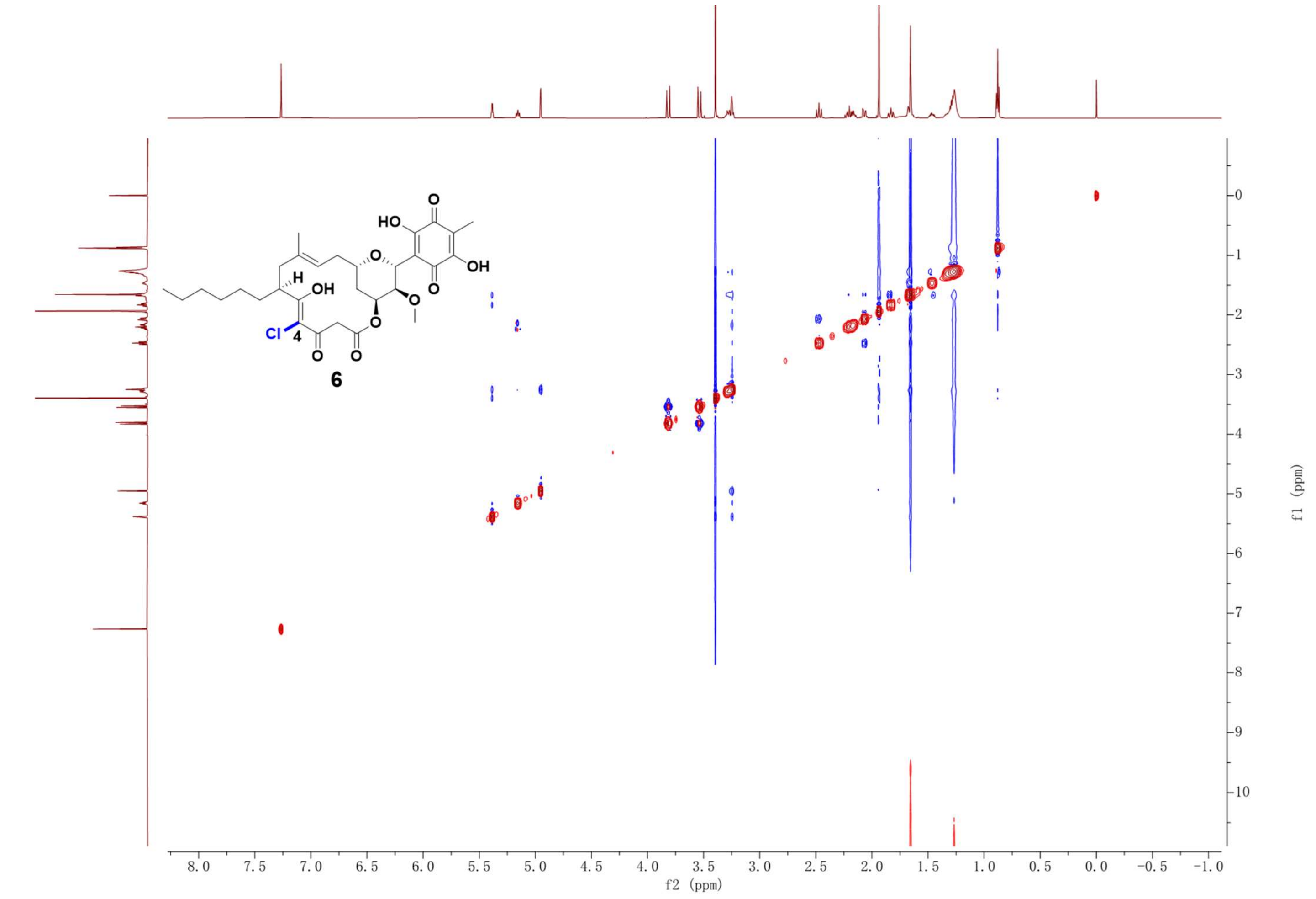

Supplementary Fig. 22. NOESY NMR spectrum (600 MHz, $CDCl_3$) of compound **6**.

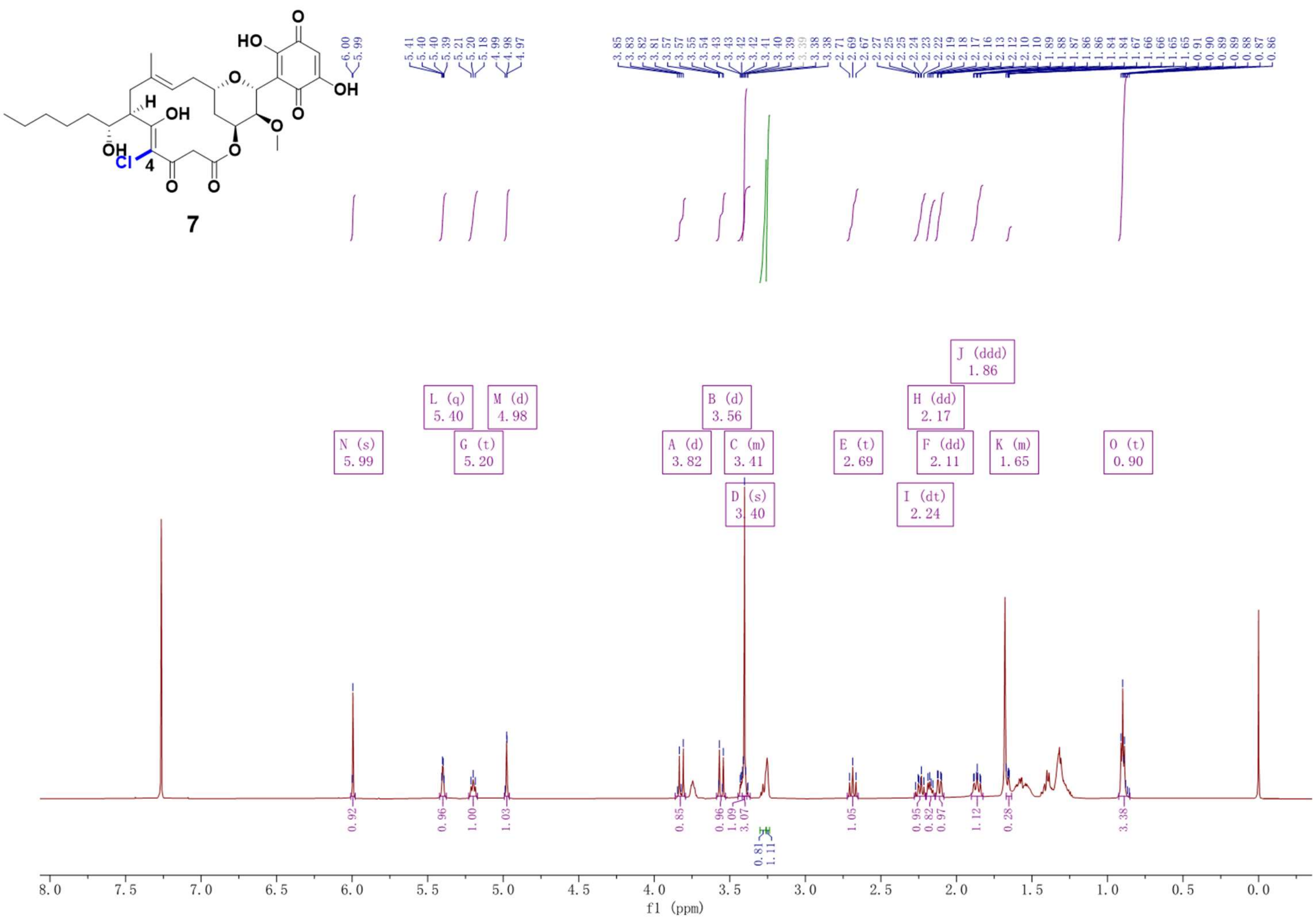

Supplementary Fig. 23. $^{1}$H NMR spectrum (600 MHz, $CDCl_3$) of compound **7**.

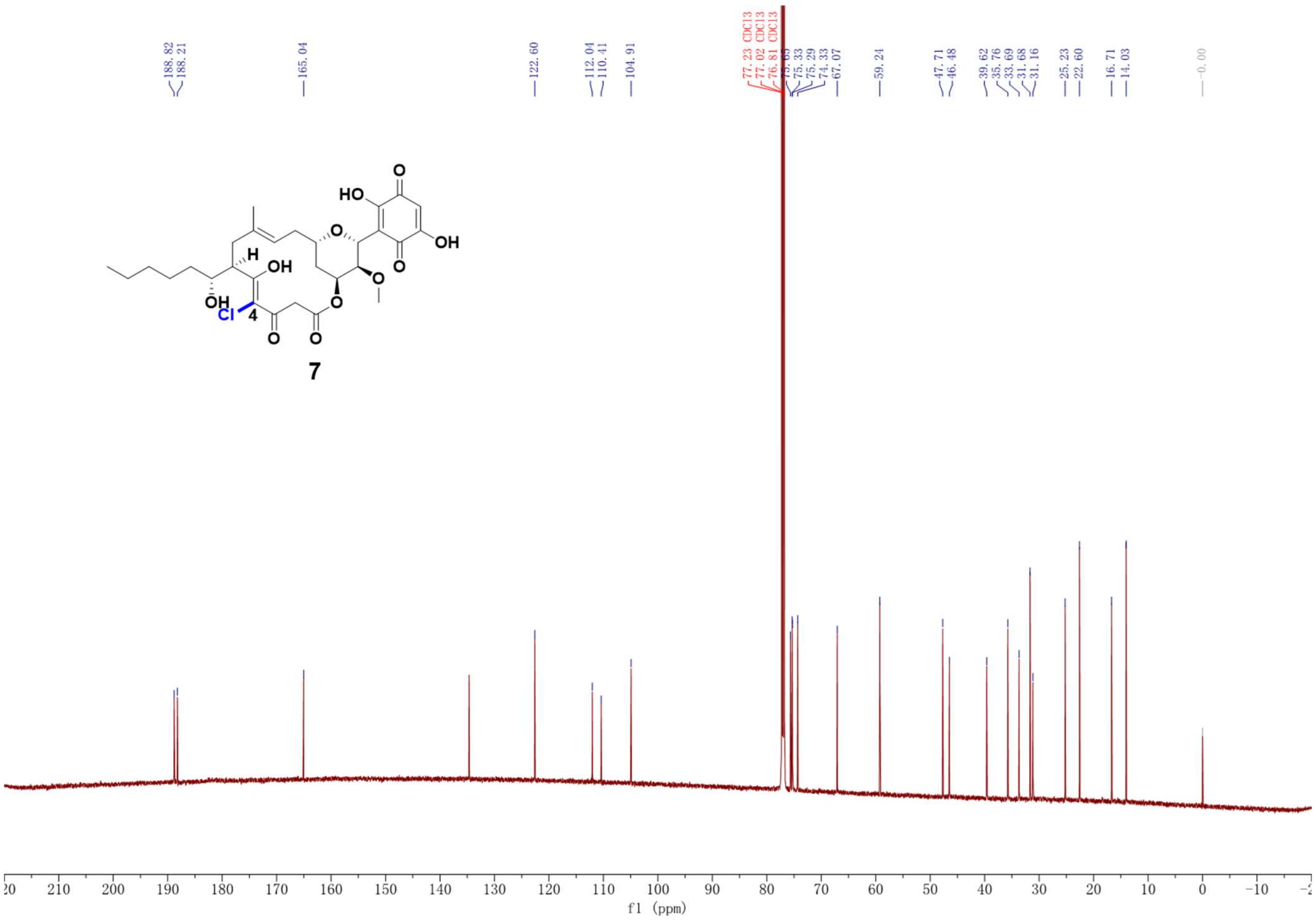

Supplementary Fig. 24. $^{13}$C NMR spectrum (150 MHz, $CDCl_3$) of compound **7**.

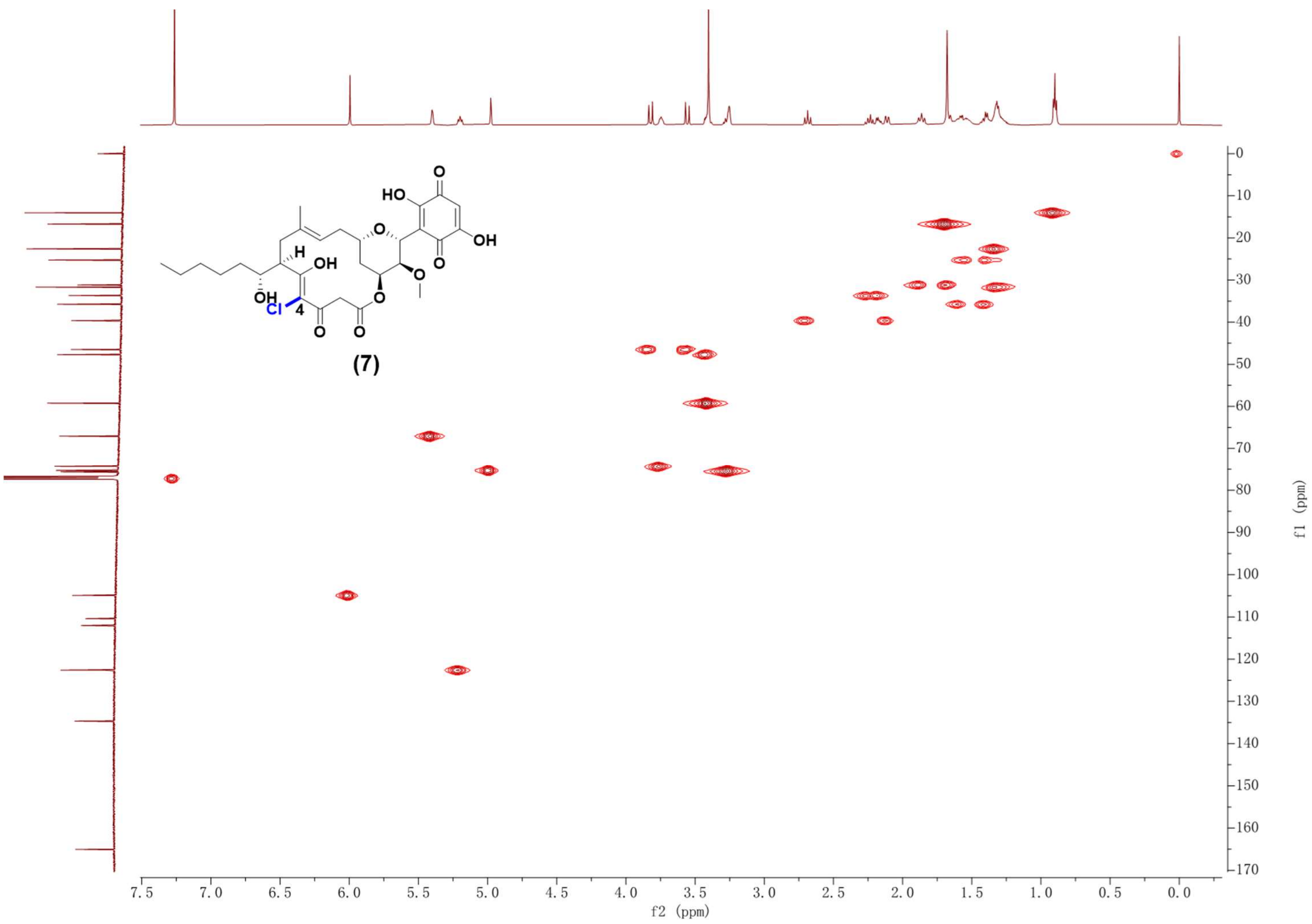

Supplementary Fig. 25. HSQC NMR spectrum (600 MHz, $CDCl_3$) of compound **7**.

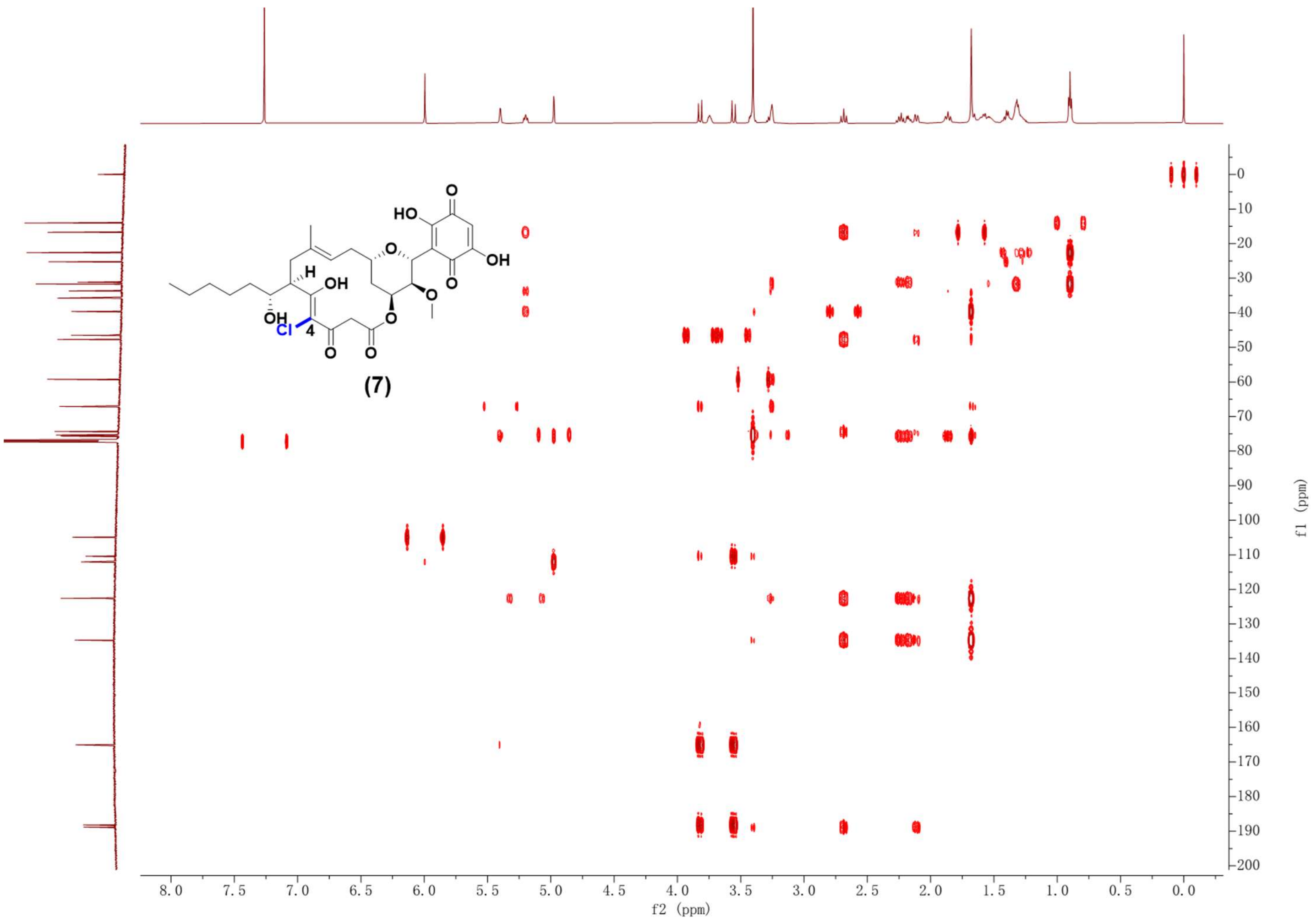

Supplementary Fig. 26. HMBC NMR spectrum (600 MHz, $CDCl_3$) of compound **7**.

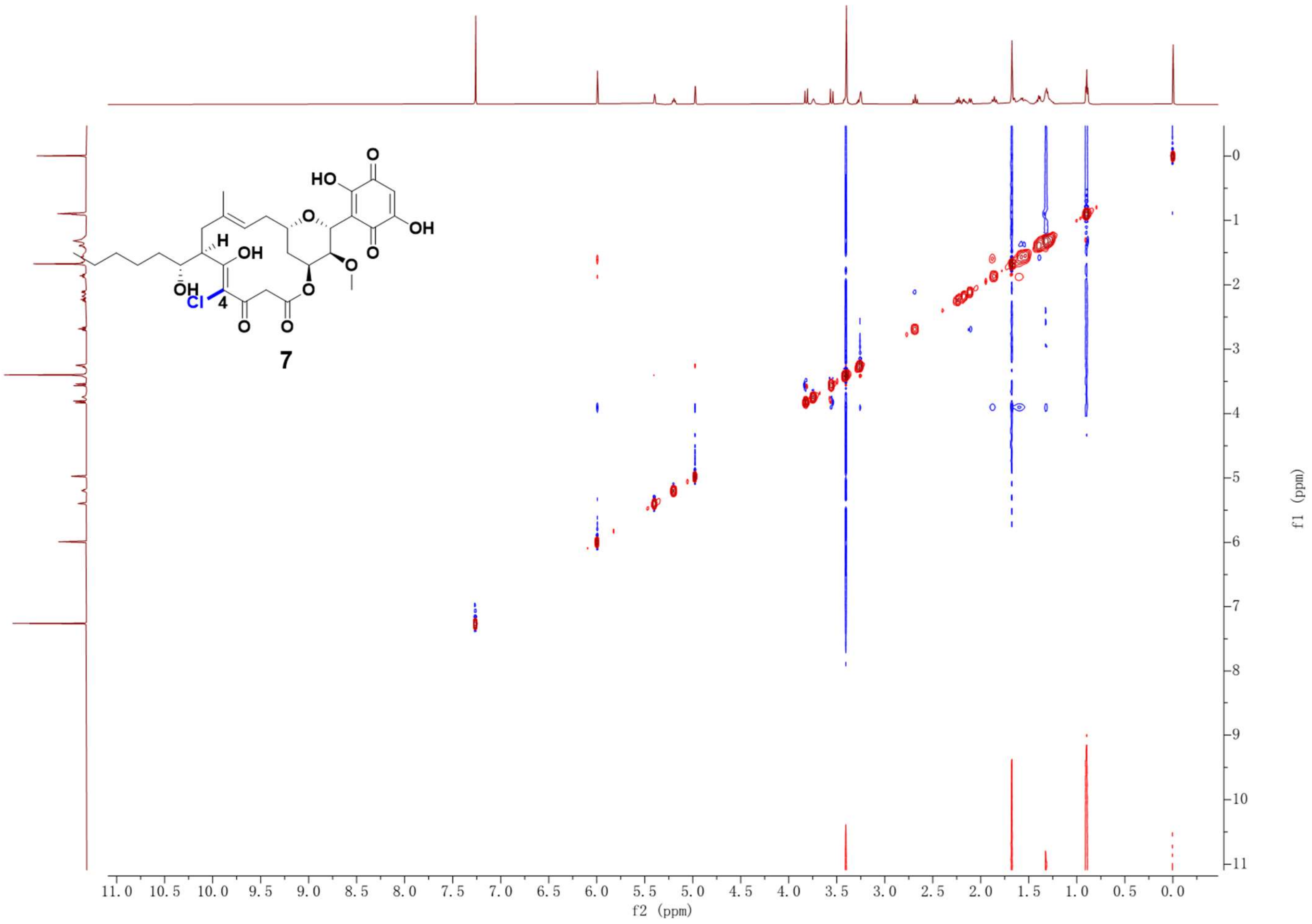

Supplementary Fig. 27. $^{1}H$-$^{1}H$ COSY NMR spectrum (600 MHz, $CDCl_3$) of compound **7**.

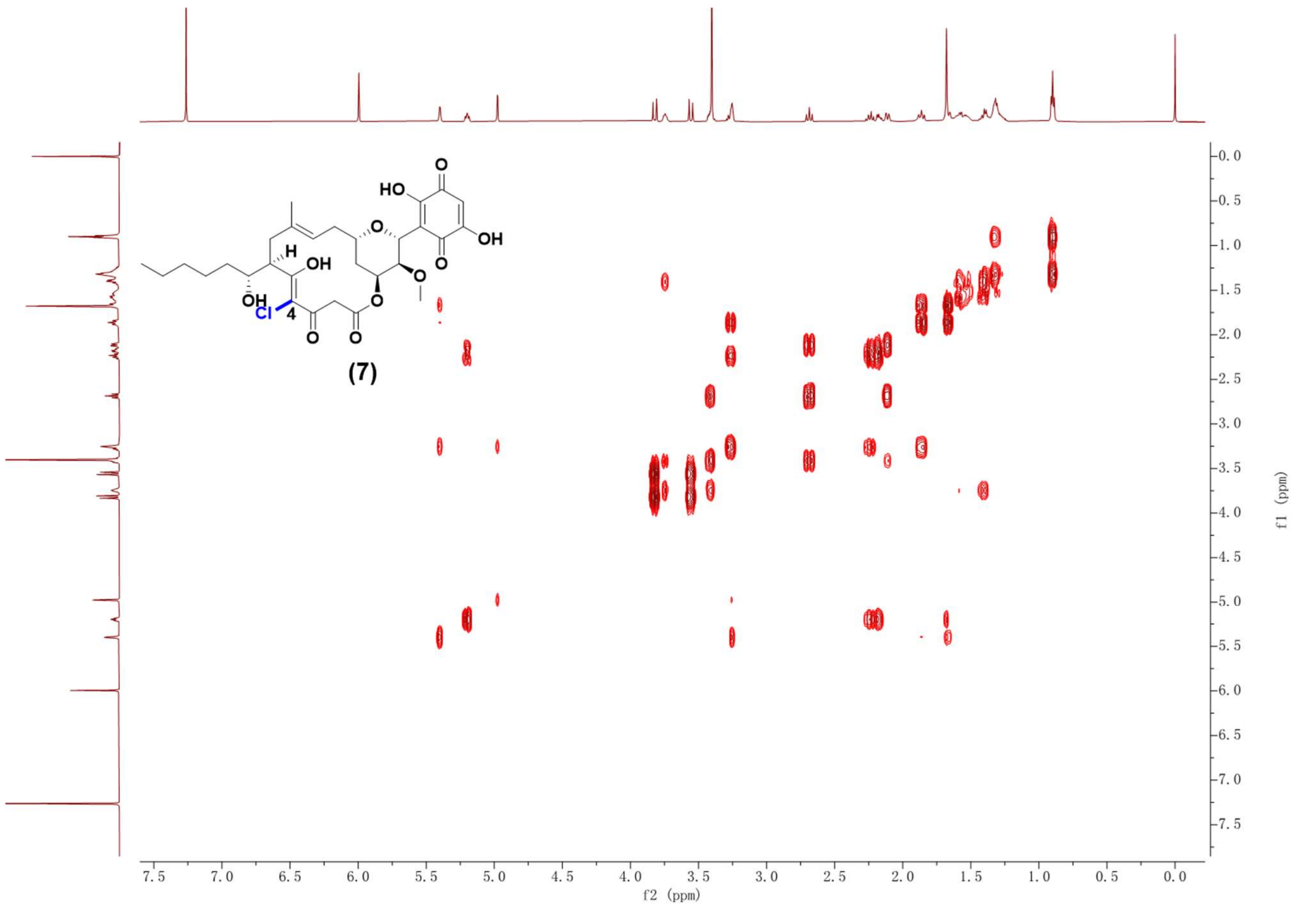

Supplementary Fig. 28. NOESY NMR spectrum (600 MHz, $CDCl_3$) of compound **7**.

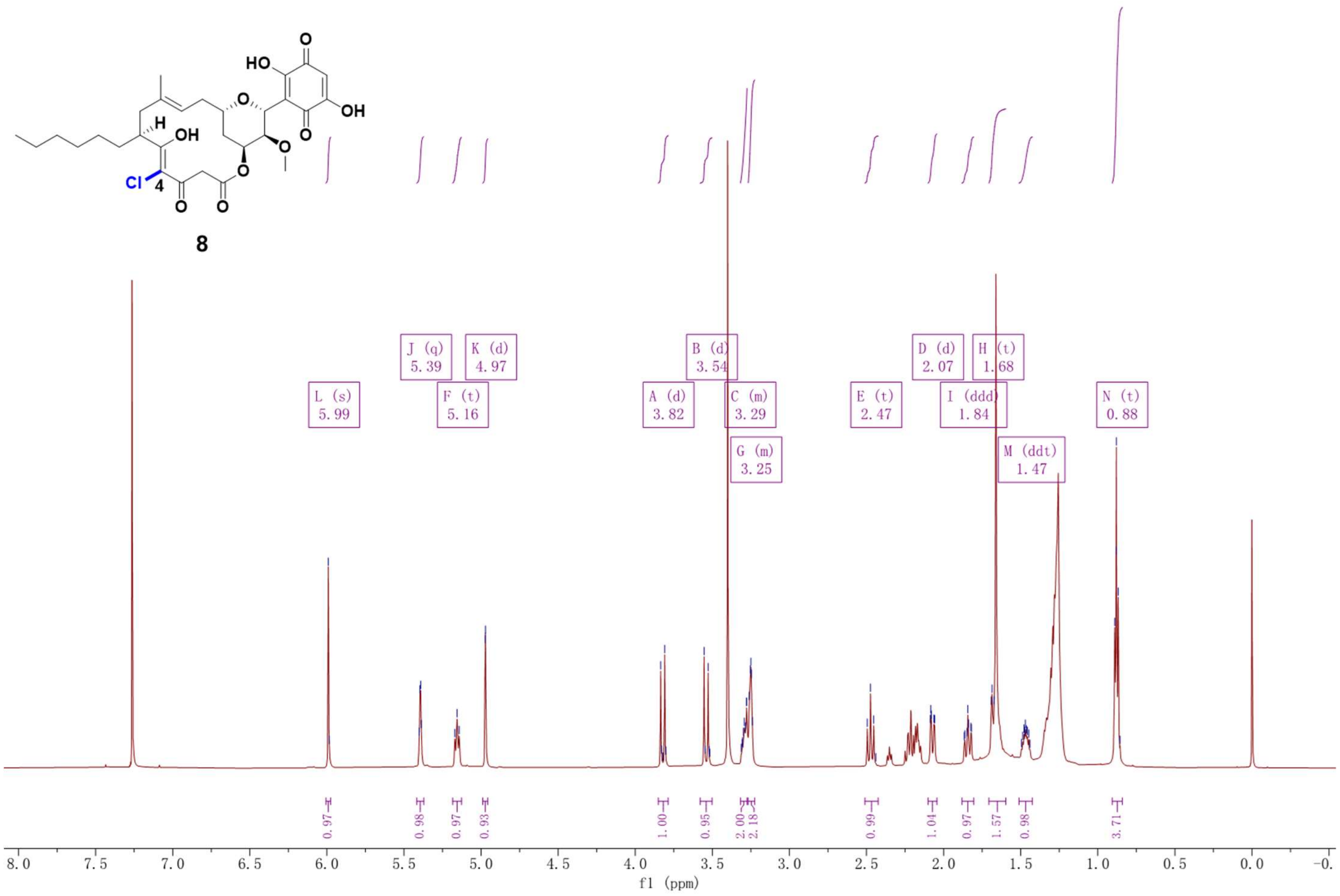

Supplementary Fig. 29. $^{1}$H NMR spectrum (600 MHz, $CDCl_3$) of compound **8**.

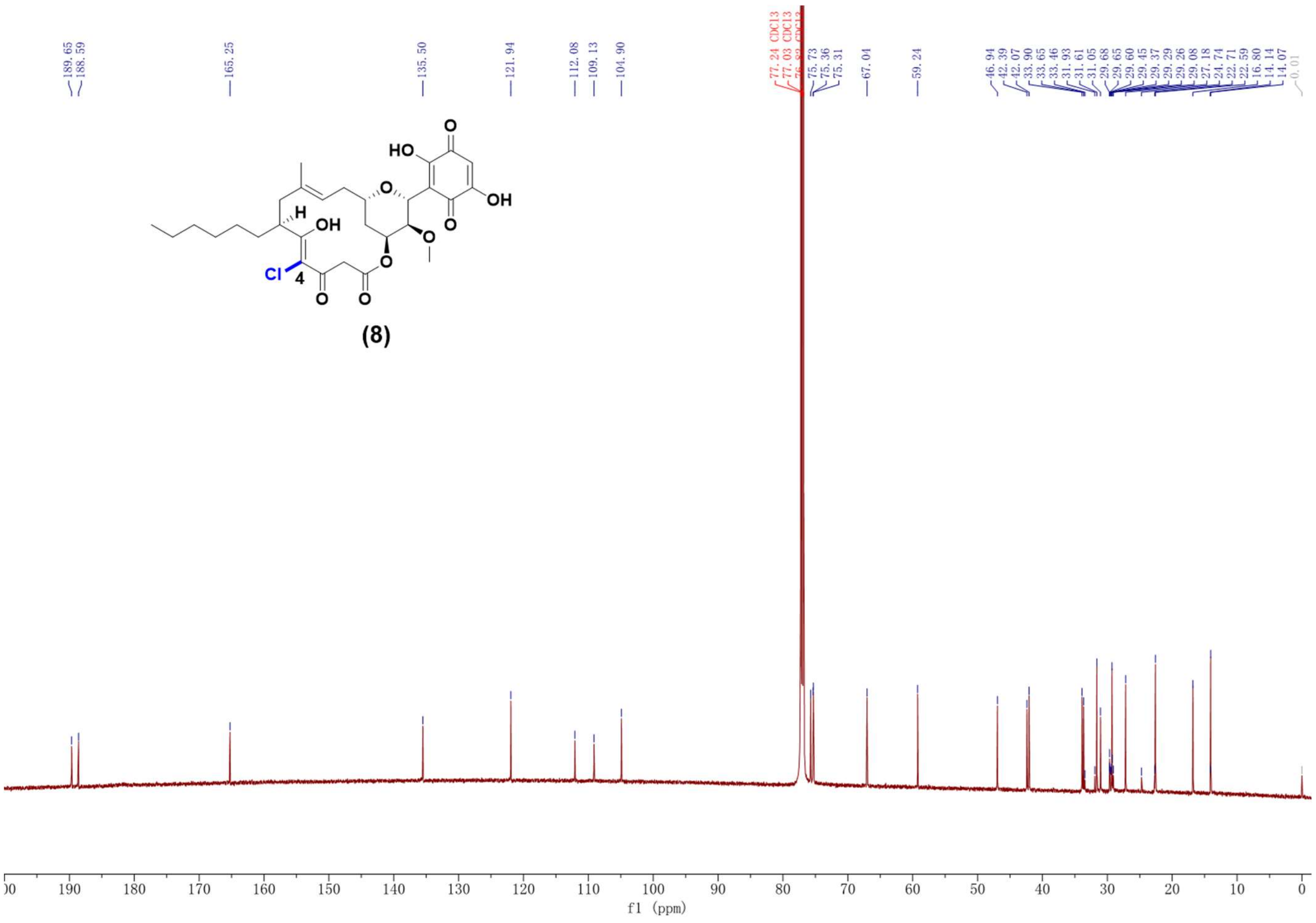

Supplementary Fig. 30. $^{13}$C NMR spectrum (150 MHz, $CDCl_3$) of compound **8**.

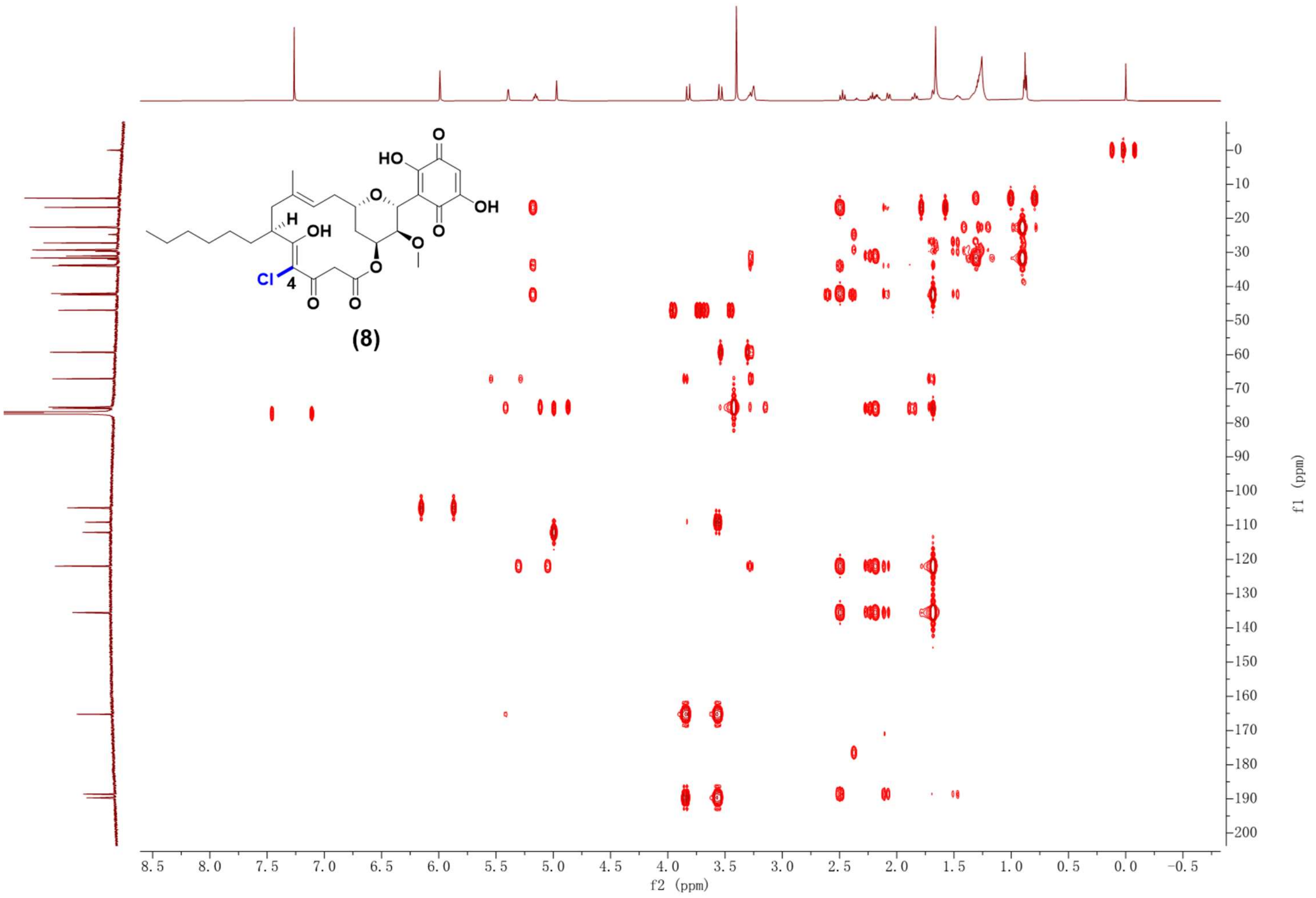

Supplementary Fig. 31. HSQC NMR spectrum (600 MHz, $CDCl_3$) of compound **8**.

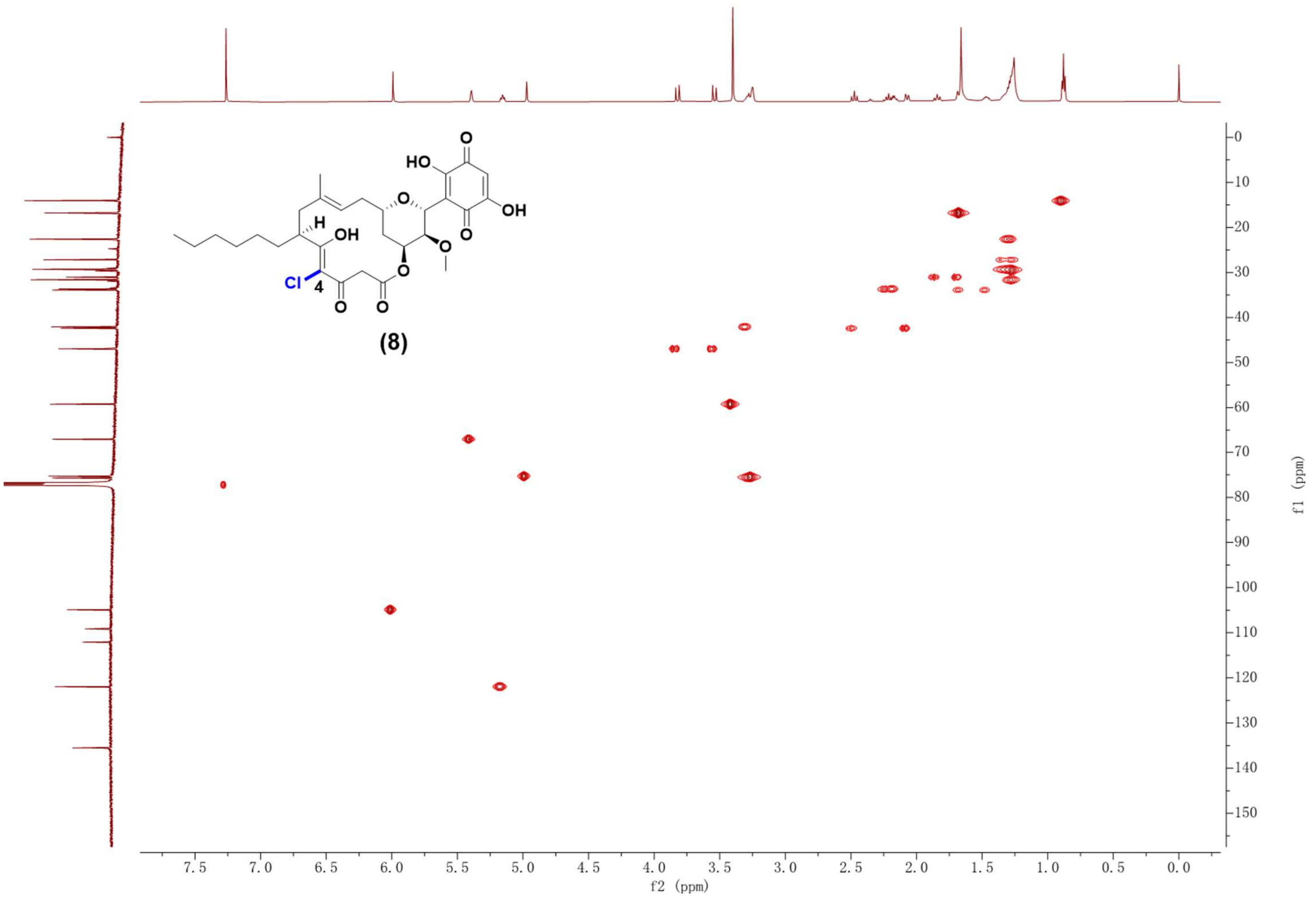

Supplementary Fig. 32. HMBC NMR spectrum (600 MHz, $CDCl_3$) of compound **8**.

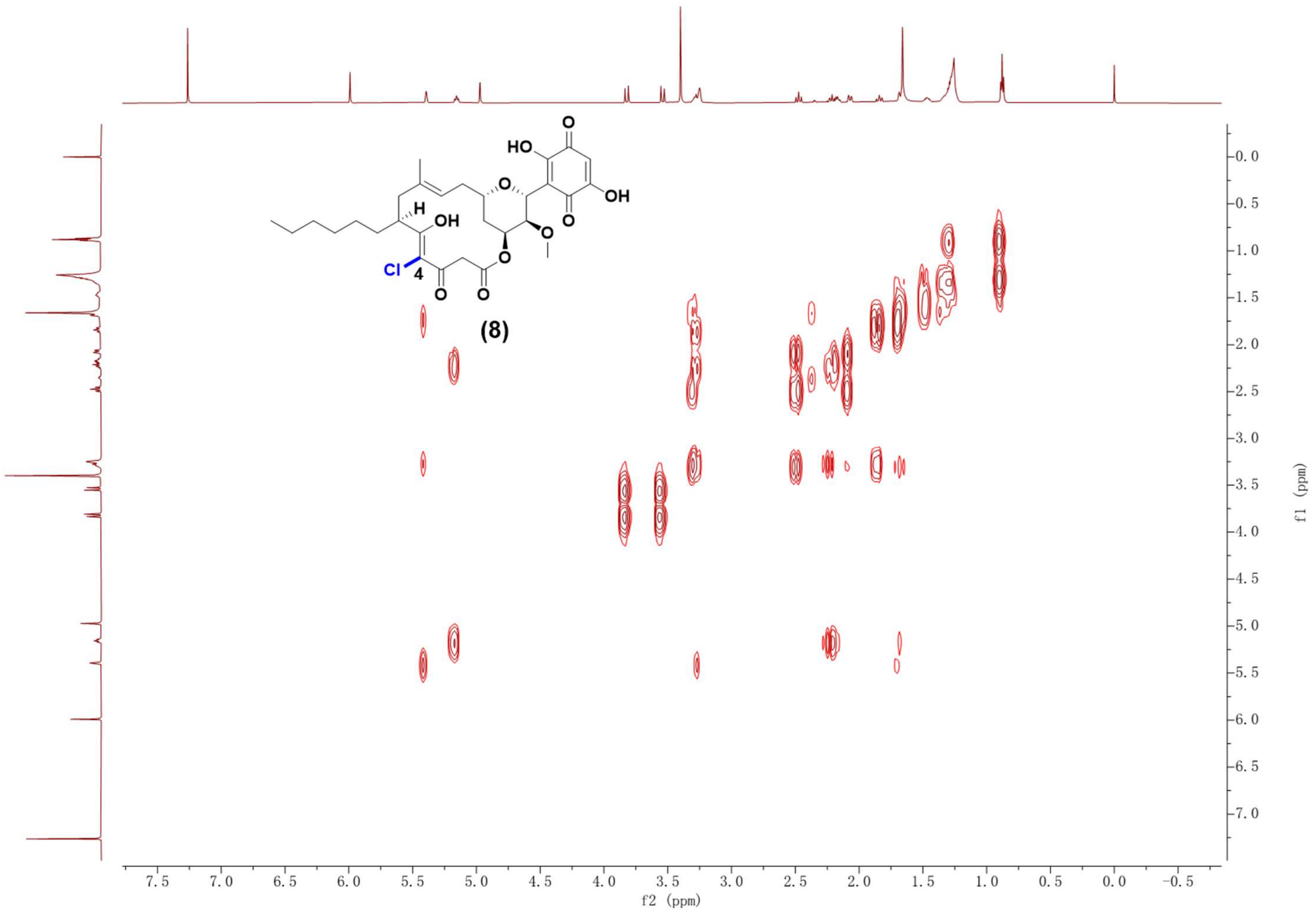

Supplementary Fig. 33. $^1H$-$^1H$ COSY NMR spectrum (600 MHz, $CDCl_3$) of compound **8**.

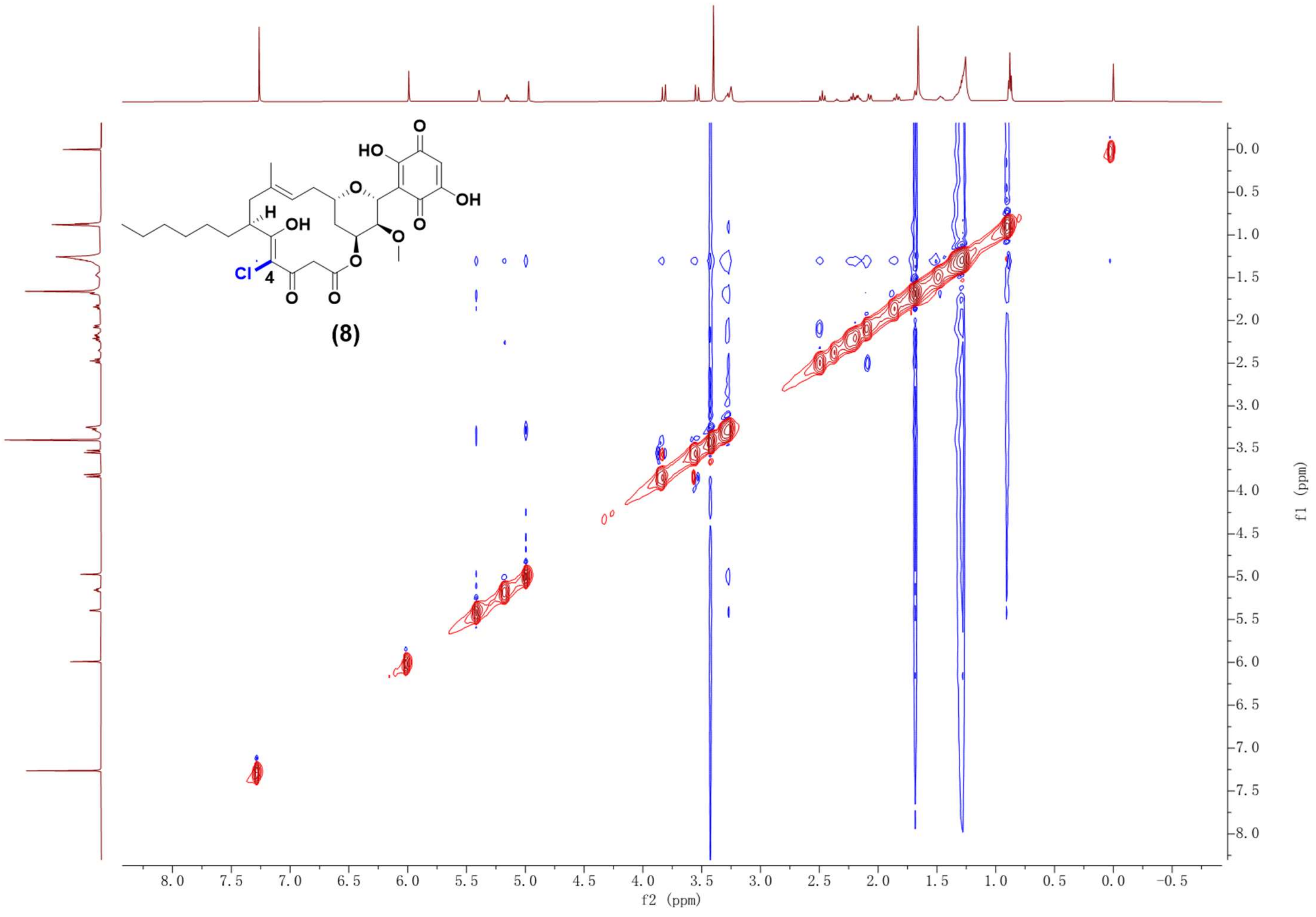

Supplementary Fig. 34. NOESY NMR spectrum (600 MHz, $CDCl_3$) of compound **8**.

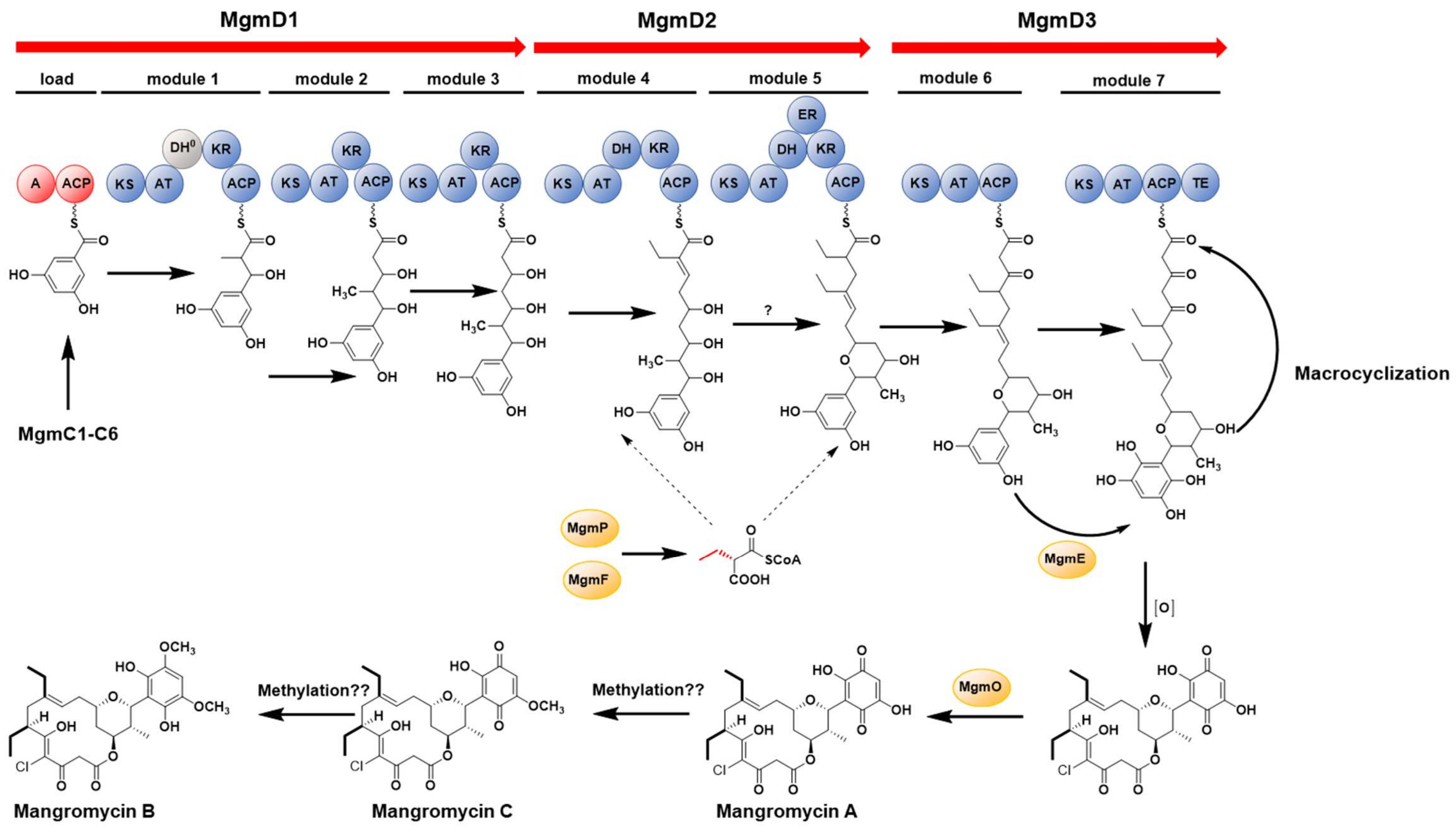

Supplementary Fig. 35. Proposed biosynthetic pathway of mangromycin **A**-**C**.

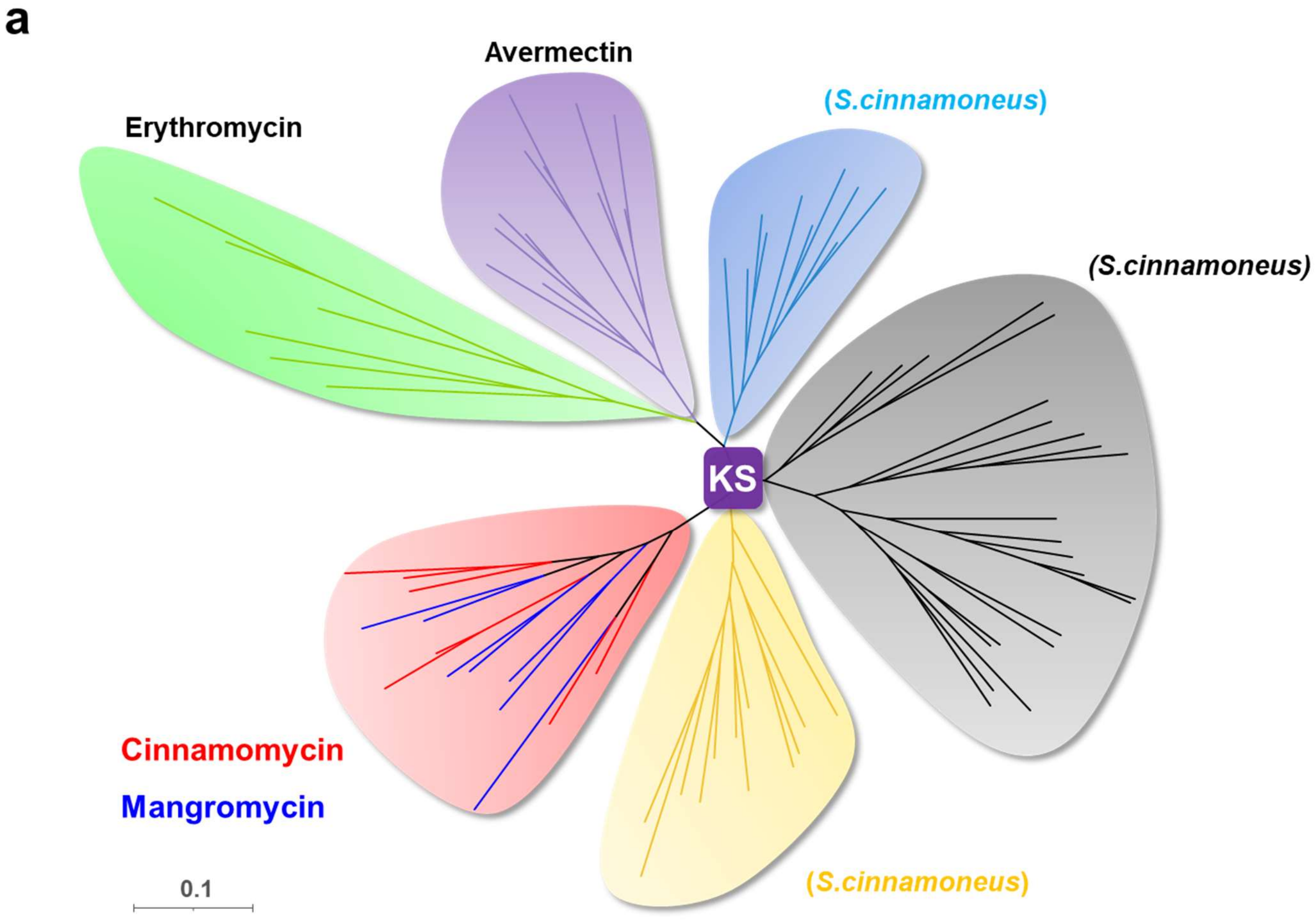

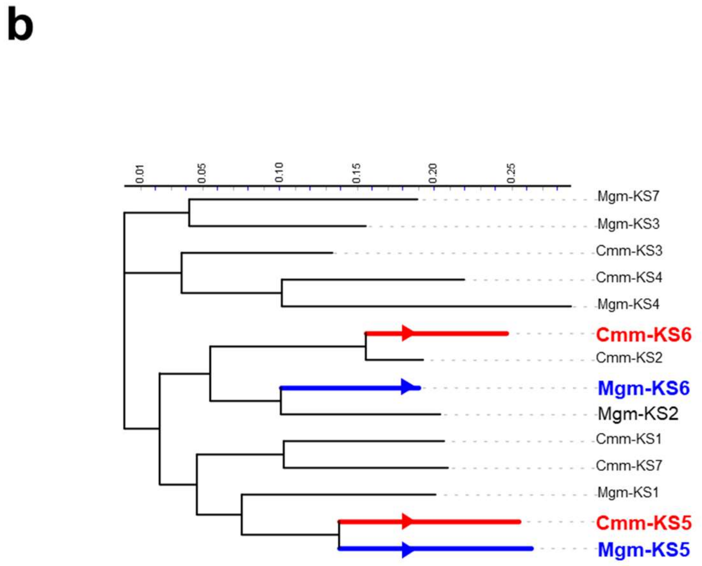

Supplementary Fig. 36. Phylogenetic tree analysis of KS domains.

**(a)** Phylogenetic tree analysis of KS domains from *cmm* and *mgm* BGCs compared to KS domains from other BGCs. **(b)** Phylogenetic tree analysis of KS domains from *cmm* and *mgm* BGCs.

**a**

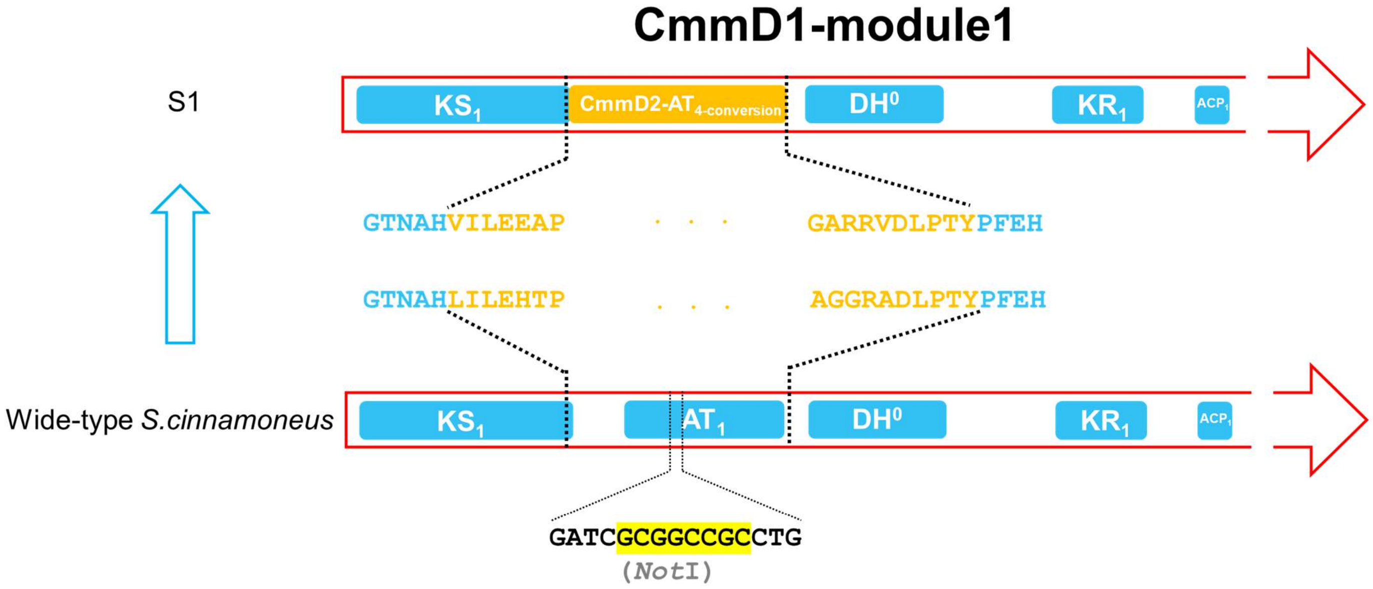

**b**

| Mutant | Origin | Primer Name | Primer Sequence (5'→3') | Screening Method |
|---|---|---|---|---|
| S1 | Wild-type *S.cinnamoneus* | S1-C1 | ACGGAAGTCGACT GGTCGTC | PCR products were digested using *Not*I |
| | | S1-C2 | ACGGACCAGTAGT GCCGGTG | |

**c**

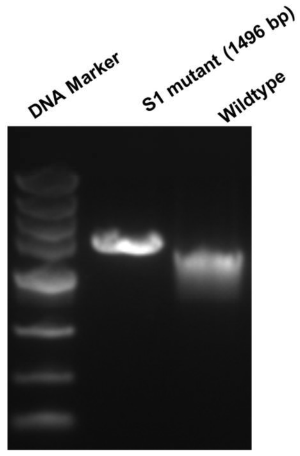

Supplementary Fig. 37. Construction and confirmation of mutant S1 strain.

(**a**): Schematic diagram of construction of mutant S1, *Not*I restriction enzyme cutting site used for screening is highlighted in yellow; (**b**) Primers and methods for mutant strain screening; (**c**) Agarose gel screening of mutant S1 by PCR and restriction enzyme digesting. The blue squares represent domains from CmmD1-module 1.

*Not*I enzyme digestion always causes fragment blurring, so S1 strain was finally confirmed by DNA sequencing.

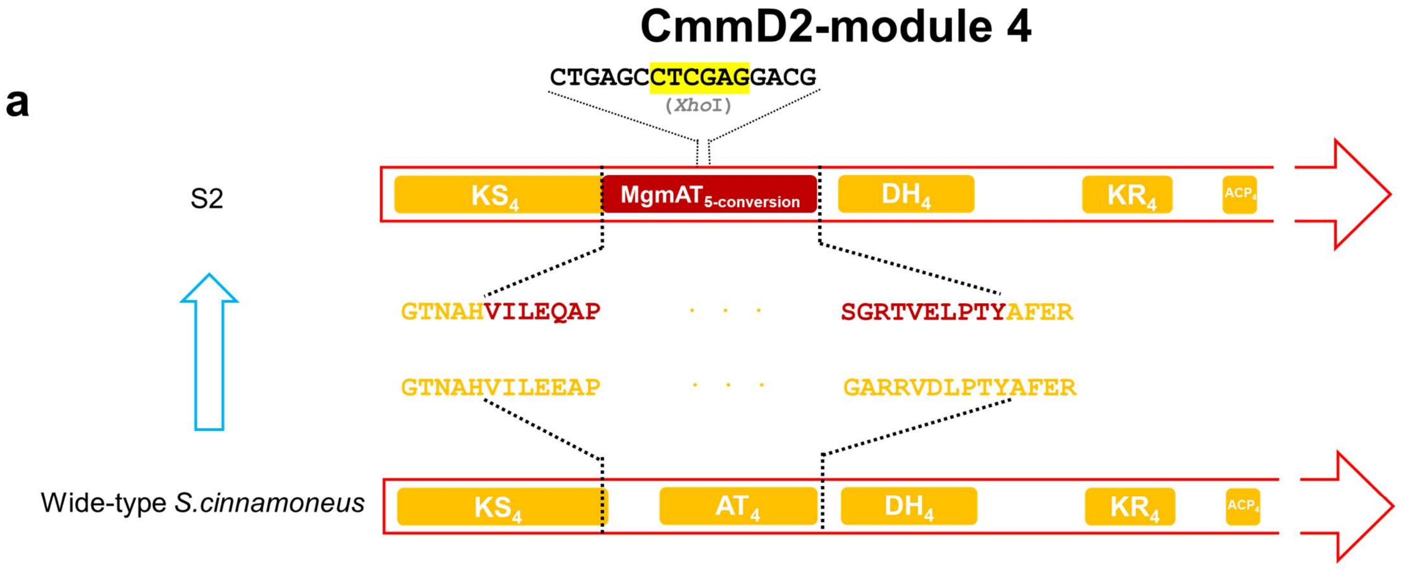

**b**

| Mutant | Origin | Primer Name | Primer Sequence (5'→3') | Screening Method |
|---|---|---|---|---|
| S2 | Wild-type *S.cinnamoneus* | S2-C1 | TGGTAAGGGCATGACGAACACG | PCR products were digested using *Xho*I |
| | | S2-C2 | TGACAGGTCCACTCCTCCTCGTC | |

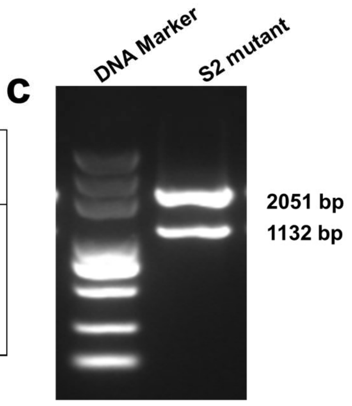

Supplementary Fig. 38. Construction and confirmation of mutant S2 strain.

(**a**): Schematic diagram of construction of mutant S2, *Xho*I restriction enzyme cutting site used for screening is highlighted in yellow; (**b**) Primers and methods for mutant strain screening; (**c**) Agarose gel screening of mutant S2 by PCR and restriction enzyme digesting. The yellow squares represent domains from CmmD2-module 4.

### CmmD2-module 5

a

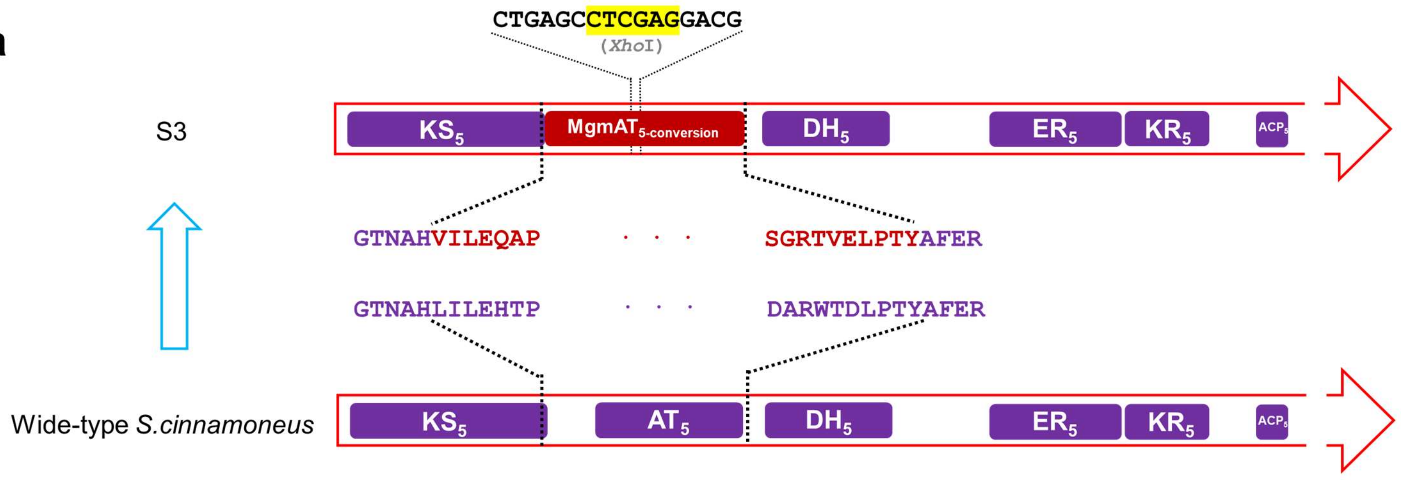

b

| Mutant | Origin | Primer Name | Primer Sequence (5'→3') | Screening Method |
|---|---|---|---|---|
| S3 | Wild-type *S.cinnamoneus* | S3-C1 | ATGTACCAGGACTACCTGCTGC | PCR products were digested using *Xho*I |
| | | S3-C2 | GTGCAGCCACAGGTGCGTCC | |

c

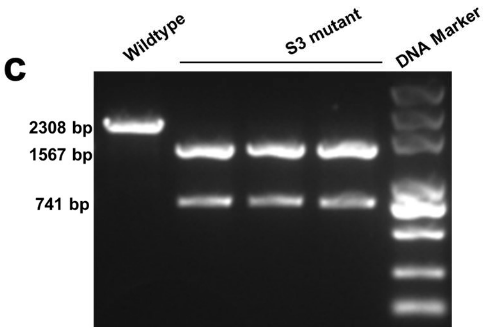

Supplementary Fig. 39. Construction and confirmation of mutant S3 strain.

(**a**): Schematic diagram of construction of mutant S3, *Xho*I restriction enzyme cutting site used for screening is highlighted in yellow; (**b**) Primers and methods for mutant strain screening; (**c**) Agarose gel screening of mutant S3 by PCR and restriction enzyme digesting. The purple squares represent domains from CmmD2-module 5.

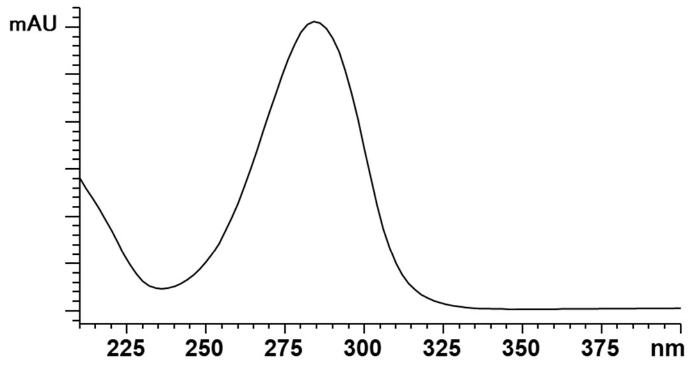

Compound
(**1-4, 9-12, 13a, 14a, 15a, 16, 17, 18a, 18c, 19a, 19c, 20**)

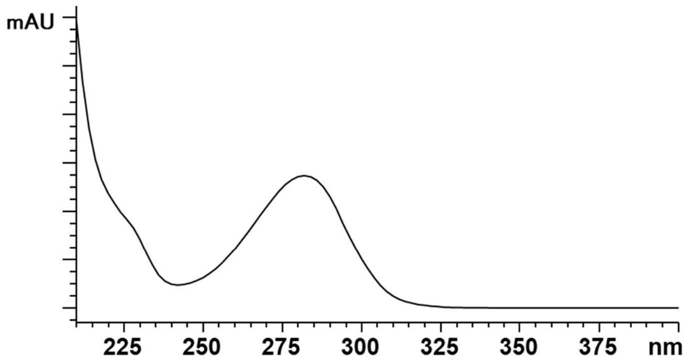

Compound
(**13b, 18b, 19b**)

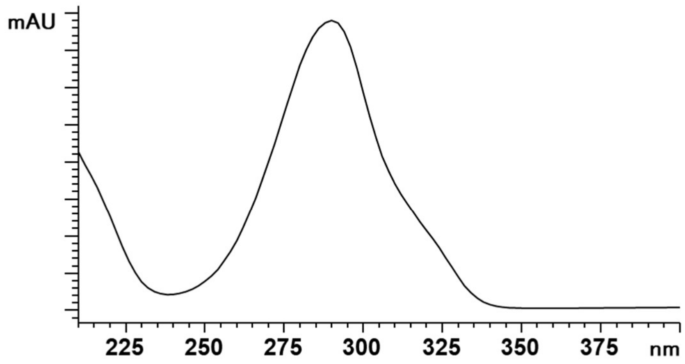

Compound
(**5-8, 21**)

Supplementary Fig. 40. UV-Vis spectra of compounds **1**-**21**.

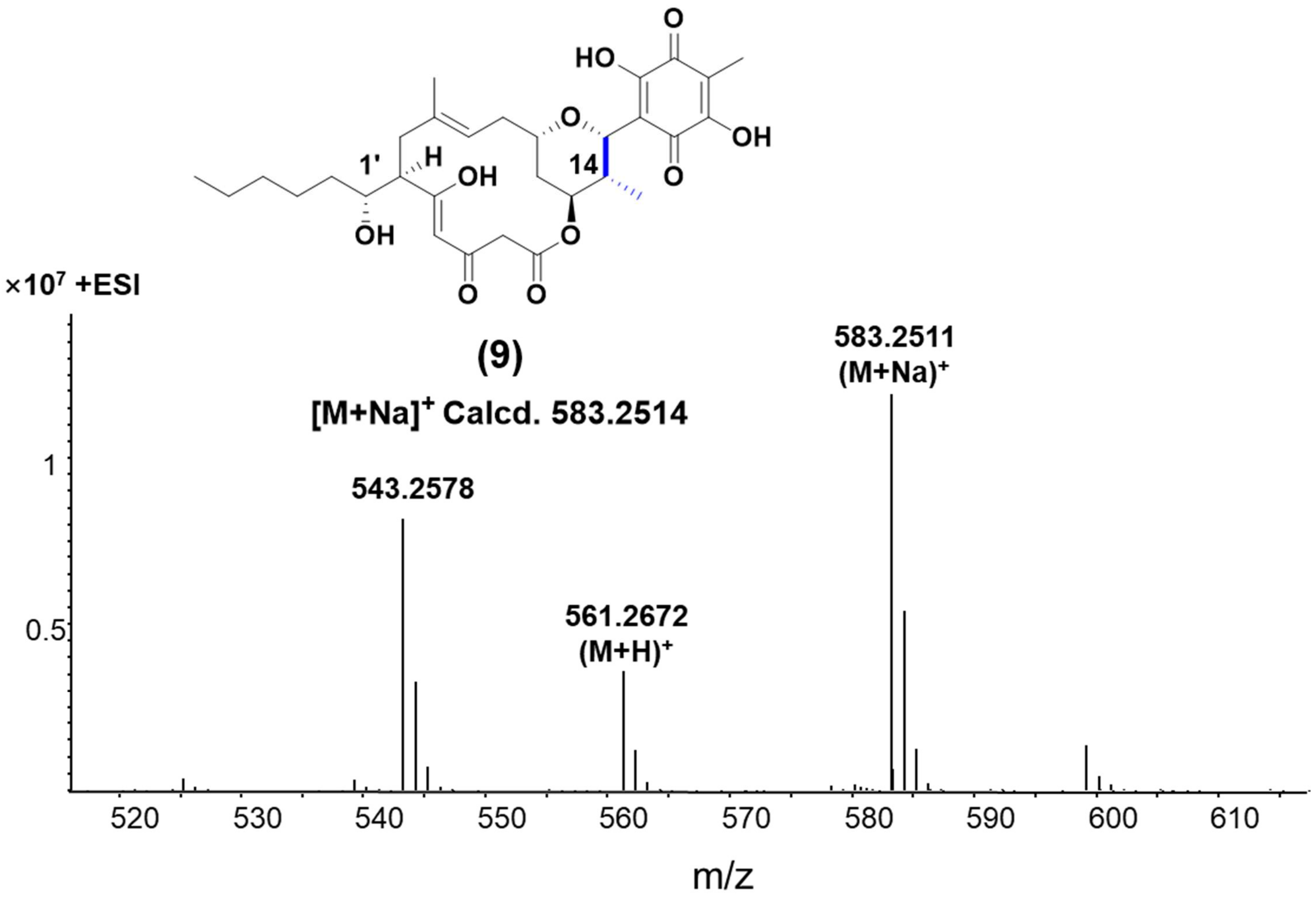

Supplementary Fig. 41. HR-ESI-MS of derivates **9**.

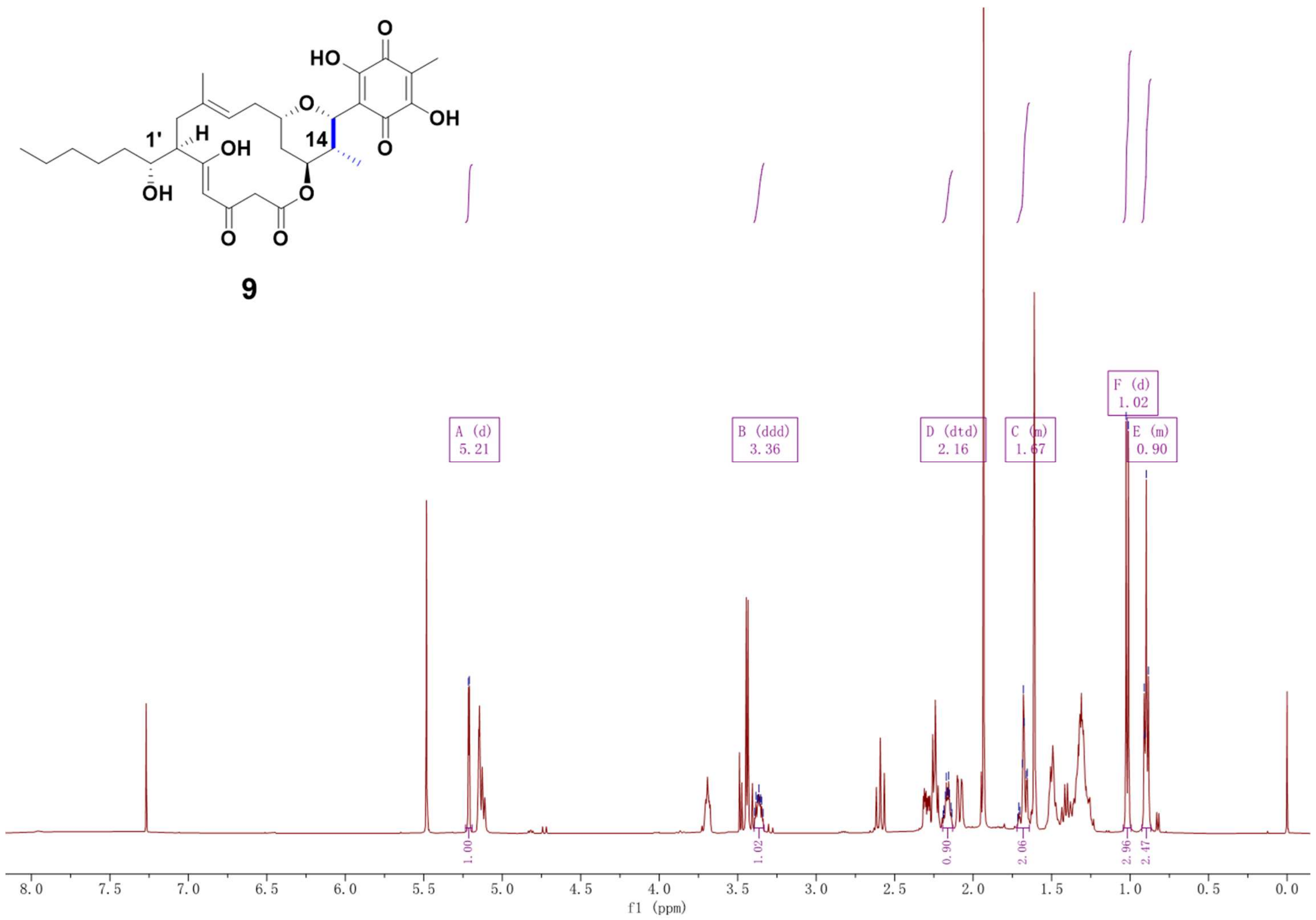

Supplementary Fig. 42. $^{1}H$ NMR spectrum (500 MHz, $CDCl_3$) of compound **9**.

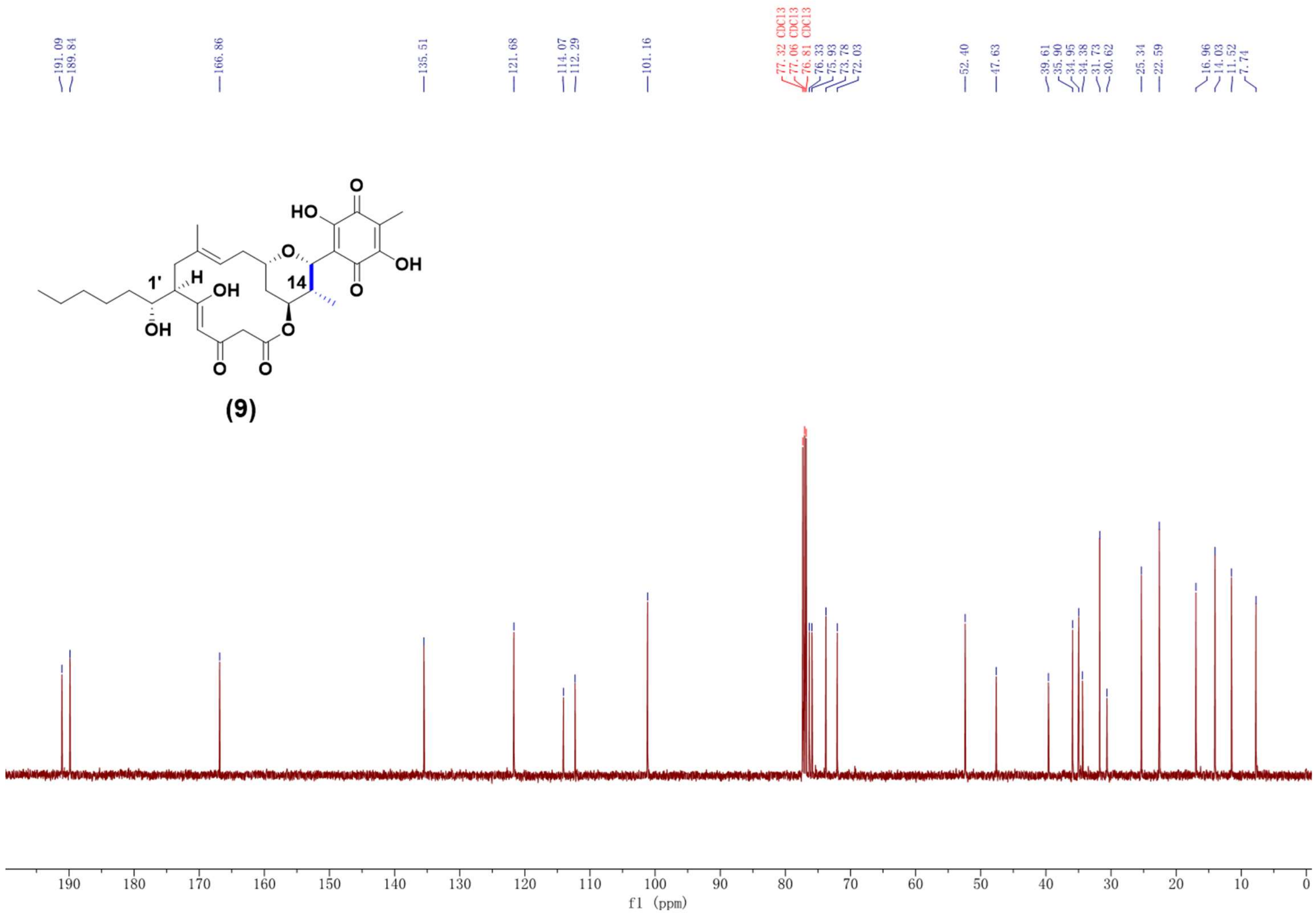

Supplementary Fig. 43. $^{13}$C NMR spectrum (125 MHz, $CDCl_3$) of compound **9**.

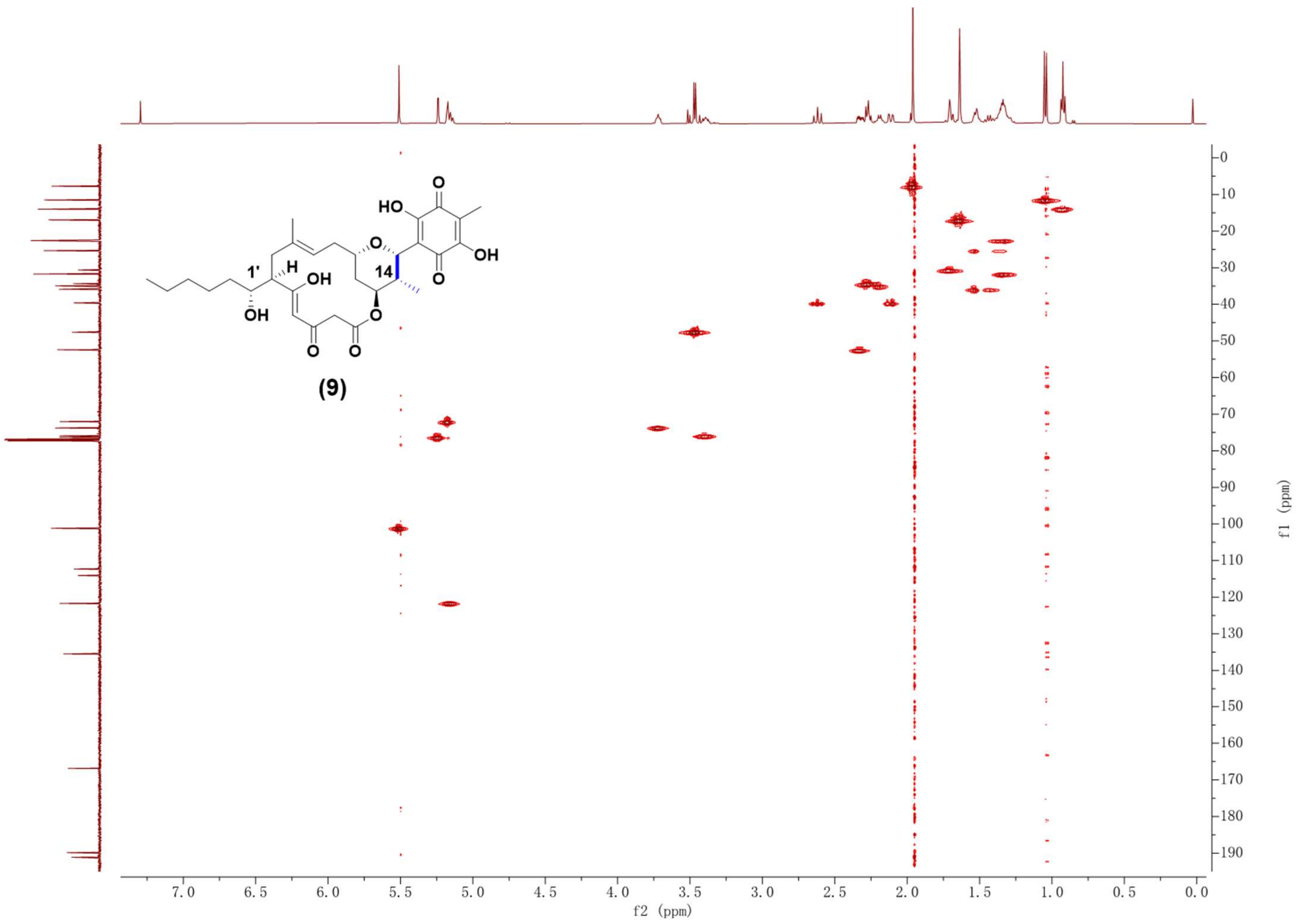

Supplementary Fig. 44. HSQC NMR spectrum (500 MHz, $CDCl_3$) of compound **9**.

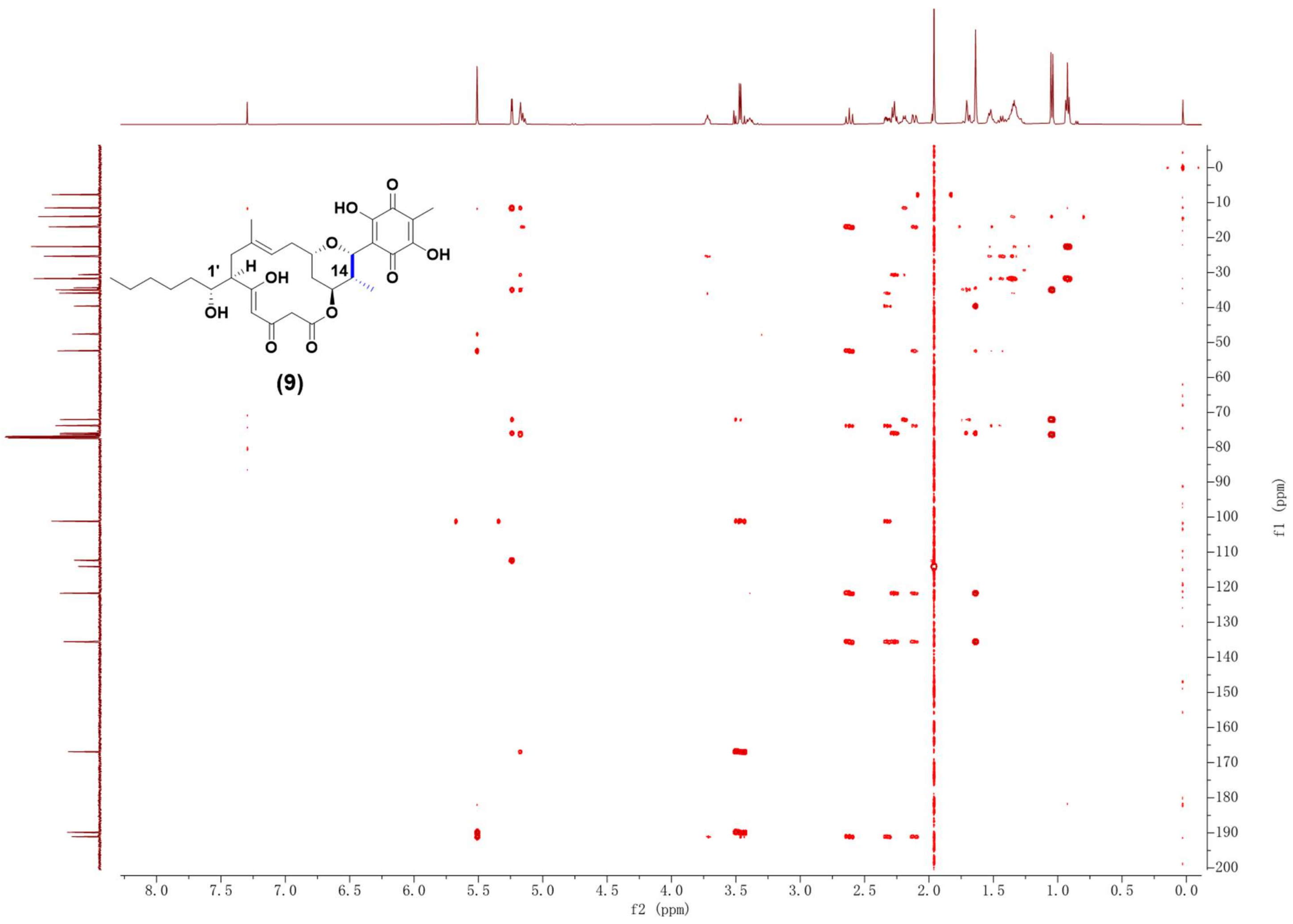

Supplementary Fig. 45. HMBC NMR spectrum (500 MHz, $CDCl_3$) of compound **9**.

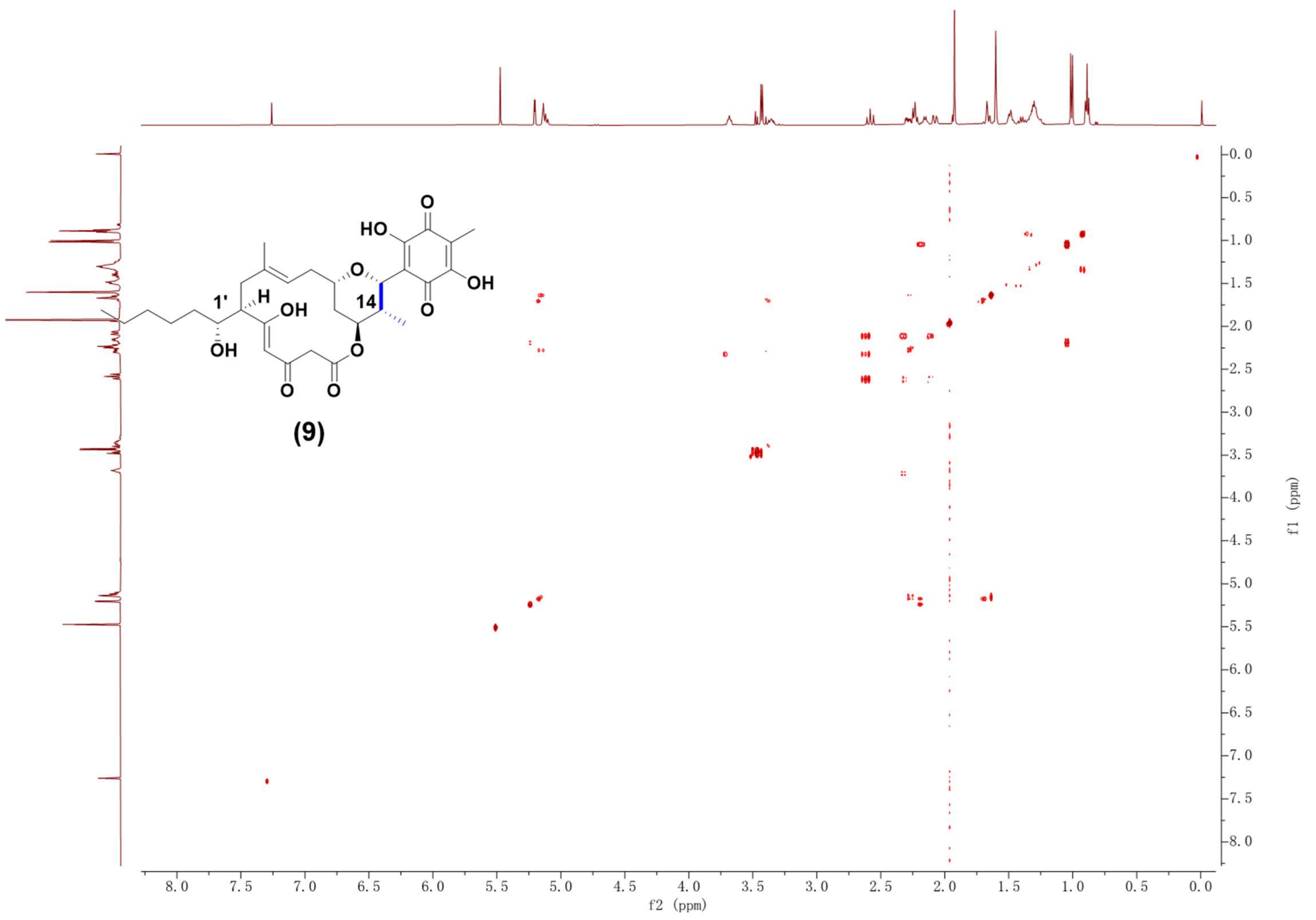

Supplementary Fig. 46. $^1H$-$^1H$ COSY NMR spectrum (500 MHz, $CDCl_3$) of compound **9**.

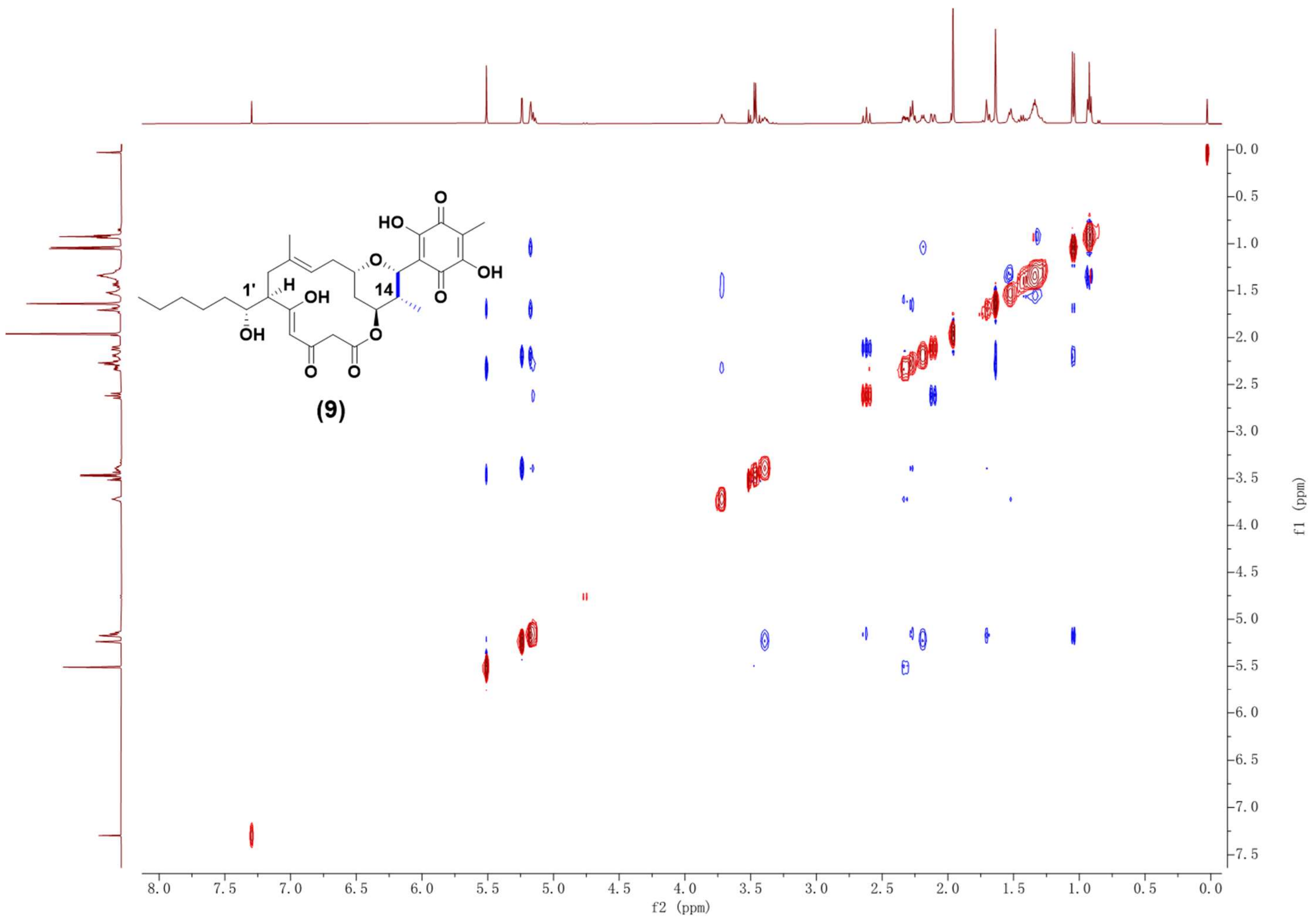

Supplementary Fig. 47. NOESY NMR spectrum (500 MHz, $CDCl_3$) of compound **9**.

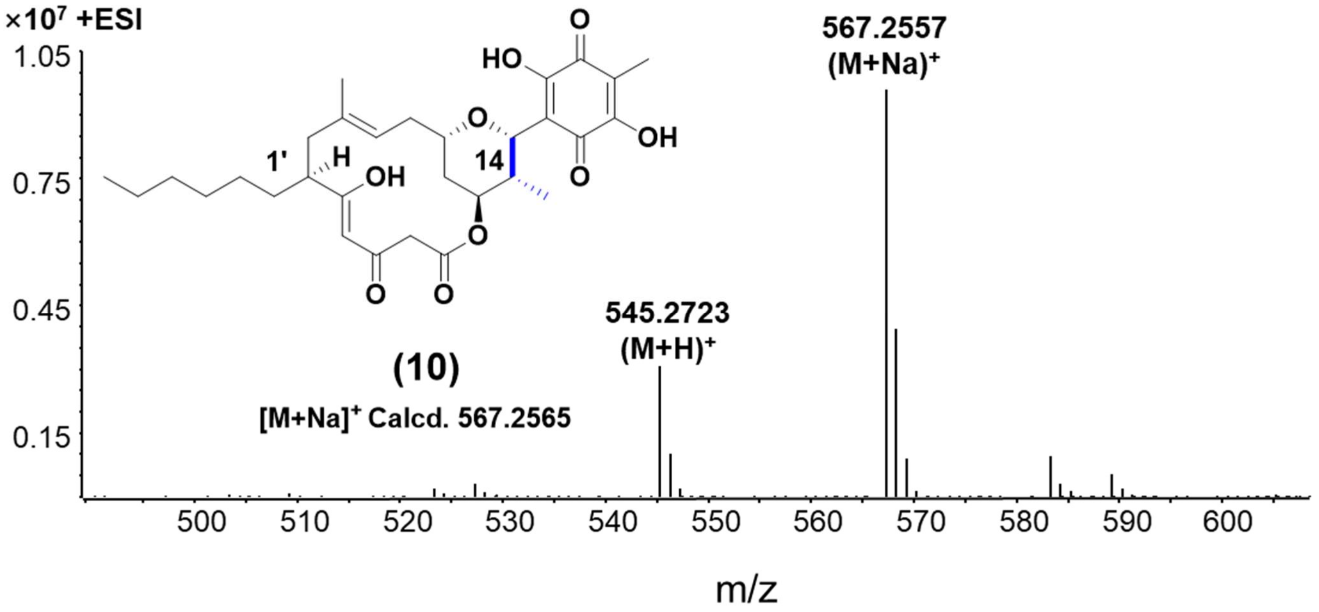

Supplementary Fig. 48. HR-ESI-MS spectrum of compound **10**.

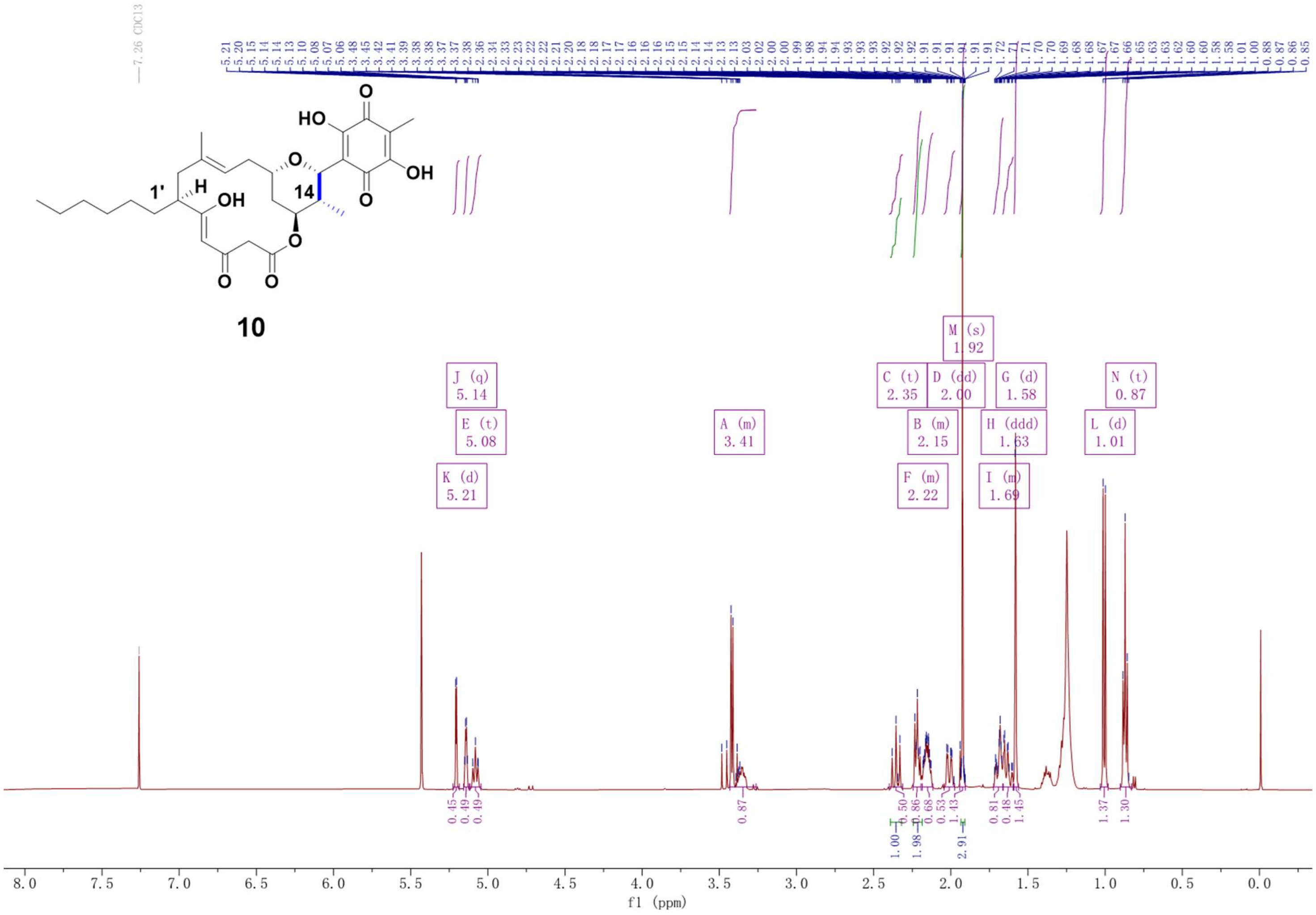

Supplementary Fig. 49. $^{1}H$ NMR spectrum (500 MHz, $CDCl_3$) of compound **10**.

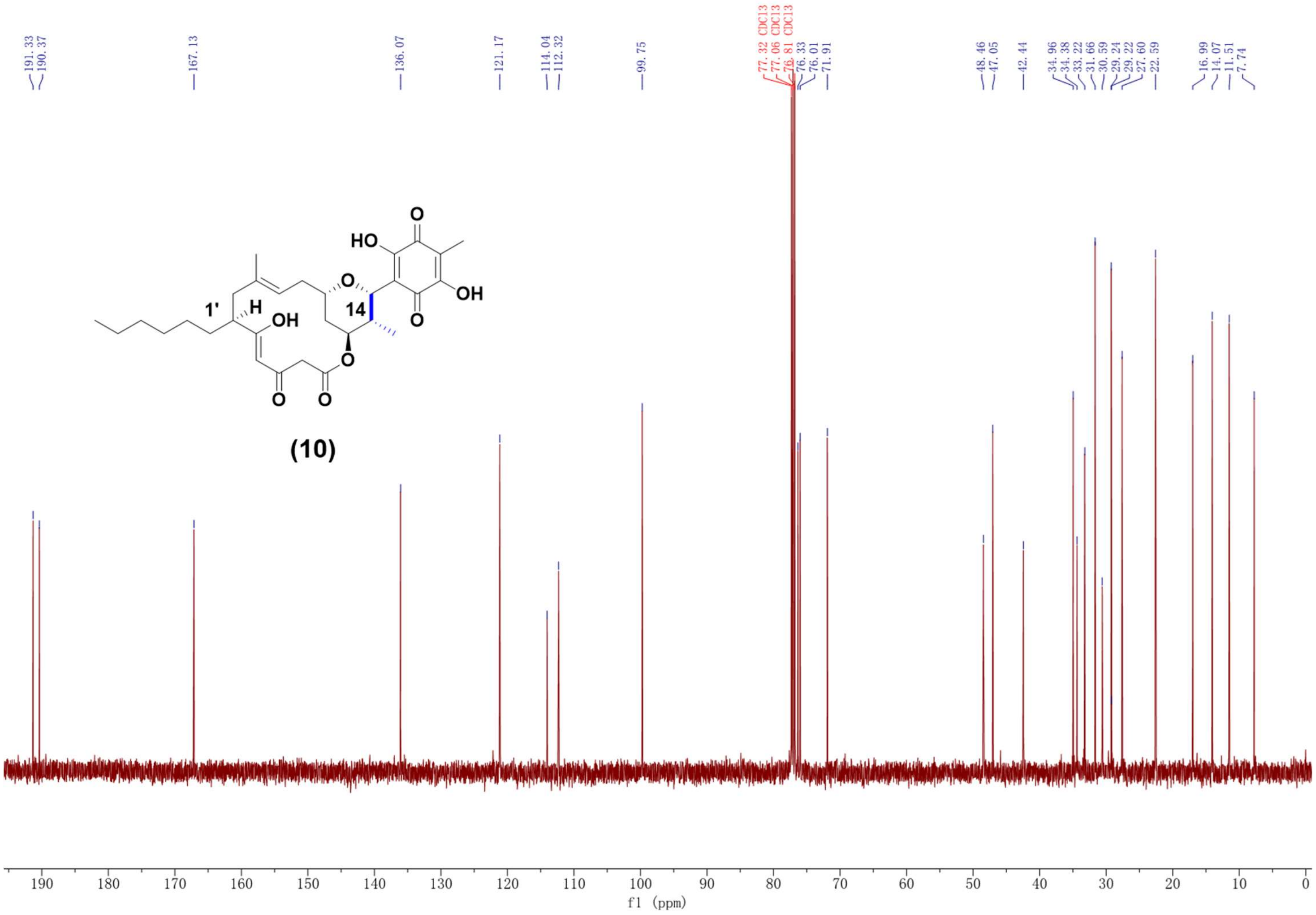

Supplementary Fig. 50. $^{13}C$ NMR spectrum (125 MHz, $CDCl_3$) of compound **10**.

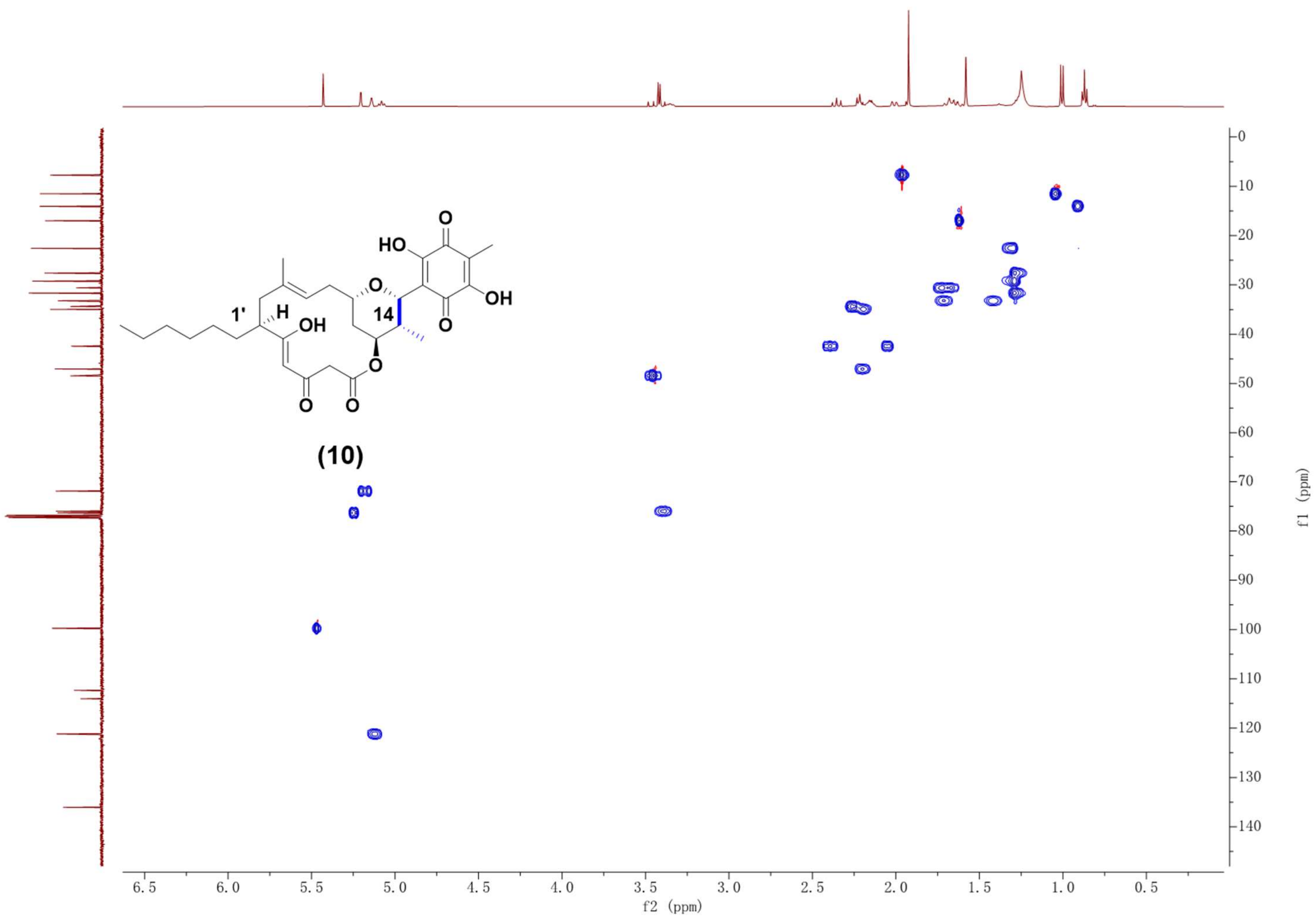

Supplementary Fig. 51. HSQC NMR spectrum (500 MHz, $CDCl_3$) of compound **10**.

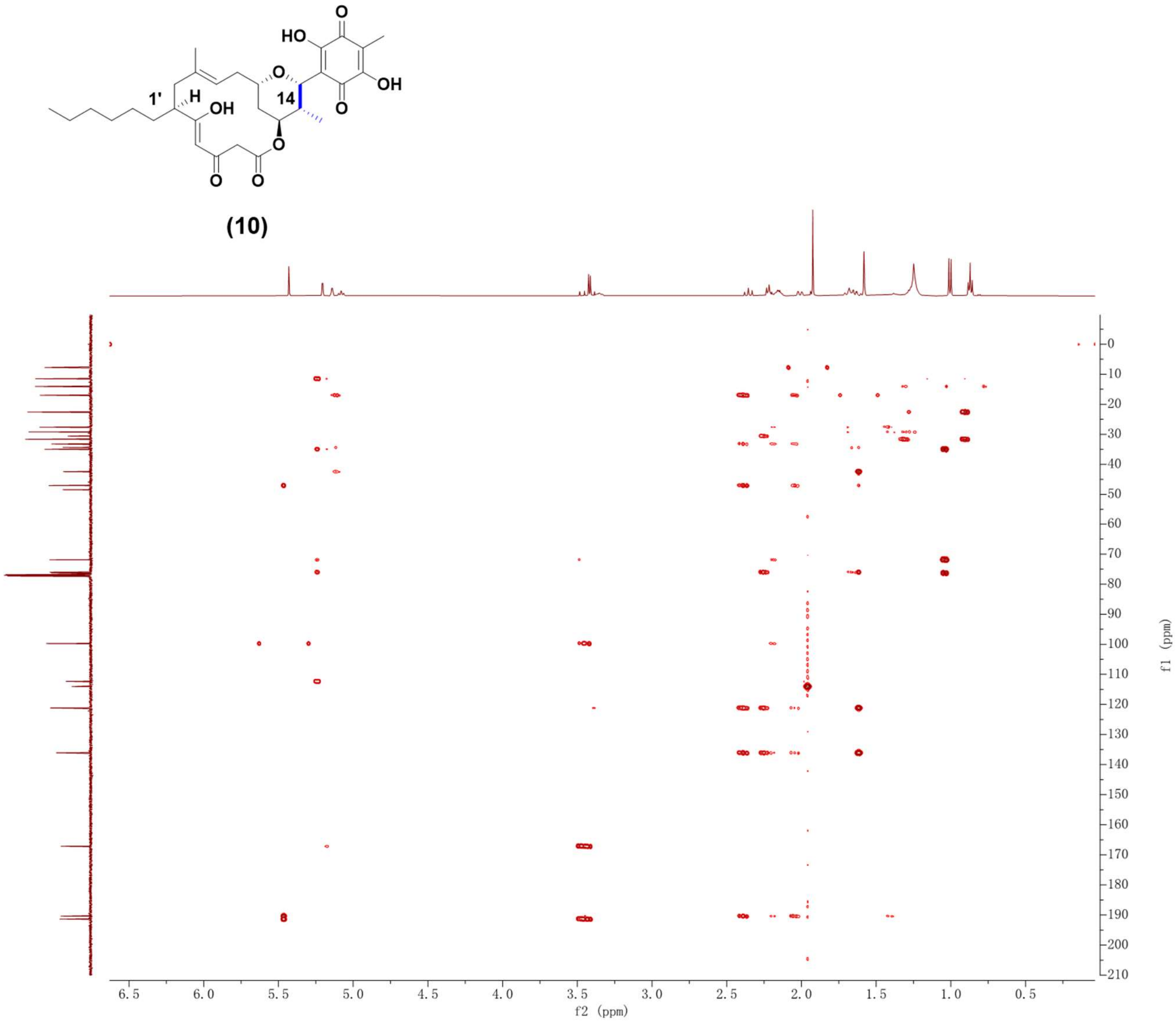

Supplementary Fig. 52. HMBC NMR spectrum (500 MHz, CDCl3) of compound **10**.

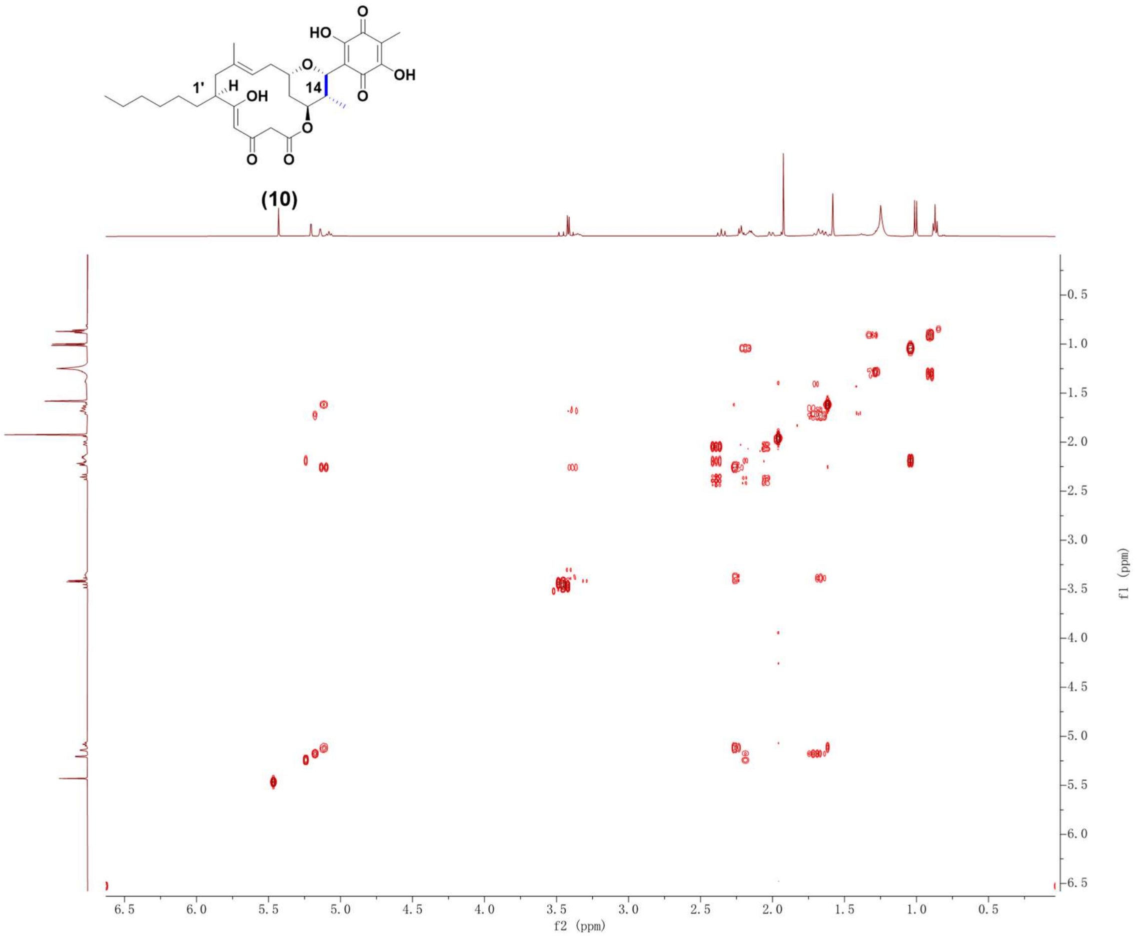

Supplementary Fig. 53. $^{1}H$-$^{1}H$ COSY NMR spectrum (500 MHz, $CDCl_3$) of compound **10**.

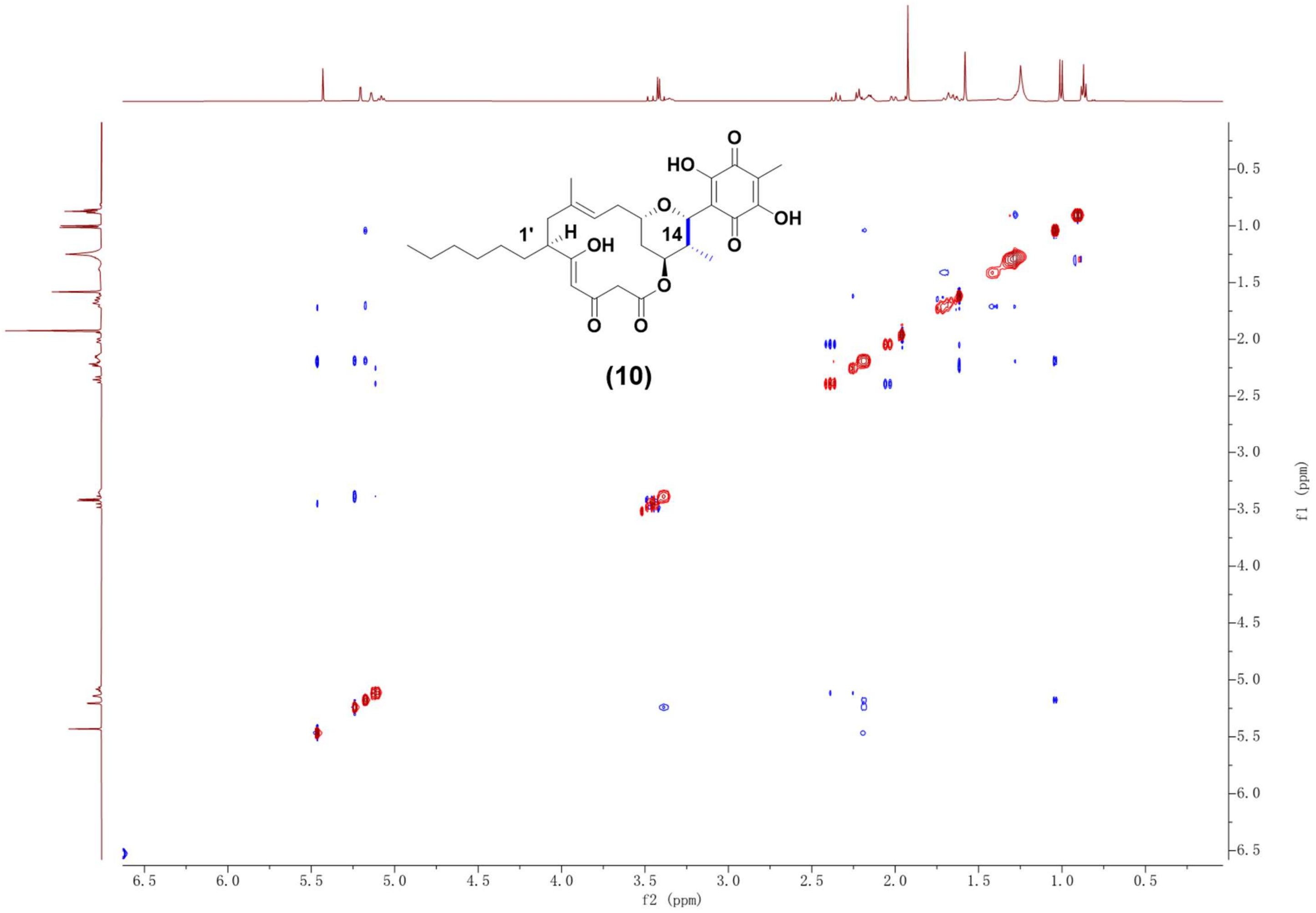

Supplementary Fig. 54. NOESY NMR spectrum (500 MHz, $CDCl_3$) of compound **10**.

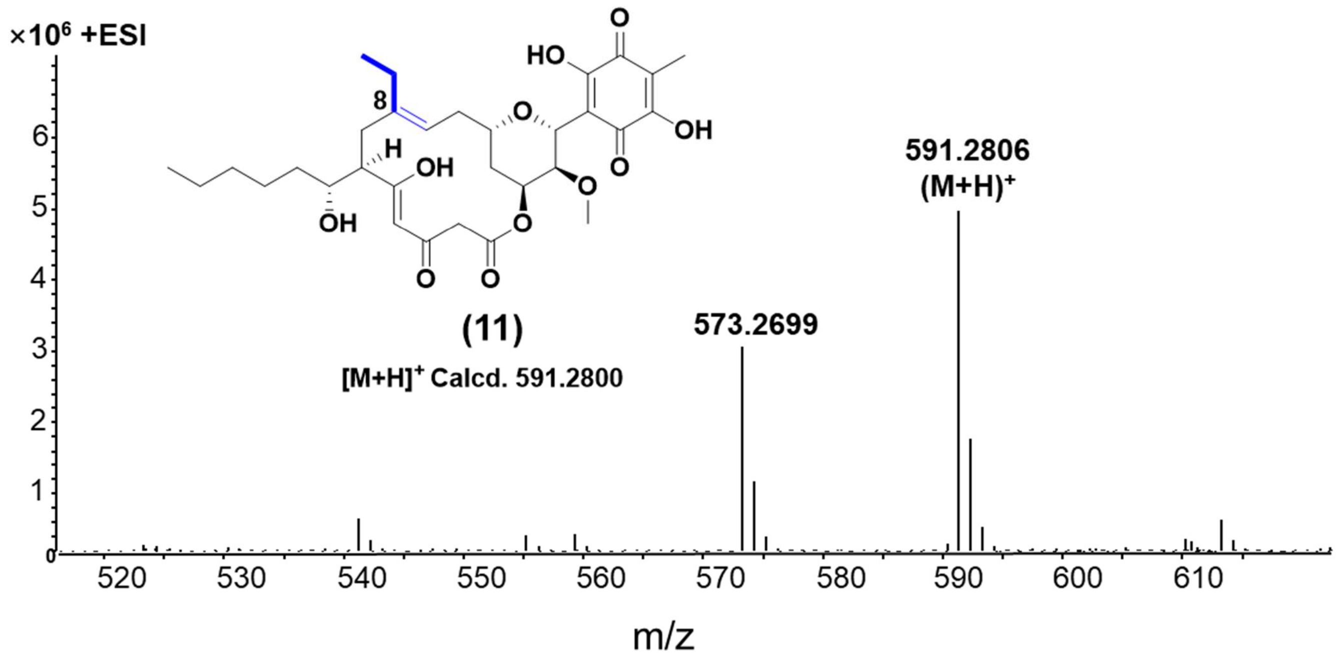

Supplementary Fig. 55. HR-ESI-MS spectrum of compound **11**.

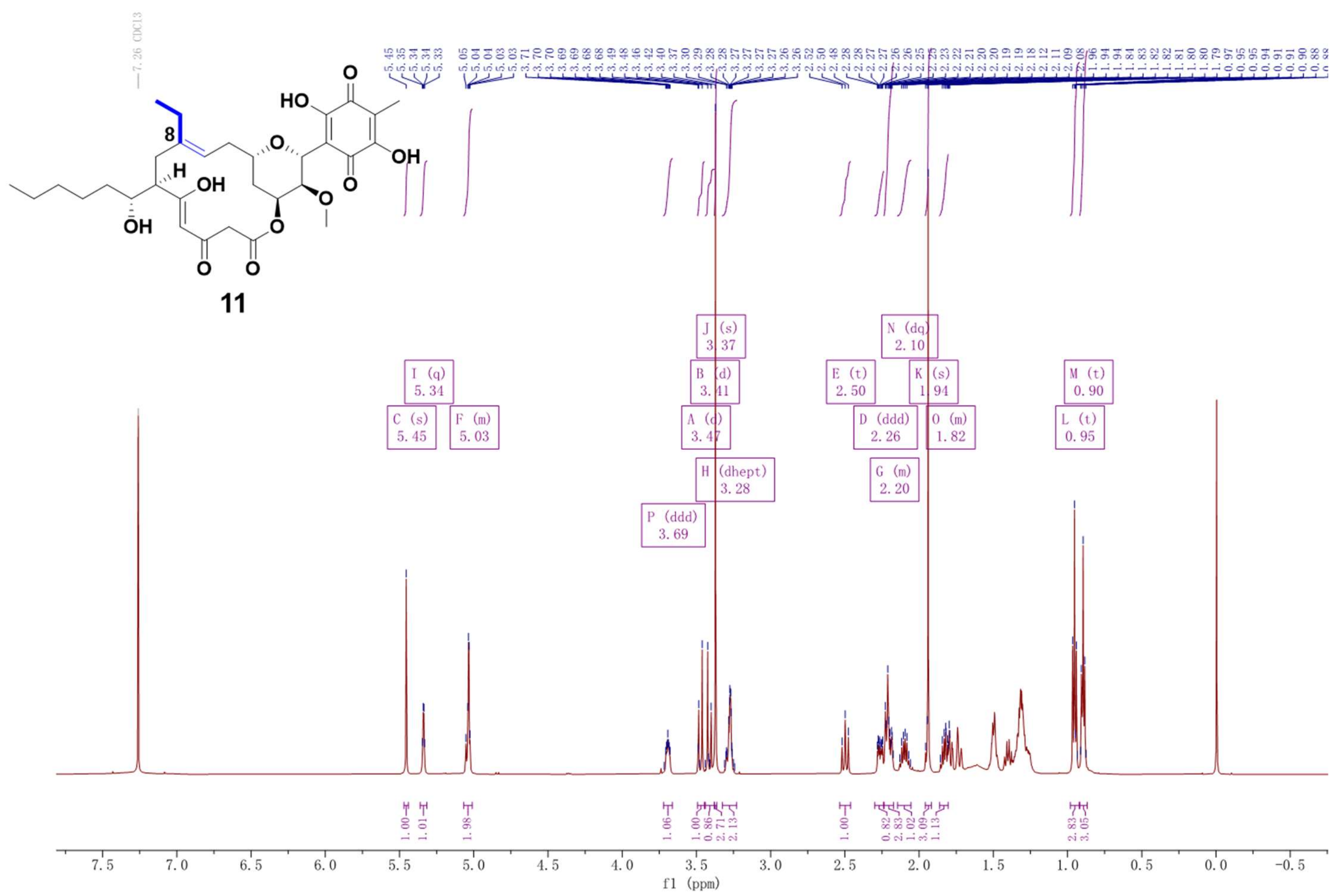

Supplementary Fig. 56. $^1$H NMR spectrum (600 MHz, $CDCl_3$) of compound **11**.

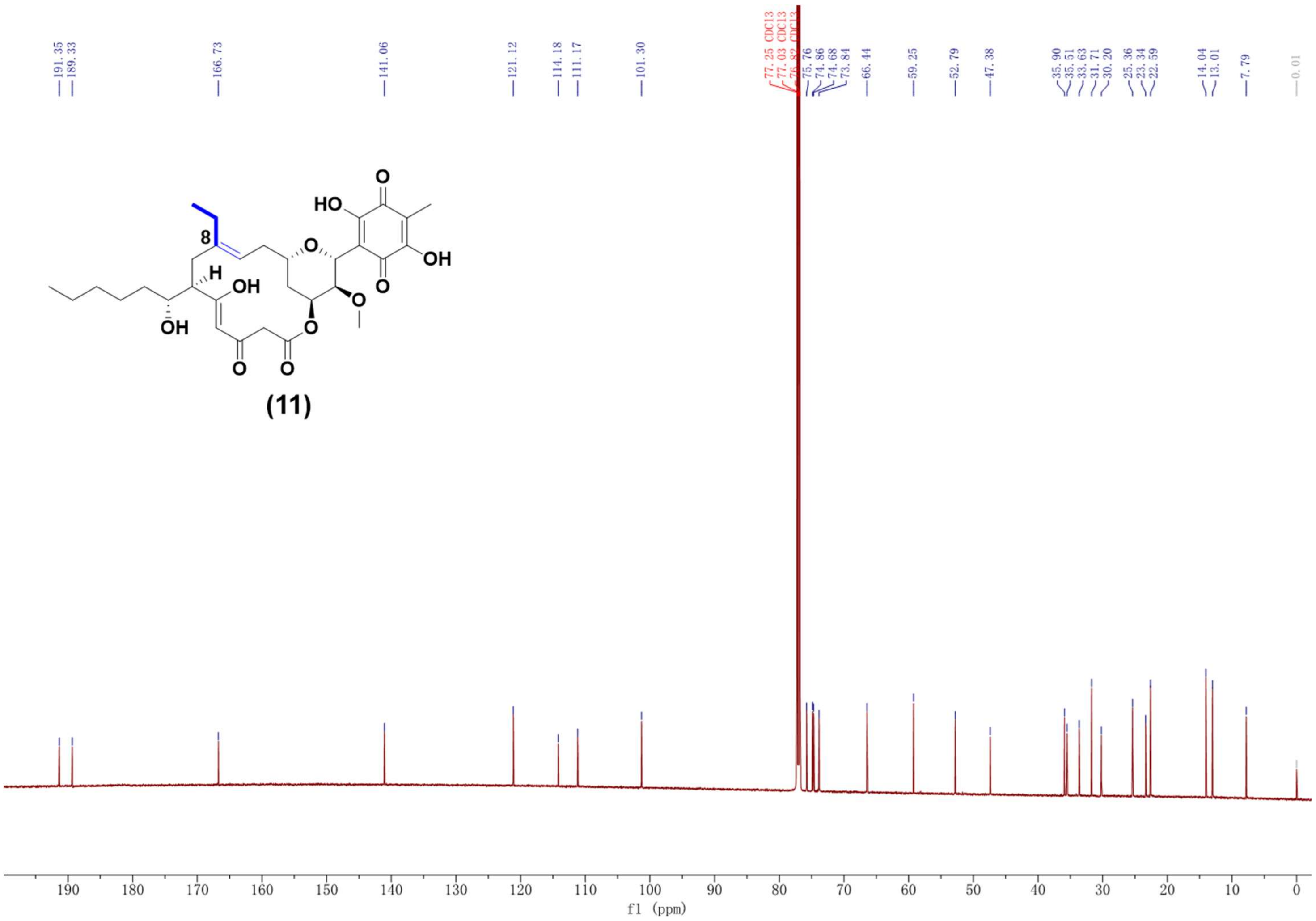

Supplementary Fig. 57. $^{13}C$ NMR spectrum (150 MHz, $CDCl_3$) of compound **11**.

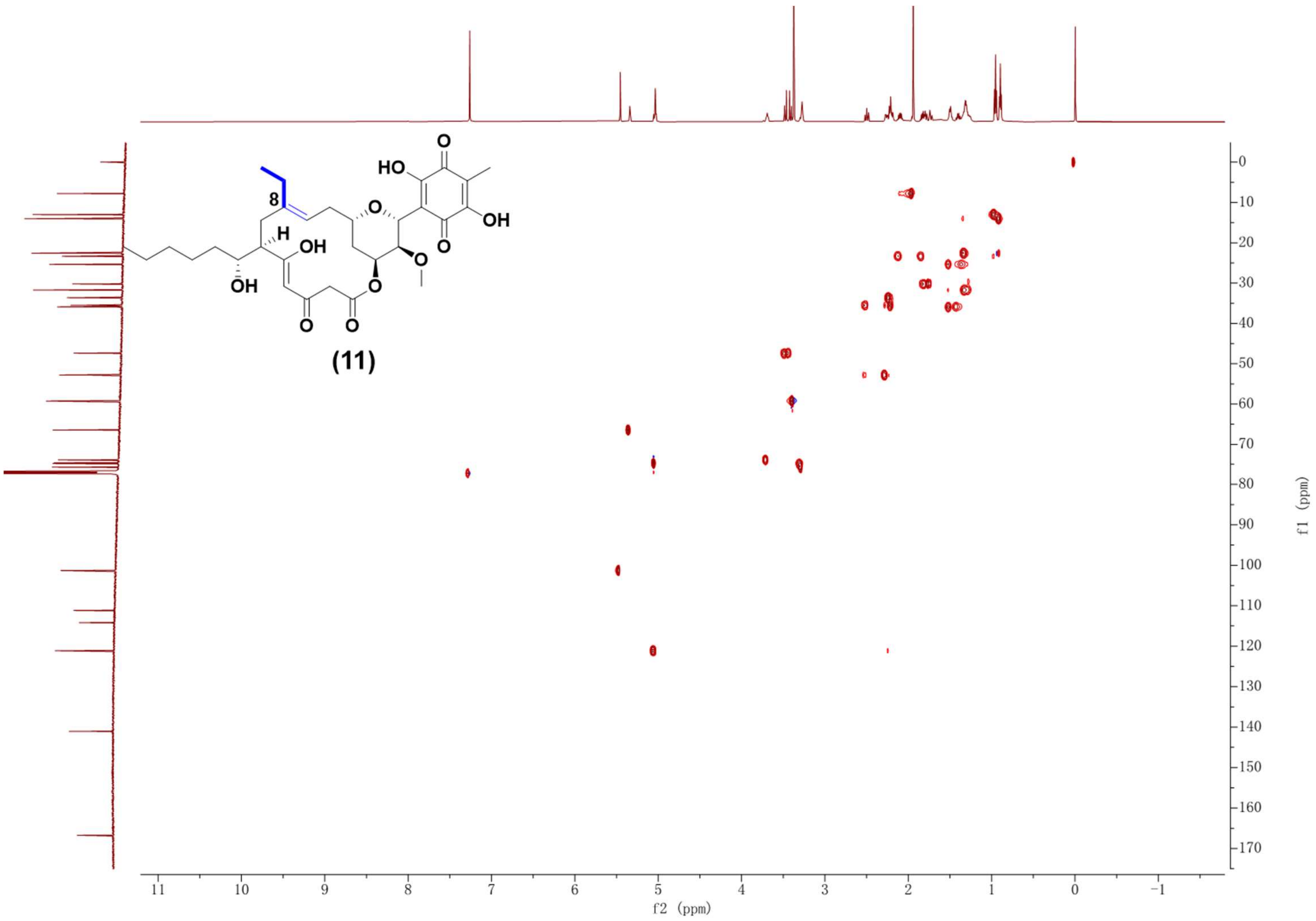

Supplementary Fig. 58. HSQC NMR spectrum (600 MHz, $CDCl_3$) of compound **11**.

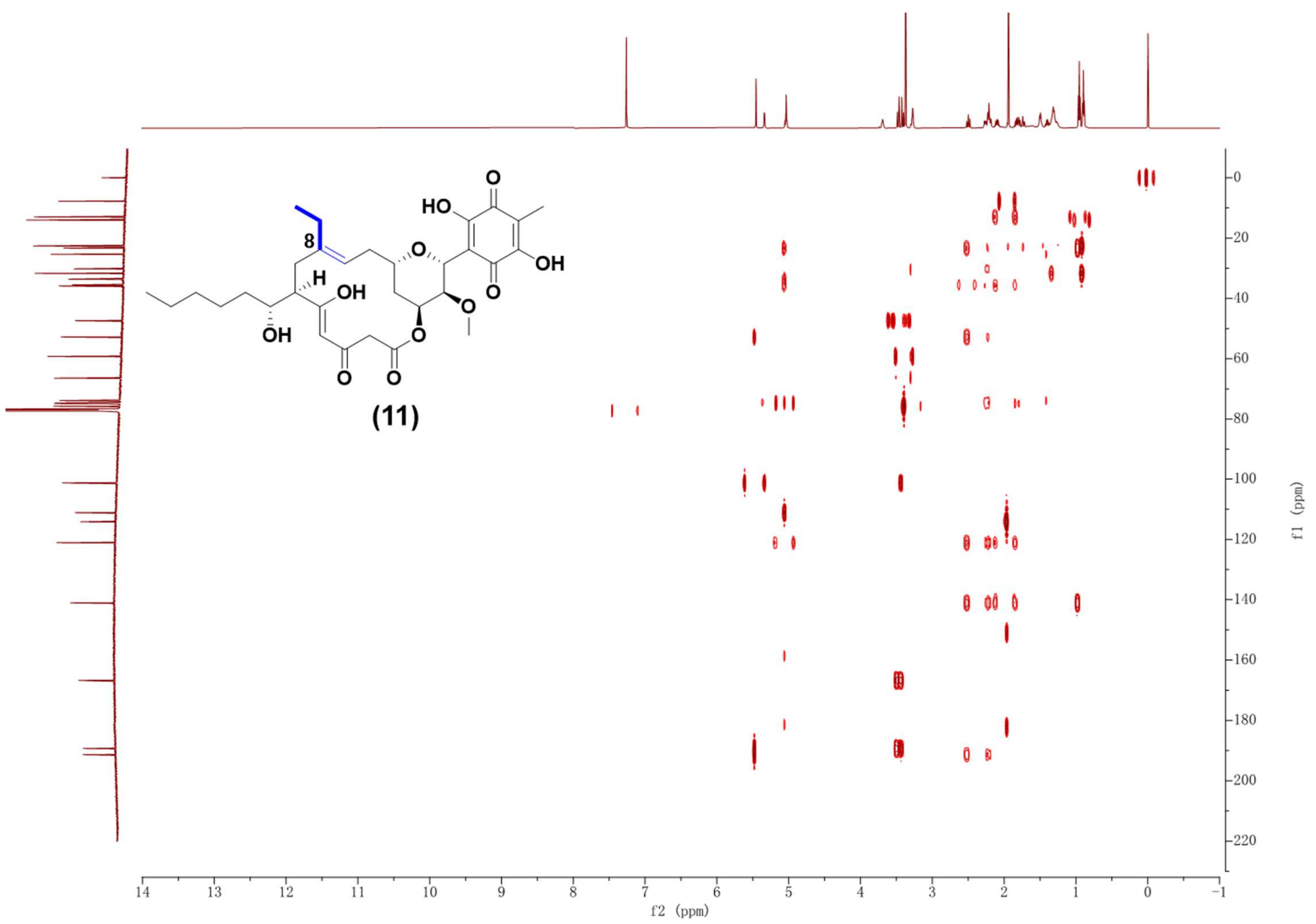

Supplementary Fig. 59. HMBC NMR spectrum (600 MHz, $CDCl_3$) of compound **11**.

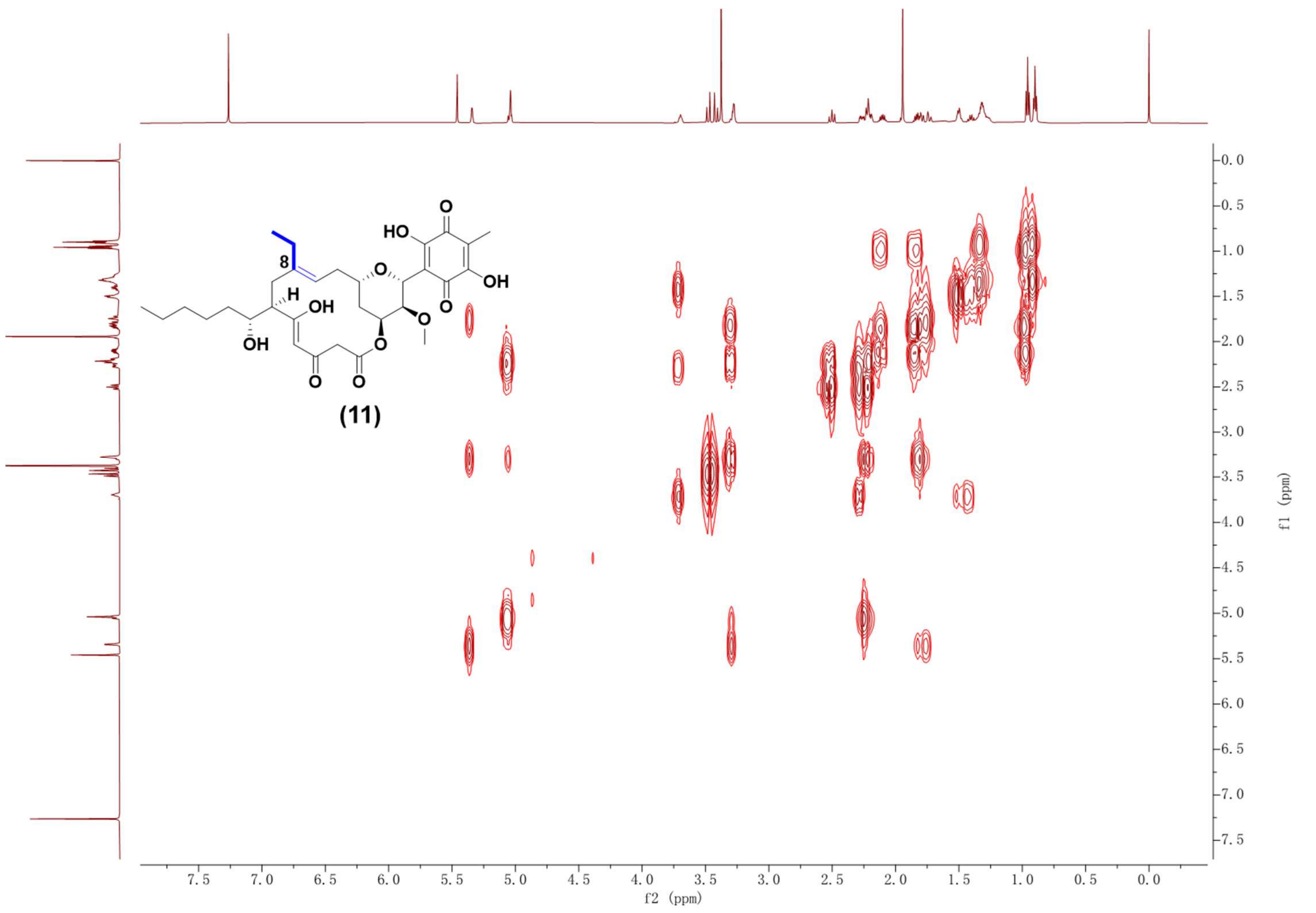

Supplementary Fig. 60. $^1H$-$^1H$ COSY NMR spectrum (600 MHz, $CDCl_3$) of compound **11**.

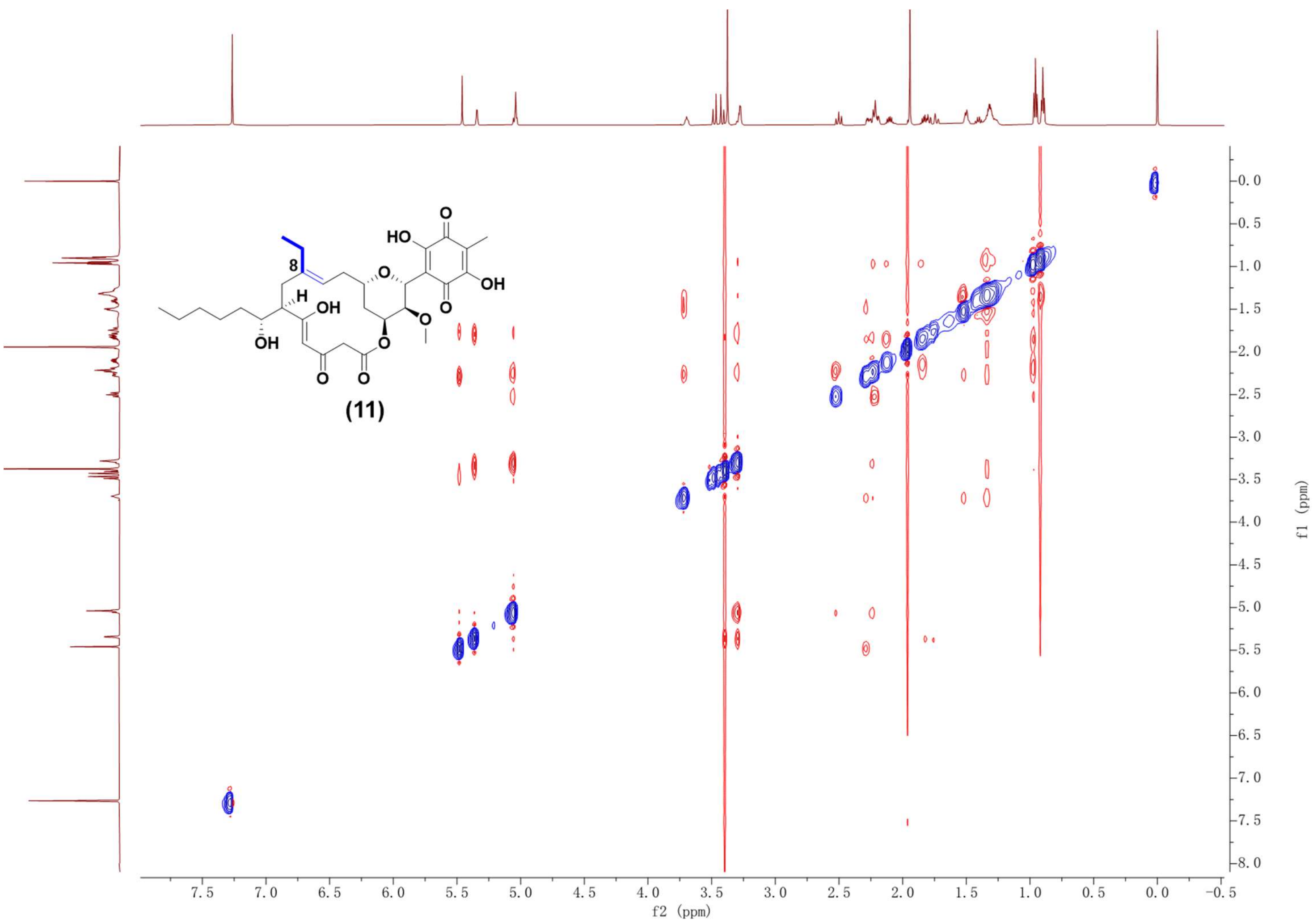

Supplementary Fig. 61. NOESY NMR spectrum (600 MHz, $CDCl_3$) of compound **11**.

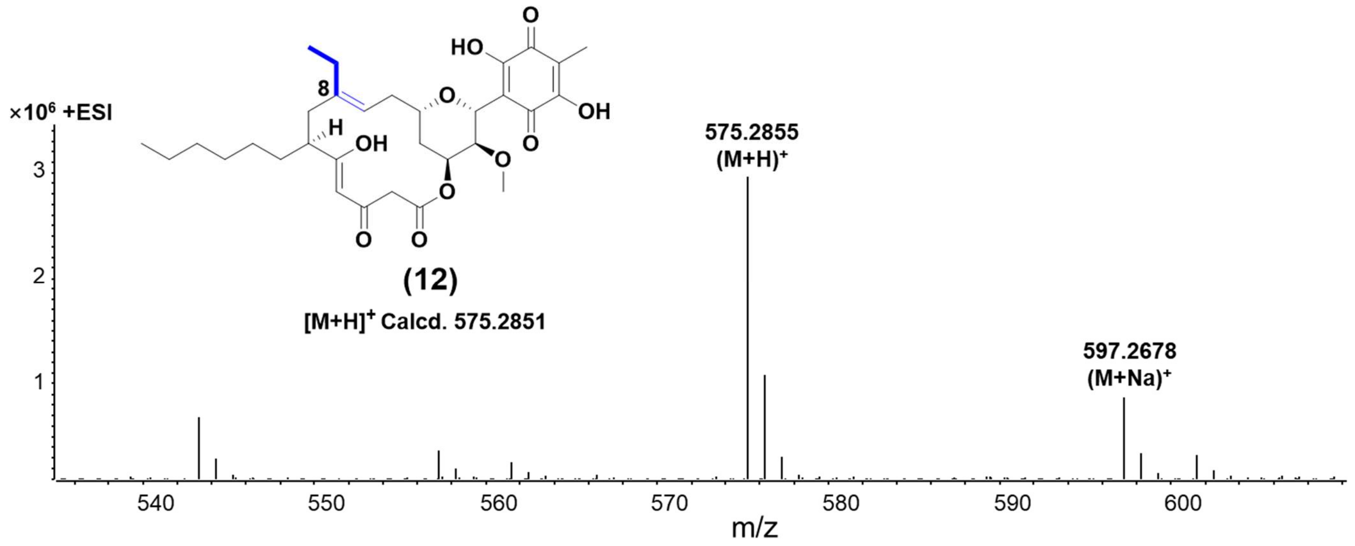

Supplementary Fig. 62. HR-ESI-MS spectrum of compound **12**.

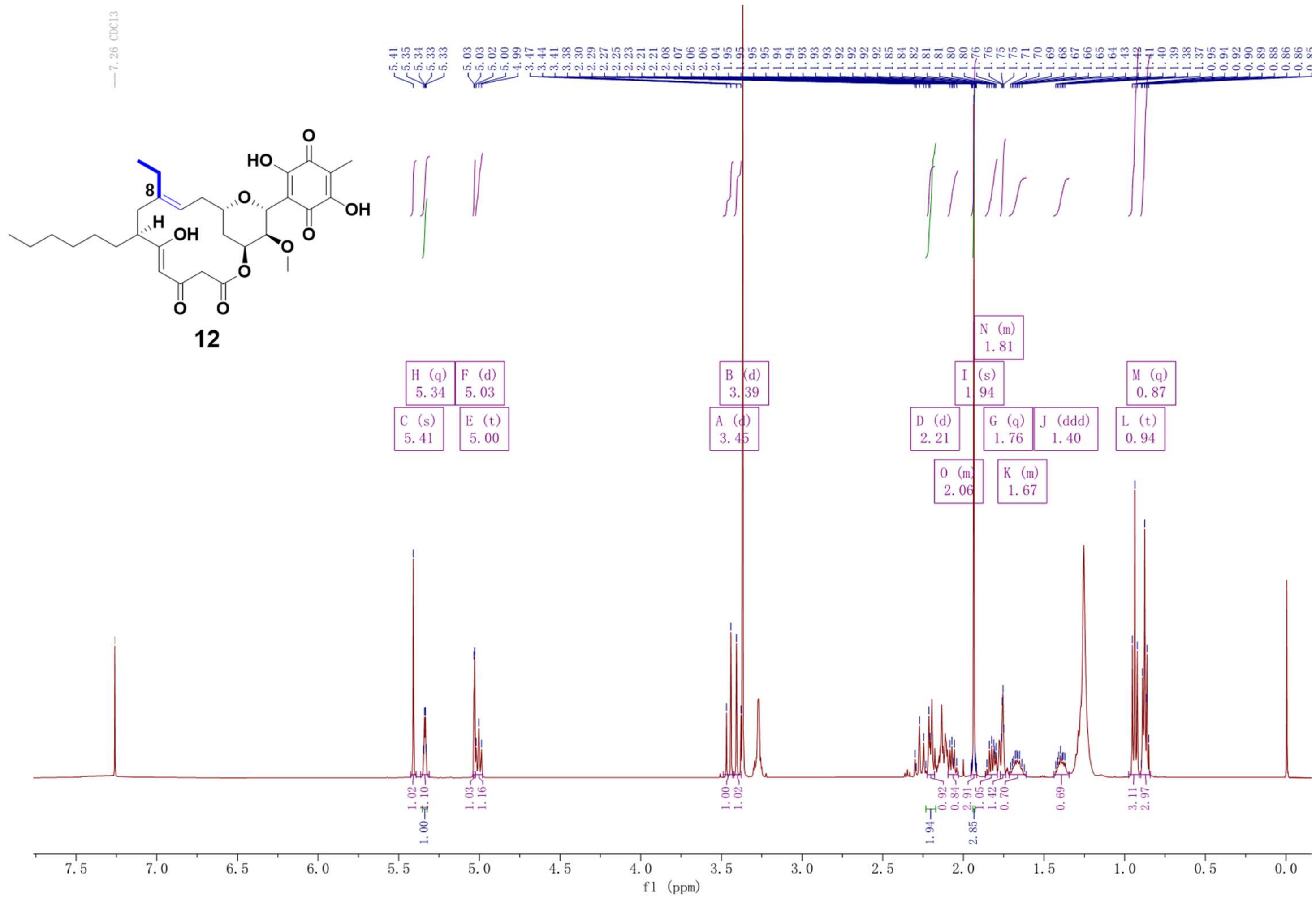

Supplementary Fig. 63. $^{1}H$ NMR spectrum (500 MHz, $CDCl_3$) of compound **12**.

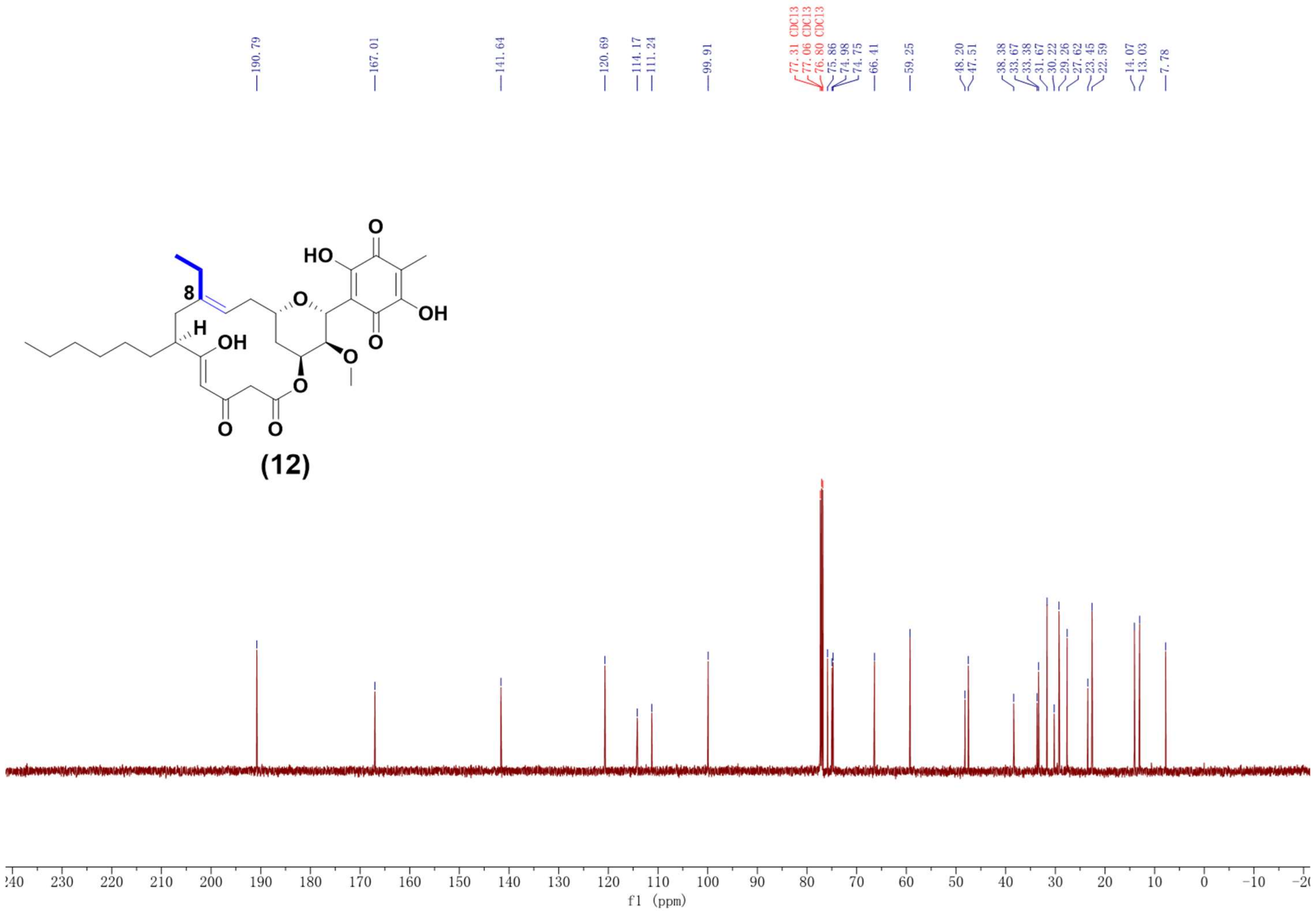

Supplementary Fig. 64. $^{13}C$ NMR spectrum (125 MHz, $CDCl_3$) of compound **12**.

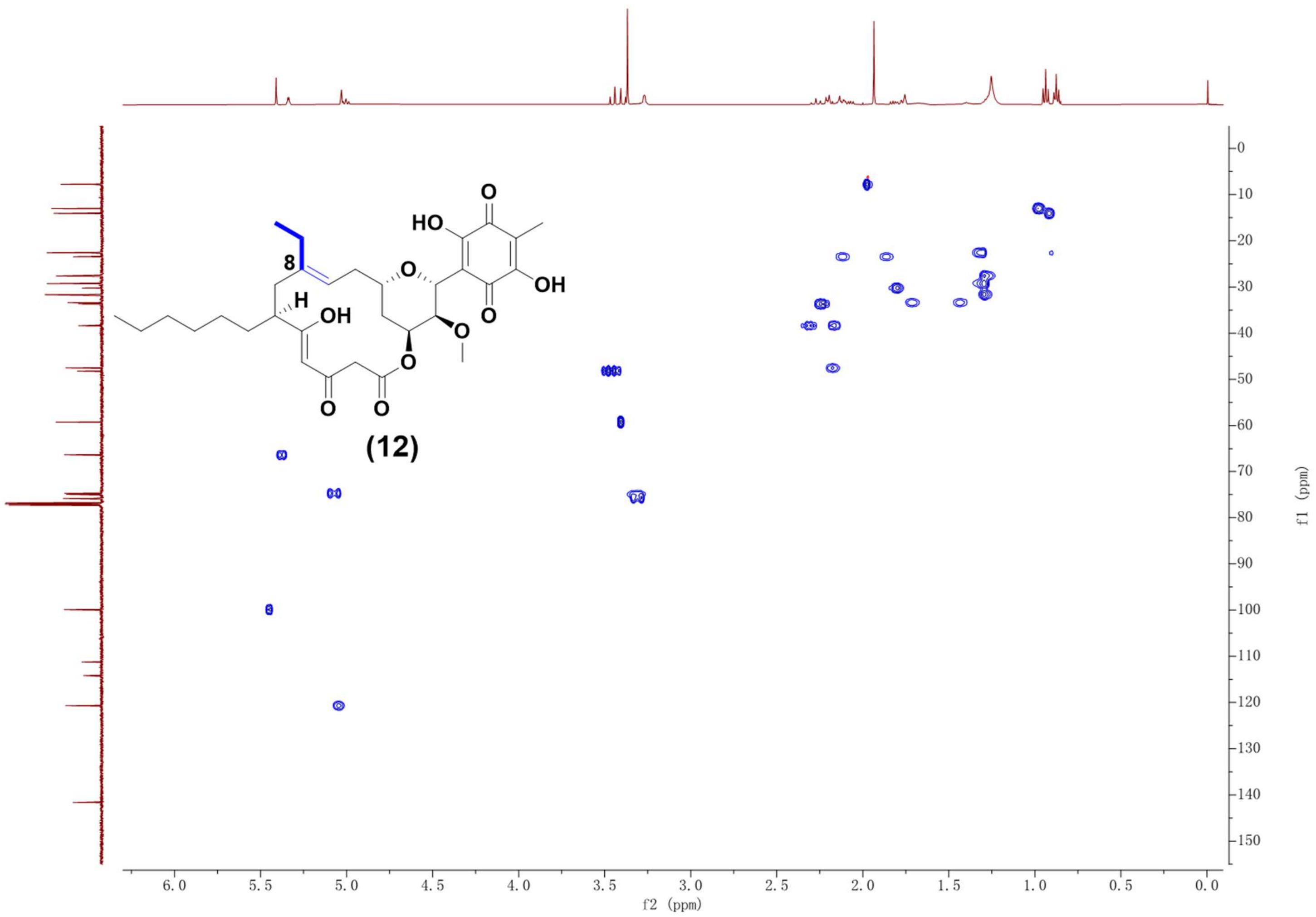

Supplementary Fig. 65. HSQC NMR spectrum (500 MHz, $CDCl_3$) of compound **12**.

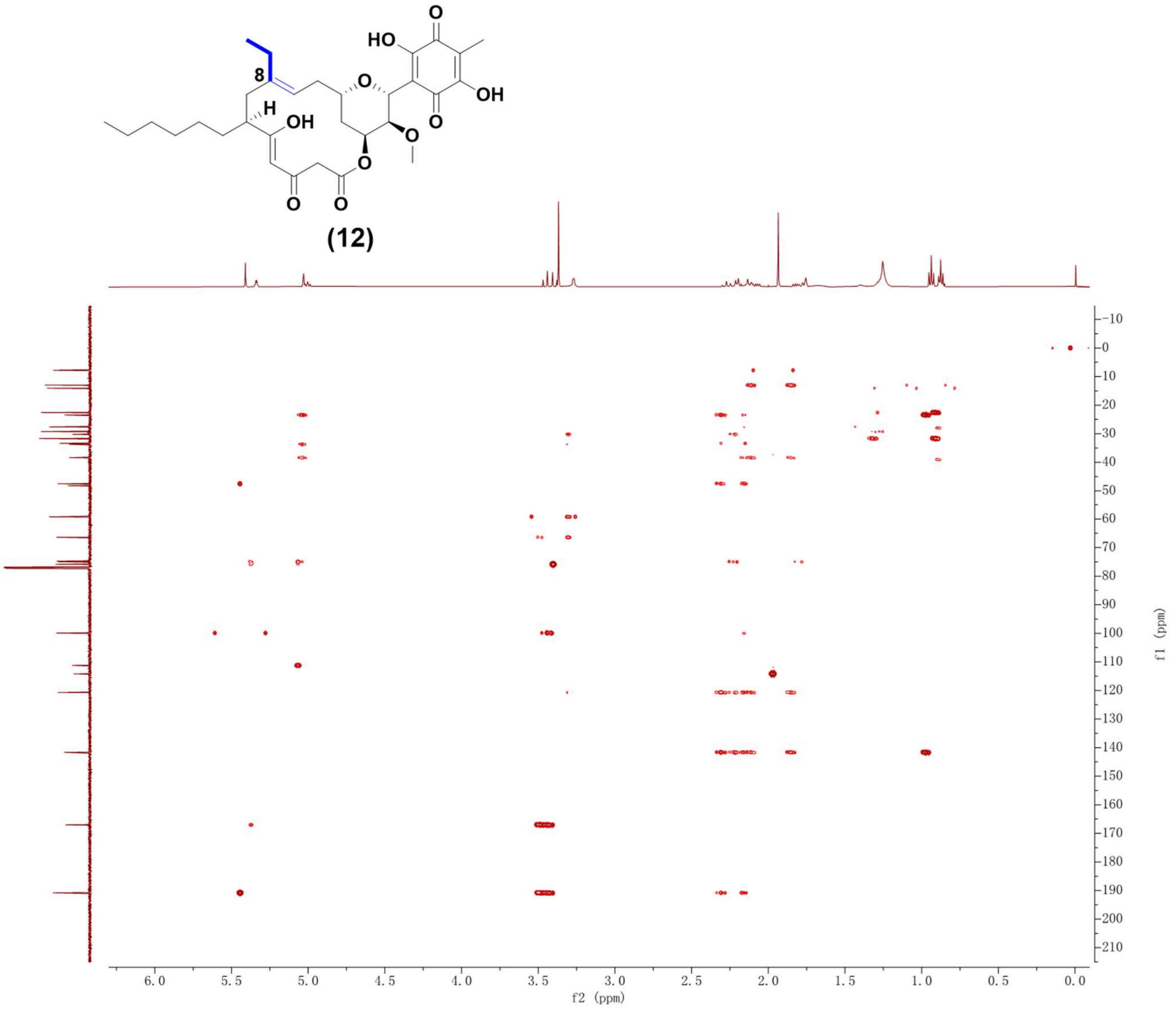

Supplementary Fig. 66. HMBC NMR spectrum (500 MHz, $CDCl_3$) of compound **12**.

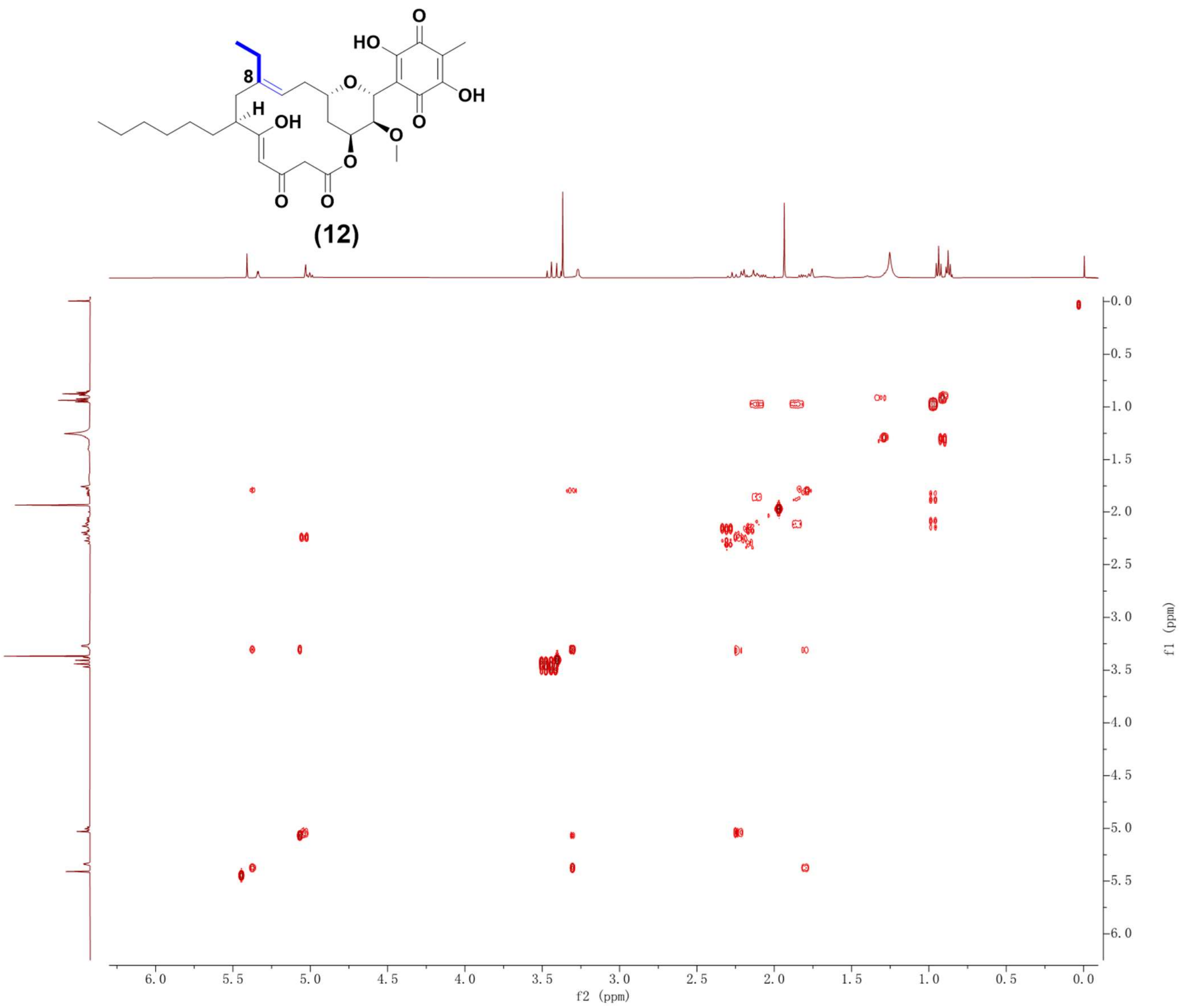

Supplementary Fig. 67. $^{1}H$-$^{1}H$ COSY NMR spectrum (500 MHz, $CDCl_3$) of compound **12**.

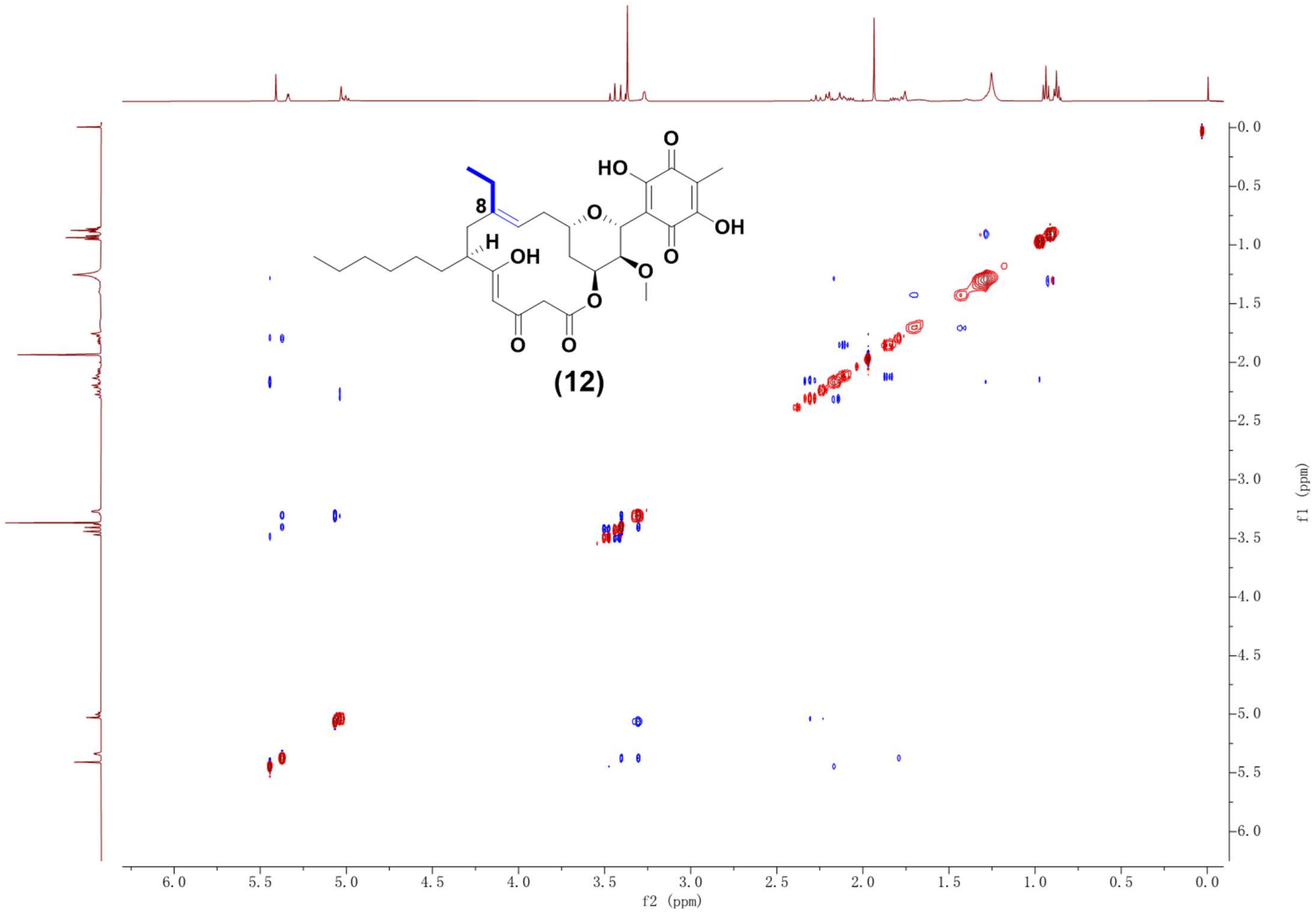

Supplementary Fig. 68. NOESY NMR spectrum (500 MHz, $CDCl_3$) of compound **12**.

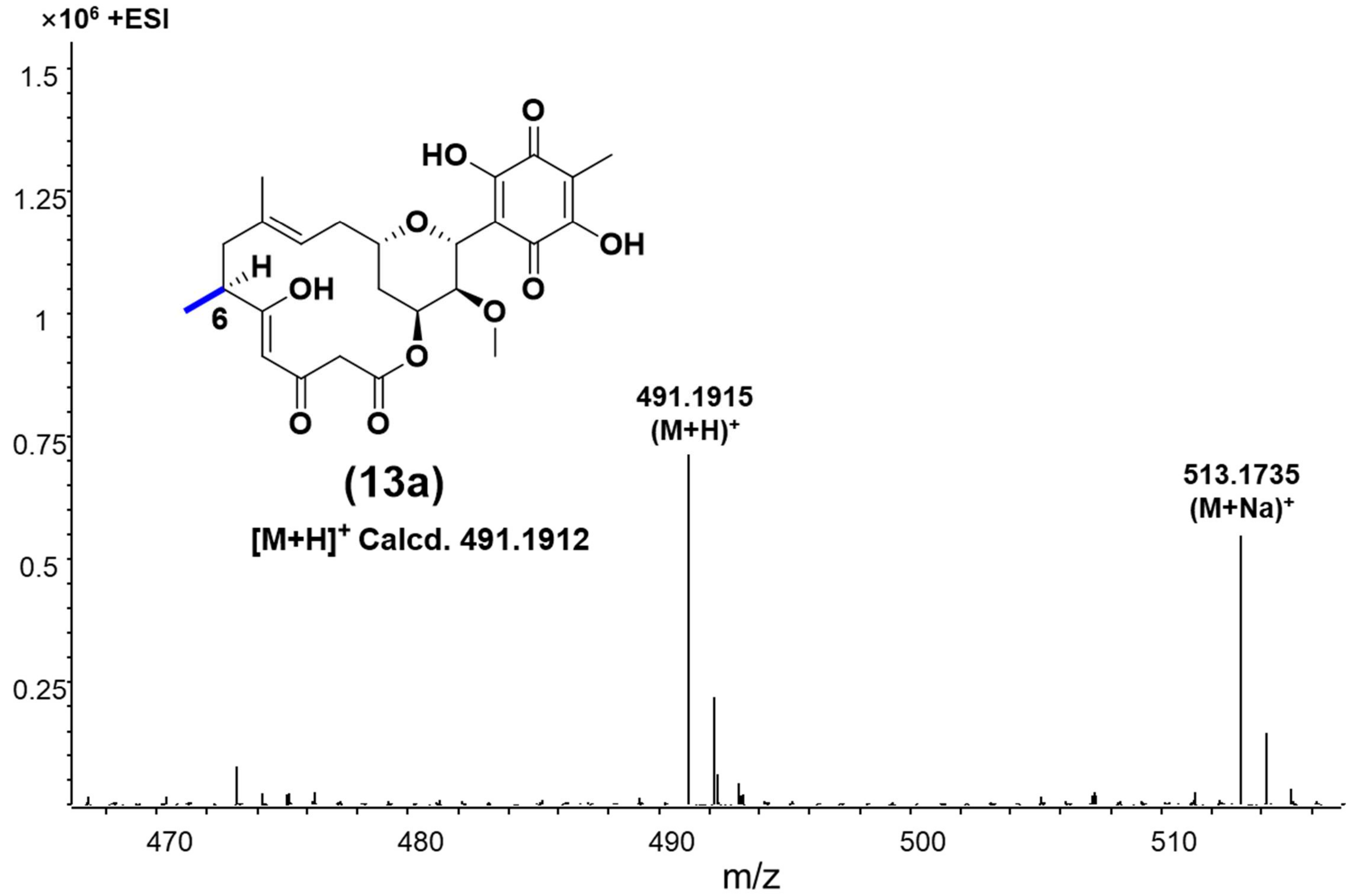

Supplementary Fig. 69. HR-ESI-MS spectrum of compound **13a**.

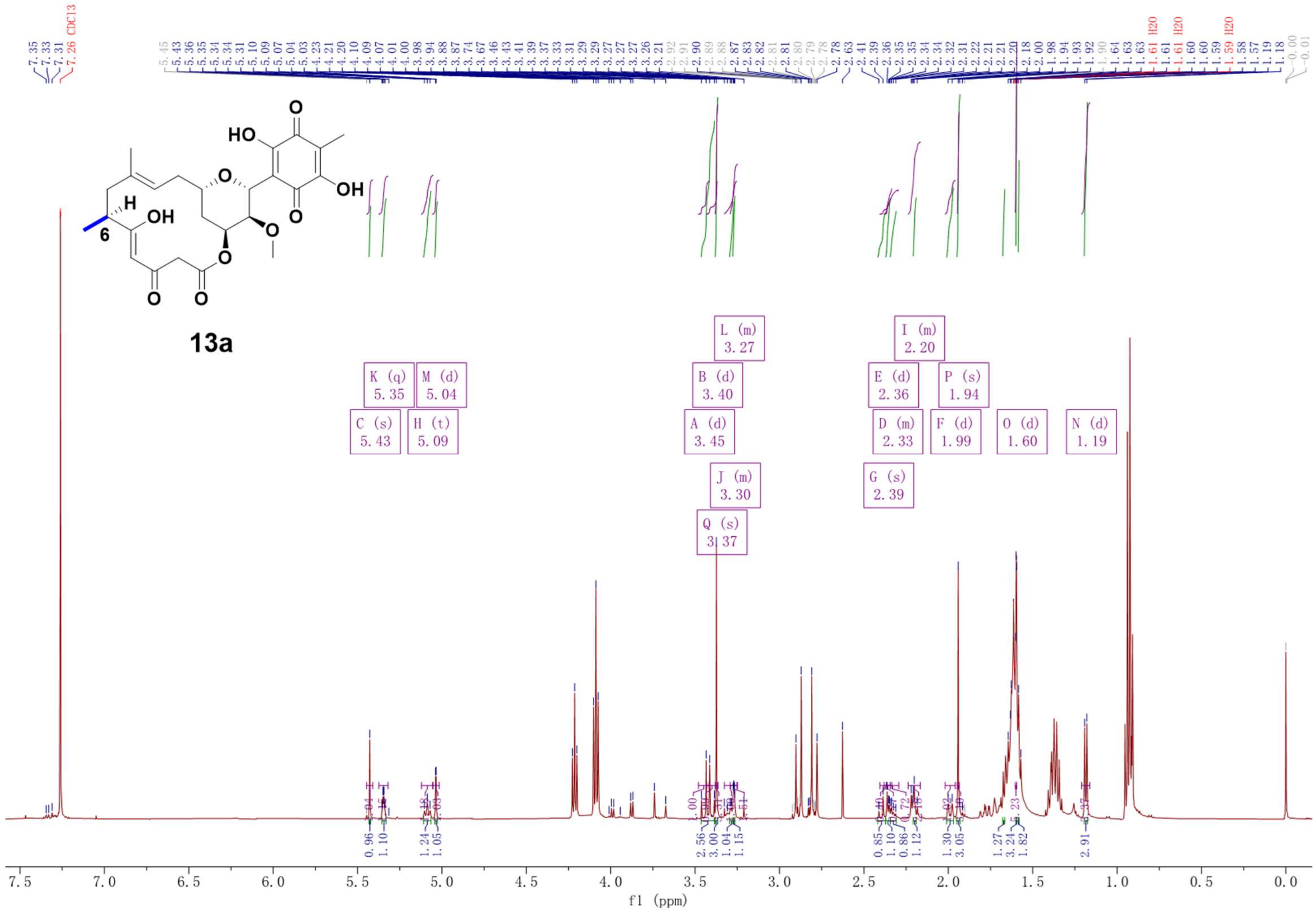

Supplementary Fig. 70. $^{1}$H NMR spectrum (600 MHz, $CDCl_3$) of compound **13a**.

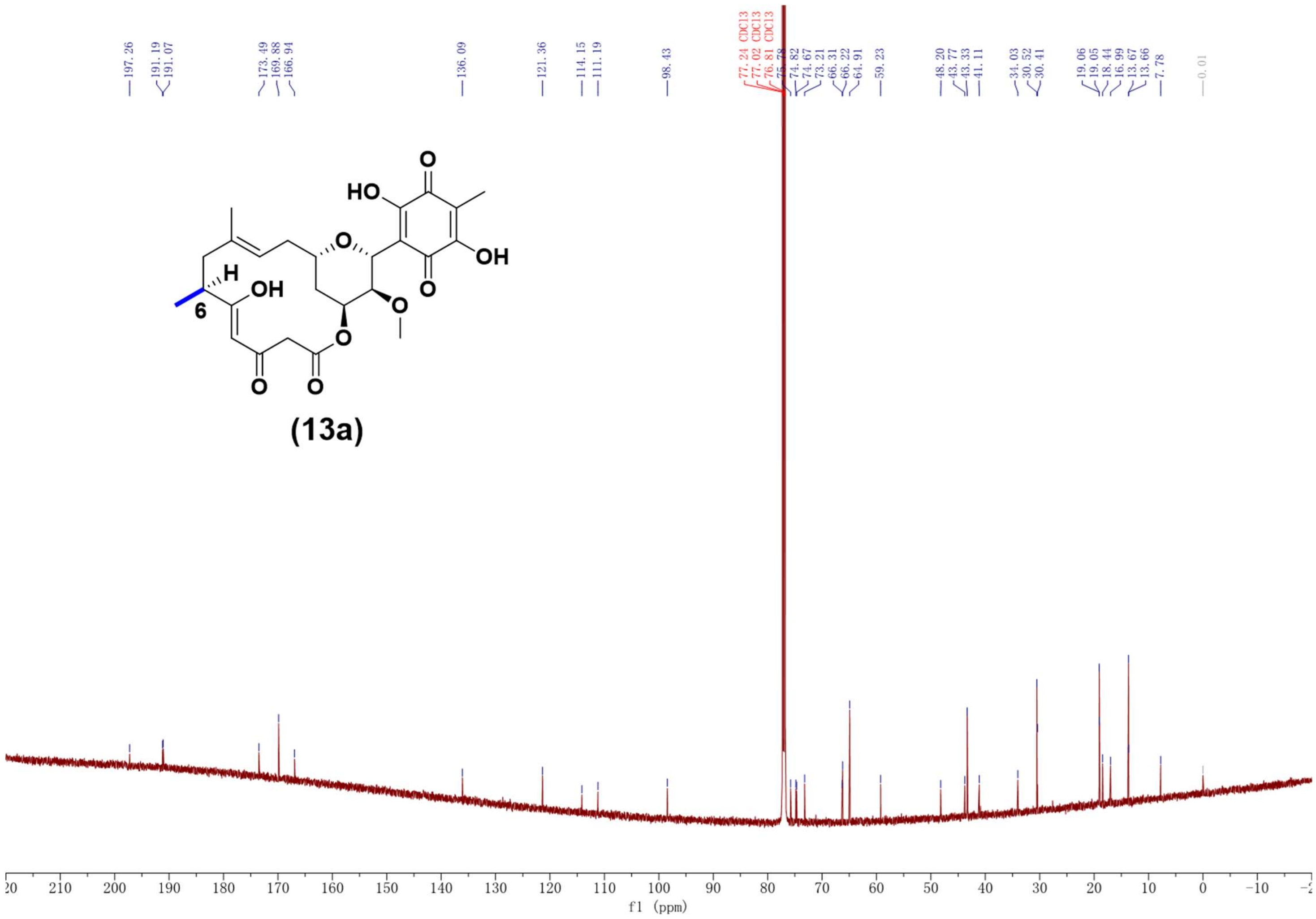

Supplementary Fig. 71. $^{13}C$ NMR spectrum (150 MHz, $CDCl_3$) of compound **13a**.

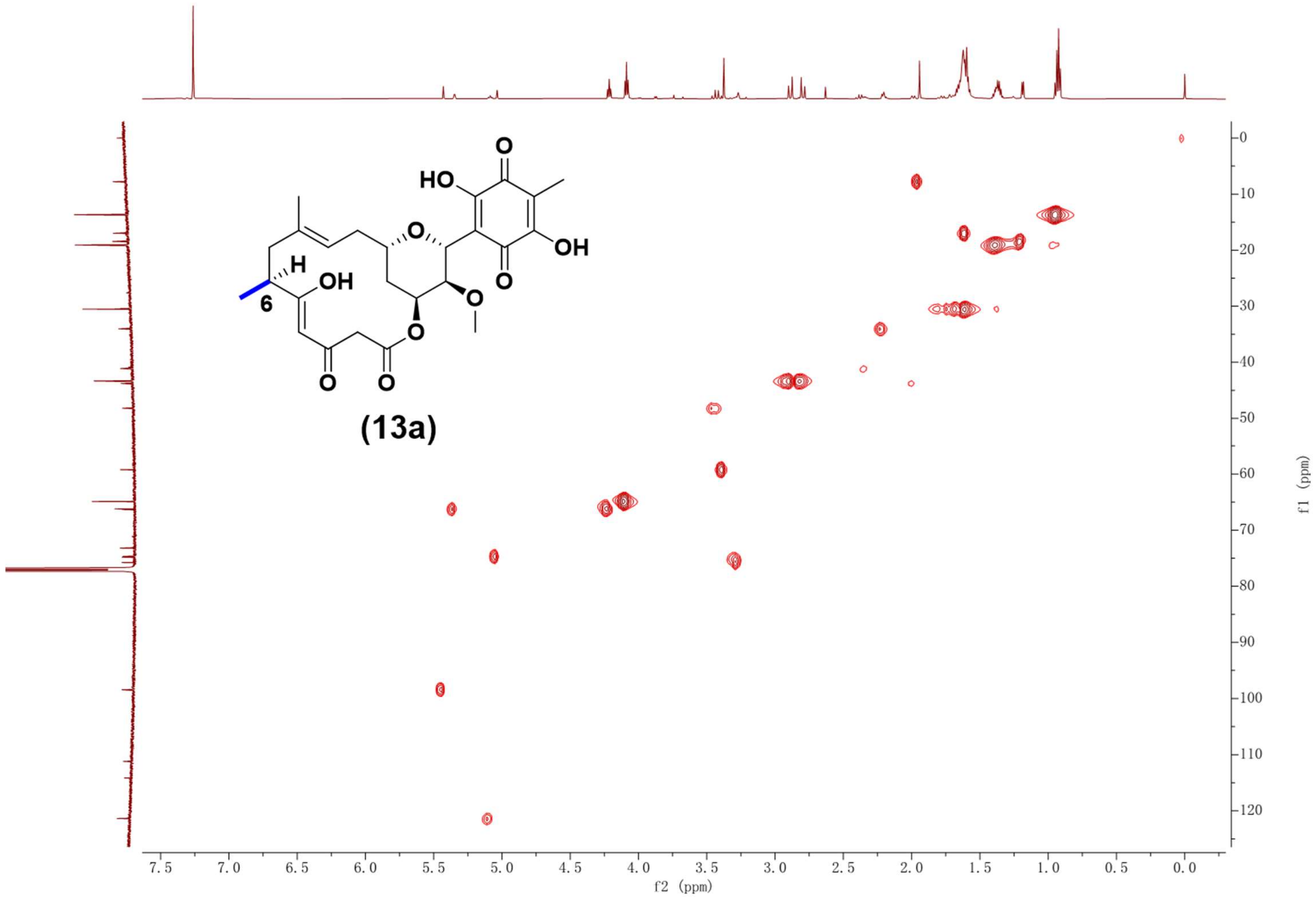

Supplementary Fig. 72. HSQC NMR spectrum (600 MHz, $CDCl_3$) of compound **13a**.

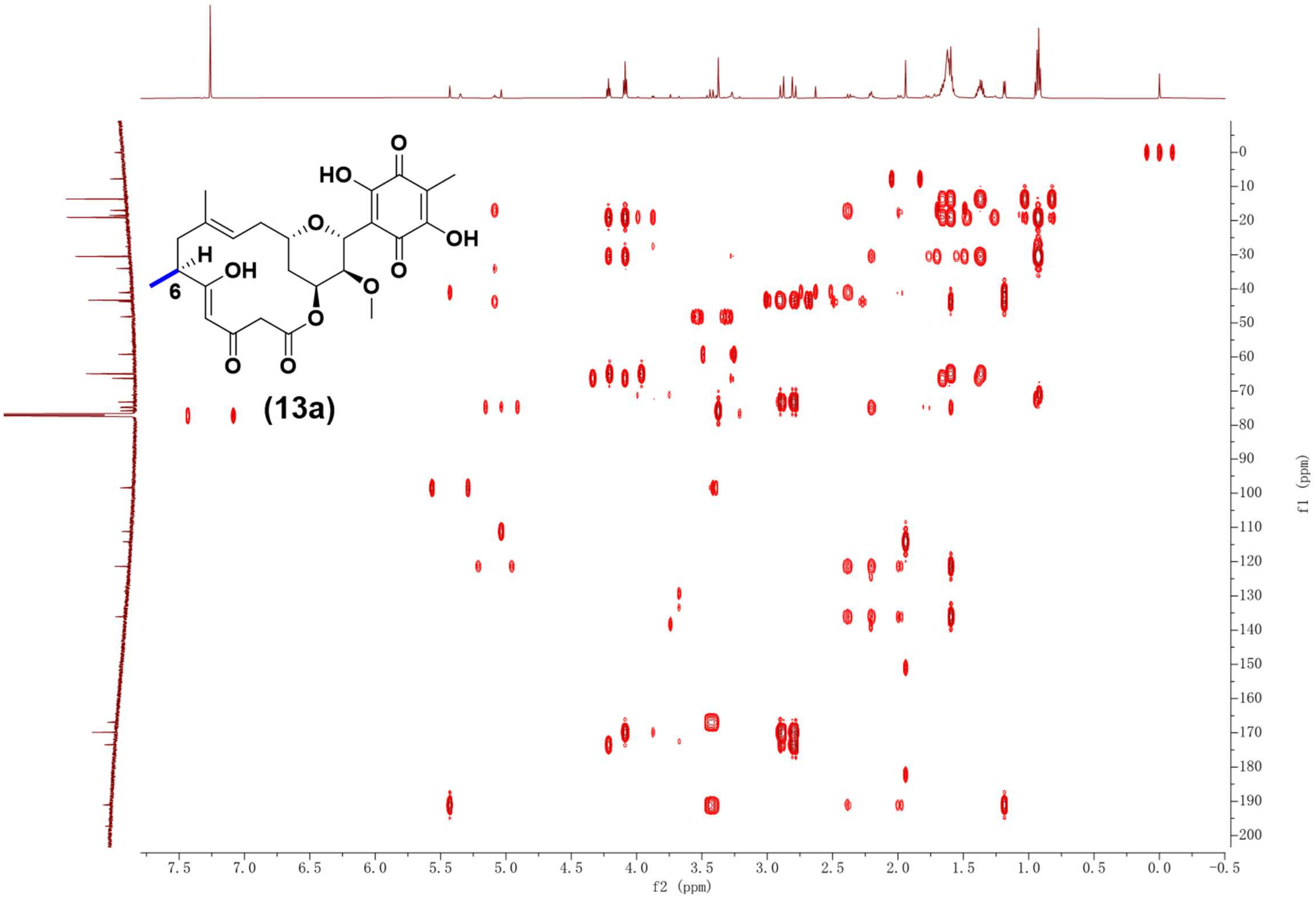

Supplementary Fig. 73. HMBC NMR spectrum (600 MHz, $CDCl_3$) of compound **13a**.

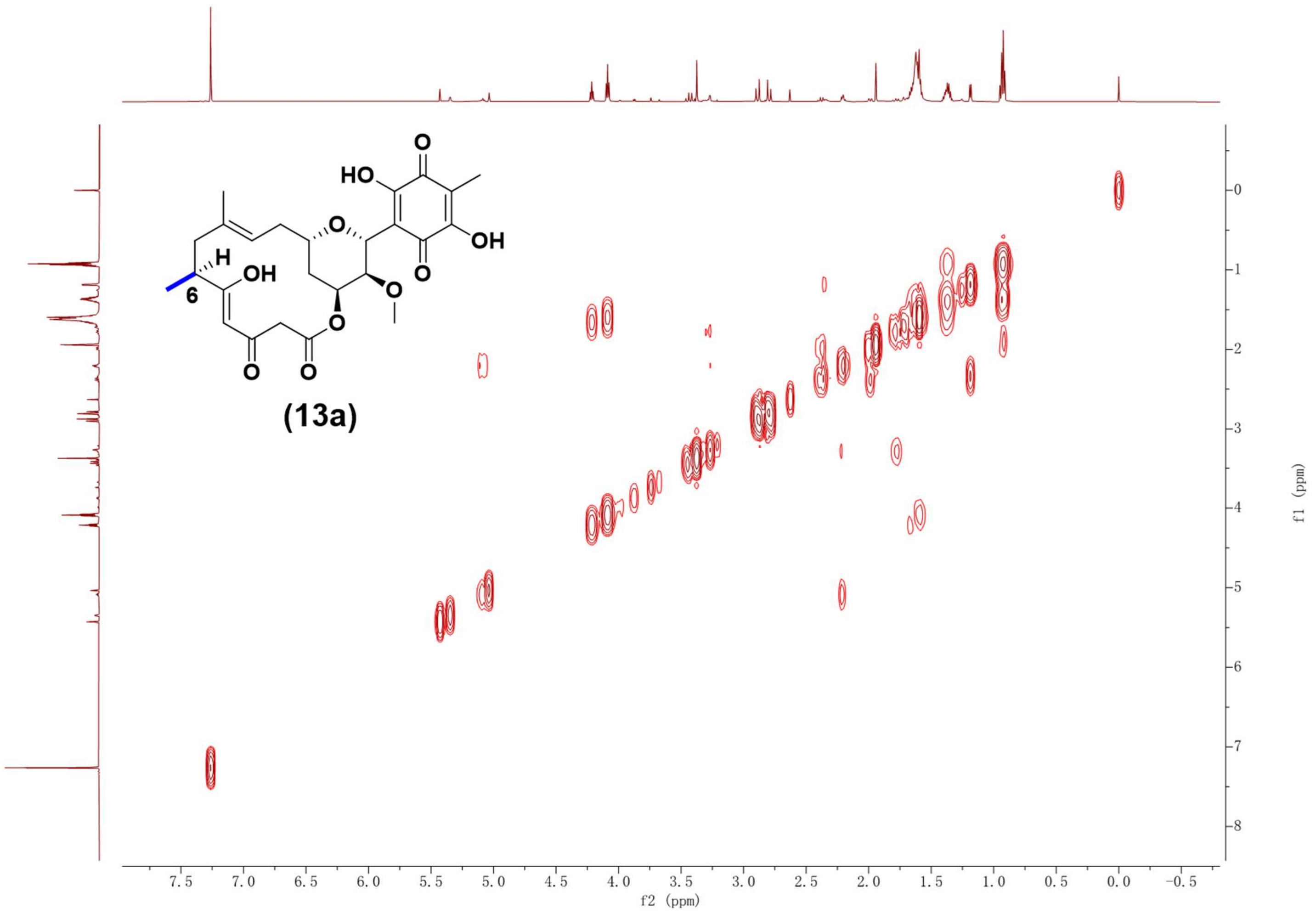

Supplementary Fig. 74. $^1H$-$^1H$ COSY NMR spectrum (600 MHz, $CDCl_3$) of compound **13a**.

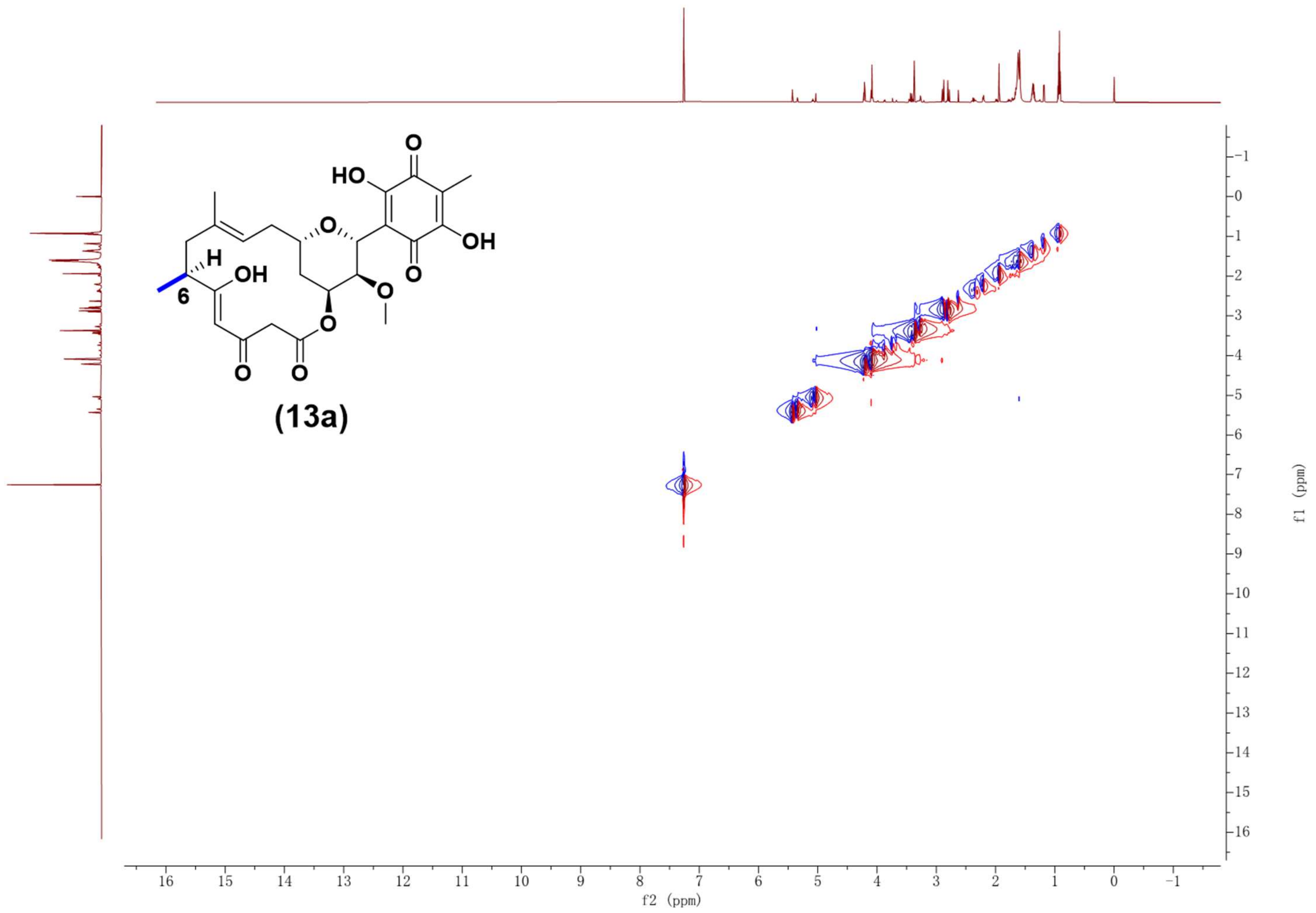

Supplementary Fig. 75. ROESY NMR spectrum (600 MHz, $CDCl_3$) of compound **13a**.

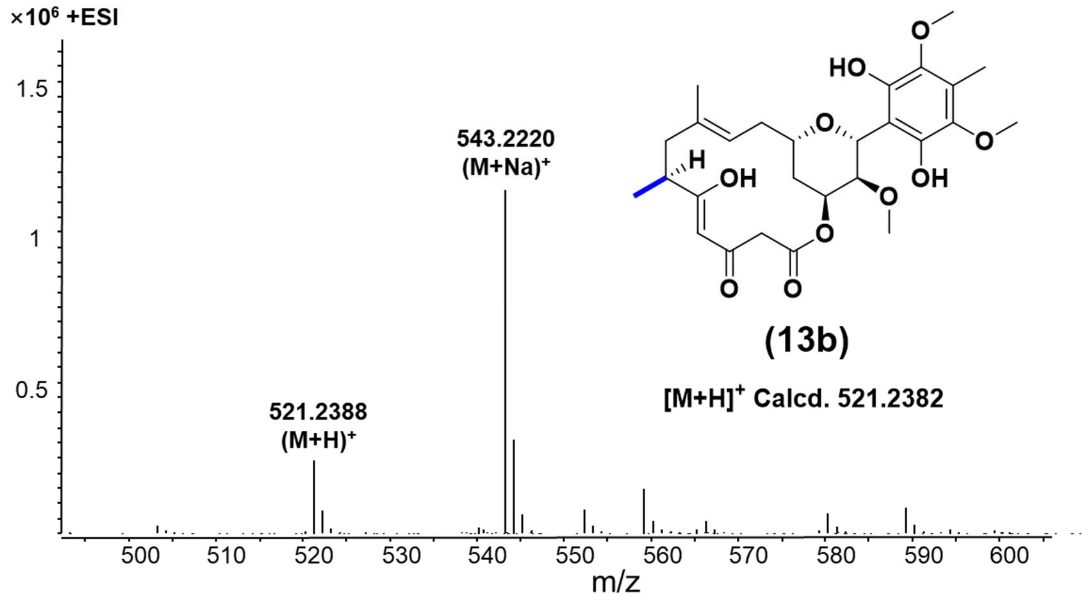

Supplementary Fig. 76. HR-ESI-MS spectrum of compound **13b**.

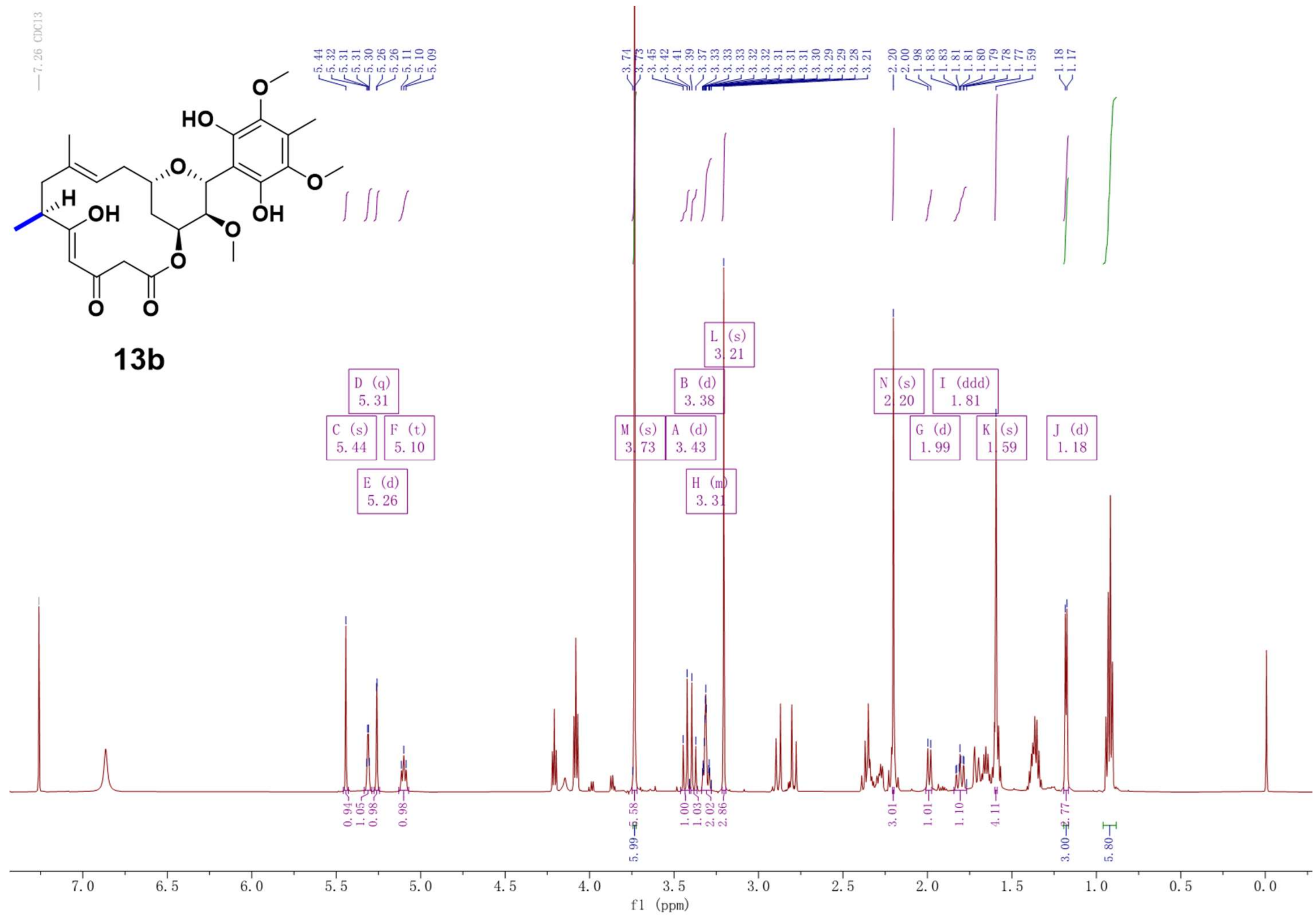

Supplementary Fig. 77. $^{1}H$ NMR spectrum (600 MHz, $CDCl_3$) of compound **13b**.

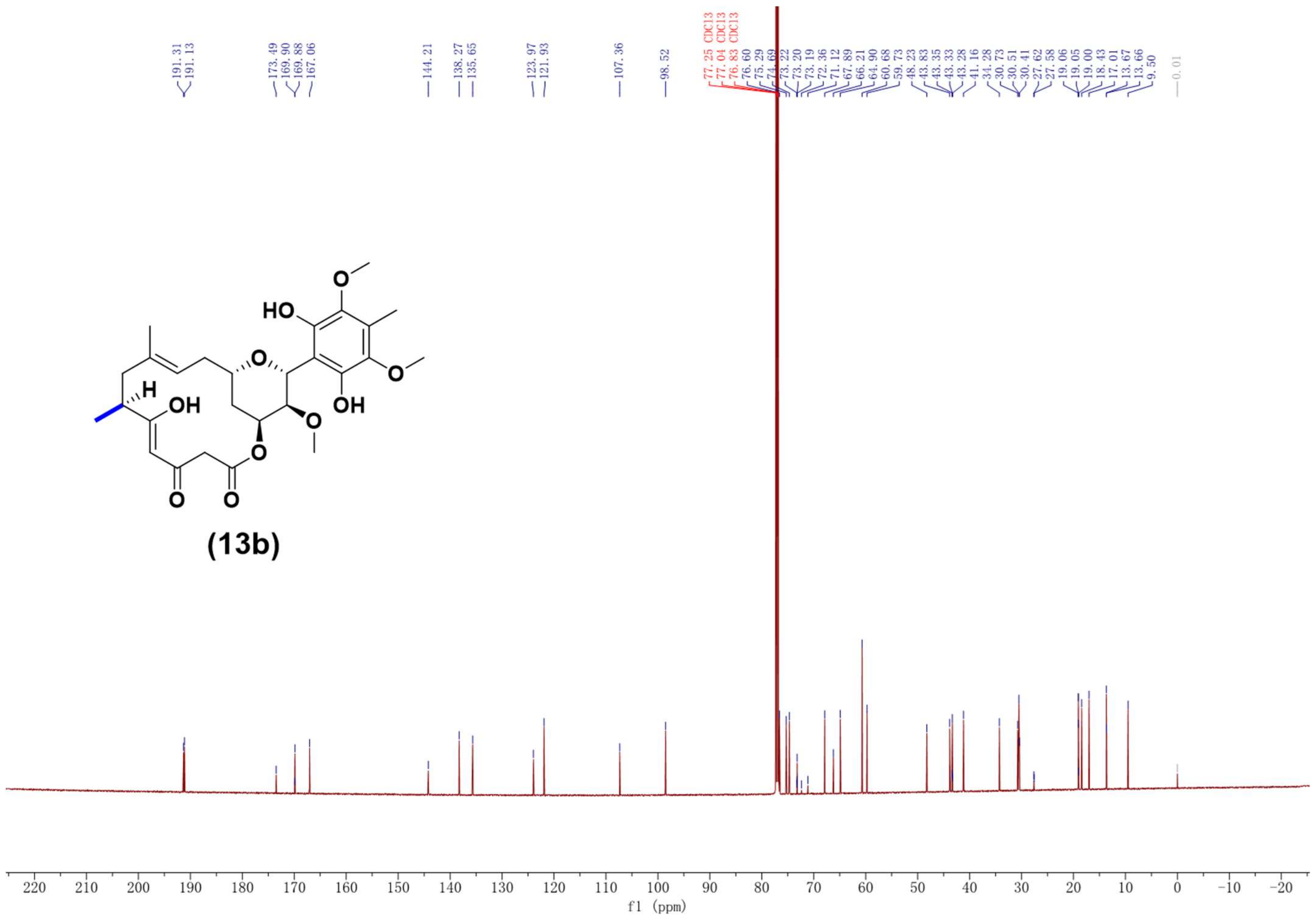

Supplementary Fig. 78. 13C NMR spectrum (150 MHz, CDCl3) of compound **13b**.

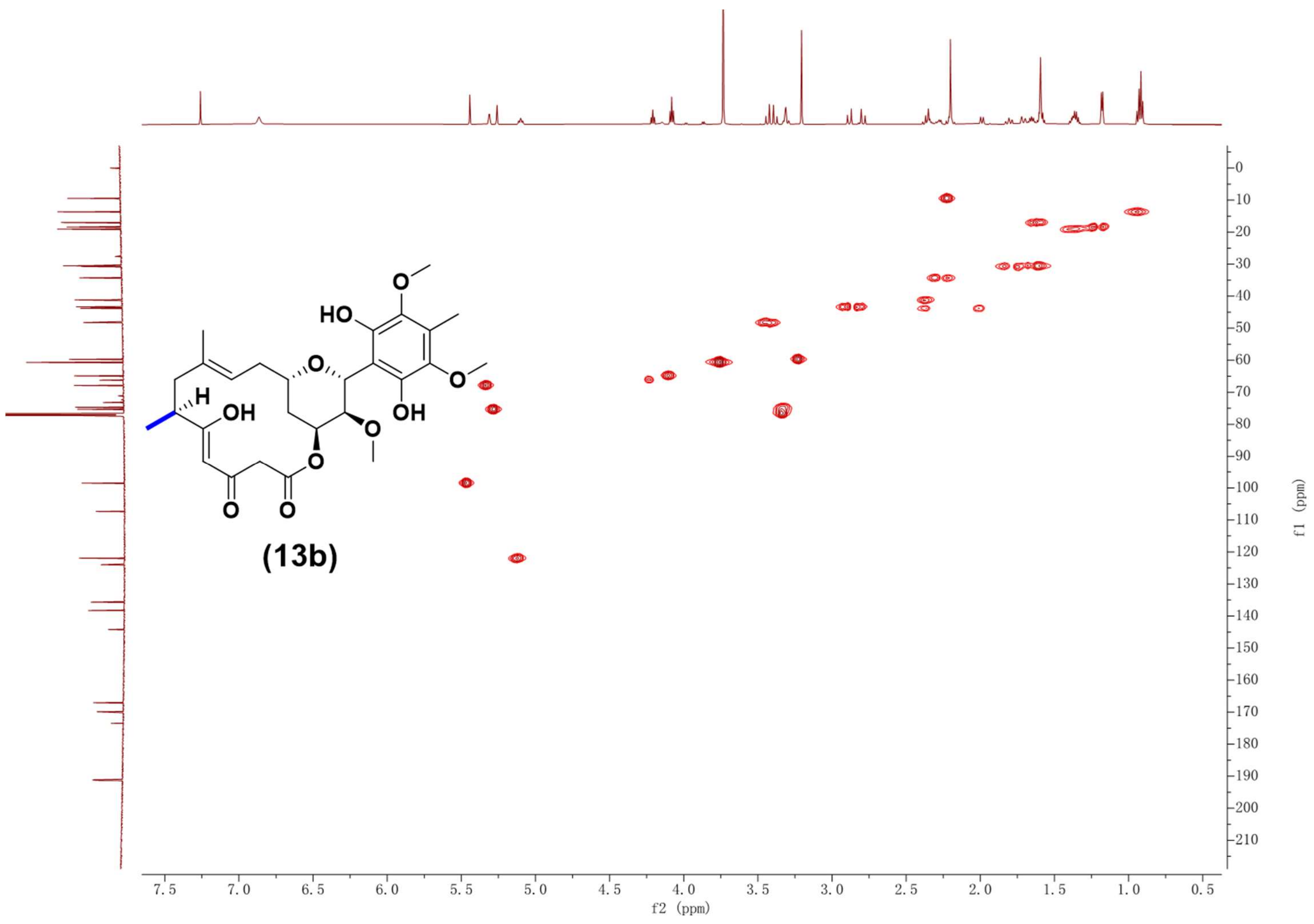

Supplementary Fig. 79. HSQC NMR spectrum (600 MHz, CDCl3) of compound **13b**.

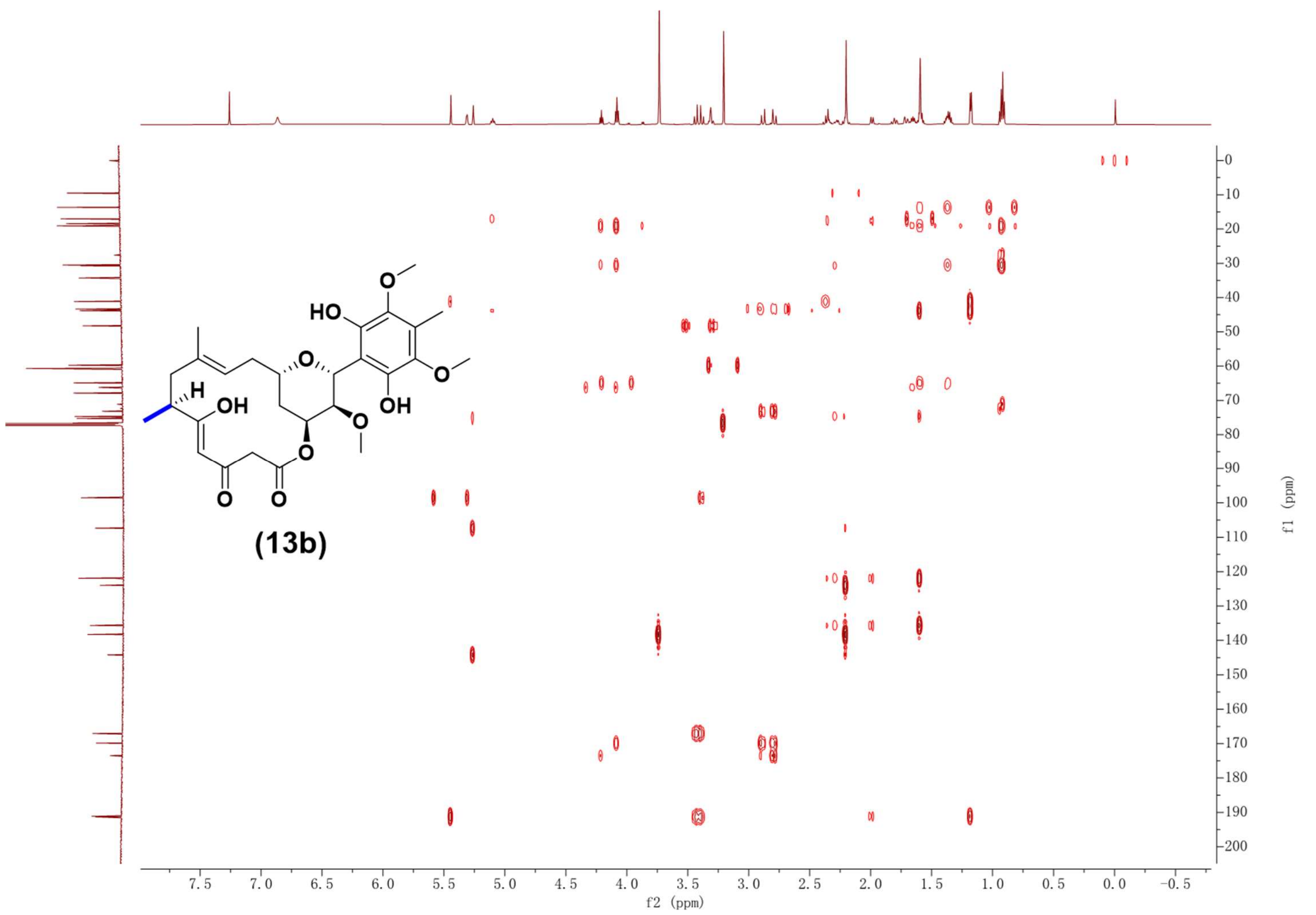

Supplementary Fig. 80. HMBC NMR spectrum (600 MHz, CDCl3) of compound **13b**.

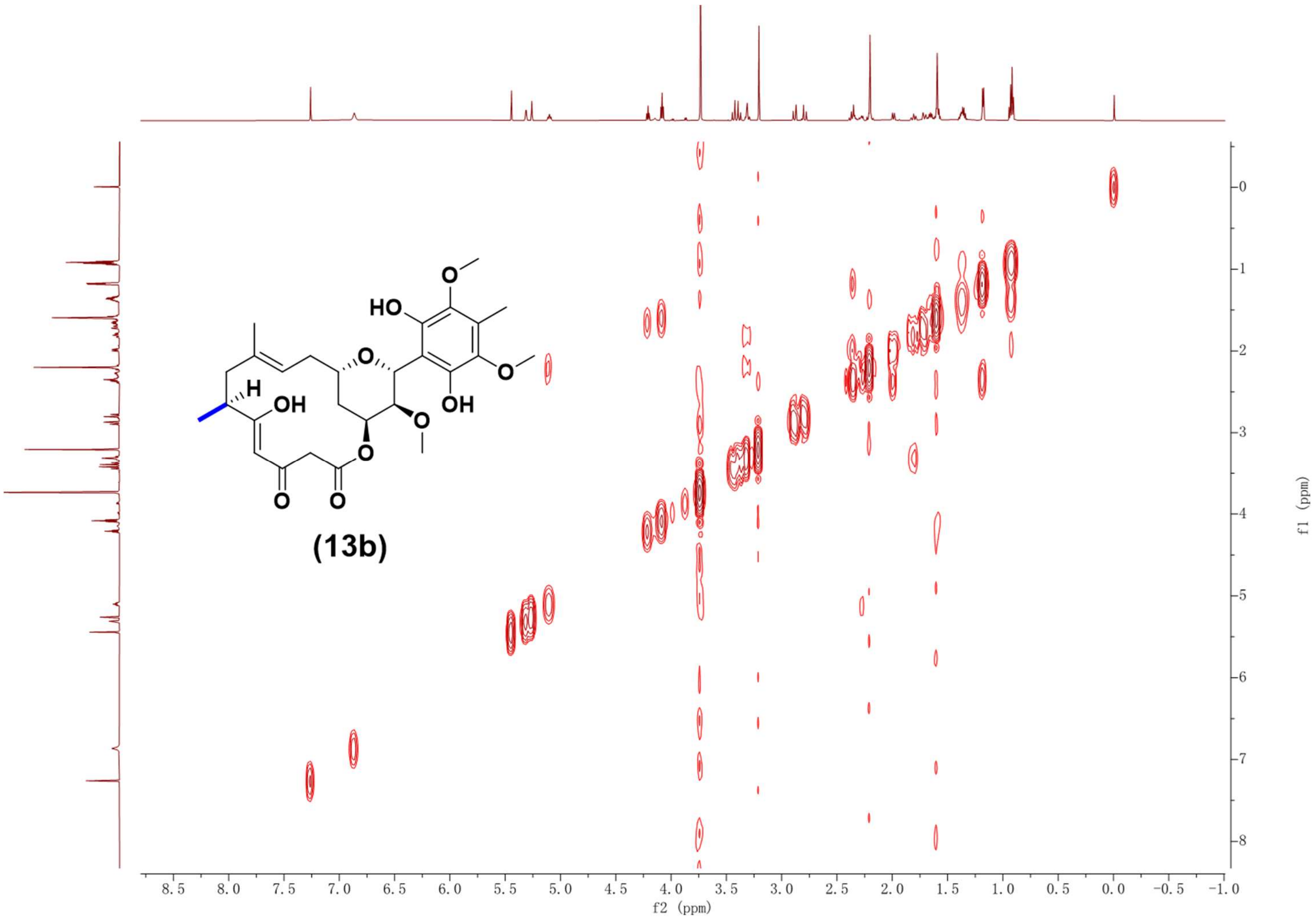

Supplementary Fig. 81. $^{1}H$-$^{1}H$ COSY NMR spectrum (600 MHz, $CDCl_3$) of compound **13b**.

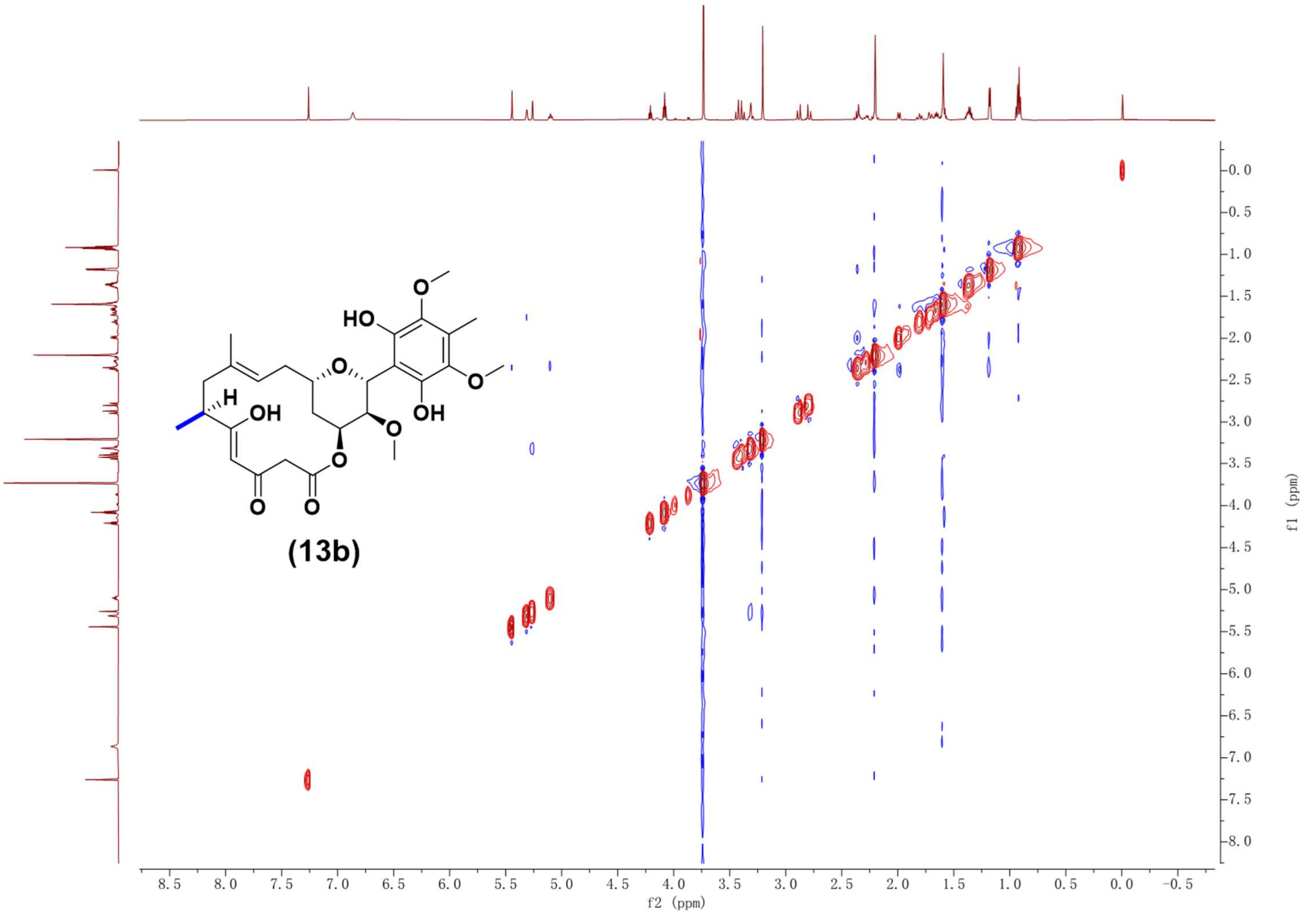

Supplementary Fig. 82. NOESY NMR spectrum (600 MHz, $CDCl_3$) of compound **13b**.

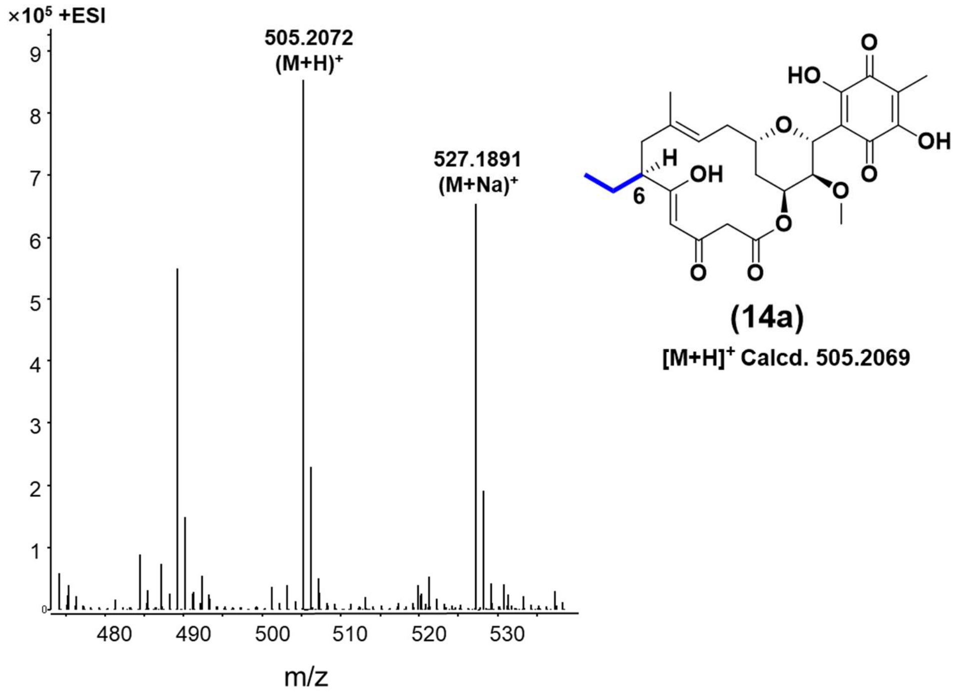

Supplementary Fig. 83. HR-ESI-MS spectrum of compound **14a**.

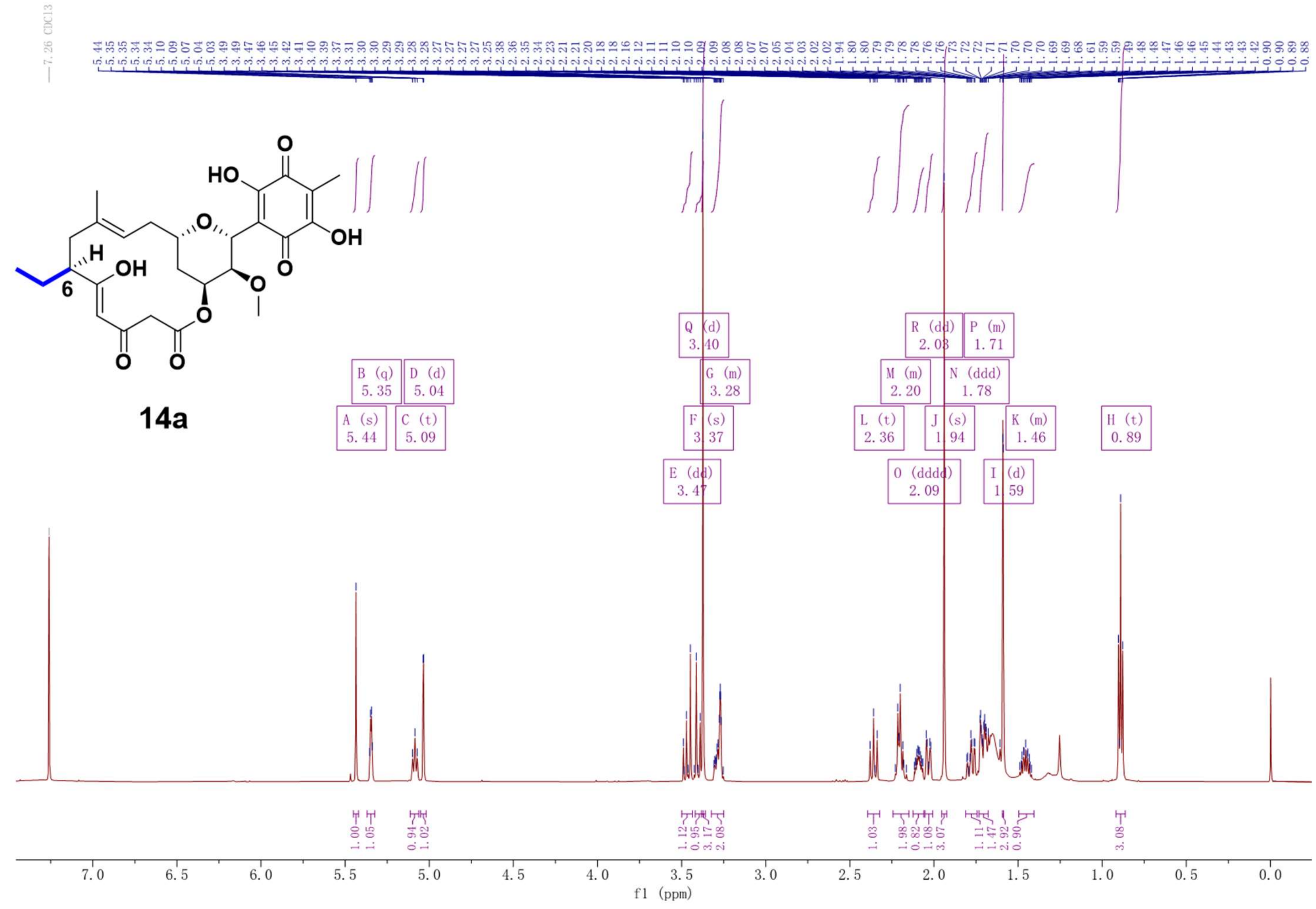

Supplementary Fig. 84. $^{1}H$ NMR spectrum (600 MHz, $CDCl_3$) of compound **14a**.

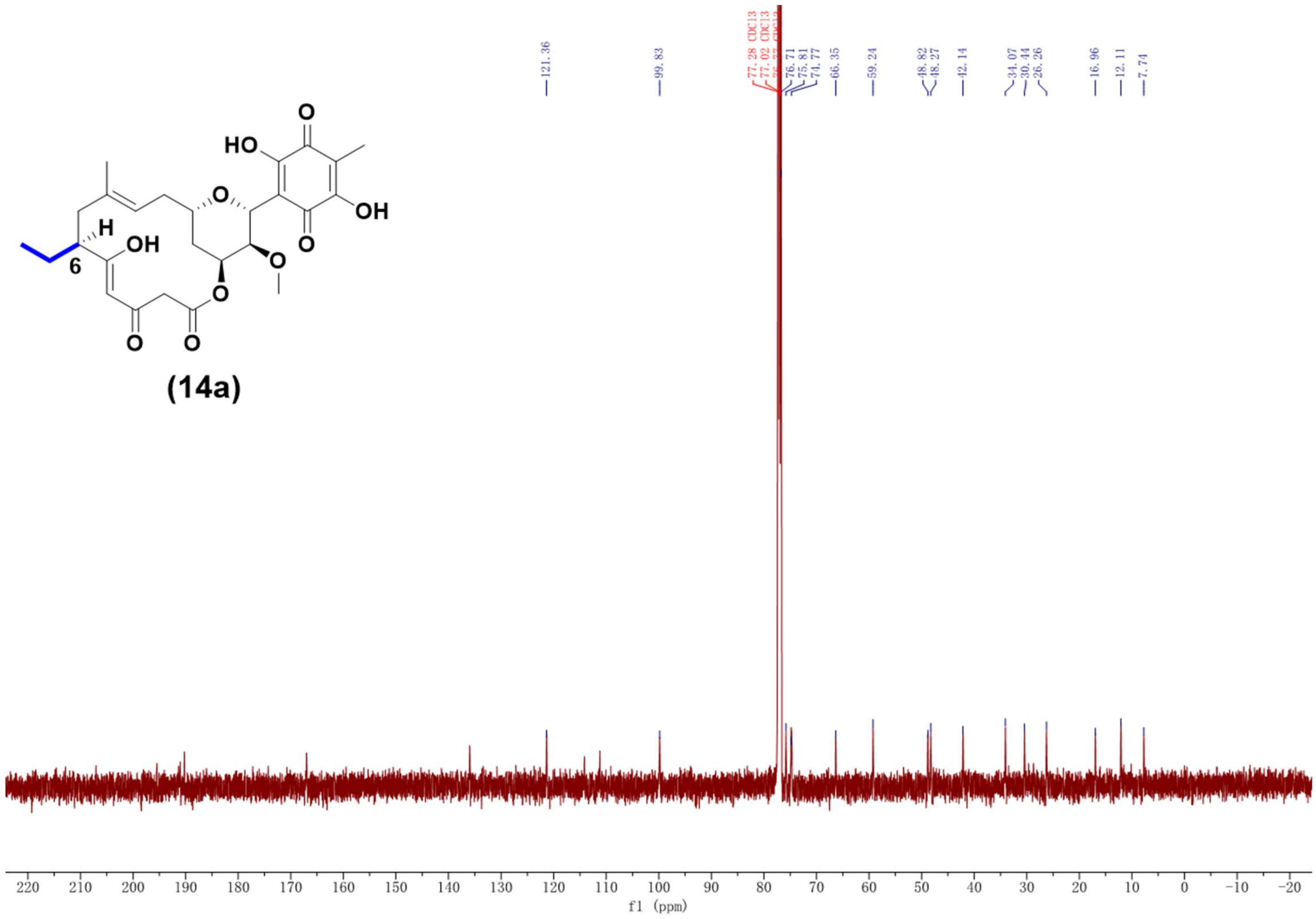

Supplementary Fig. 85. $^{13}$C NMR spectrum (125 MHz, $CDCl_3$) of compound **14a**.

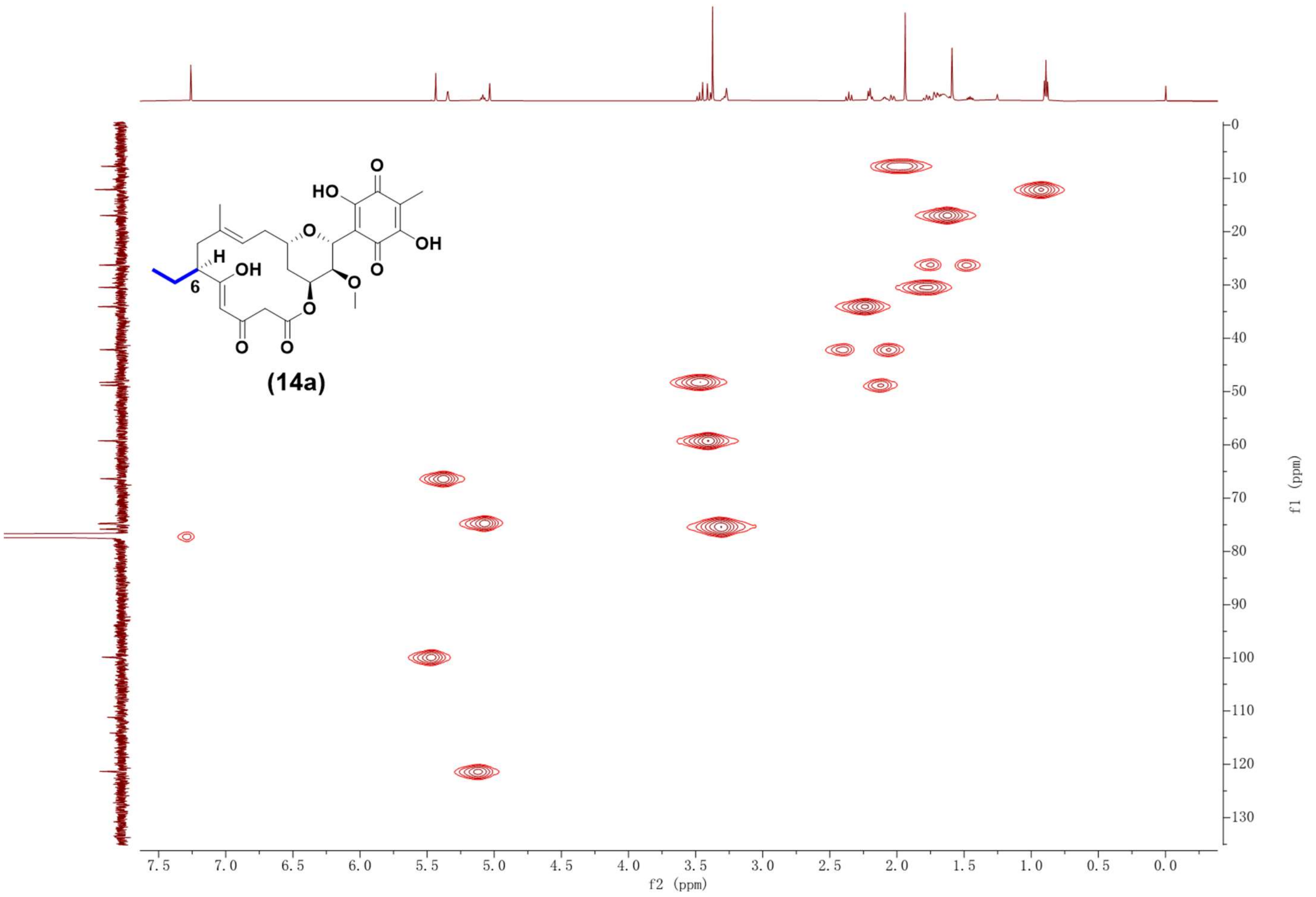

Supplementary Fig. 86. HSQC NMR spectrum (600 MHz, $CDCl_3$) of compound **14a**.

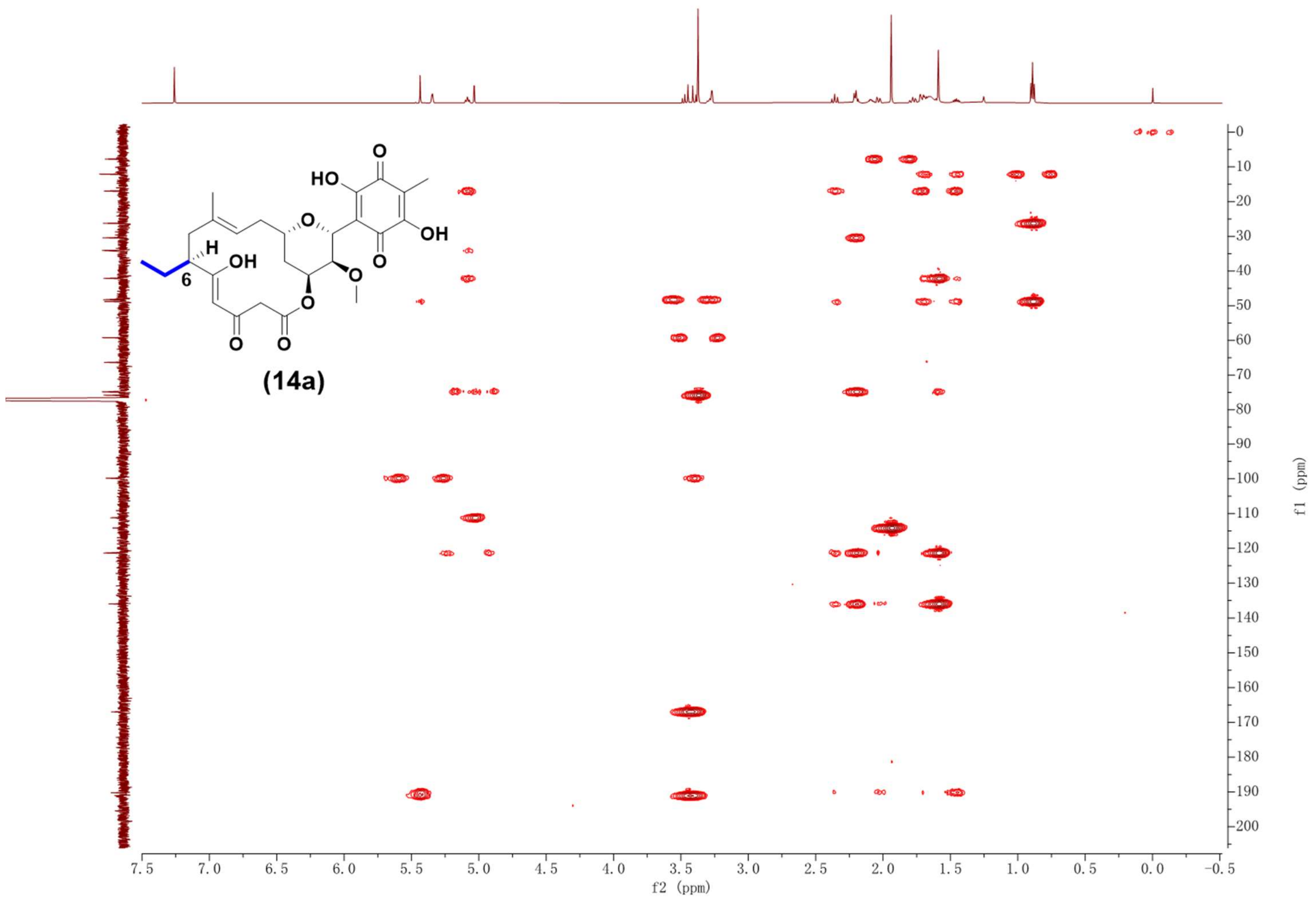

Supplementary Fig. 87. [1]HMBC NMR spectrum (600 MHz, $CDCl_3$) of compound **14a**.

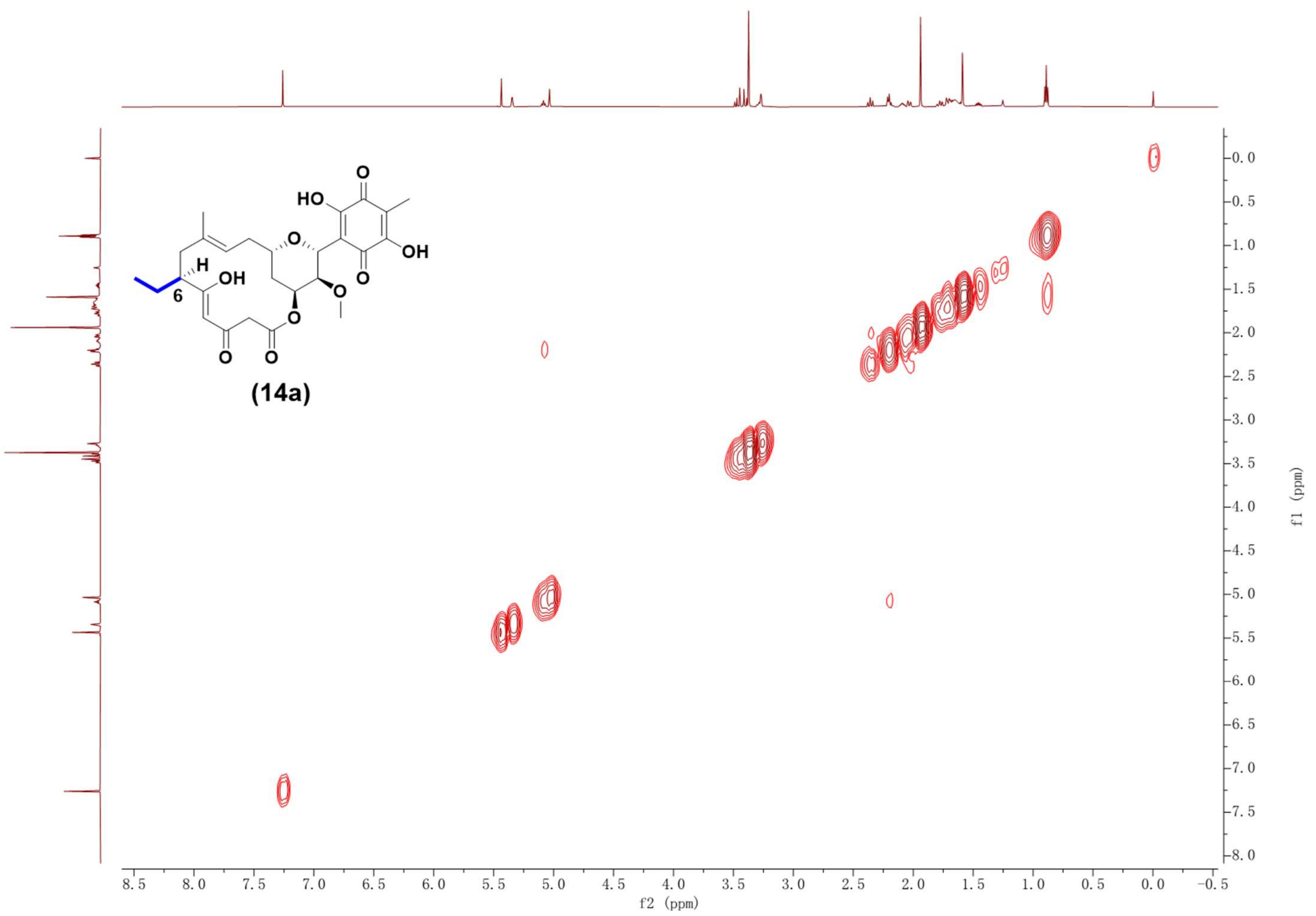

Supplementary Fig. 88. $^1H$-$^1H$ COSY NMR spectrum (600 MHz, $CDCl_3$) of compound **14a**.

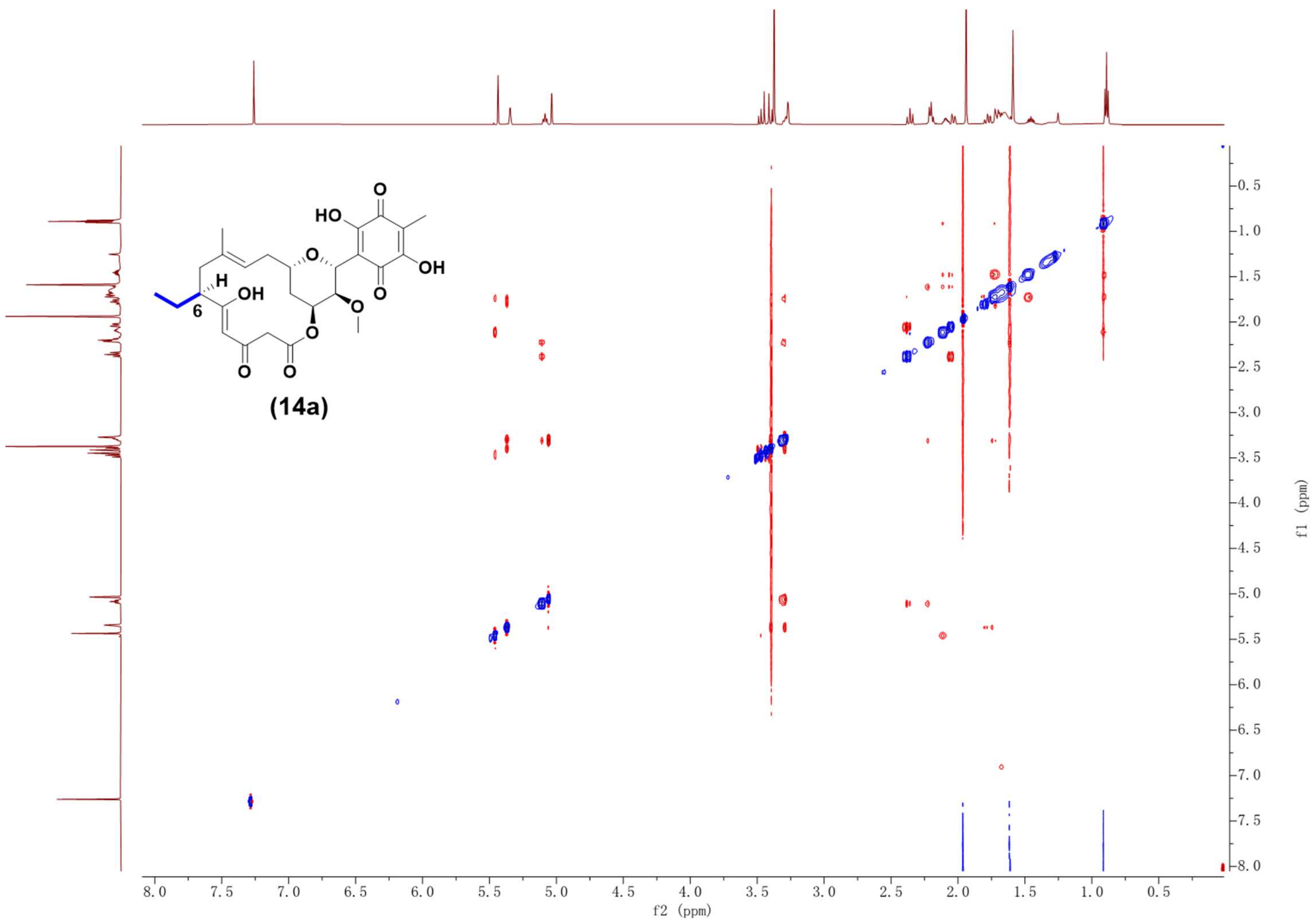

Supplementary Fig. 89. NOESY NMR spectrum (600 MHz, $CDCl_3$) of compound **14a**.

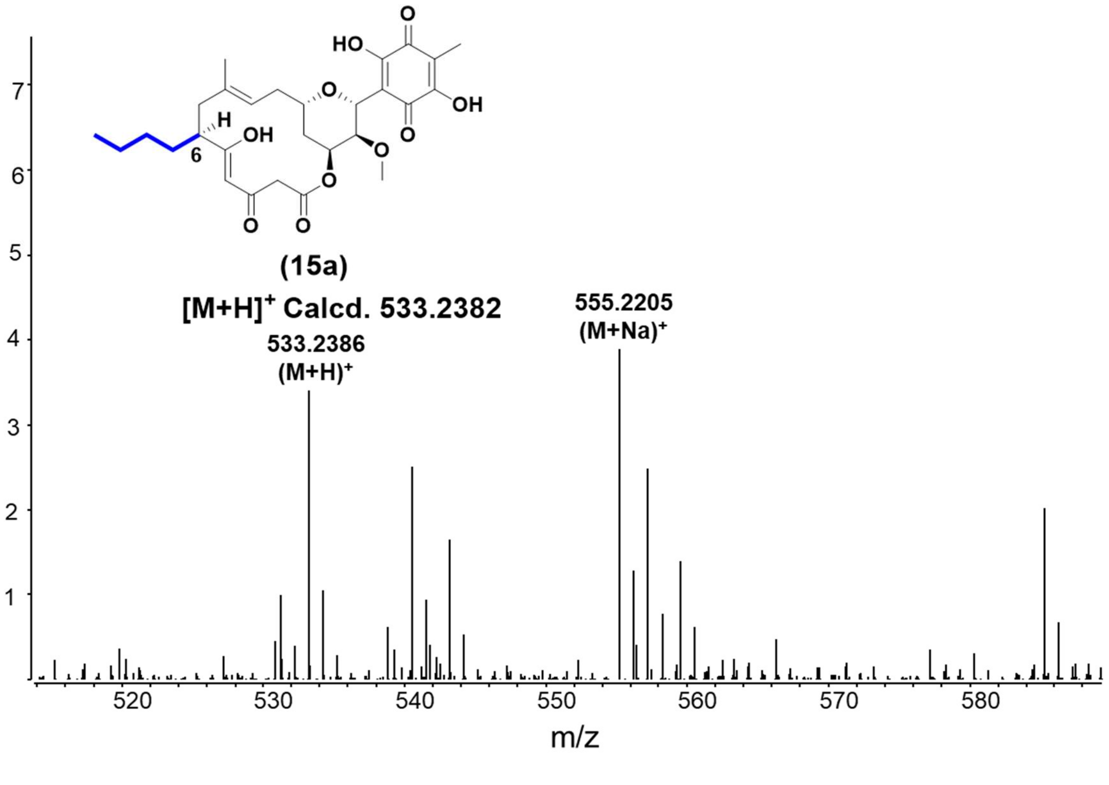

Supplementary Fig. 90. HR-ESI-MS spectrum of compound **15a**.

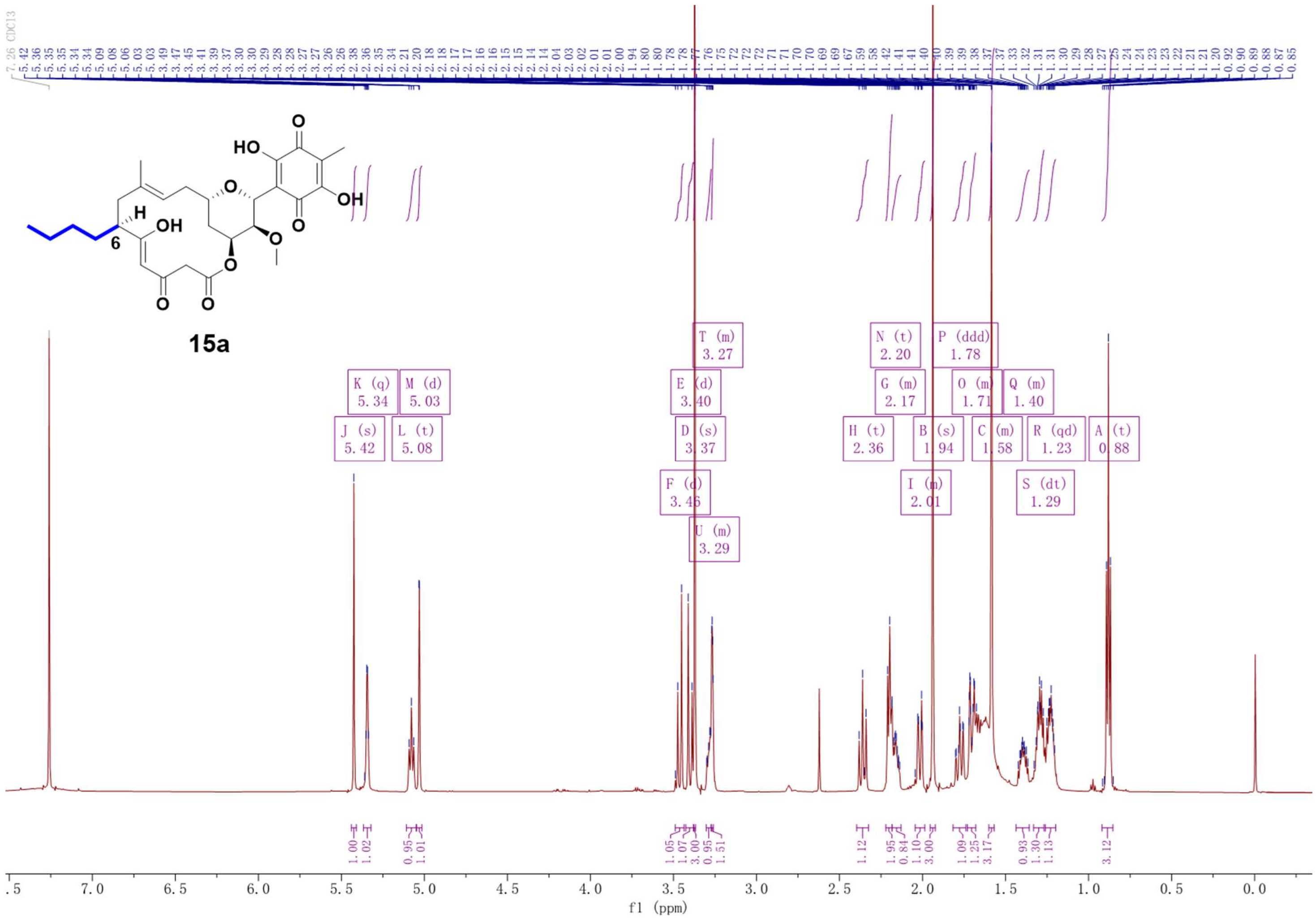

Supplementary Fig. 91. $^{1}$H NMR spectrum (600 MHz, $CDCl_3$) of compound **15a.**

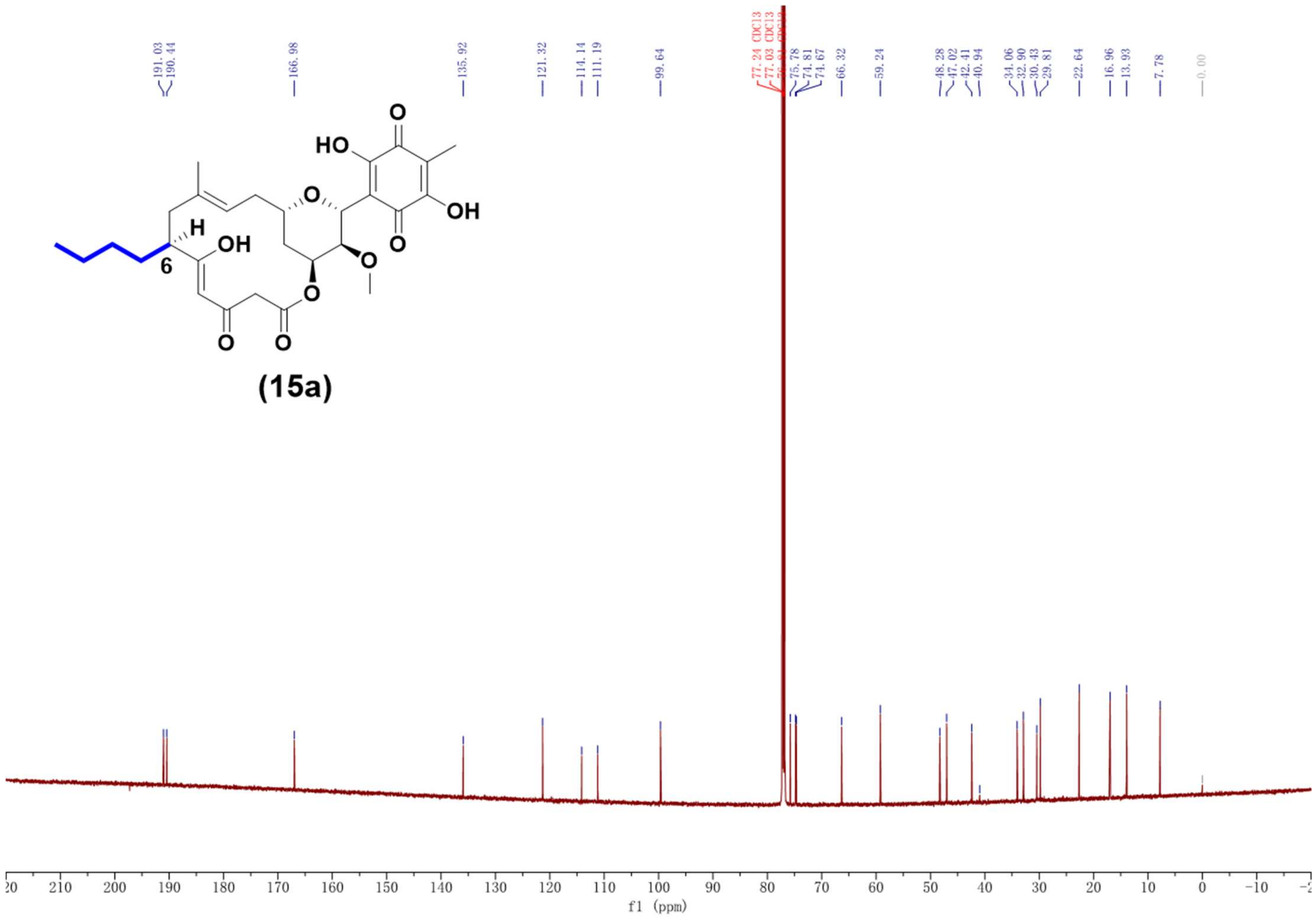

Supplementary Fig. 92. 13C NMR spectrum (150 MHz, CDCl3) of compound **15a**.

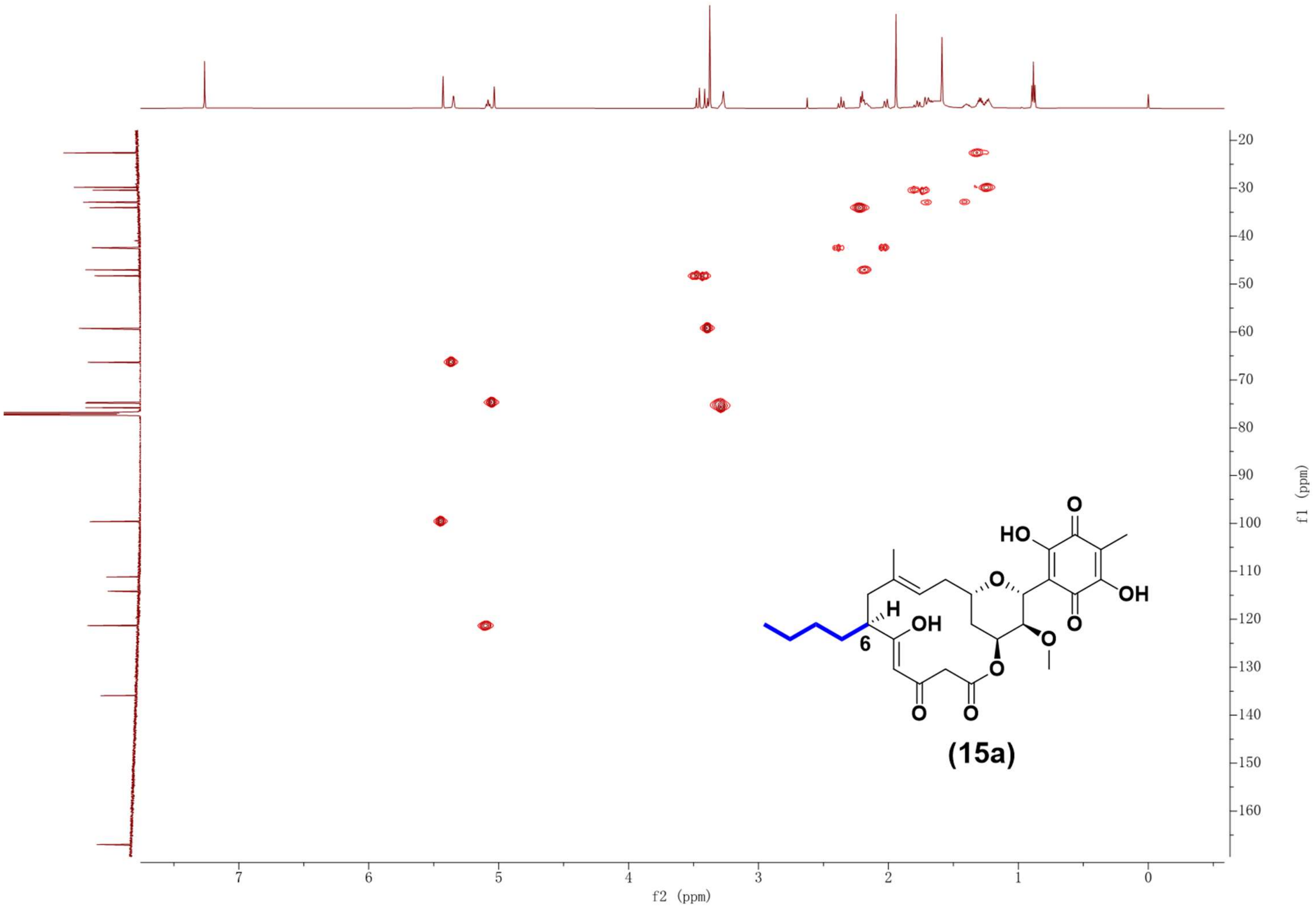

Supplementary Fig. 93. HSQC NMR spectrum (600 MHz, $CDCl_3$) of compound **15a**.

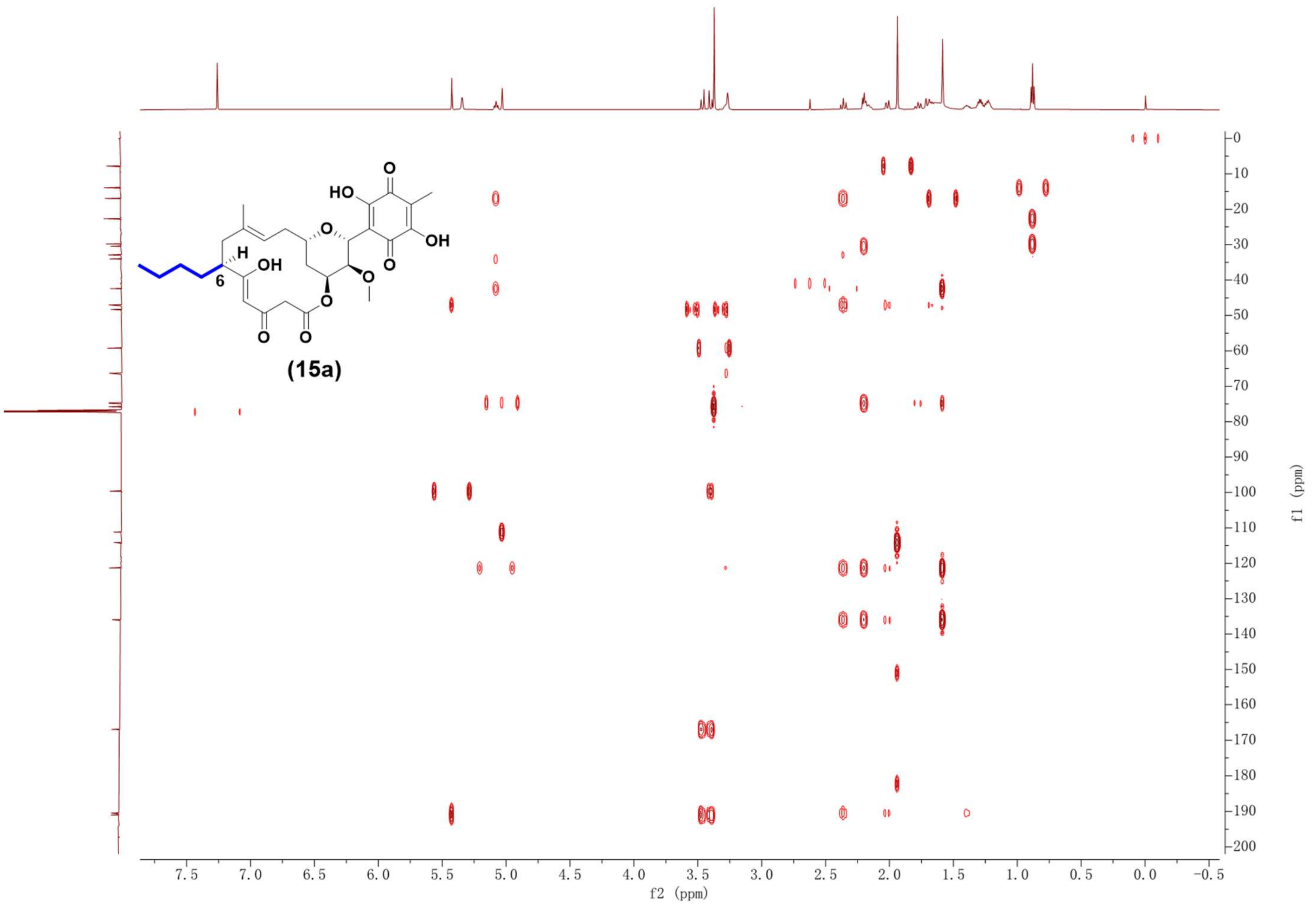

Supplementary Fig. 94. HMBC NMR spectrum (600 MHz, $CDCl_3$) of compound **15a**.

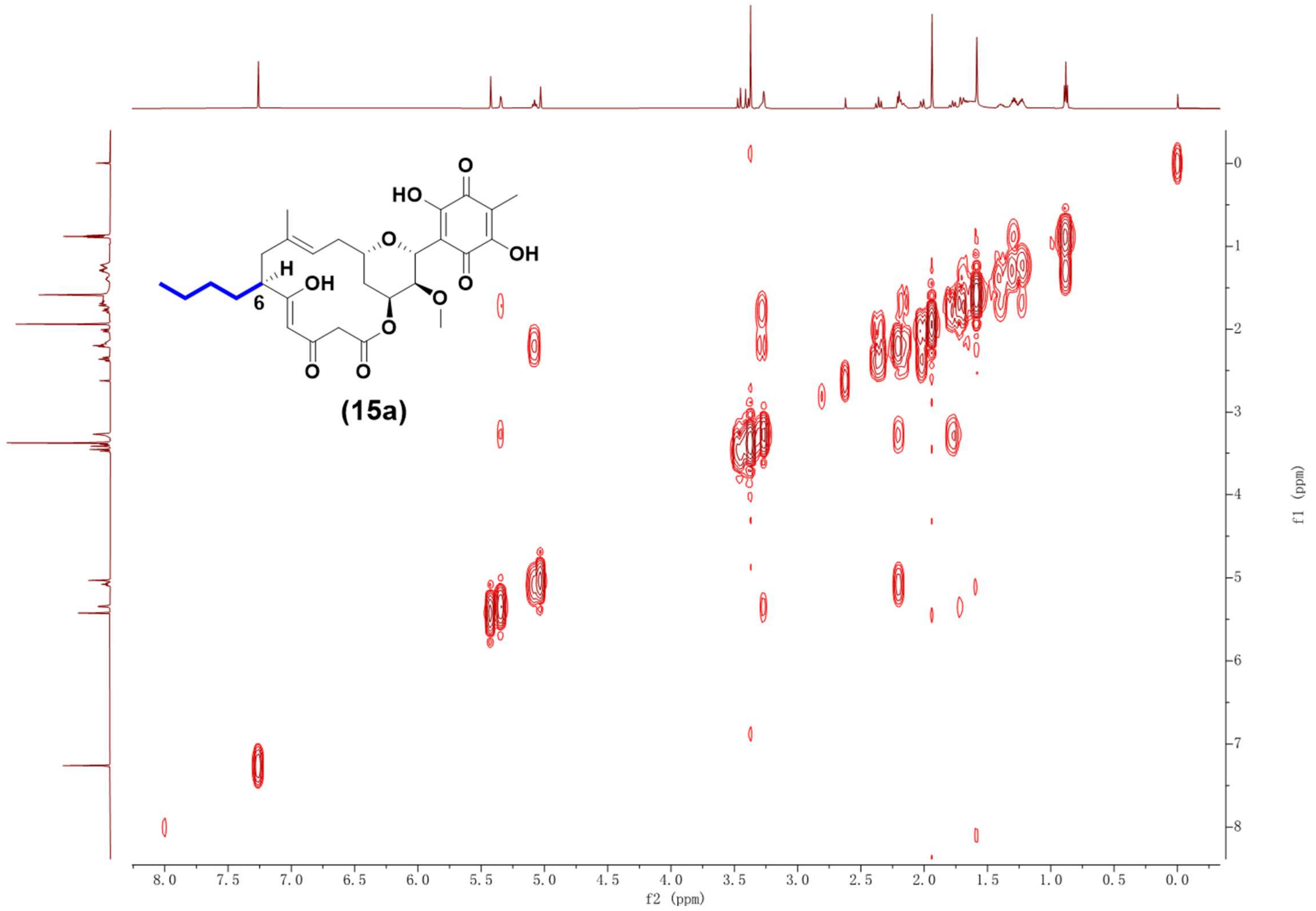

Supplementary Fig. 95. $^1H$-$^1H$ COSY NMR spectrum (600 MHz, $CDCl_3$) of compound **15a**.

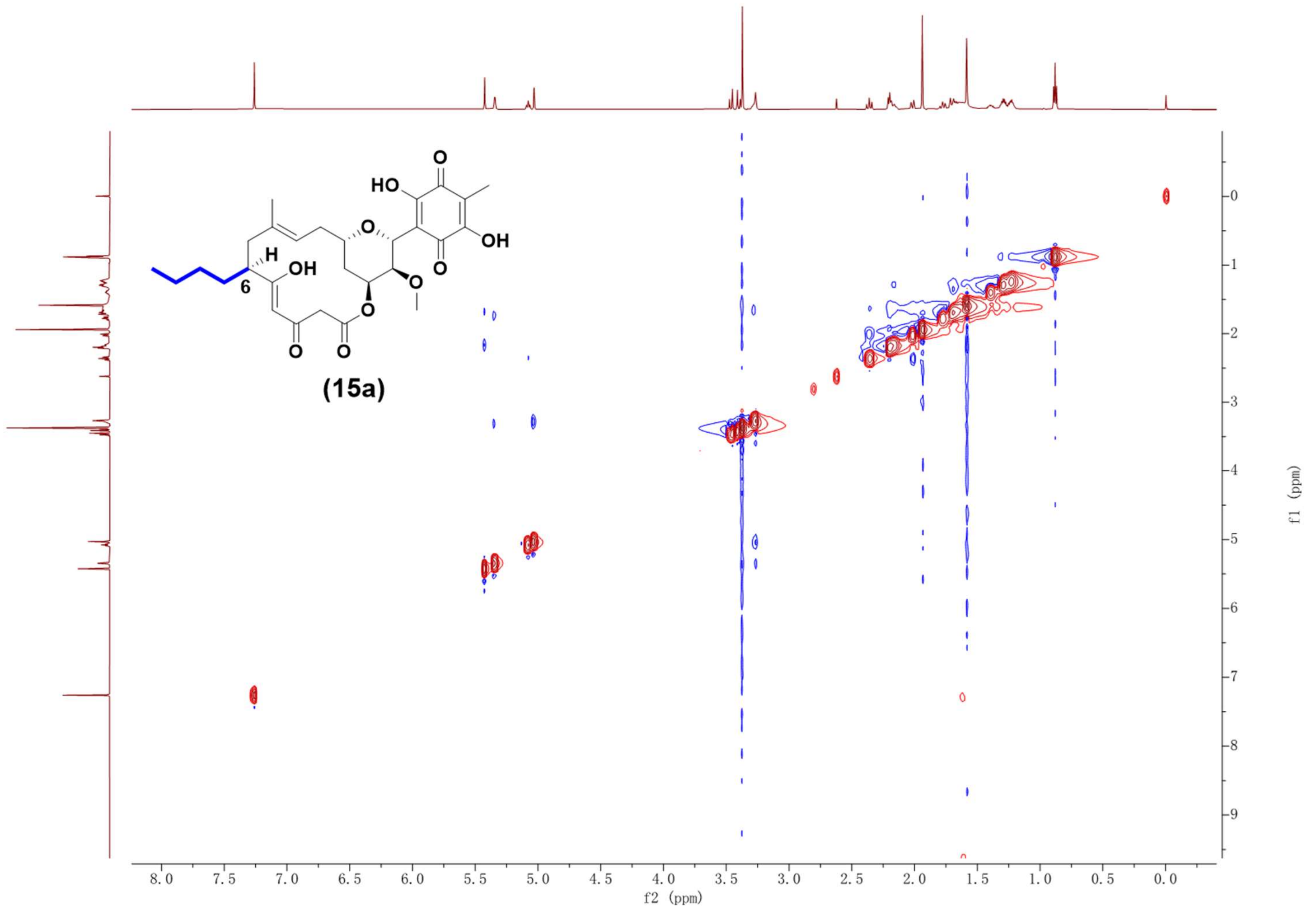

Supplementary Fig. 96. NOESY NMR spectrum (600 MHz, $CDCl_3$) of compound **15a**.

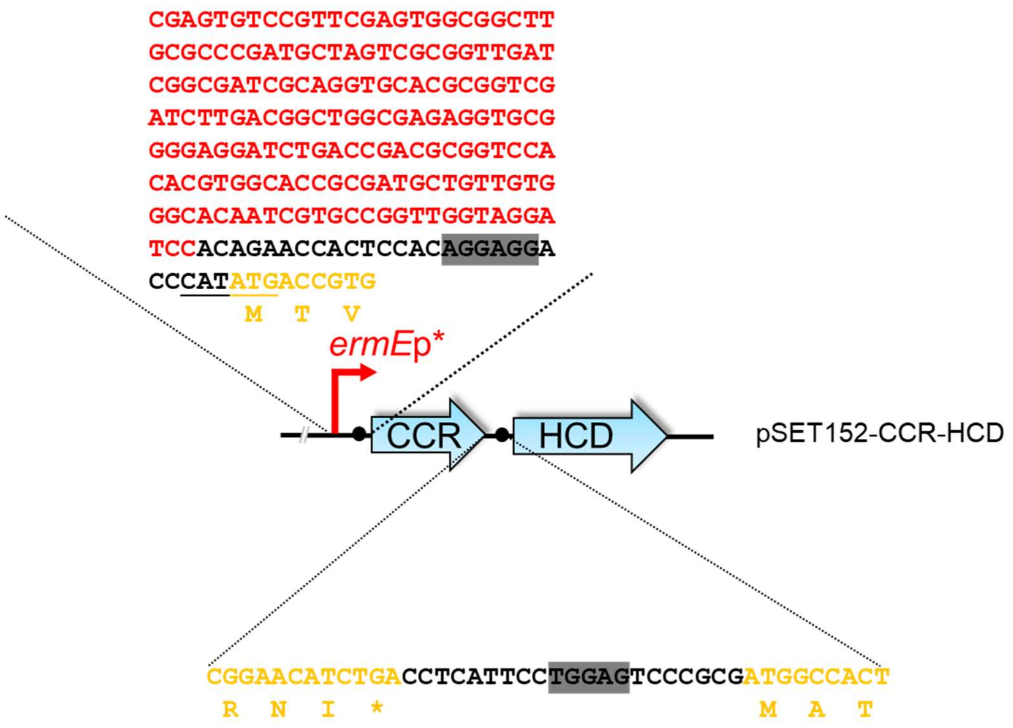

Supplementary Fig. 97. Nucleotide sequence of plasmid pSET1252-CCR-HCD**.**

The "*ermE**p" nucleotide sequence is displayed in red bold font. Black dots represent ribosome- binding sites. The nucleotide sequences of CCR (WP_019328626.1) and HCD (WP_011030948.1) are displayed in yellow bold font.

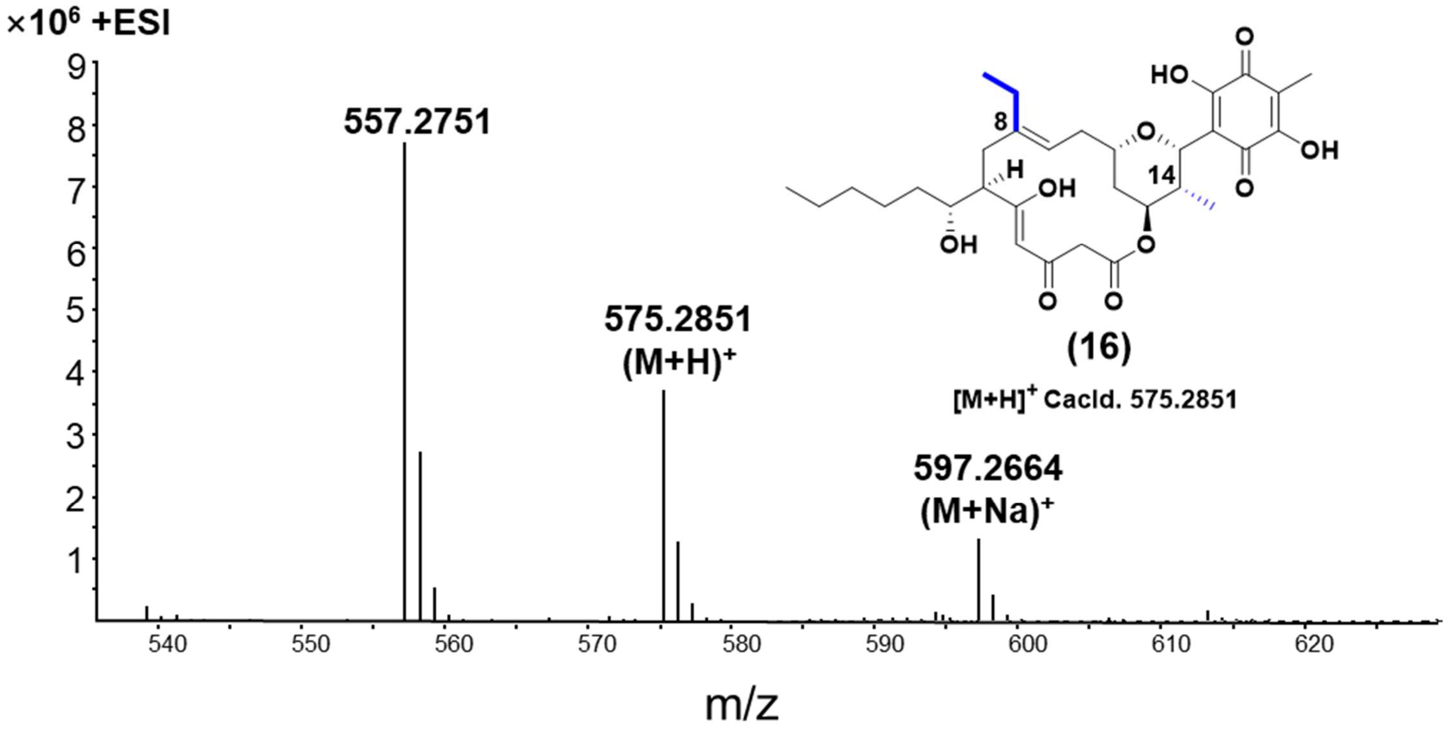

Supplementary Fig. 98. HR-ESI-MS spectrum of compound **16**.

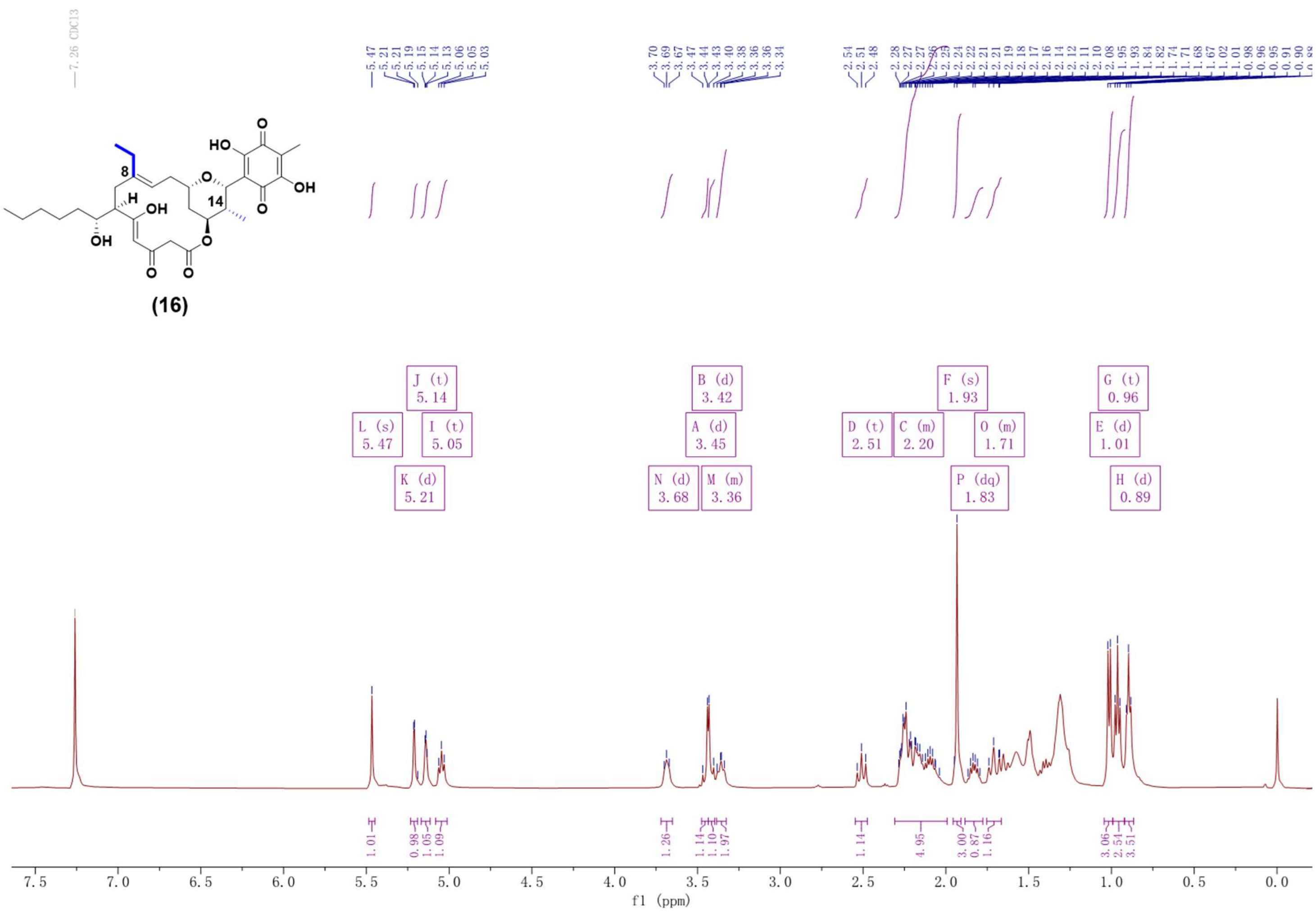

Supplementary Fig. 99. $^{1}$H NMR spectrum (500 MHz, $CDCl_3$) of compound **16**.

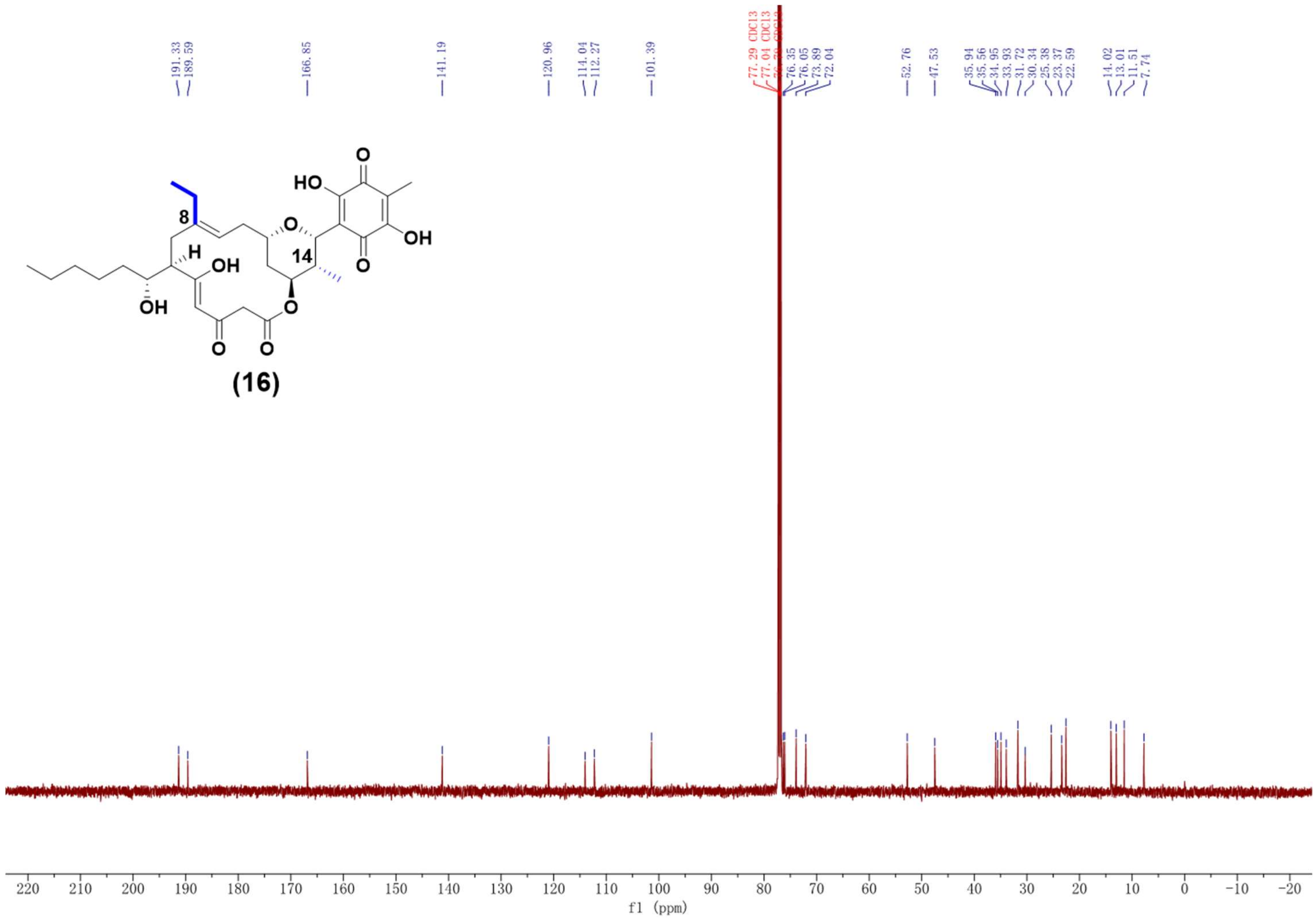

Supplementary Fig. 100. $^{13}C$ NMR spectrum (125 MHz, $CDCl_3$) of compound **16**.

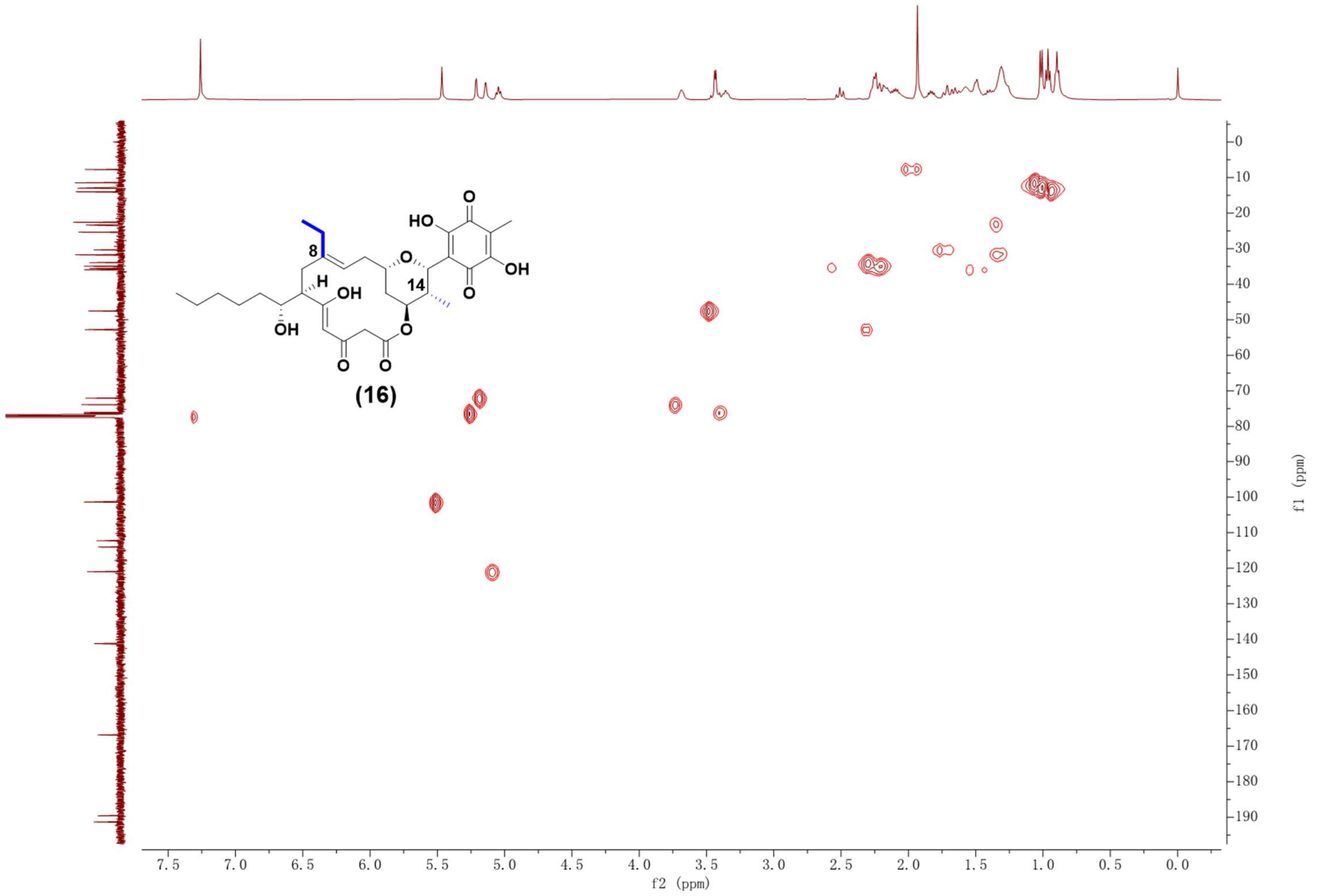

Supplementary Fig. 101. HSQC NMR spectrum (500 MHz, $CDCl_3$) of compound **16**.

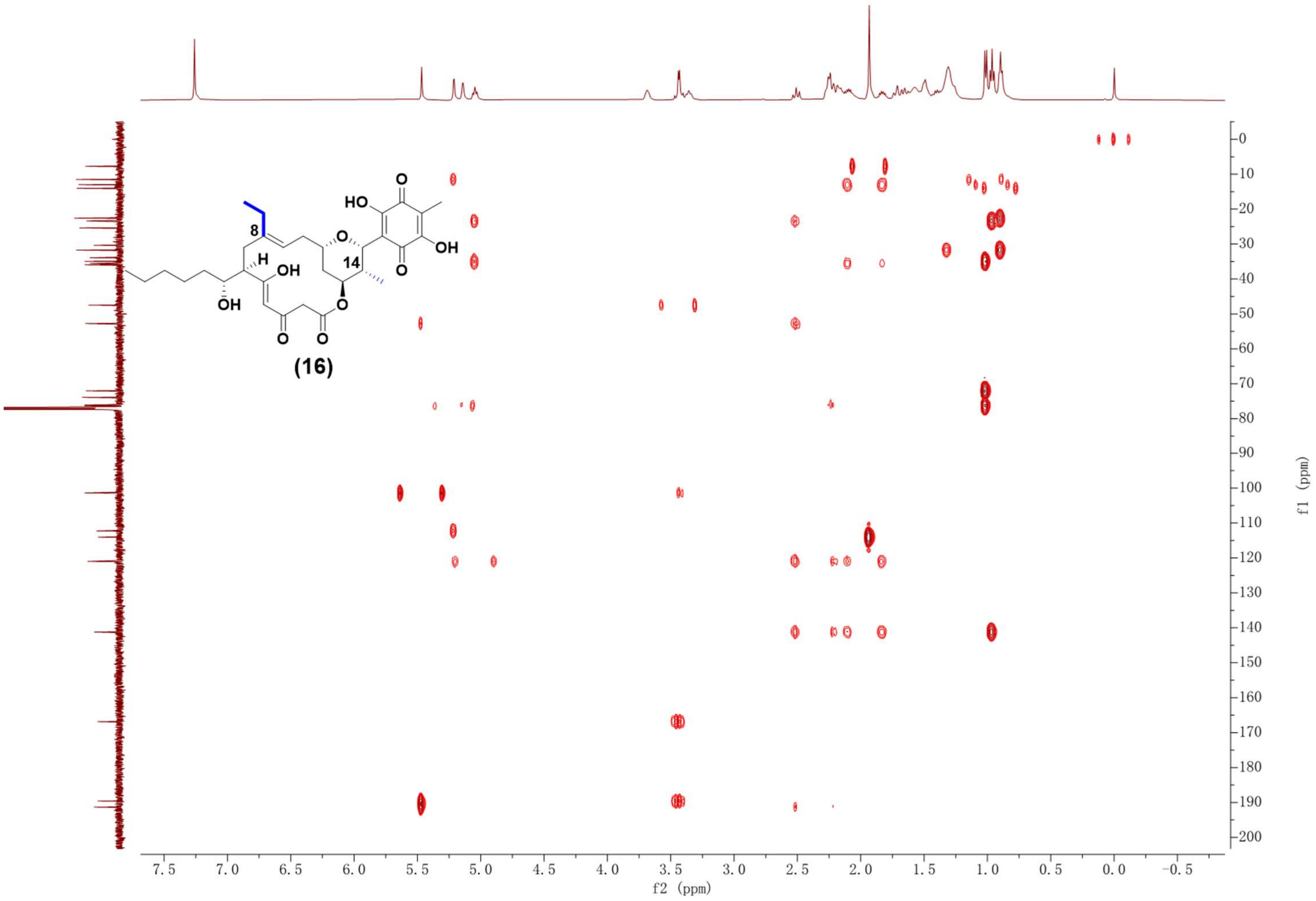

Supplementary Fig. 102. HMBC NMR spectrum (500 MHz, $CDCl_3$) of compound **16**.

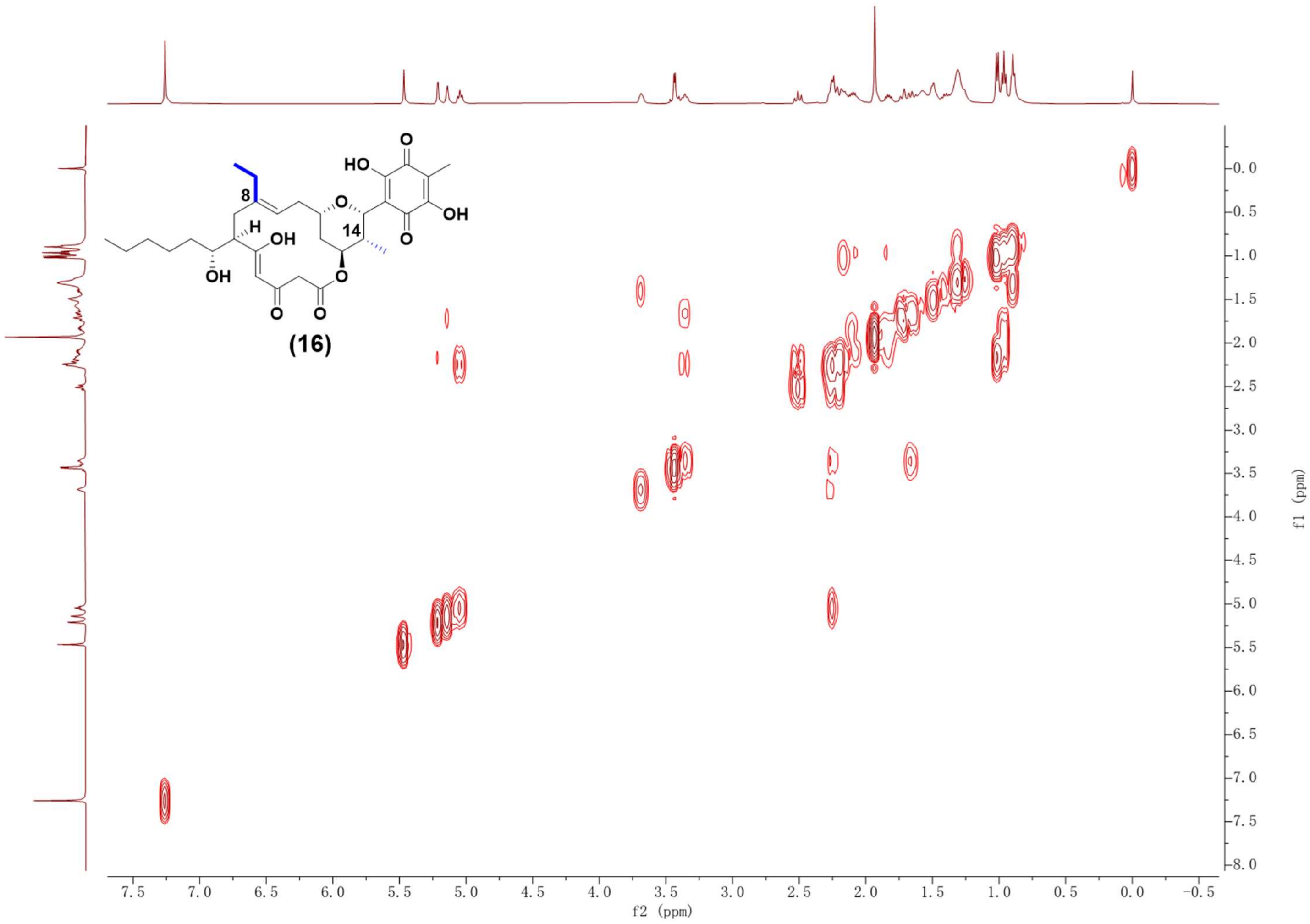

Supplementary Fig. 103. $^1H$-$^1H$ COSY NMR spectrum (500 MHz, $CDCl_3$) of compound **16**.

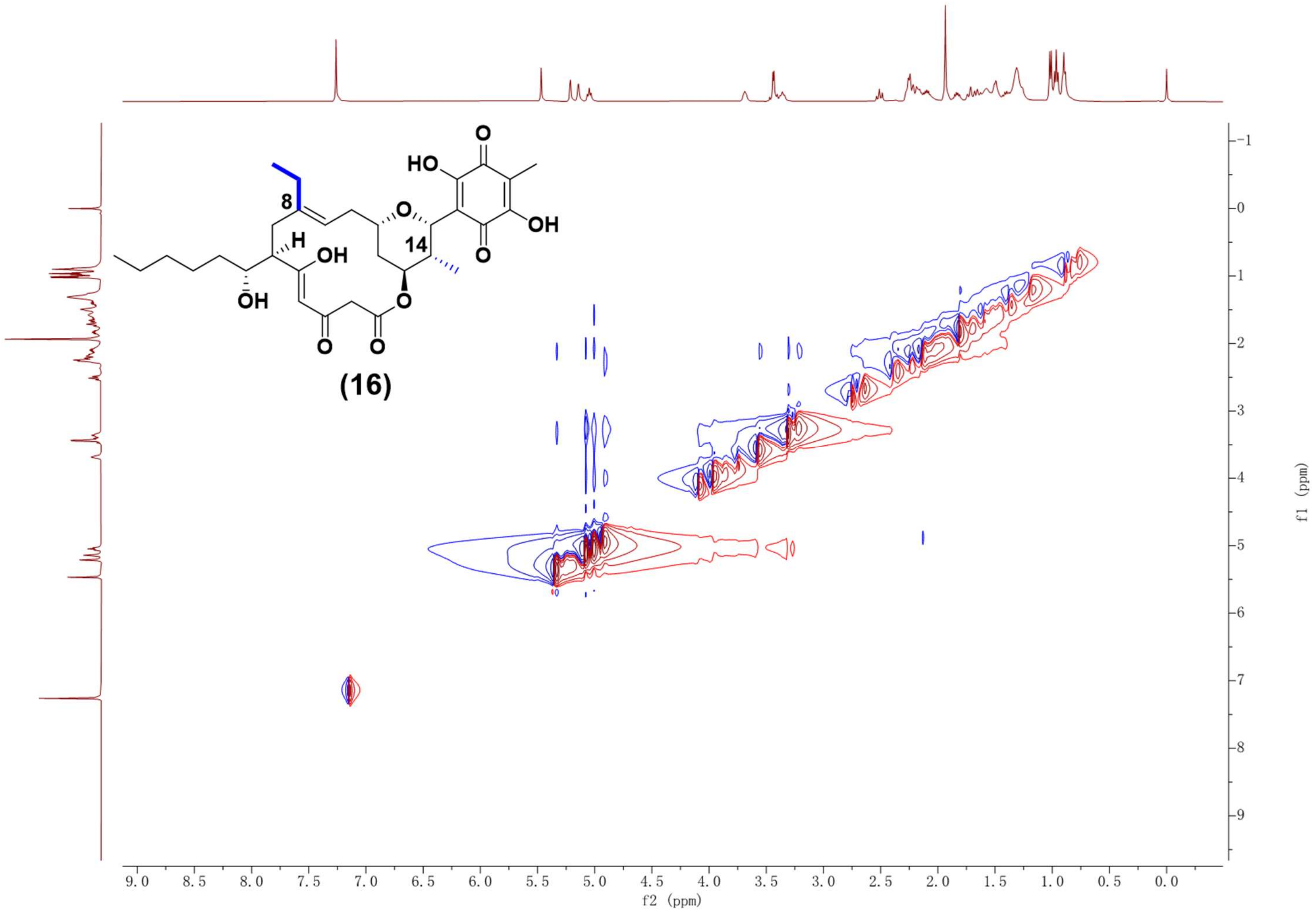

Supplementary Fig. 104. NOESY NMR spectrum (500 MHz, CDCl₃) of compound **16**.

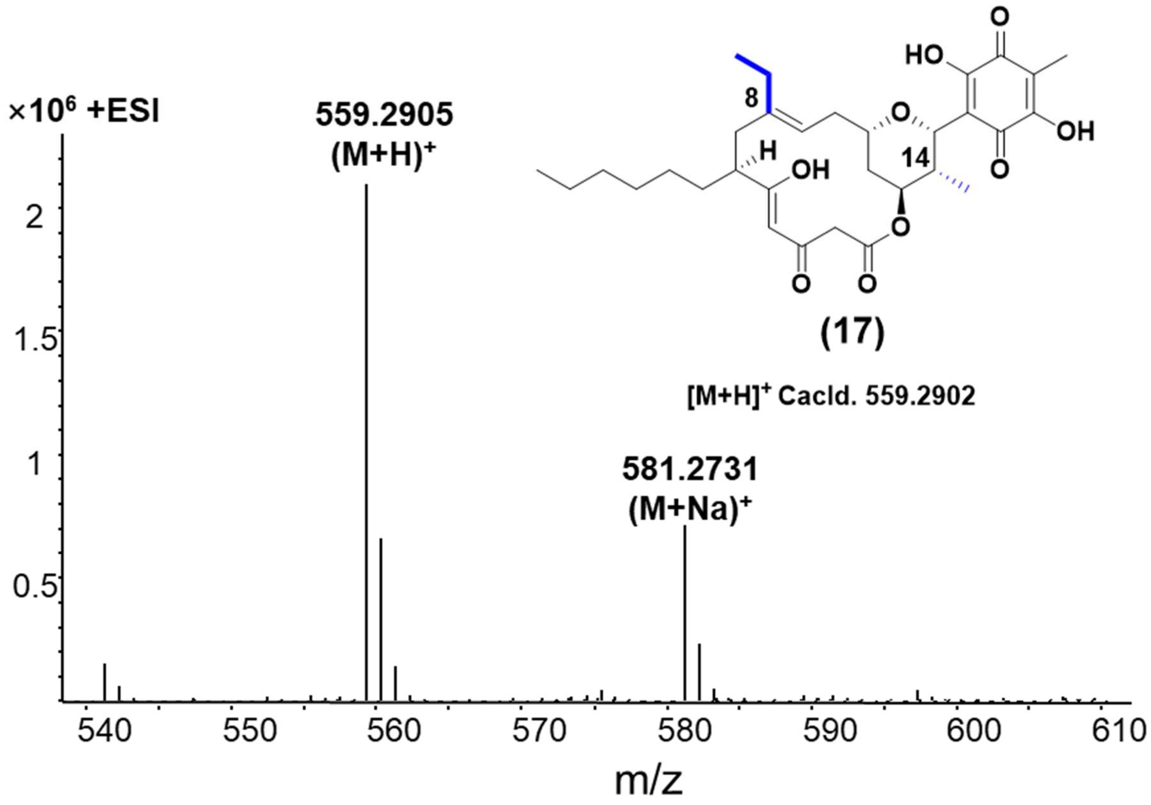

Supplementary Fig. 105. HR-ESI-MS spectrum of compound **17**.

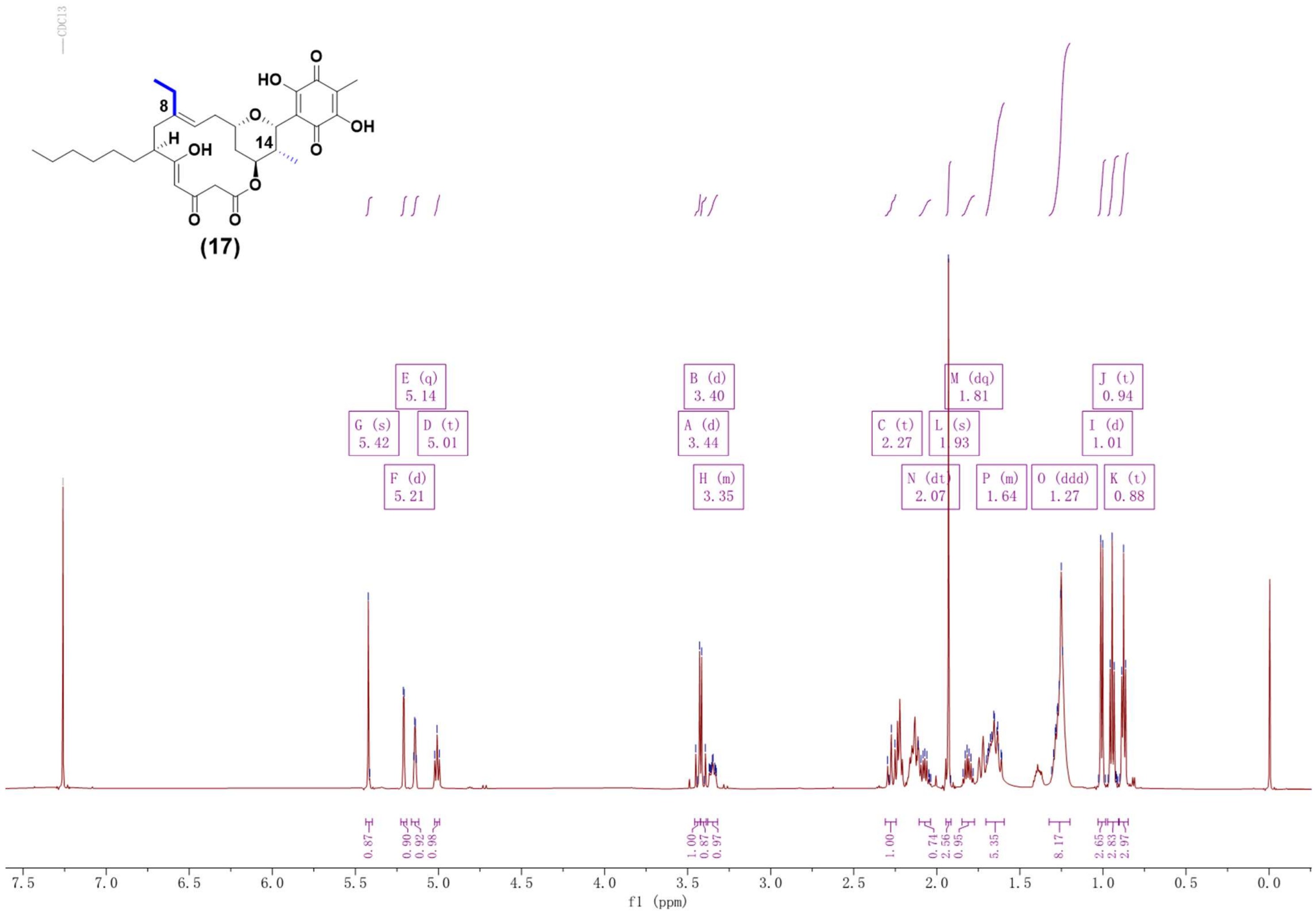

Supplementary Fig. 106. $^{1}H$ NMR spectrum (600 MHz, $CDCl_3$) of compound **17**.

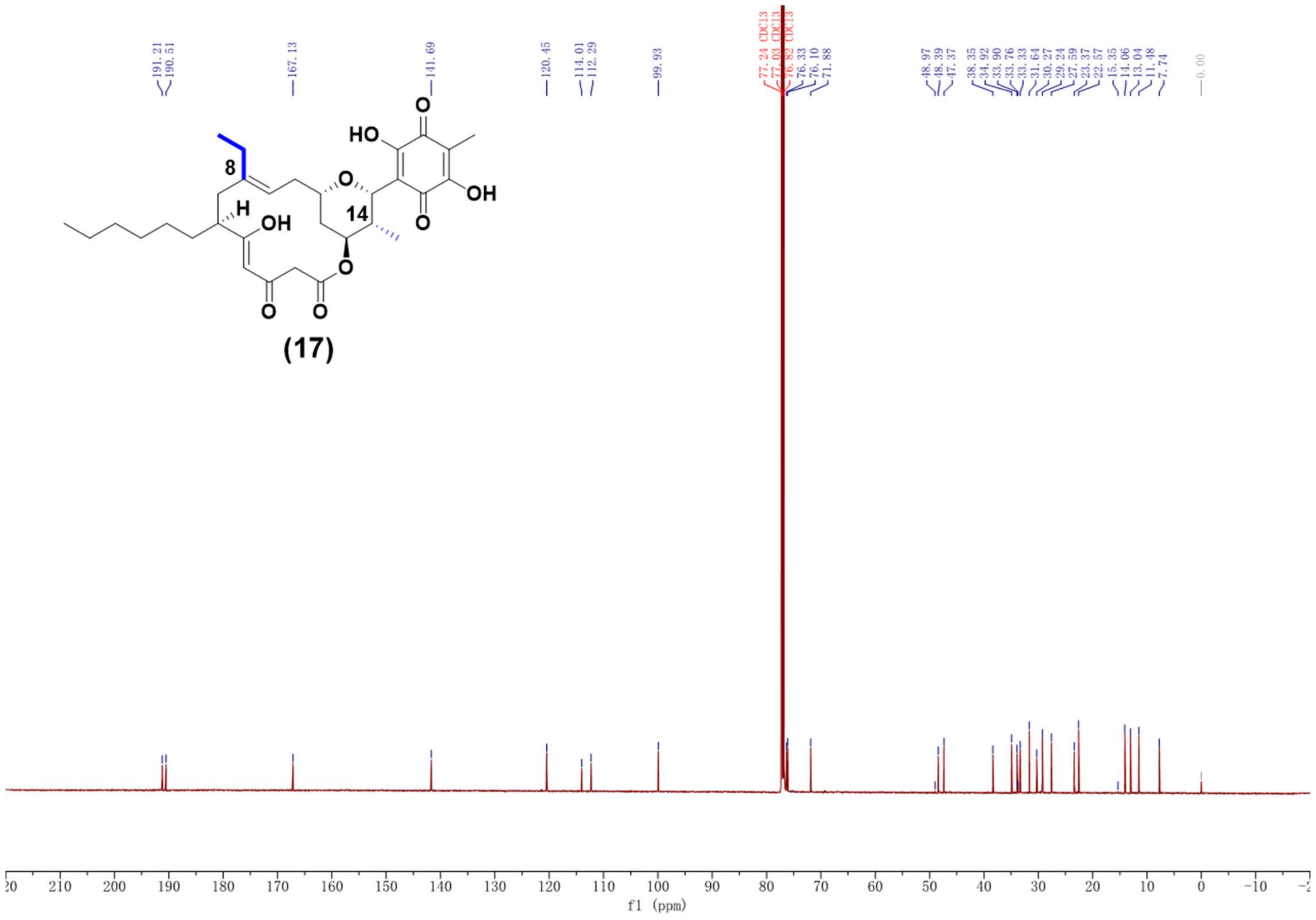

Supplementary Fig. 107. $^{13}C$ NMR spectrum (600 MHz, $CDCl_3$) of compound **17**.

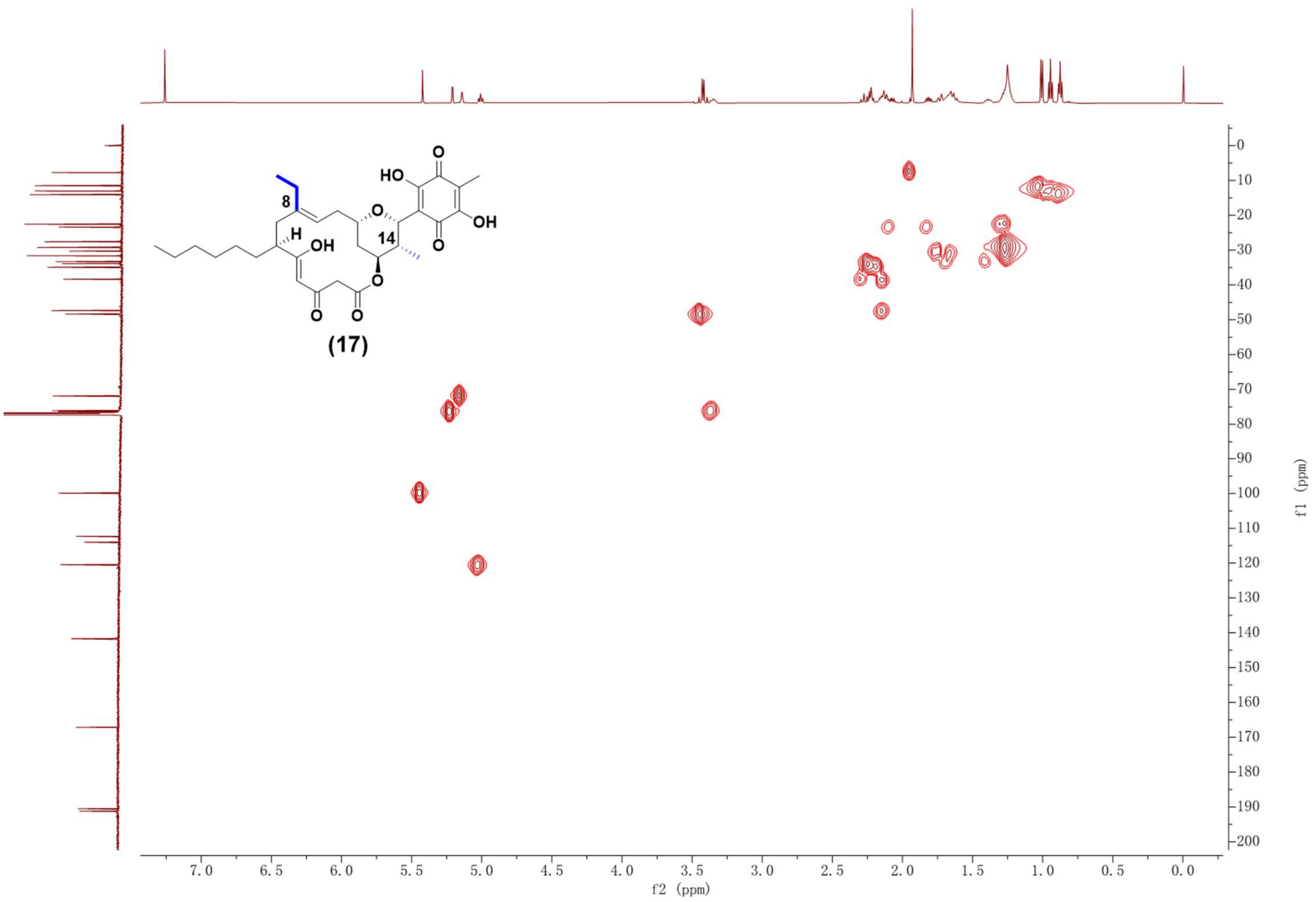

Supplementary Fig. 108. HSQC NMR spectrum (600 MHz, $CDCl_3$) of compound **17**.

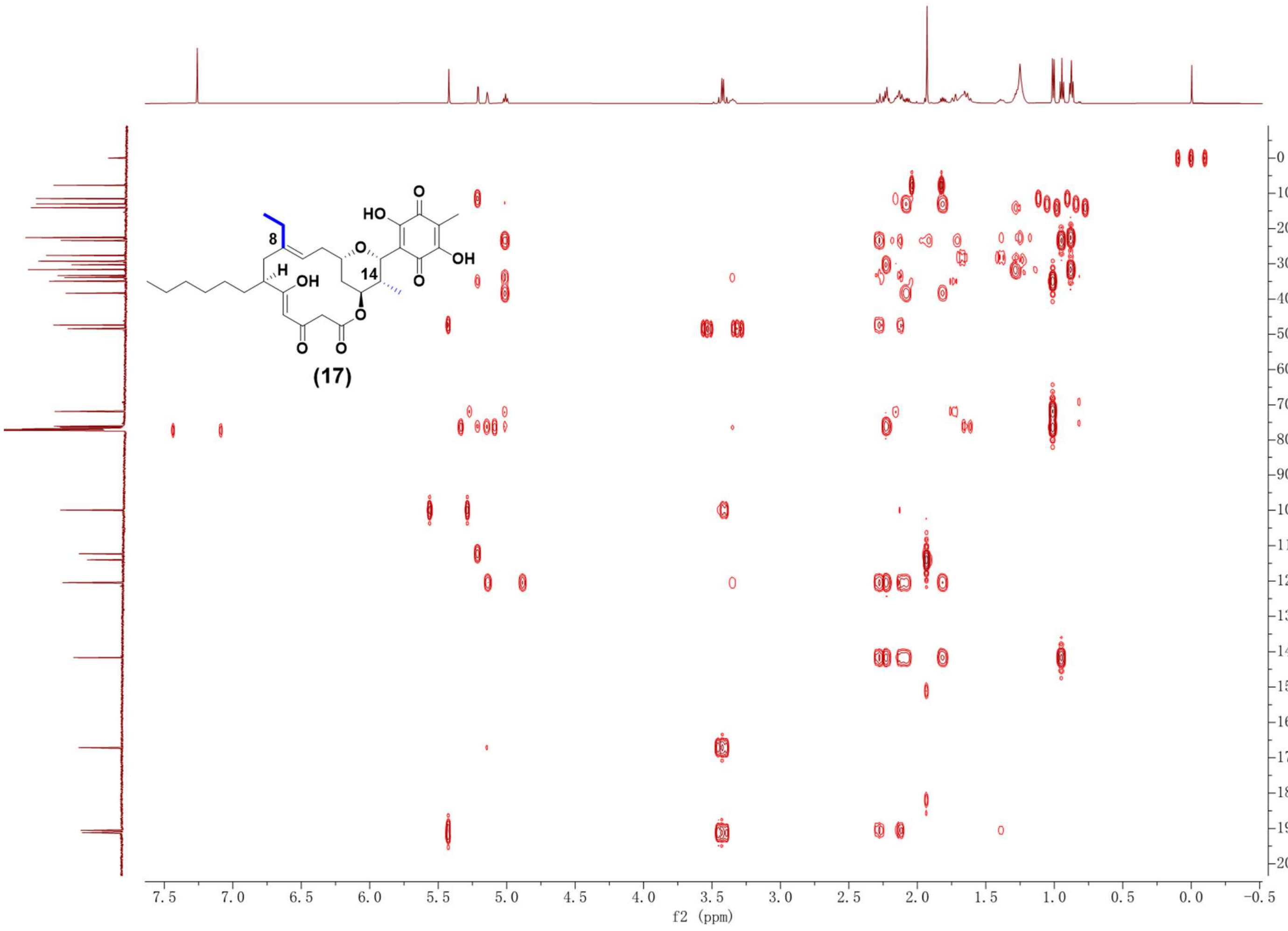

Supplementary Fig. 109**.** HMBC NMR spectrum (600 MHz, $CDCl_3$) of compound **17**.

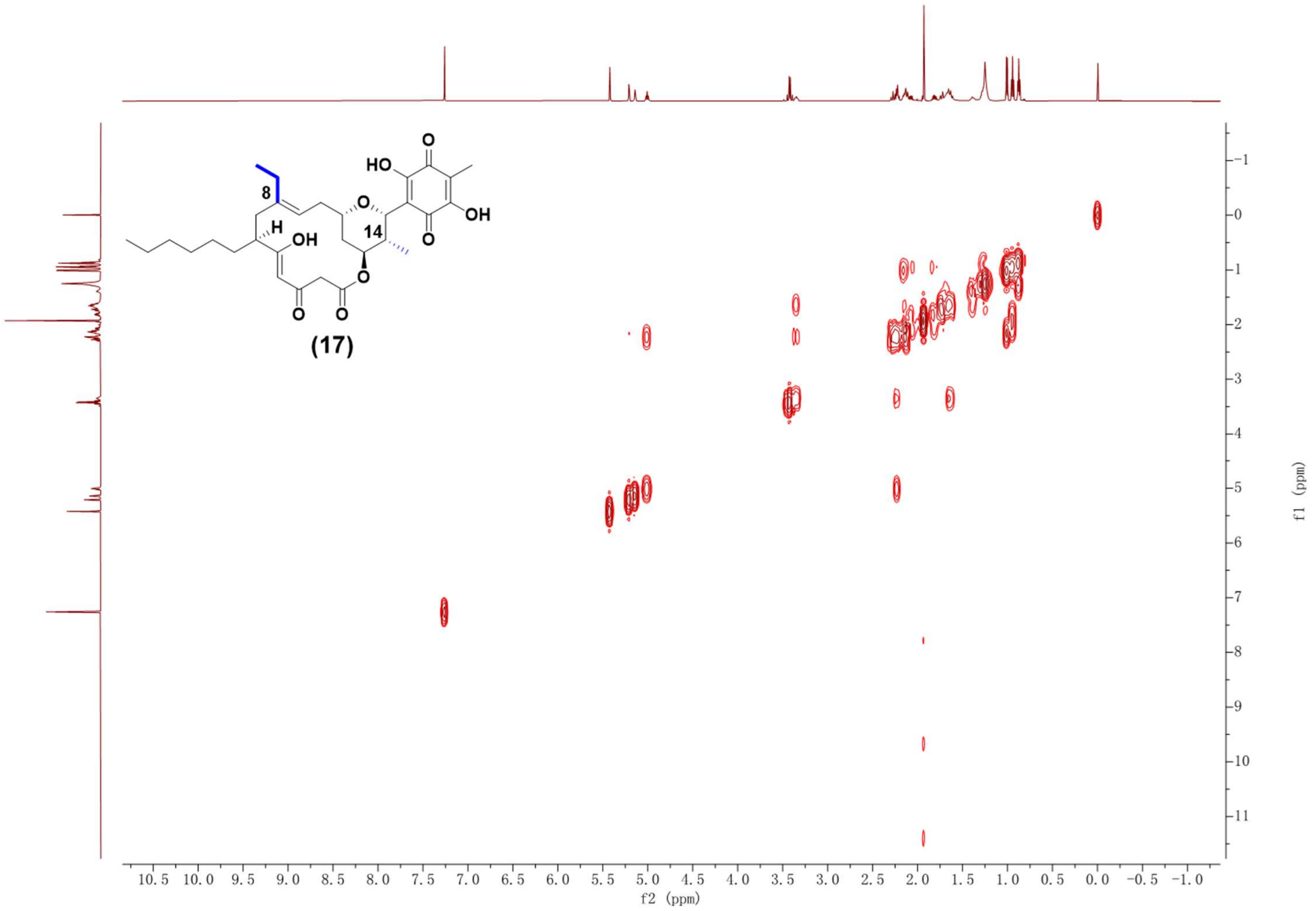

Supplementary Fig. 110. $^{1}H$-$^{1}H$ COSY NMR spectrum (600 MHz, $CDCl_3$) of compound **17**.

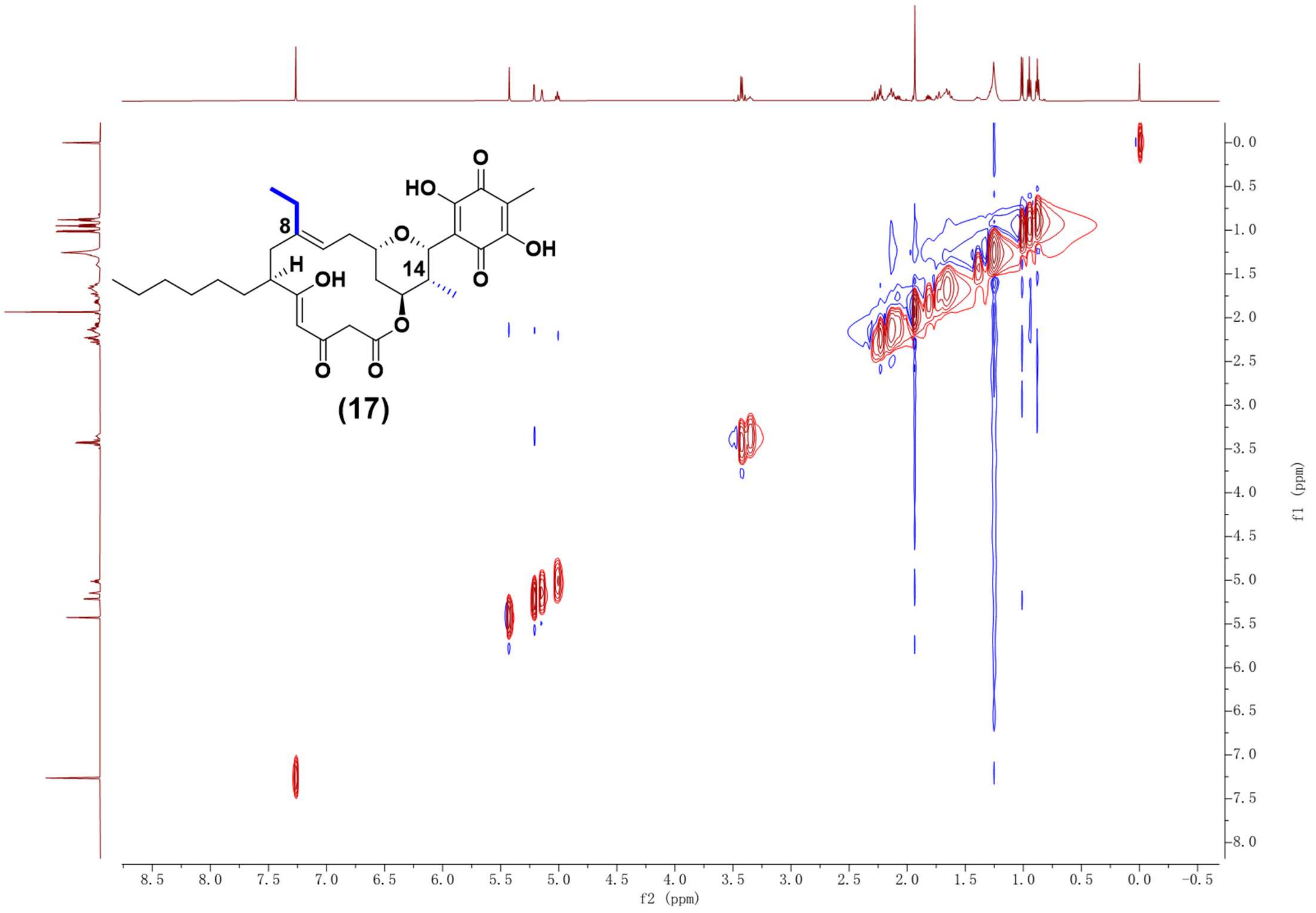

Supplementary Fig. 111. NOESY NMR spectrum (600 MHz, $CDCl_3$) of compound **17**.

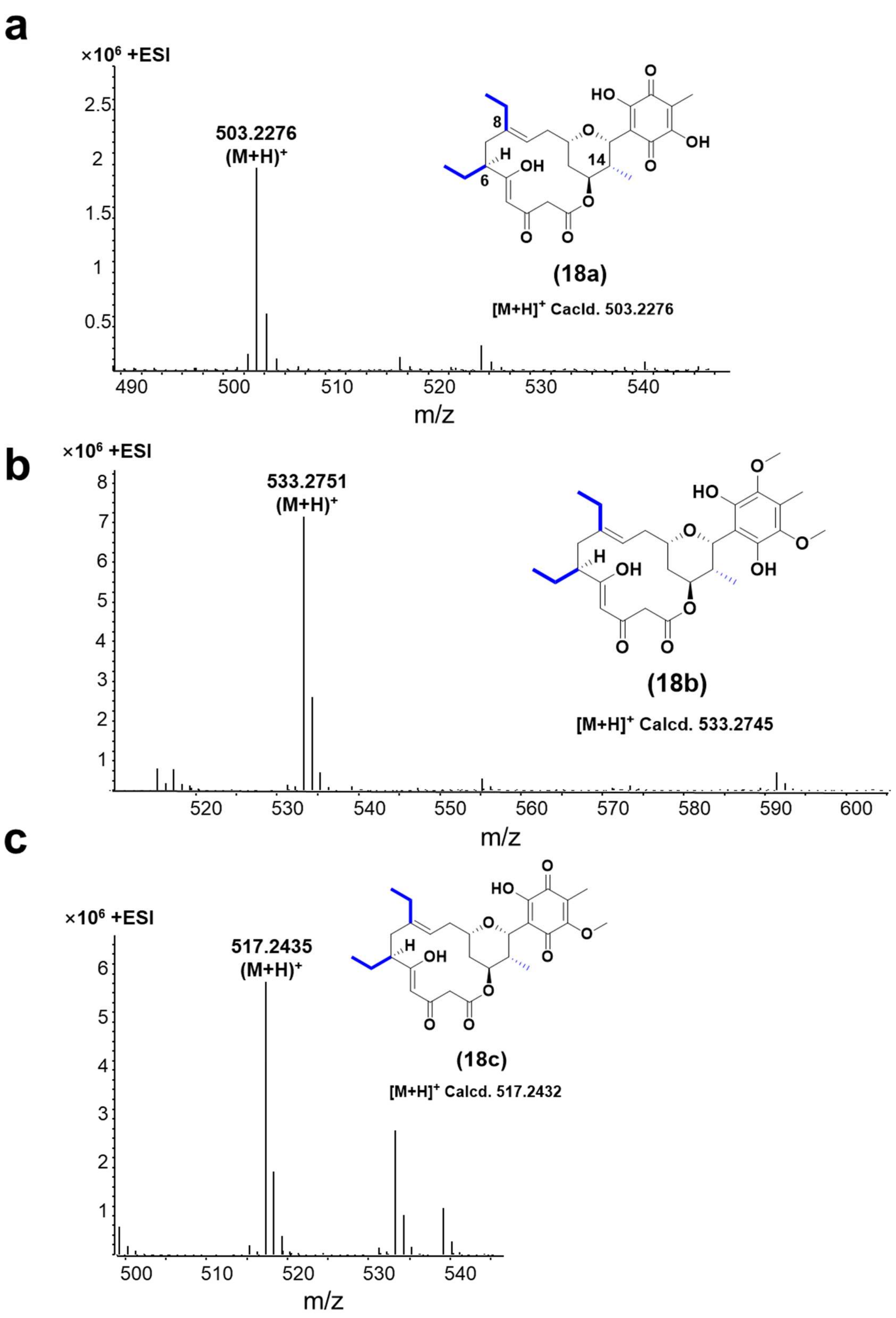

Supplementary Fig. 112. HR-ESI-MS spectrums of compounds **18a-18c**.

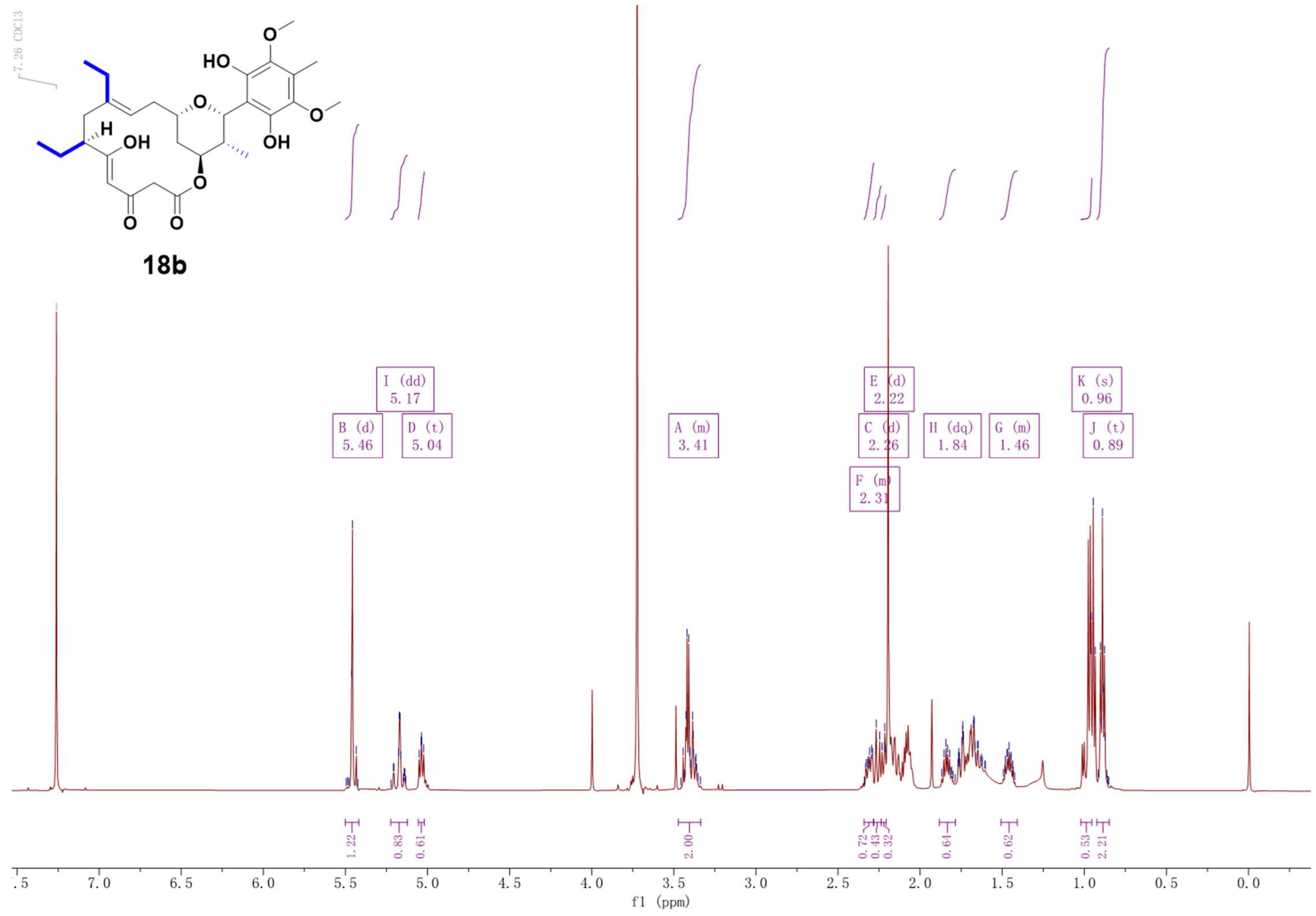

Supplementary Fig. 113. $^{1}$H NMR spectrum (600 MHz, $CDCl_3$) of compound **18b**.

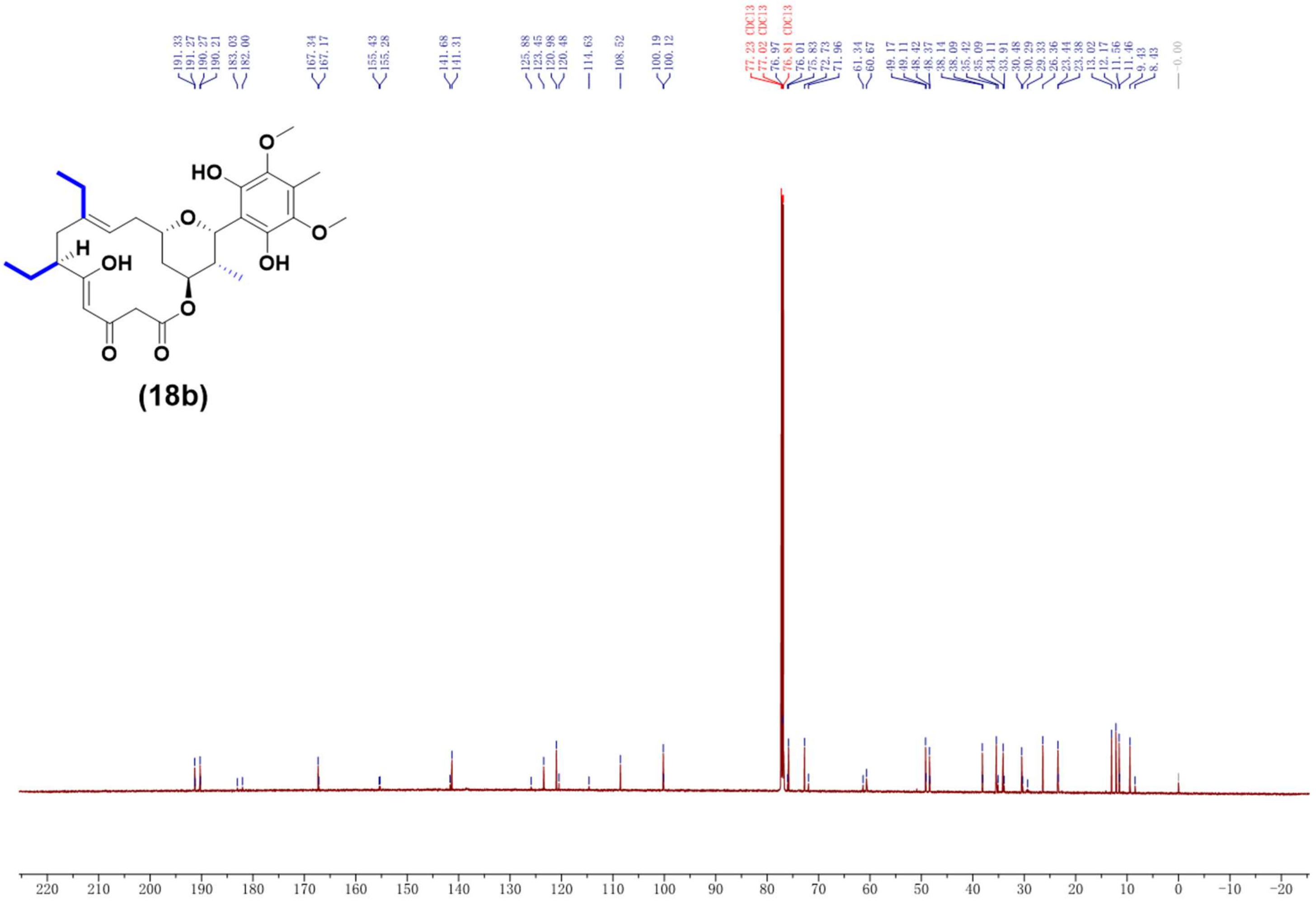

Supplementary Fig. 114. $^{13}$C NMR spectrum (150 MHz, $CDCl_3$) of compound **18b**.

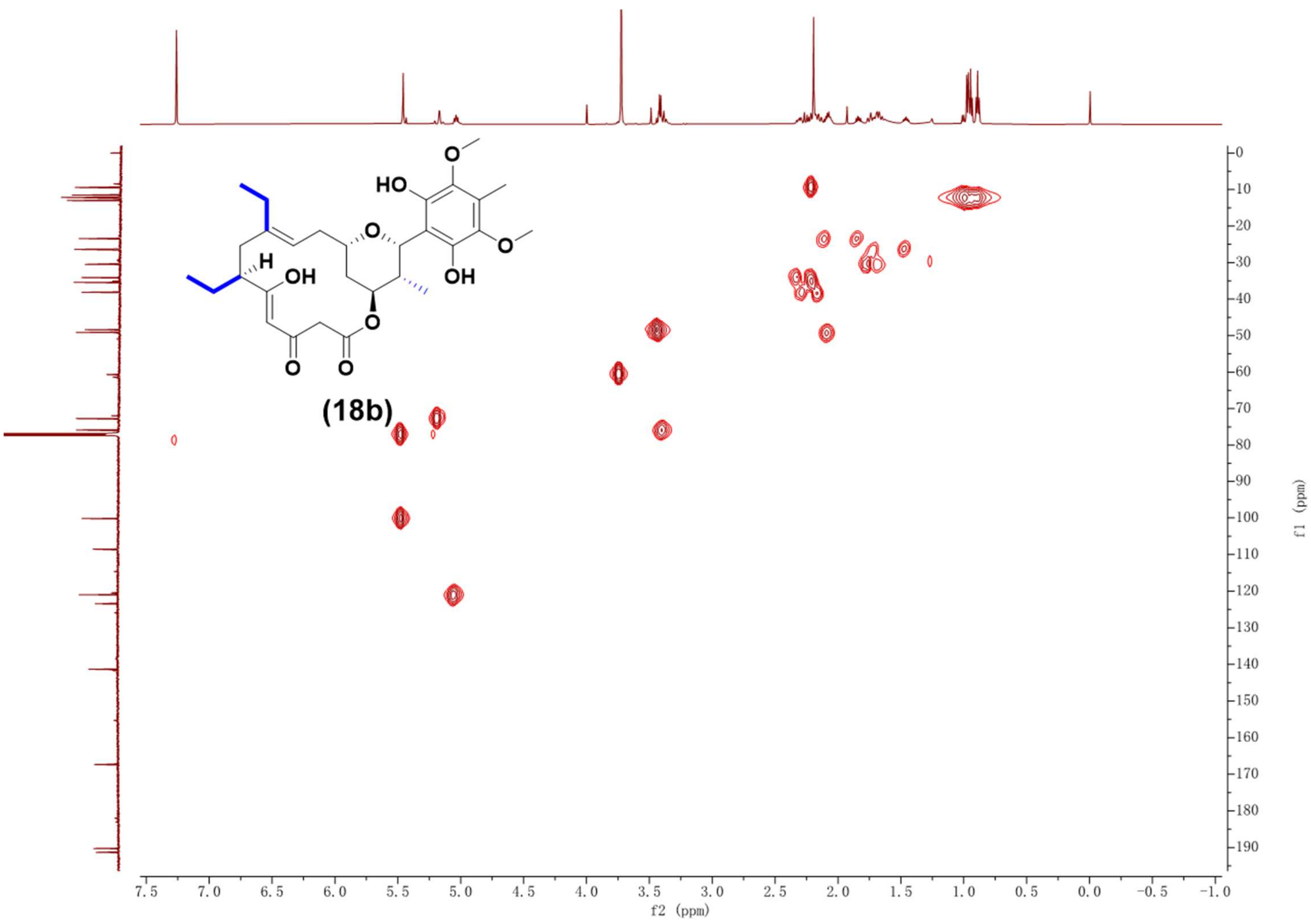

Supplementary Fig. 115. HSQC NMR spectrum (600 MHz, $CDCl_3$) of compound **18b**.

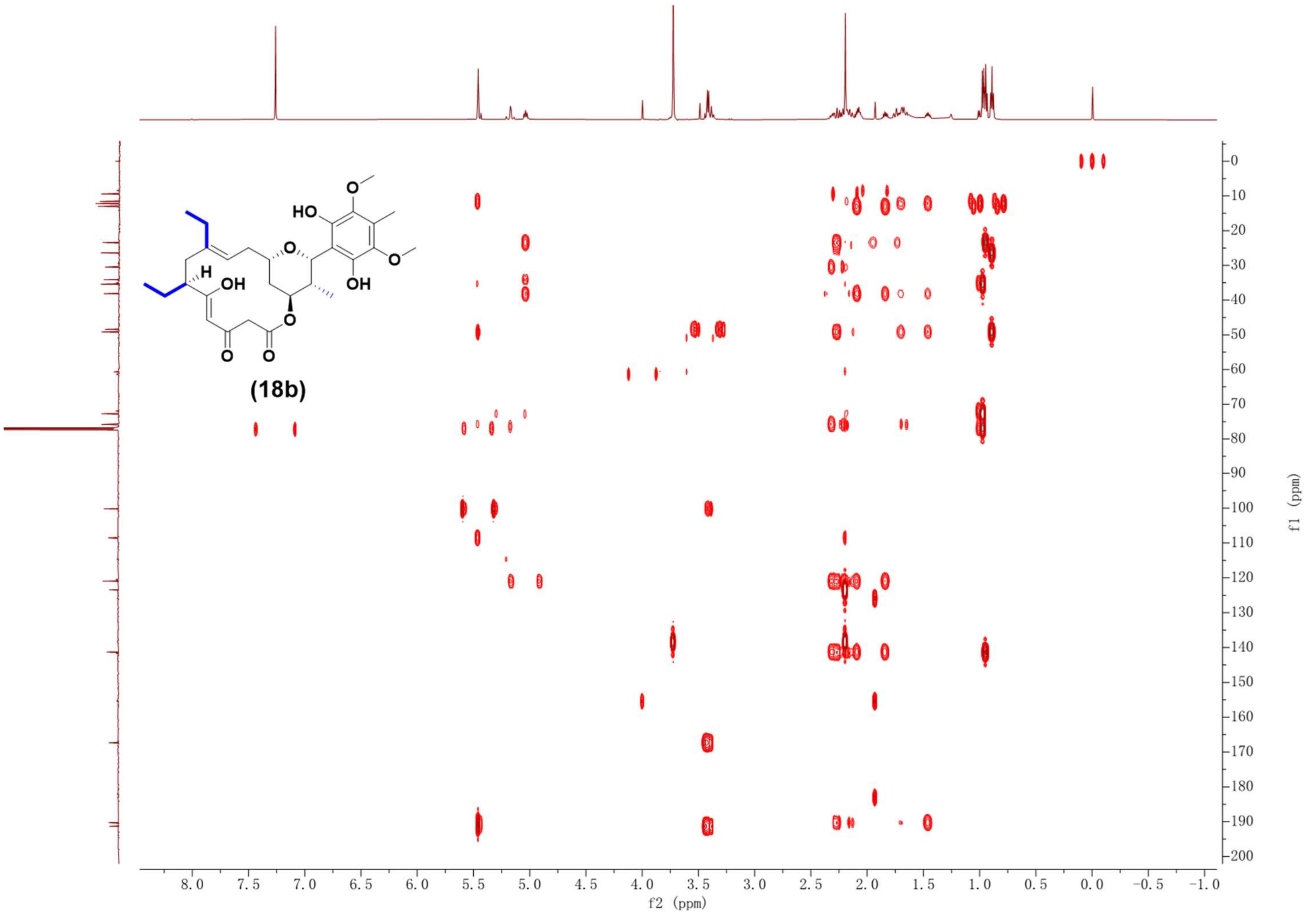

Supplementary Fig. 116. HMBC NMR spectrum (600 MHz, $CDCl_3$) of compound **18b**.

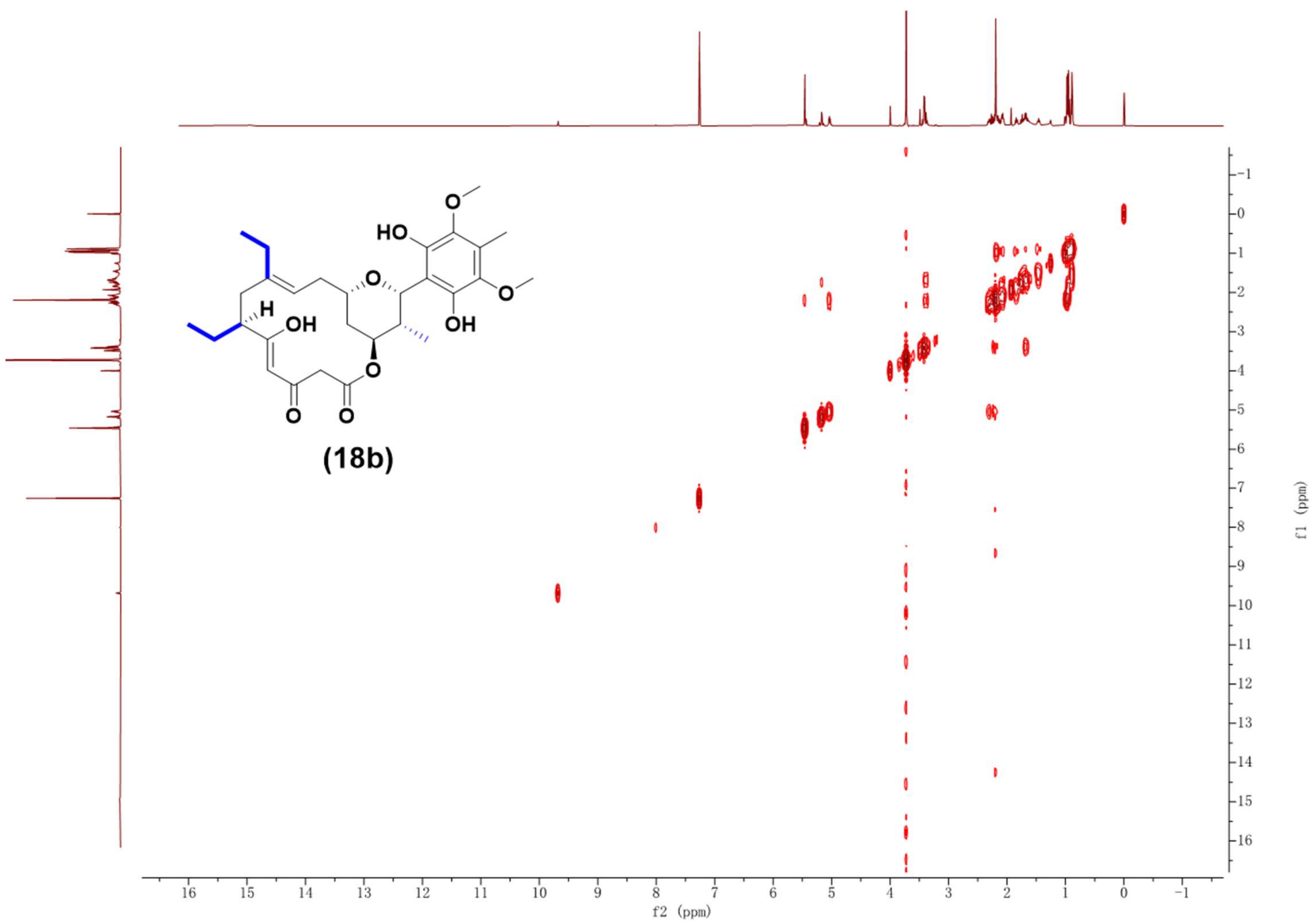

Supplementary Fig. 117. $^1$H-$^1$H COSY NMR spectrum (600 MHz, $CDCl_3$) of compound **18b**.

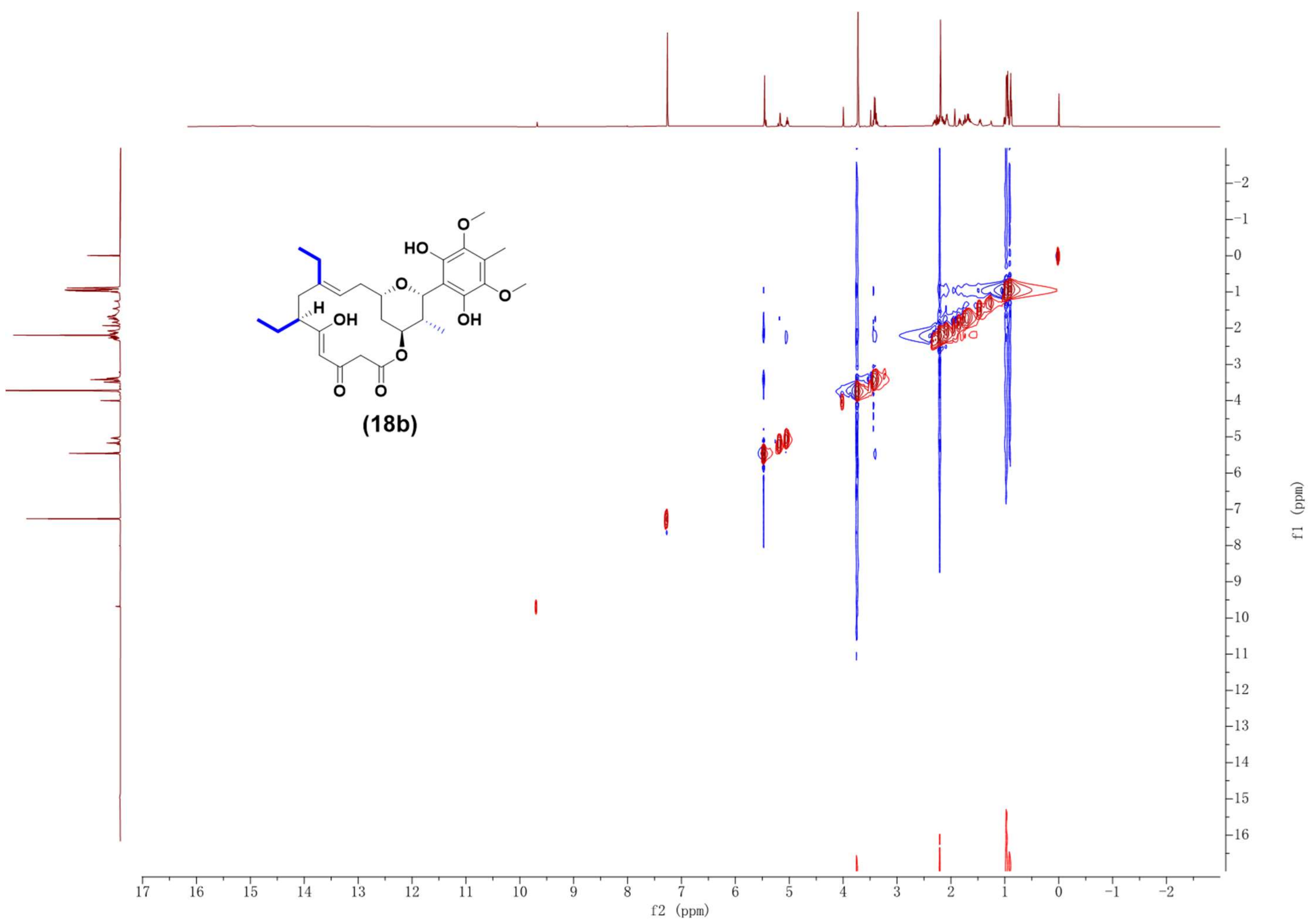

Supplementary Fig. 118. NOESY NMR spectrum (600 MHz, $CDCl_3$) of compound **18b**.

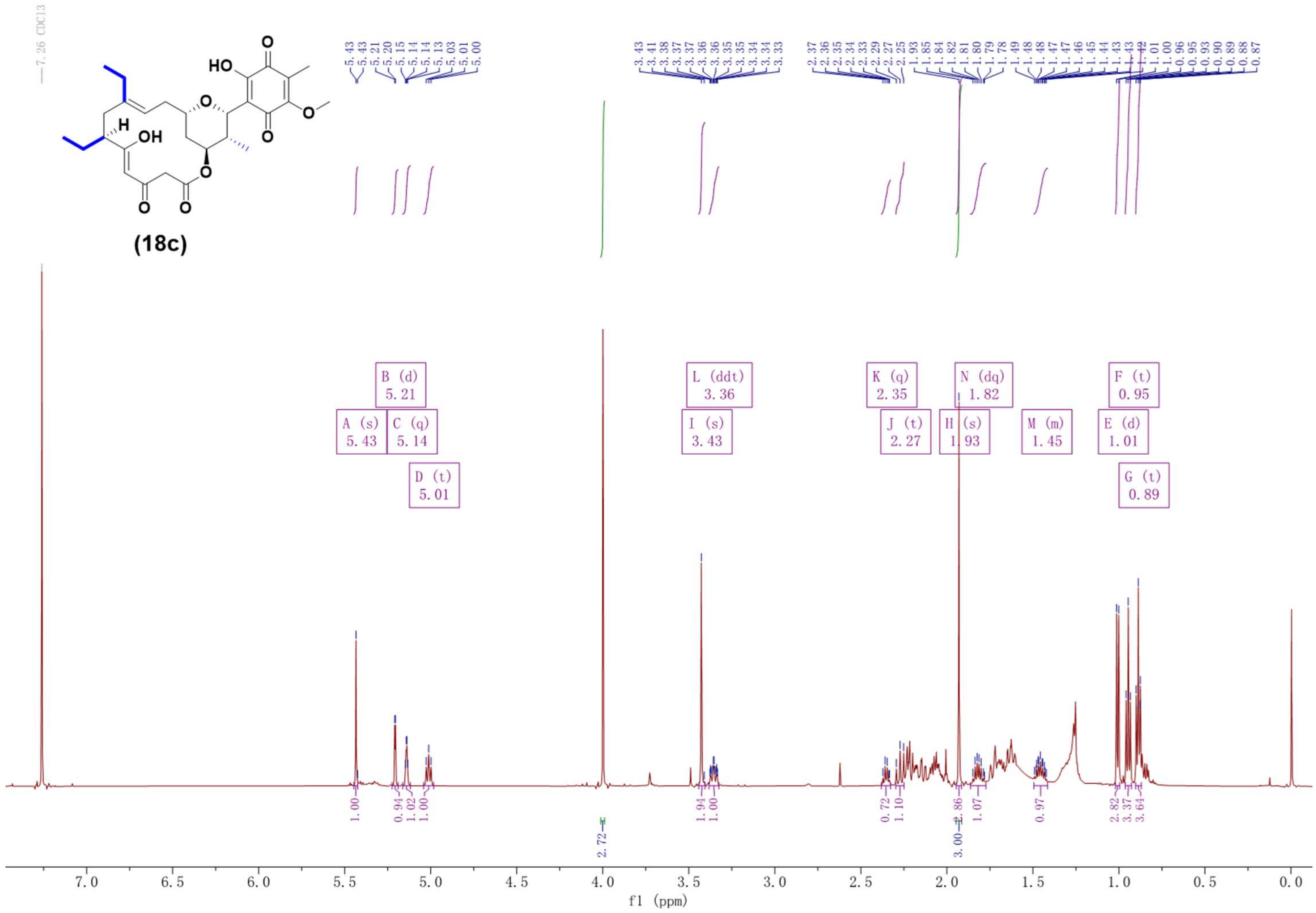

Supplementary Fig. 119. $^{1}H$ NMR spectrum (600 MHz, $CDCl_3$) of compound **18c**.

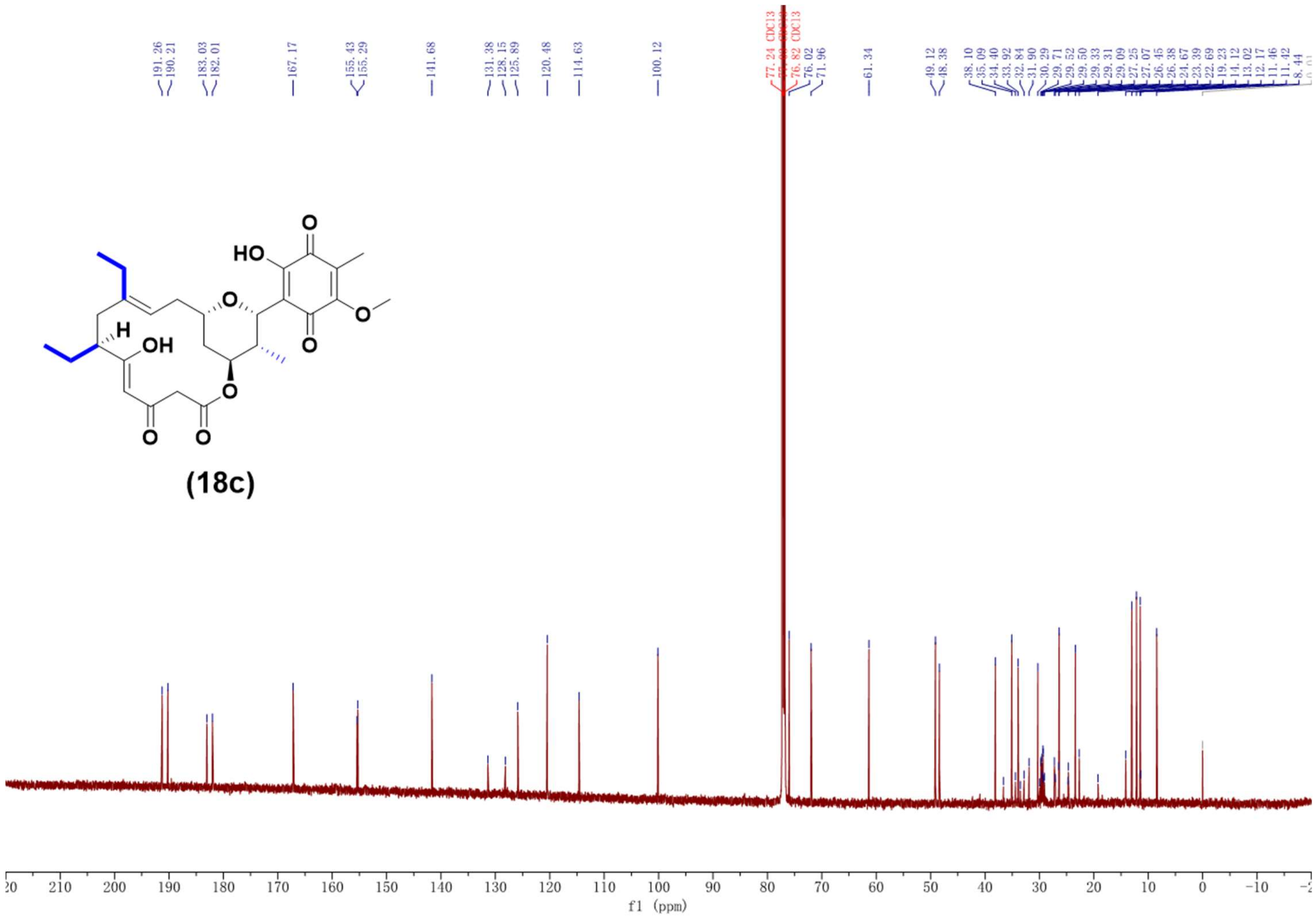

Supplementary Fig. 120. $^{13}C$ NMR spectrum (150 MHz, $CDCl_3$) of compound **18c**.

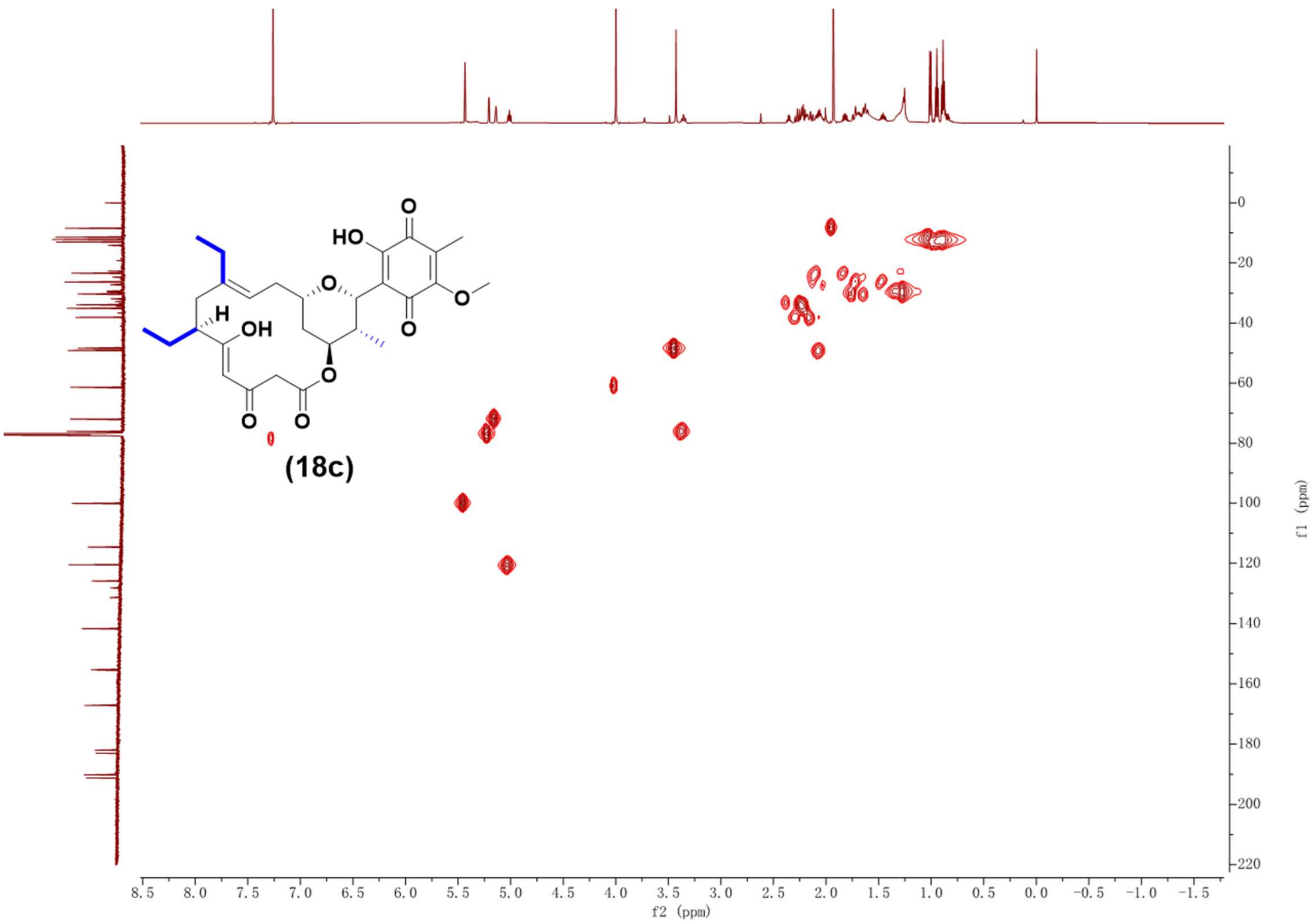

Supplementary Fig. 121. HSQC NMR spectrum (600 MHz, $CDCl_3$) of compound **18c**.

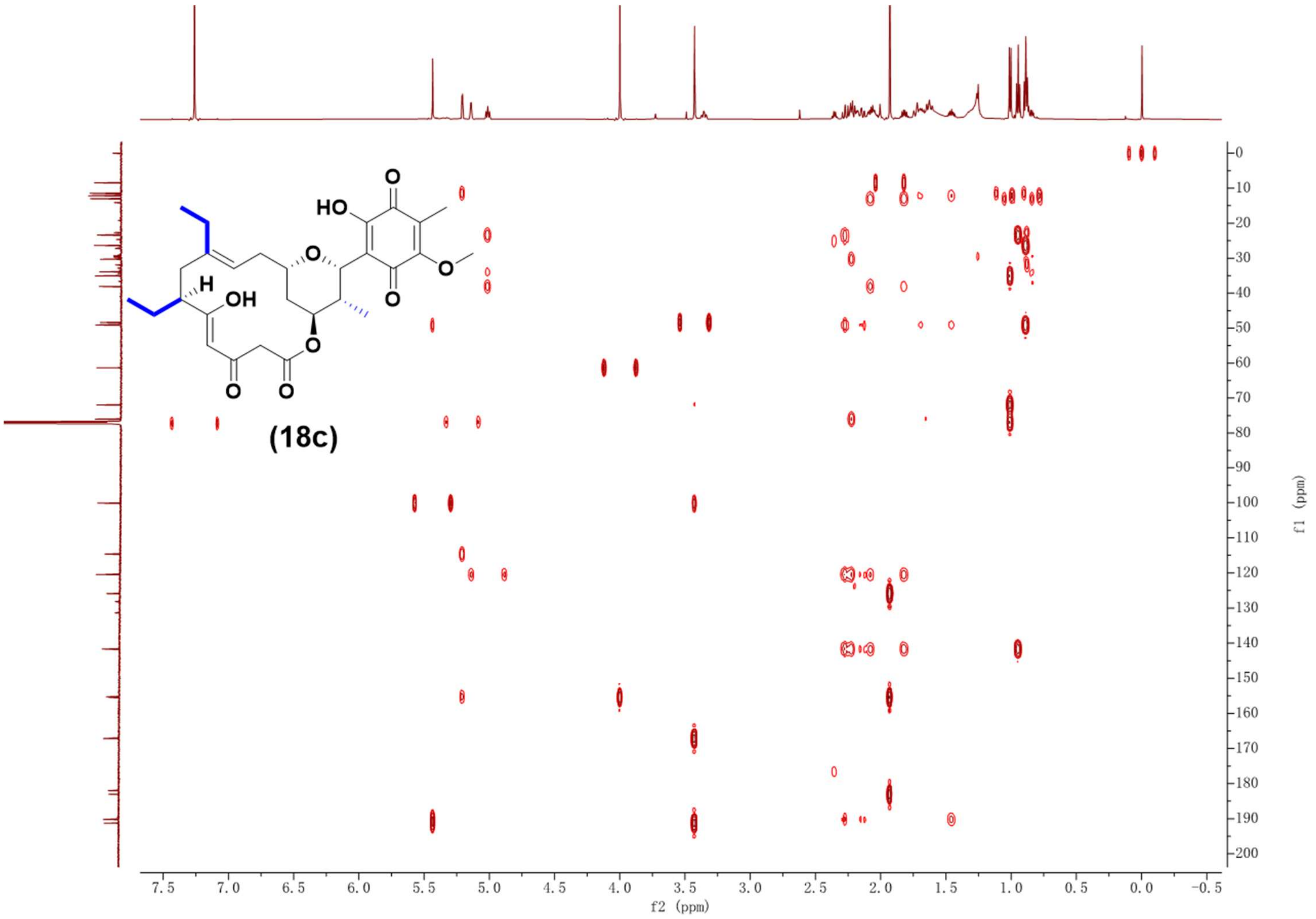

Supplementary Fig. 122. HMBC NMR spectrum (600 MHz, $CDCl_3$) of compound **18c**.

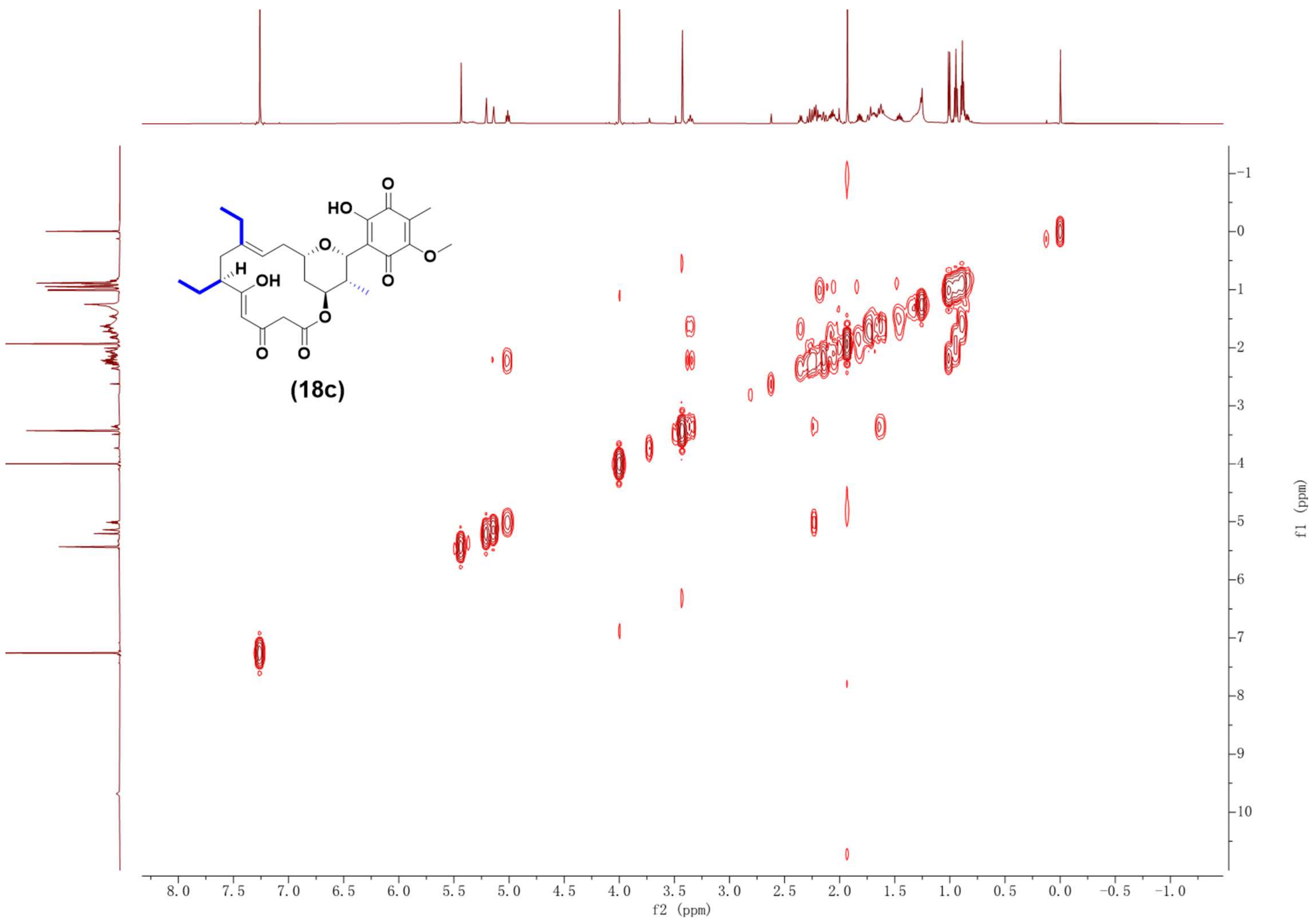

Supplementary Fig. 123. $^{1}H$-$^{1}H$ COSY NMR spectrum (600 MHz, $CDCl_3$) of compound **18c**.

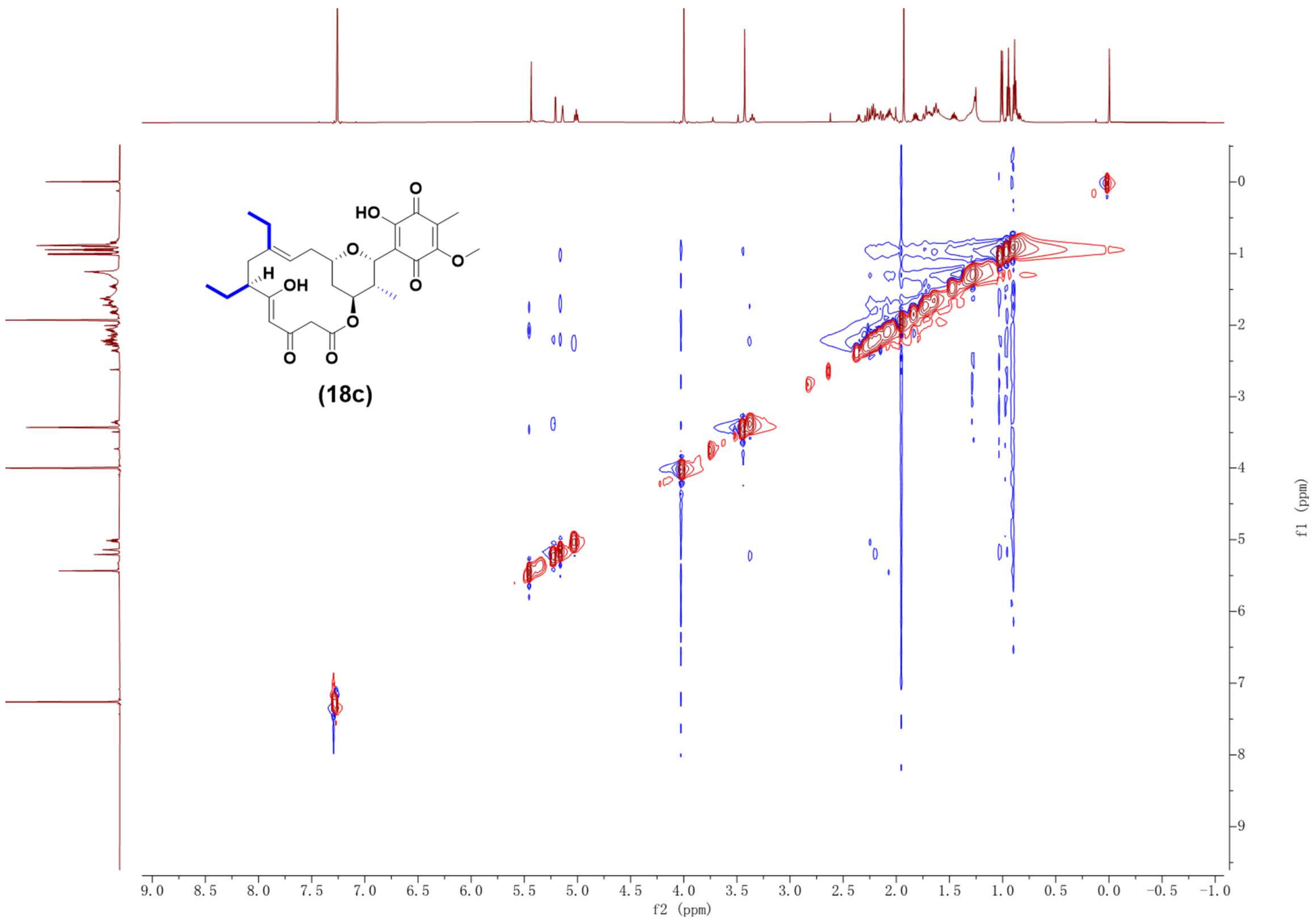

Supplementary Fig. 124. NOESY NMR spectrum (600 MHz, $CDCl_3$) of compound **18c**.

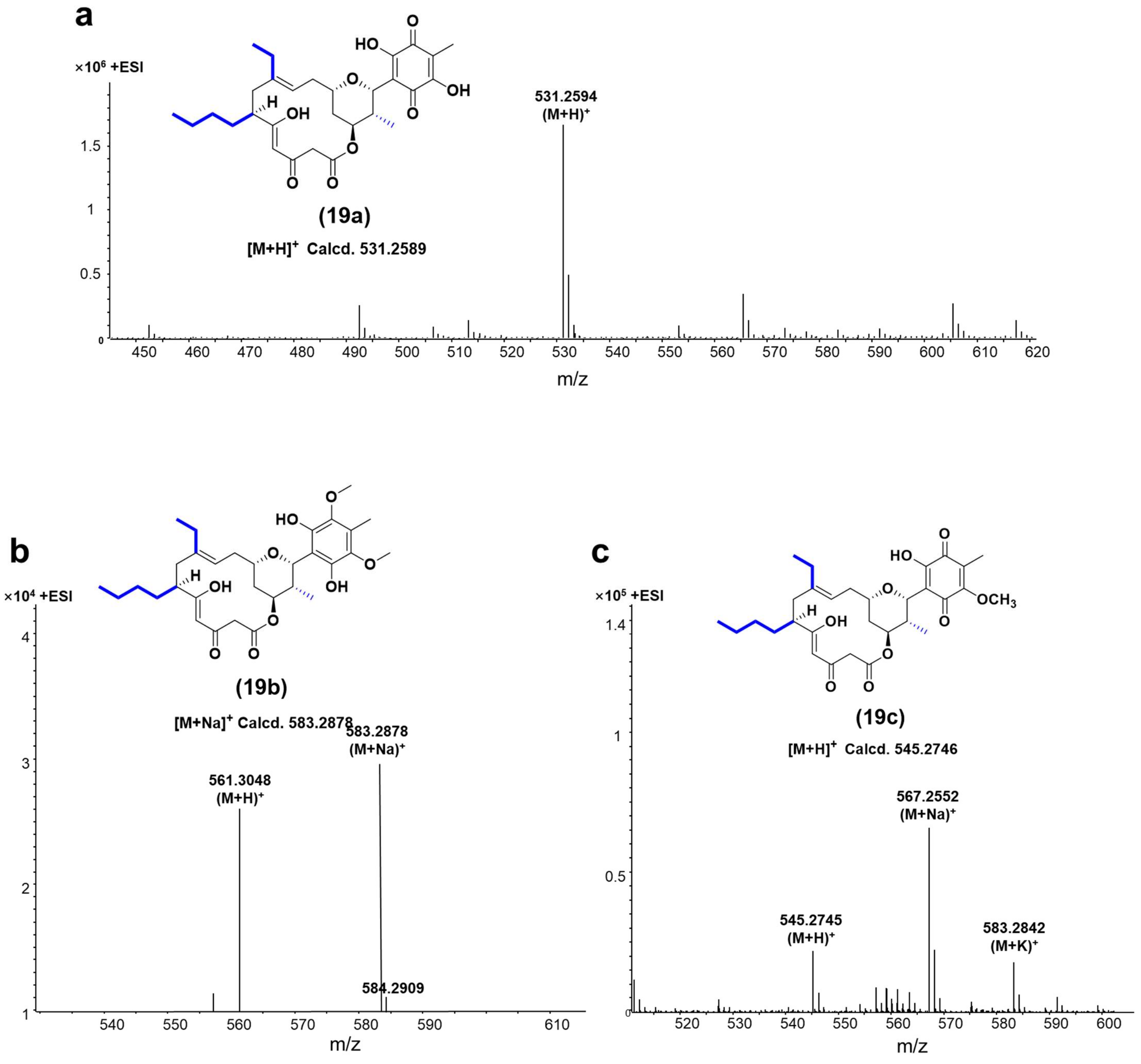

Supplementary Fig. 125. HR-ESI-MS spectrums of compounds **19a-19c**.

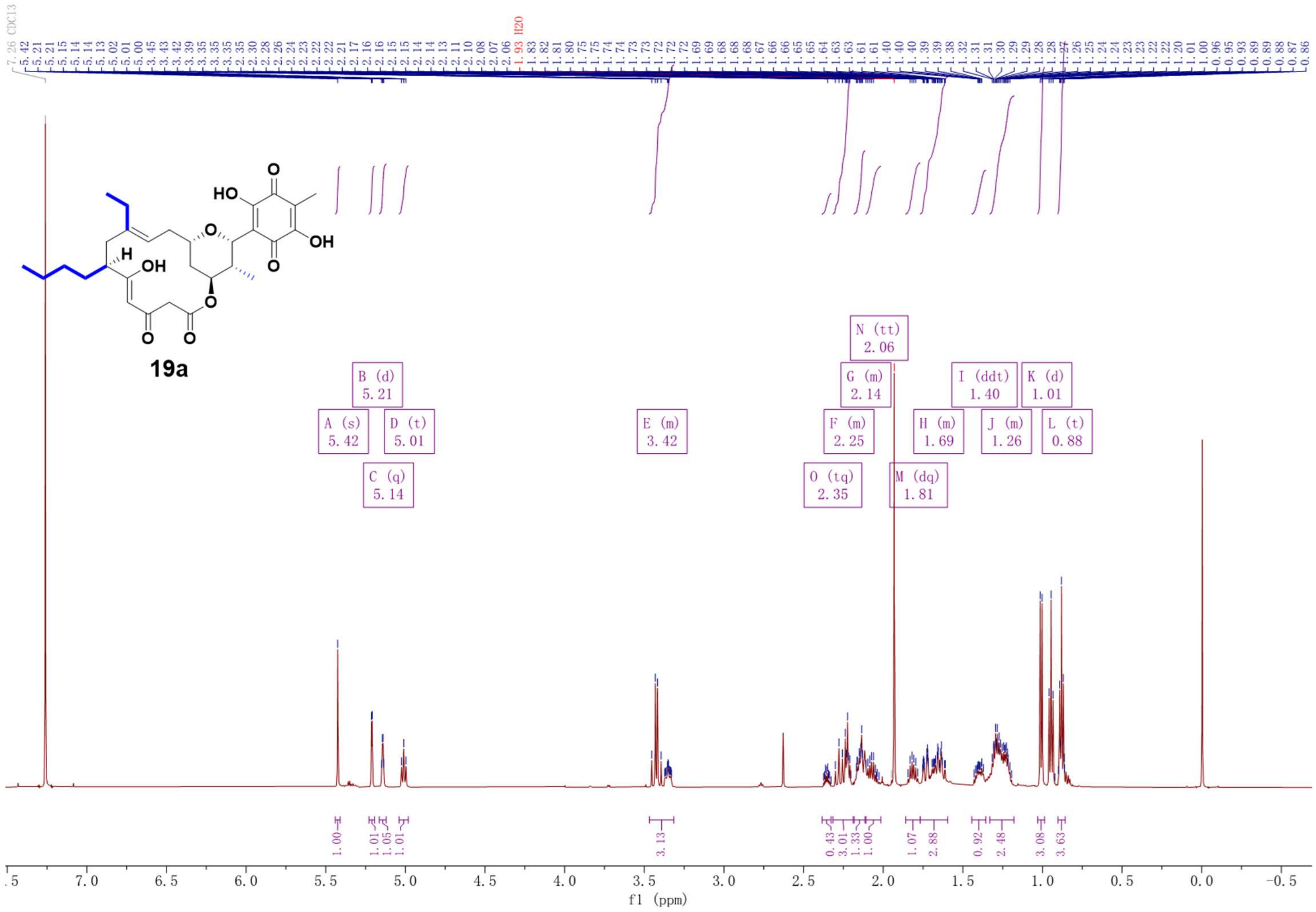

Supplementary Fig. 126. $^{1}H$ NMR spectrum (600 MHz, $CDCl_3$) of compound **19a**.

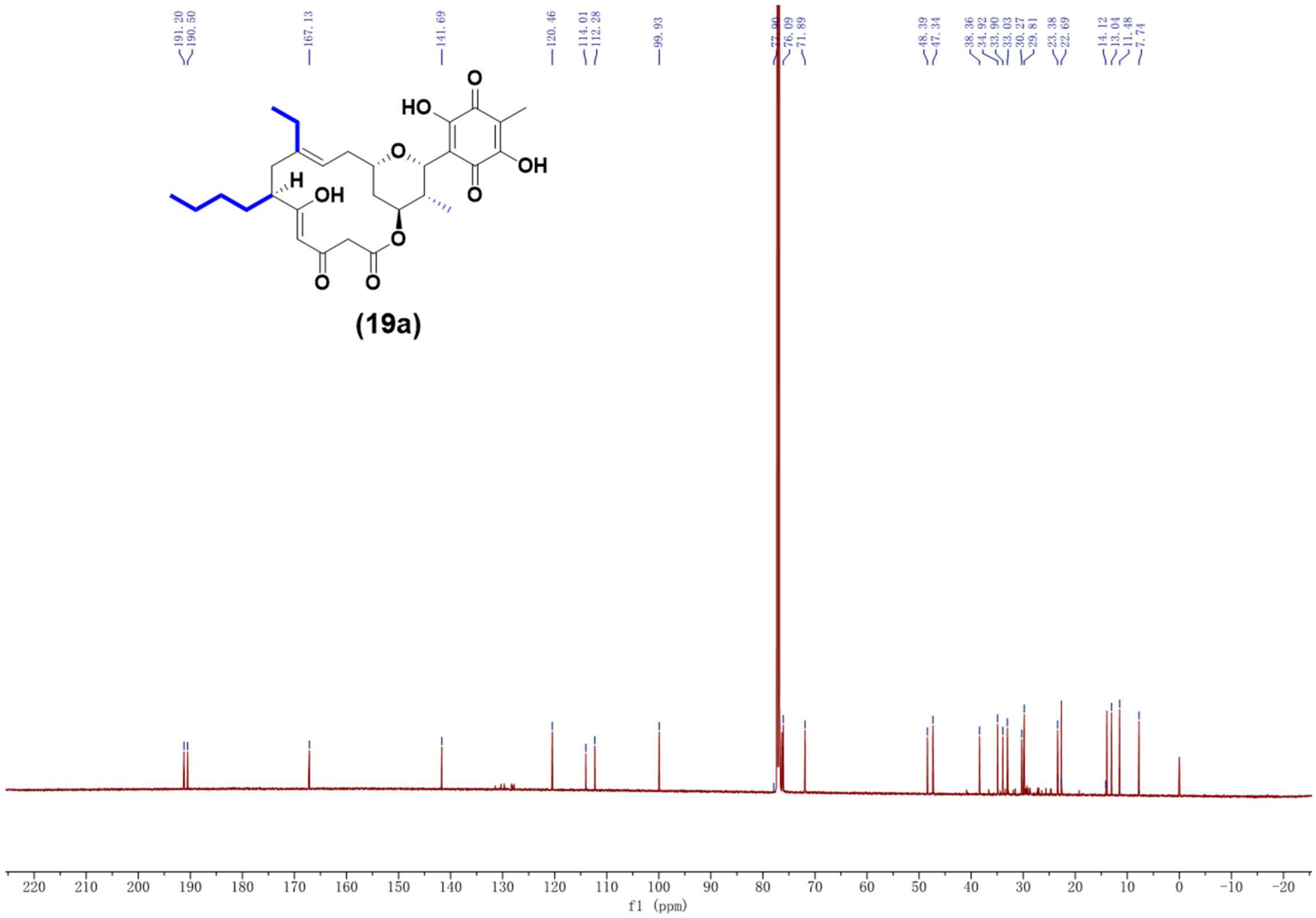

Supplementary Fig. 127. $^{13}C$ NMR spectrum (150 MHz, $CDCl_3$) of compound **19a**.

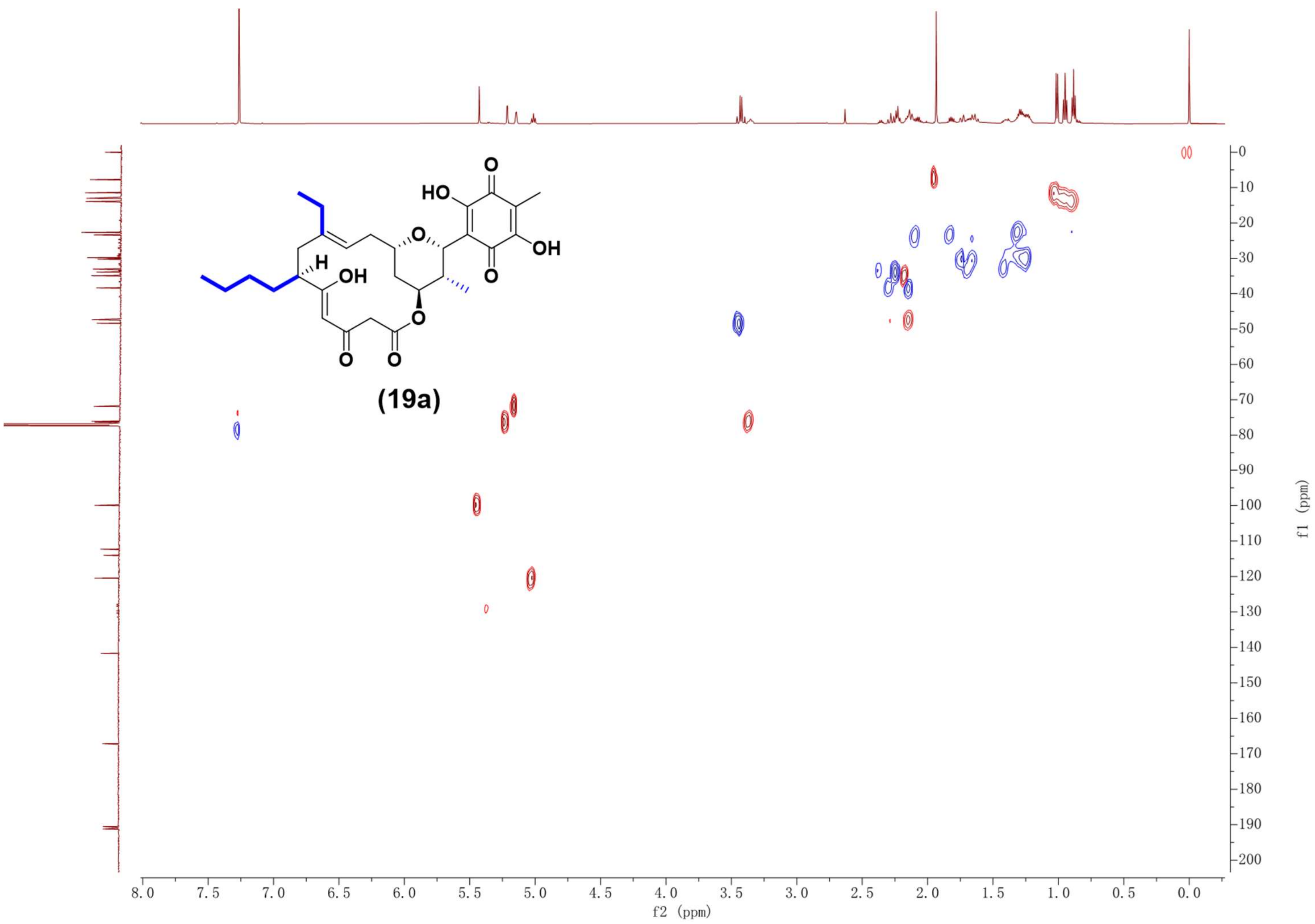

Supplementary Fig. 128. HSQC NMR spectrum (600 MHz, $CDCl_3$) of compound **19a**.

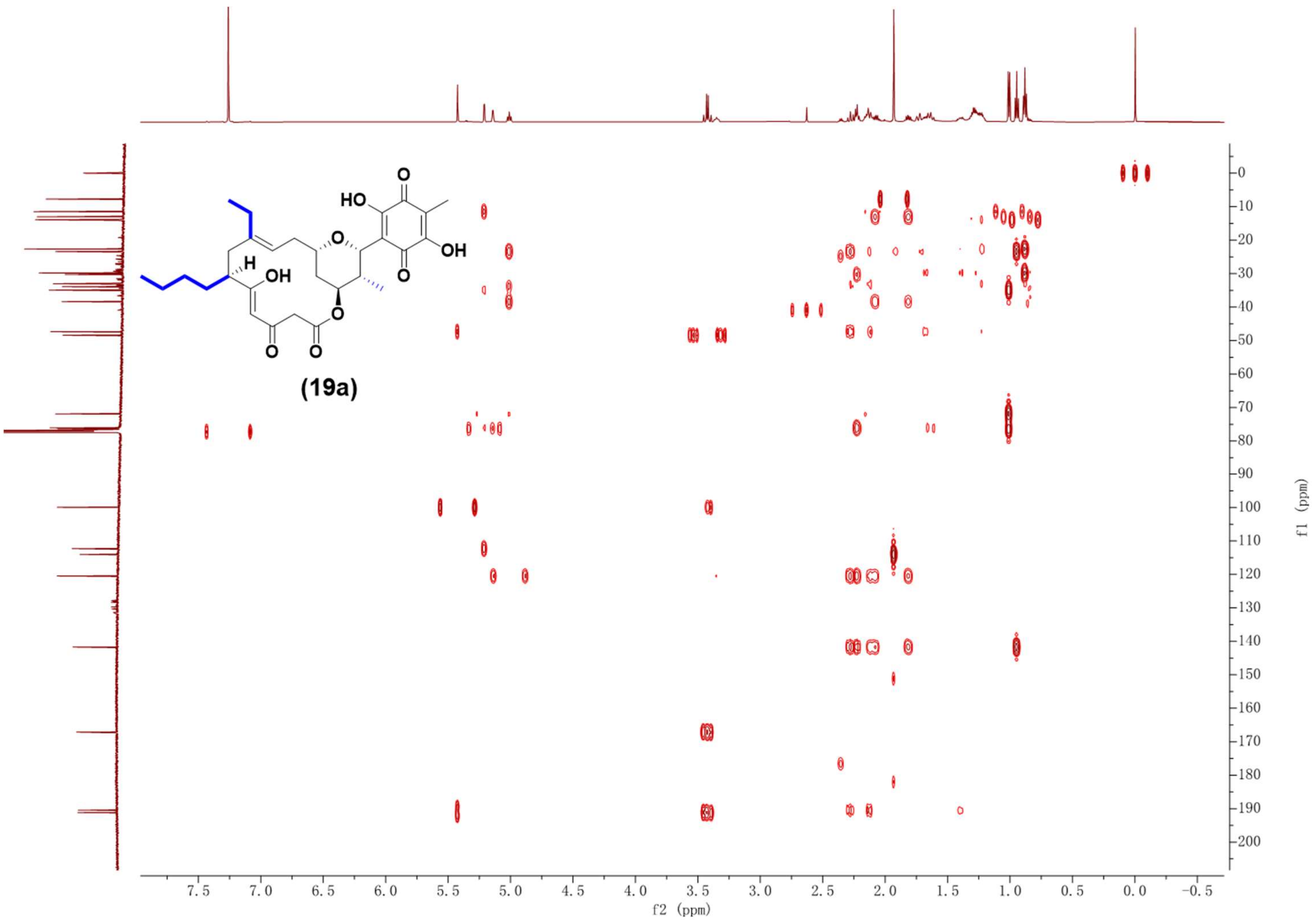

Supplementary Fig. 129. HMBC NMR spectrum (600 MHz, $CDCl_3$) of compound **19a**.

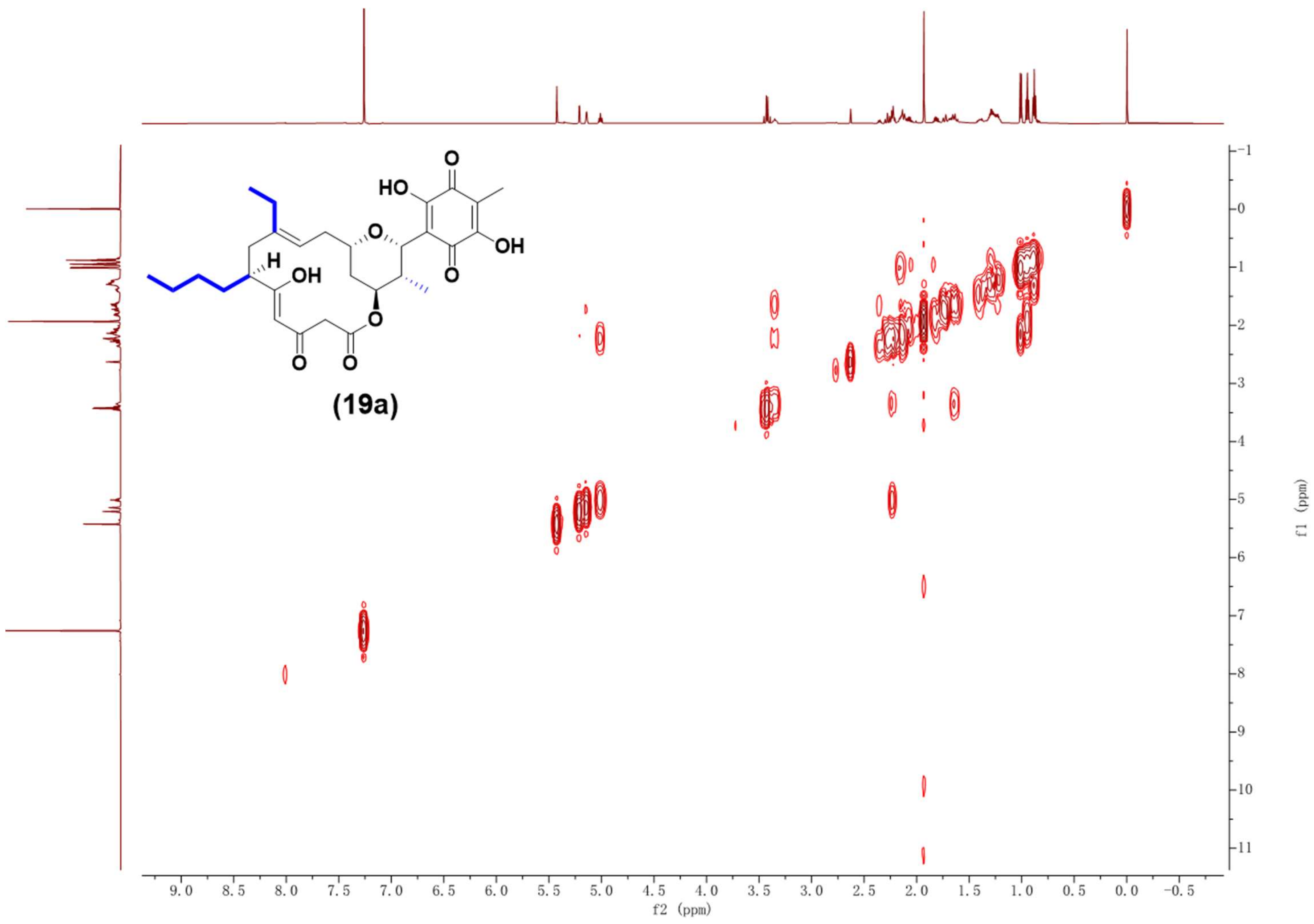

Supplementary Fig. 130. $^{1}H$-$^{1}H$ COSY NMR spectrum (600 MHz, $CDCl_3$) of compound **19a**.

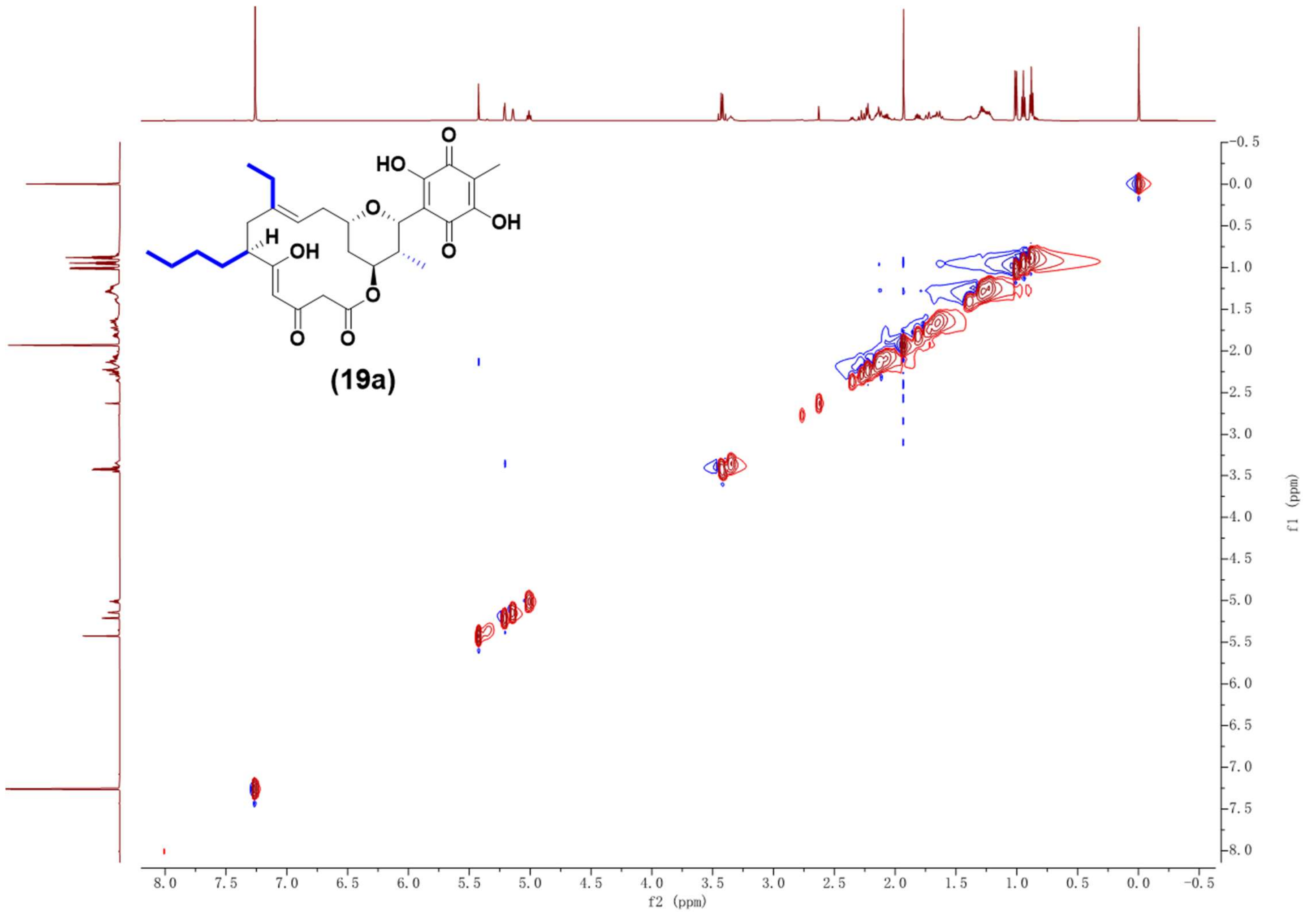

Supplementary Fig. 131. NOESY NMR spectrum (600 MHz, $CDCl_3$) of compound **19a**.

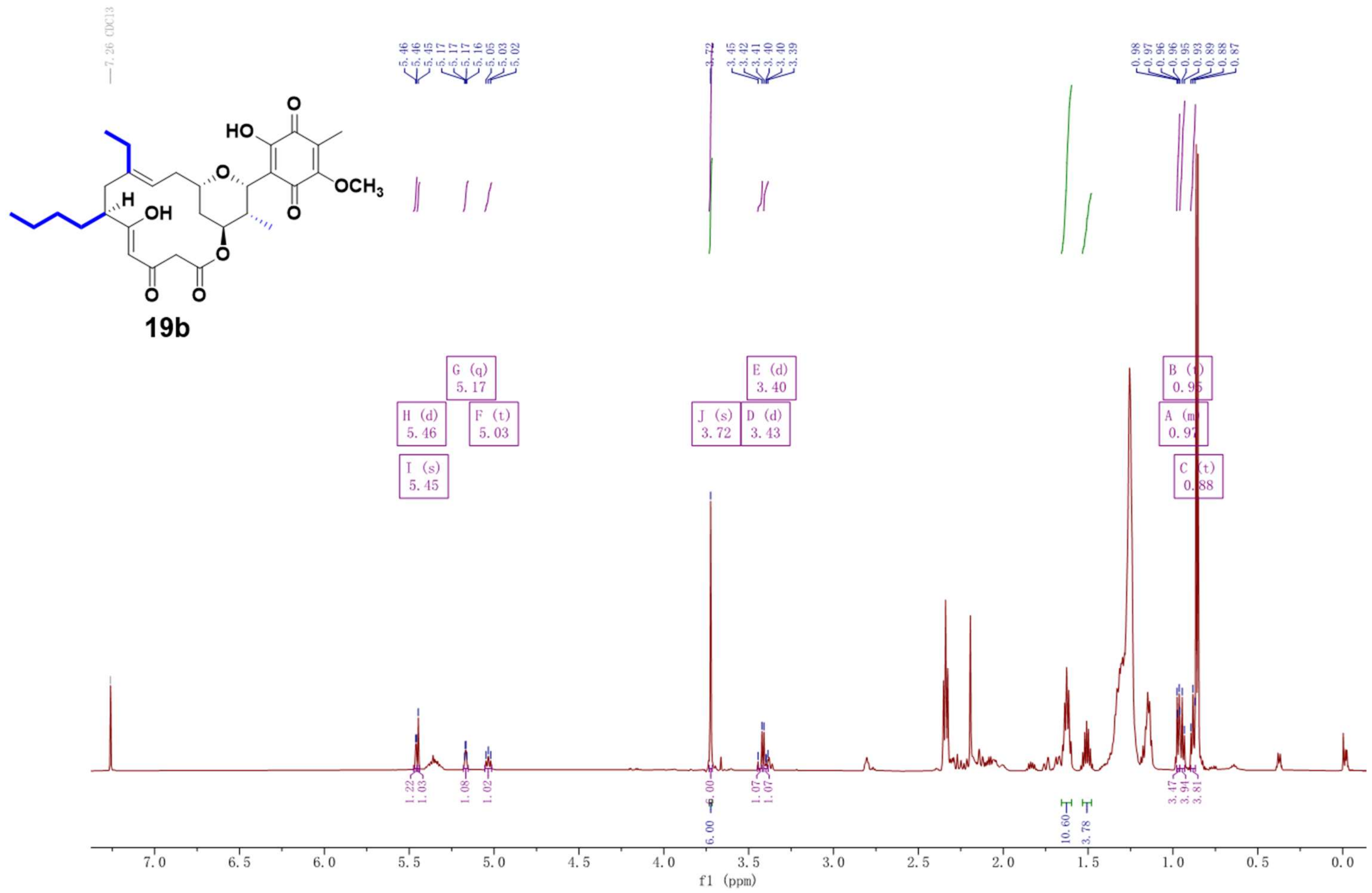

Supplementary Fig. 132. $^{1}H$ NMR spectrum (600 MHz, $CDCl_3$) of compound **19b**.

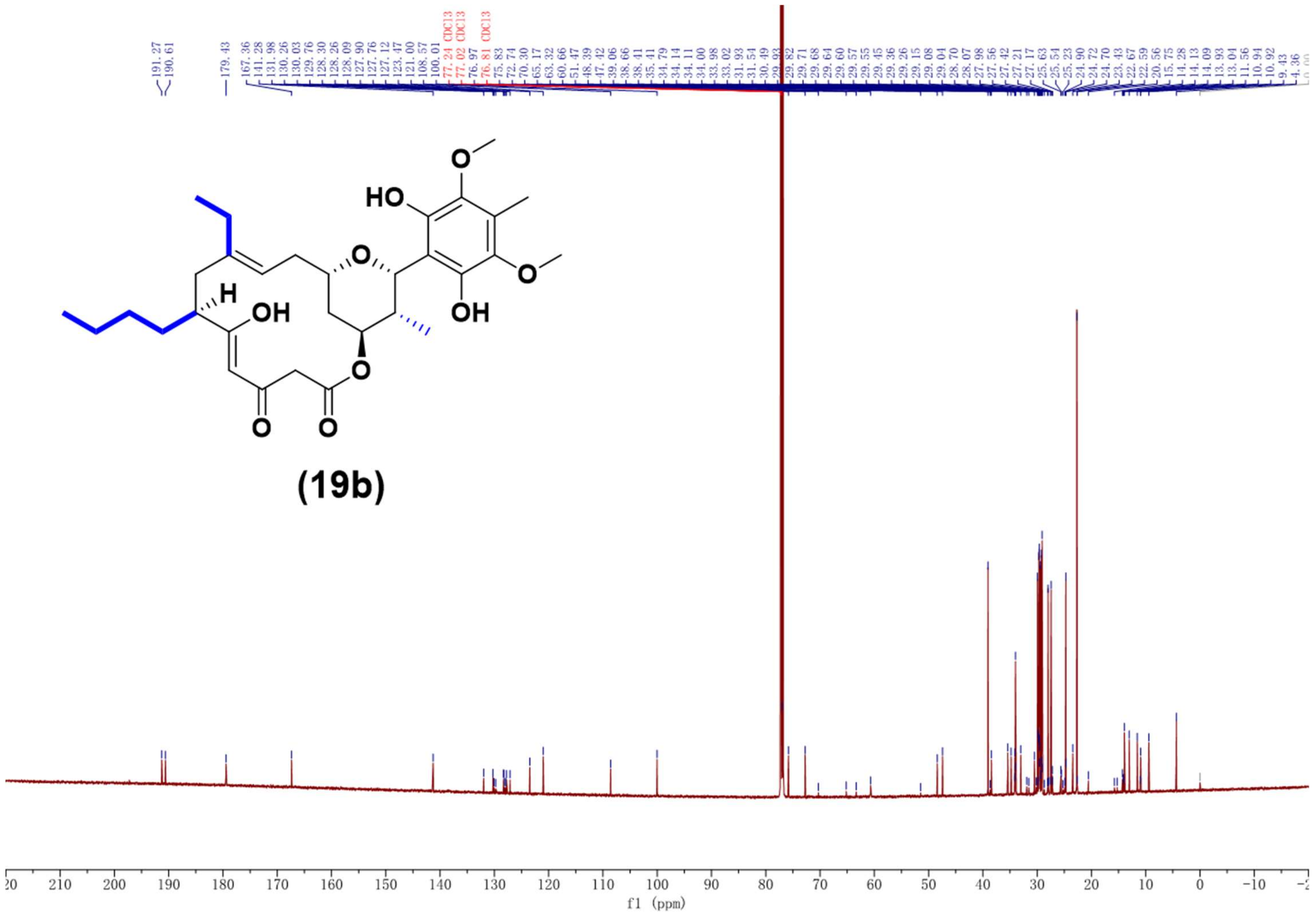

Supplementary Fig. 133. $^{13}C$ NMR spectrum (150 MHz, $CDCl_3$) of compound **19b**.

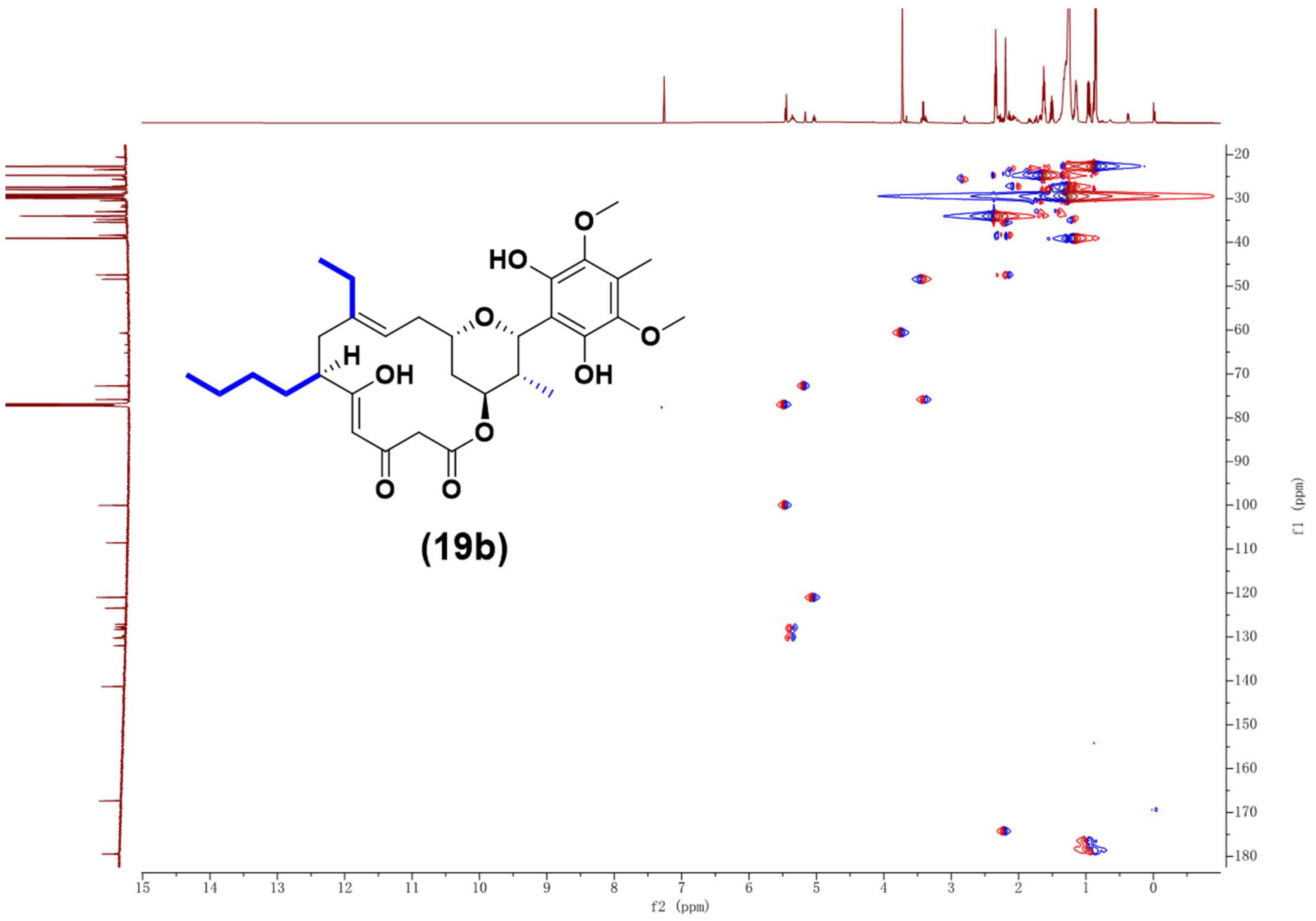

Supplementary Fig. 134. HSQC NMR spectrum (600 MHz, $CDCl_3$) of compound **19b**.

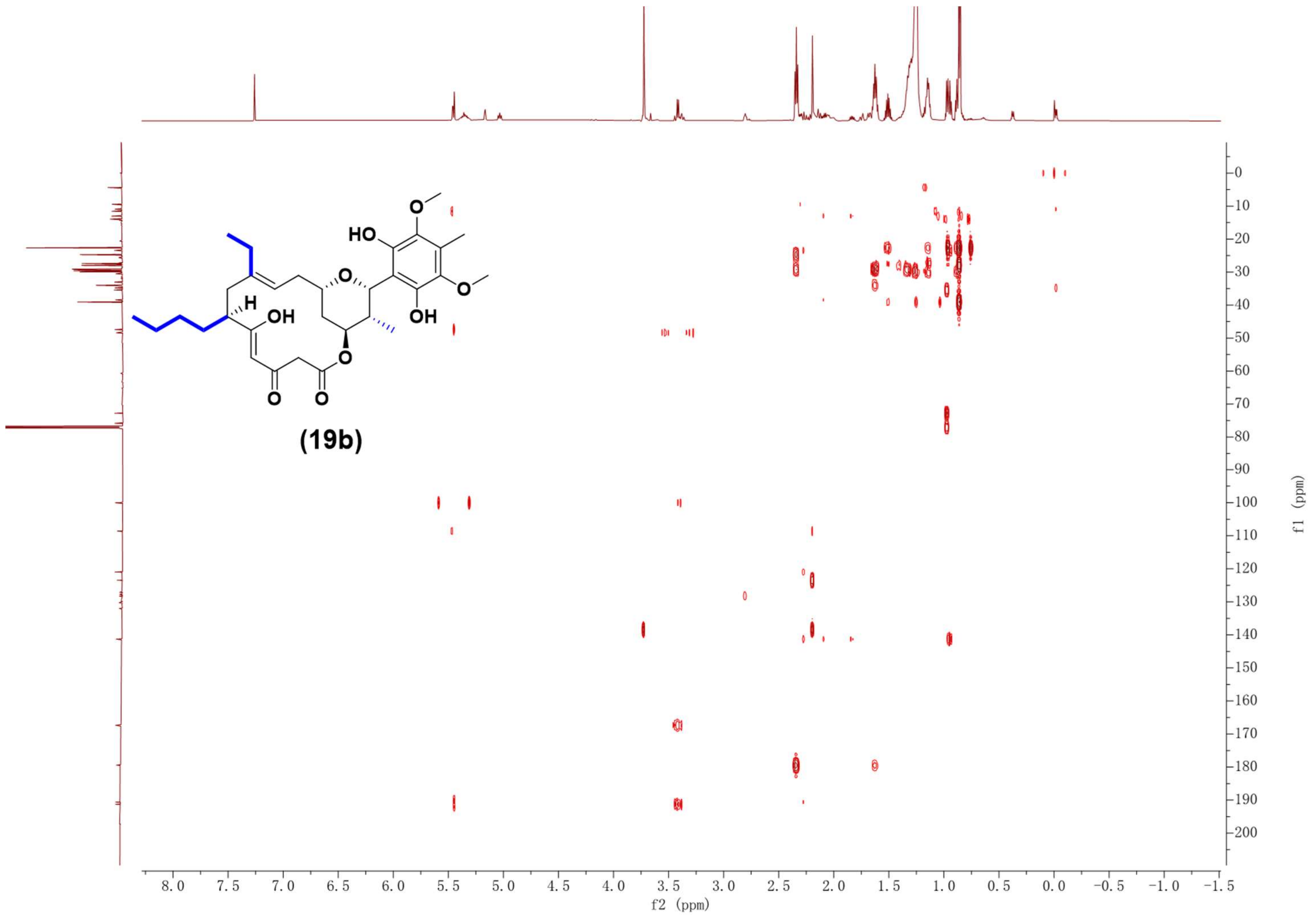

Supplementary Fig. 135. HMBC NMR spectrum (600 MHz, $CDCl_3$) of compound **19b**.

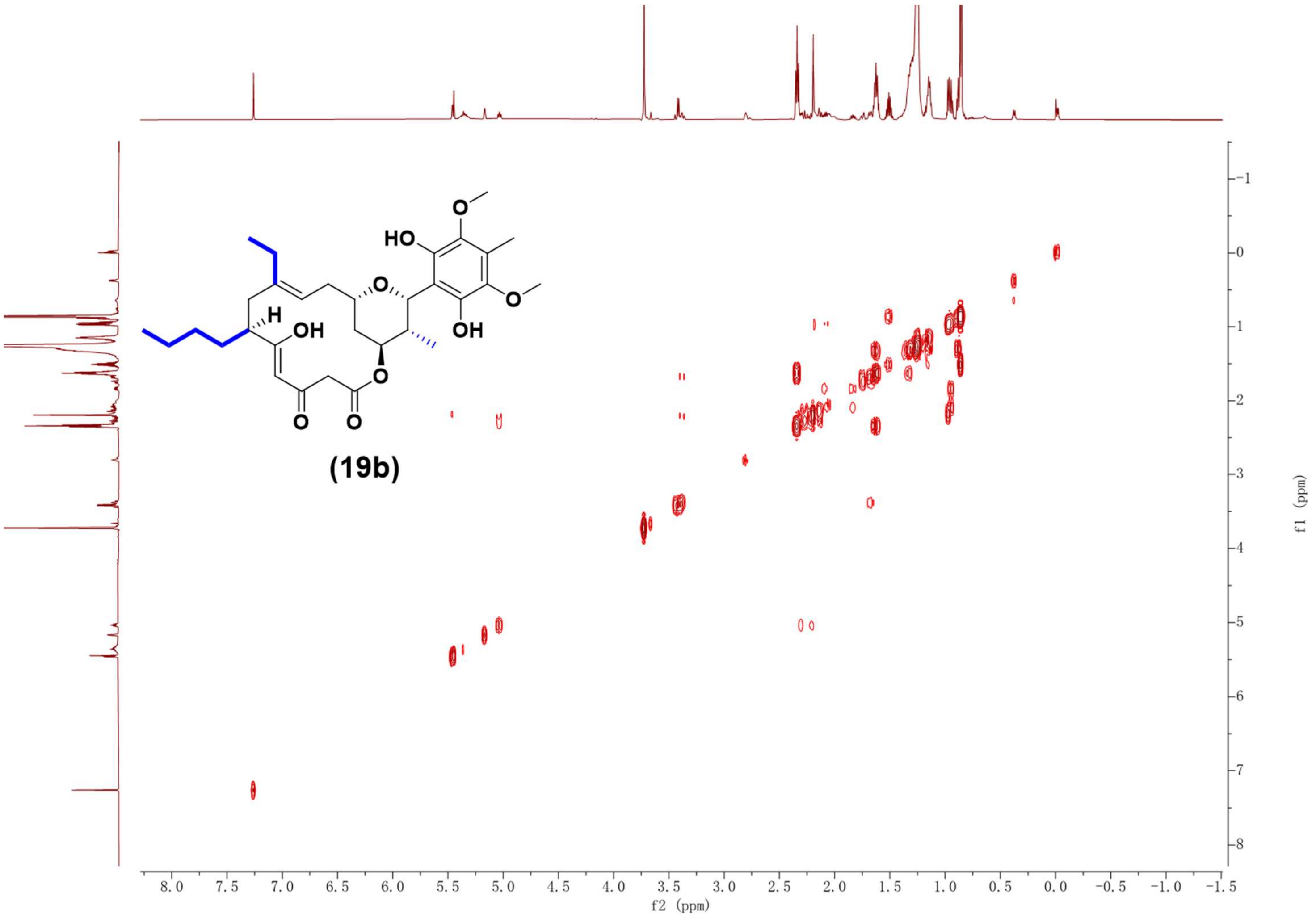

Supplementary Fig. 136. $^{1}H$-$^{1}H$ COSY NMR spectrum (600 MHz, $CDCl_3$) of compound **19b**.

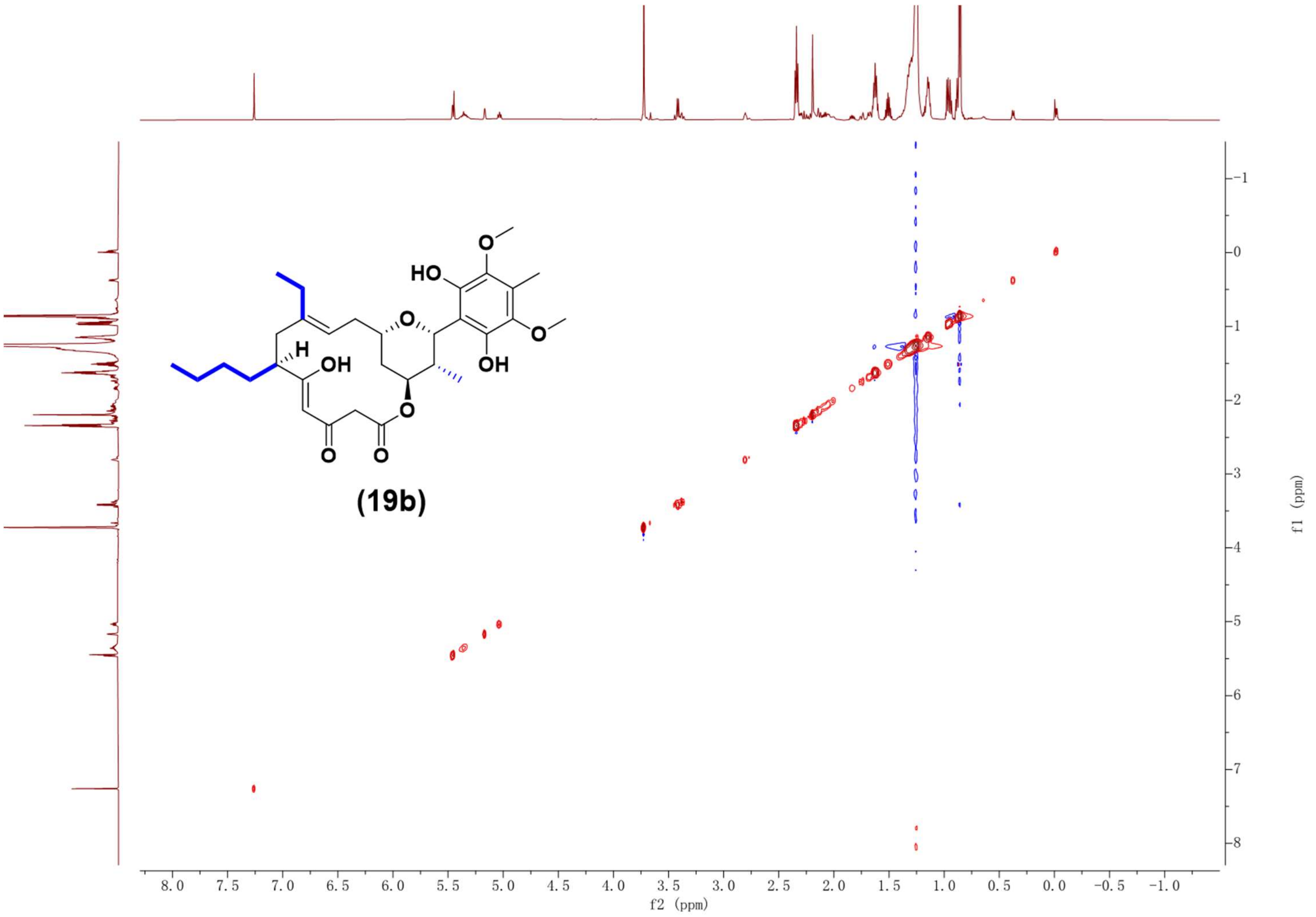

Supplementary Fig. 137. NOESY NMR spectrum (600 MHz, $CDCl_3$) of compound **19b**.

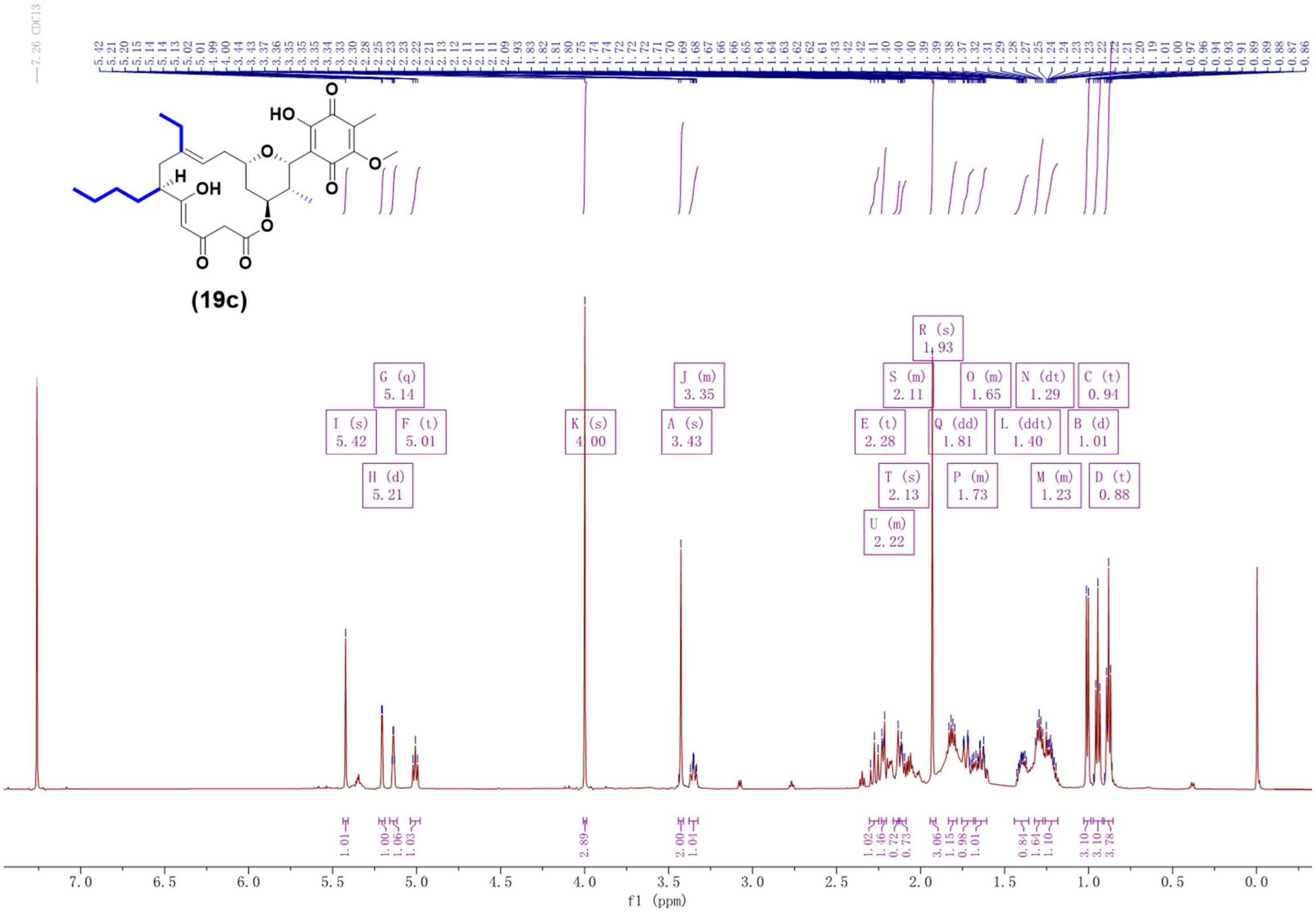

Supplementary Fig. 138. $^{1}H$ NMR spectrum (600 MHz, $CDCl_3$) of compound **19c**.

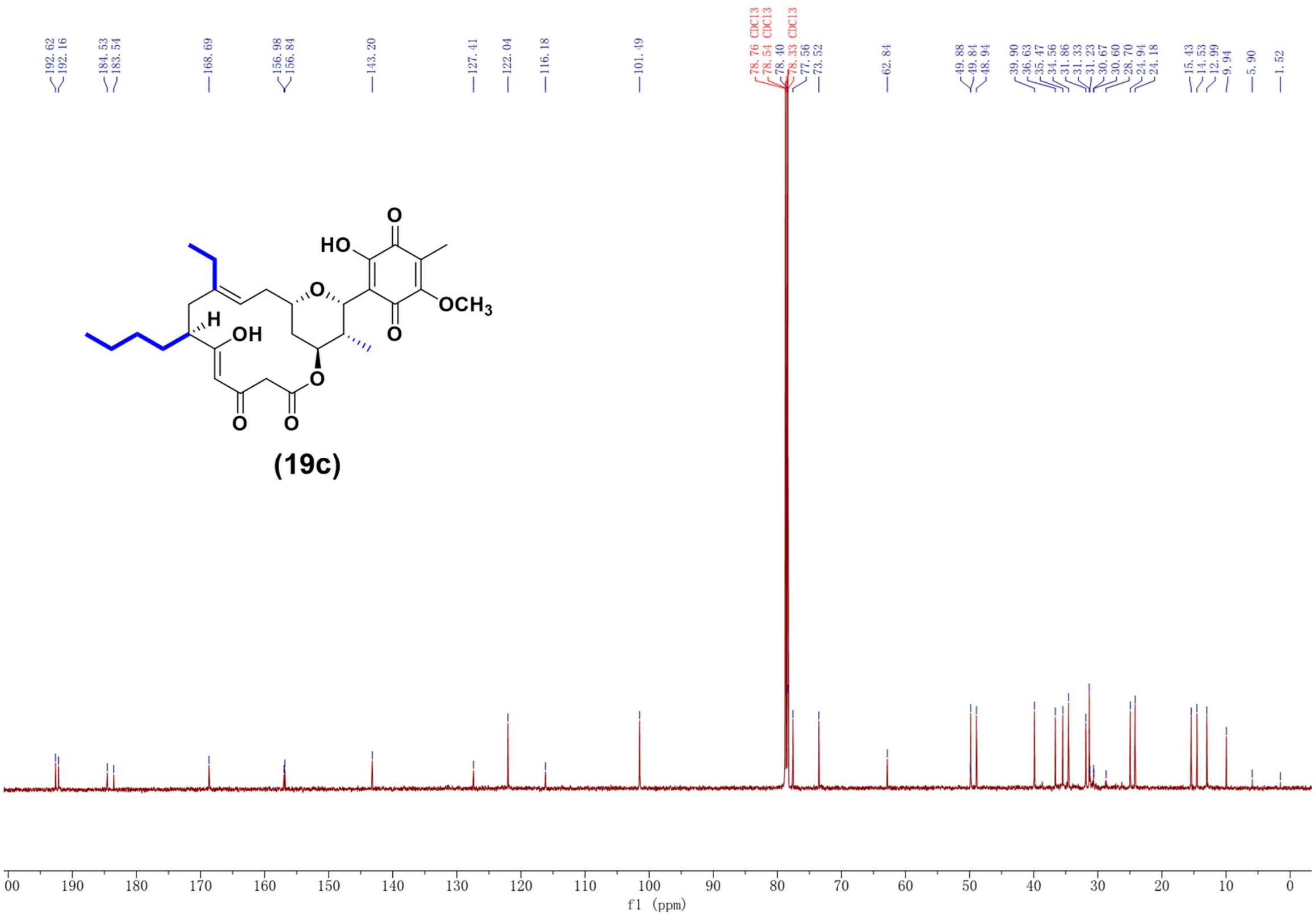

Supplementary Fig. 139. $^{13}C$ NMR spectrum (150 MHz, $CDCl_3$) of compound **19c**.

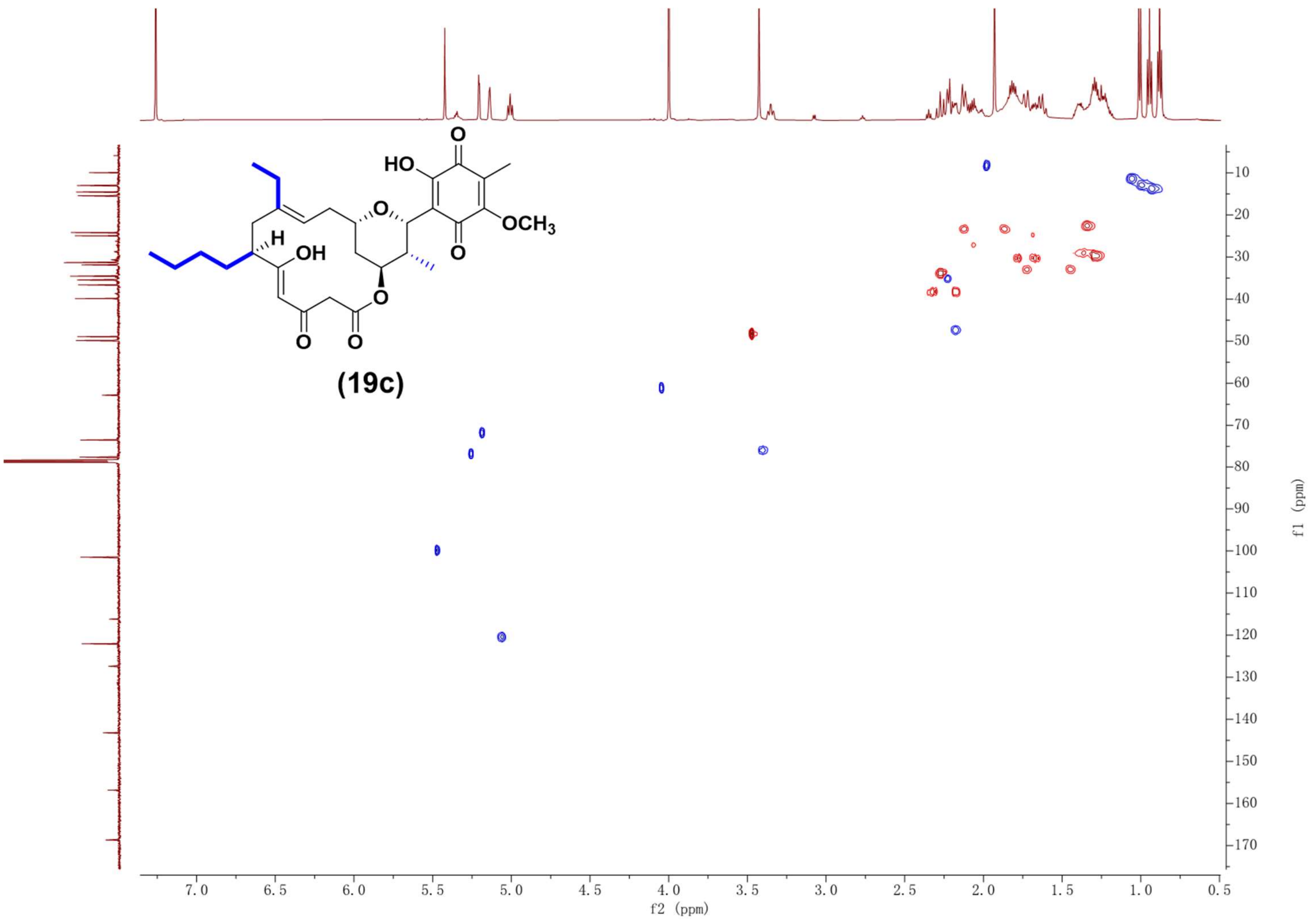

Supplementary Fig. 140. HSQC NMR spectrum (600 MHz, $CDCl_3$) of compound **19c**.

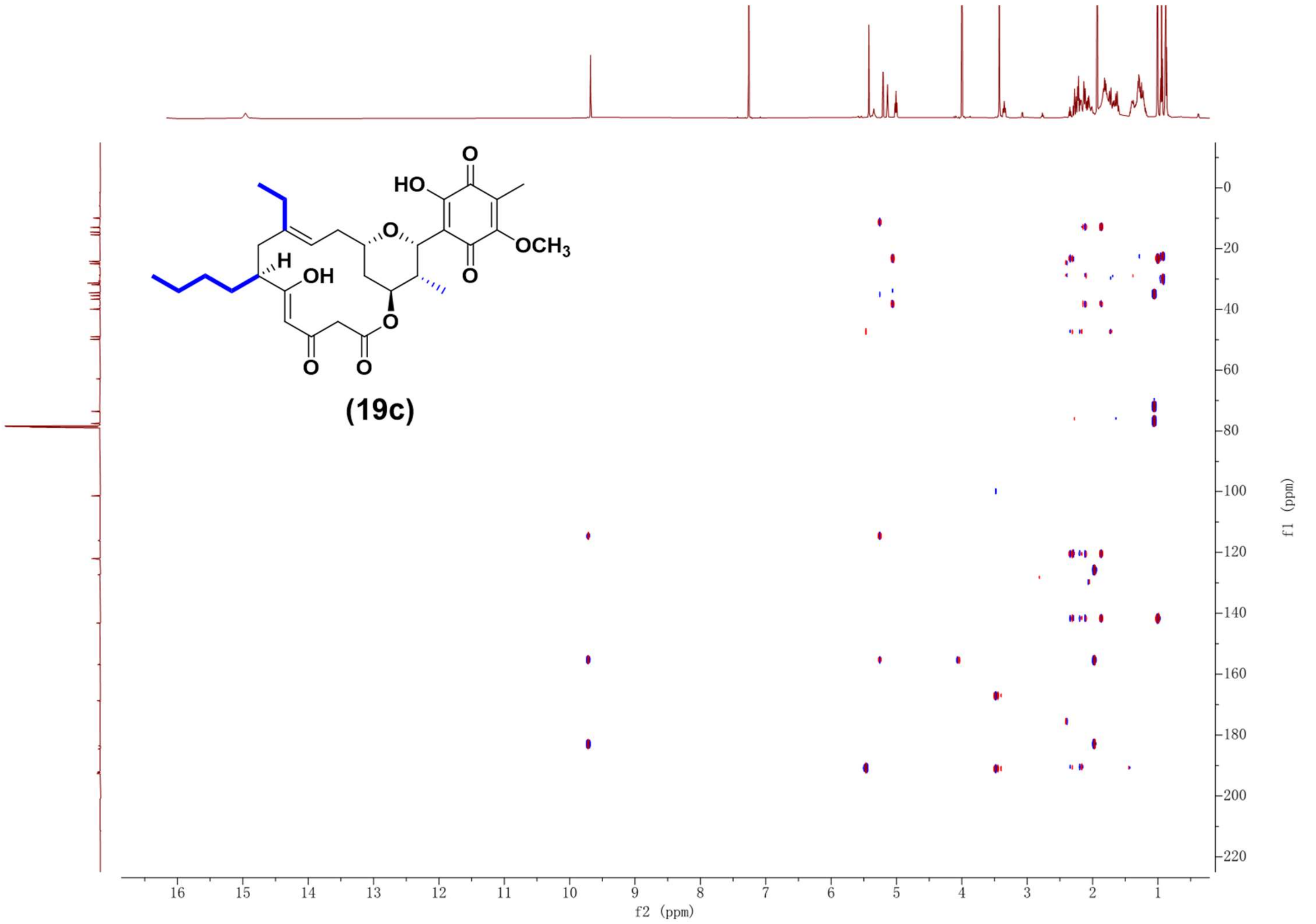

Supplementary Fig. 141. HMBC NMR spectrum (600 MHz, $CDCl_3$) of compound **19c**.

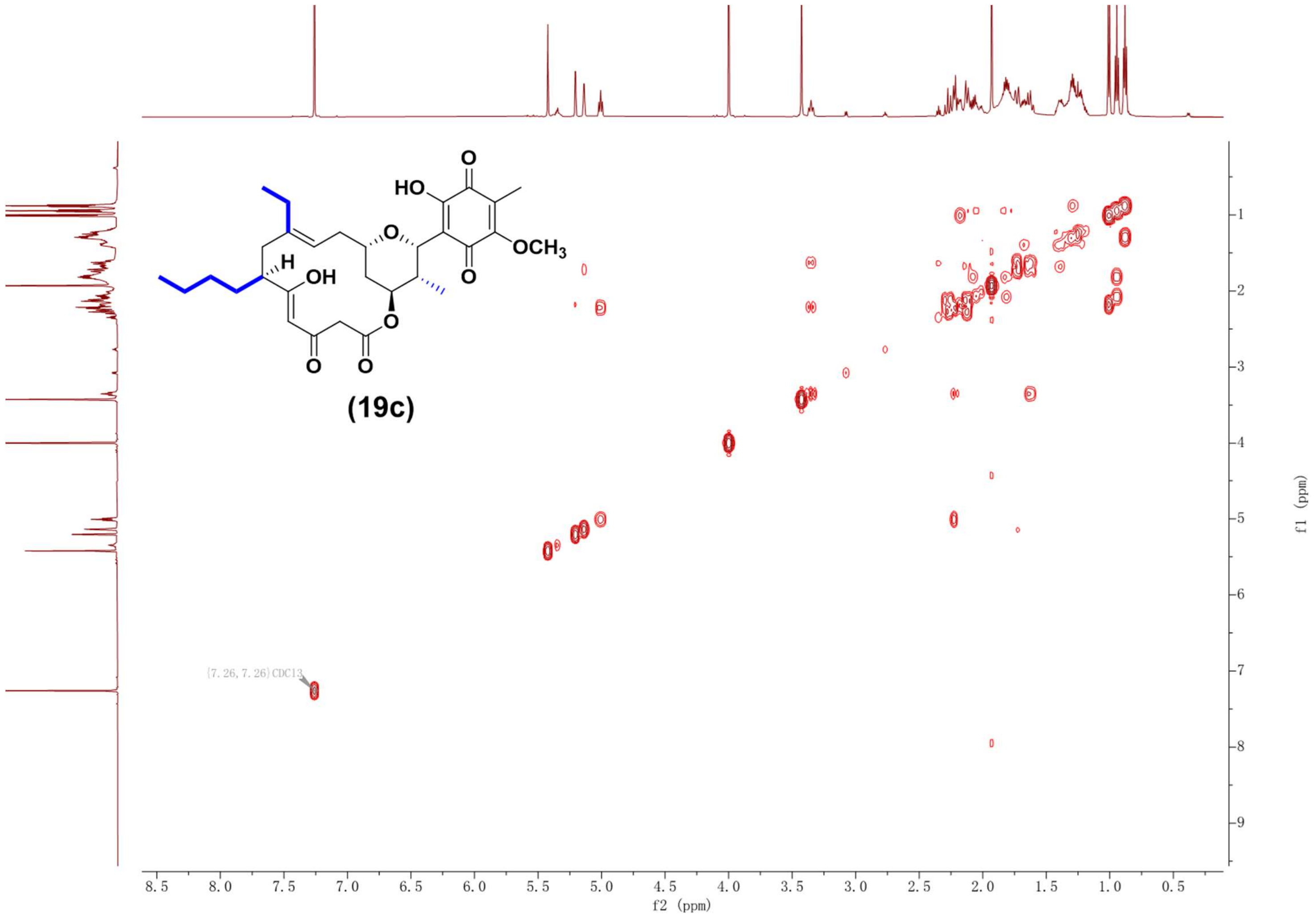

Supplementary Fig. 142. $^1H$-$^1H$ COSY NMR spectrum (600 MHz, $CDCl_3$) of compound **19c**.

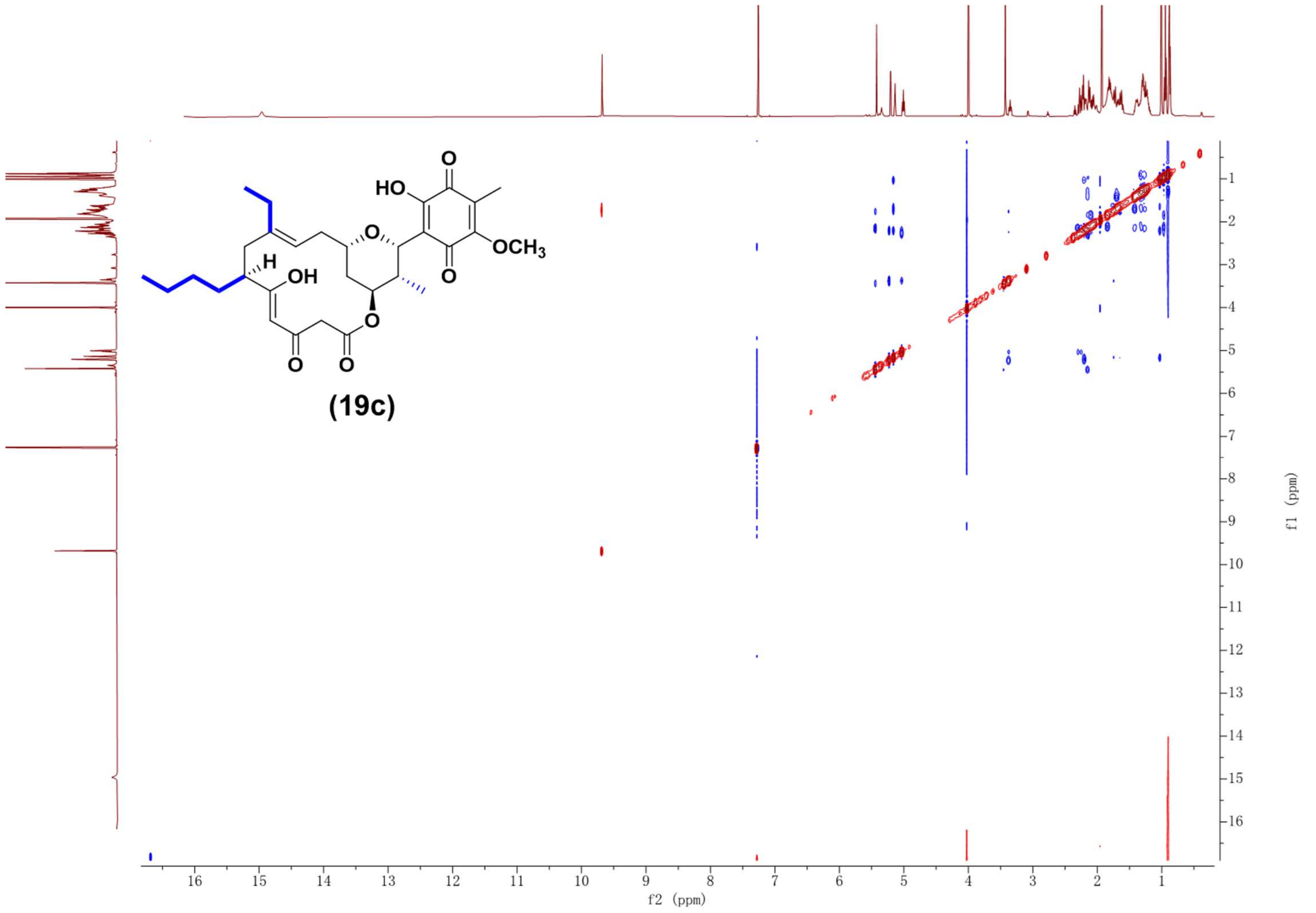

Supplementary Fig. 143. NOESY NMR spectrum (600 MHz, $CDCl_3$) of compound **19c**.

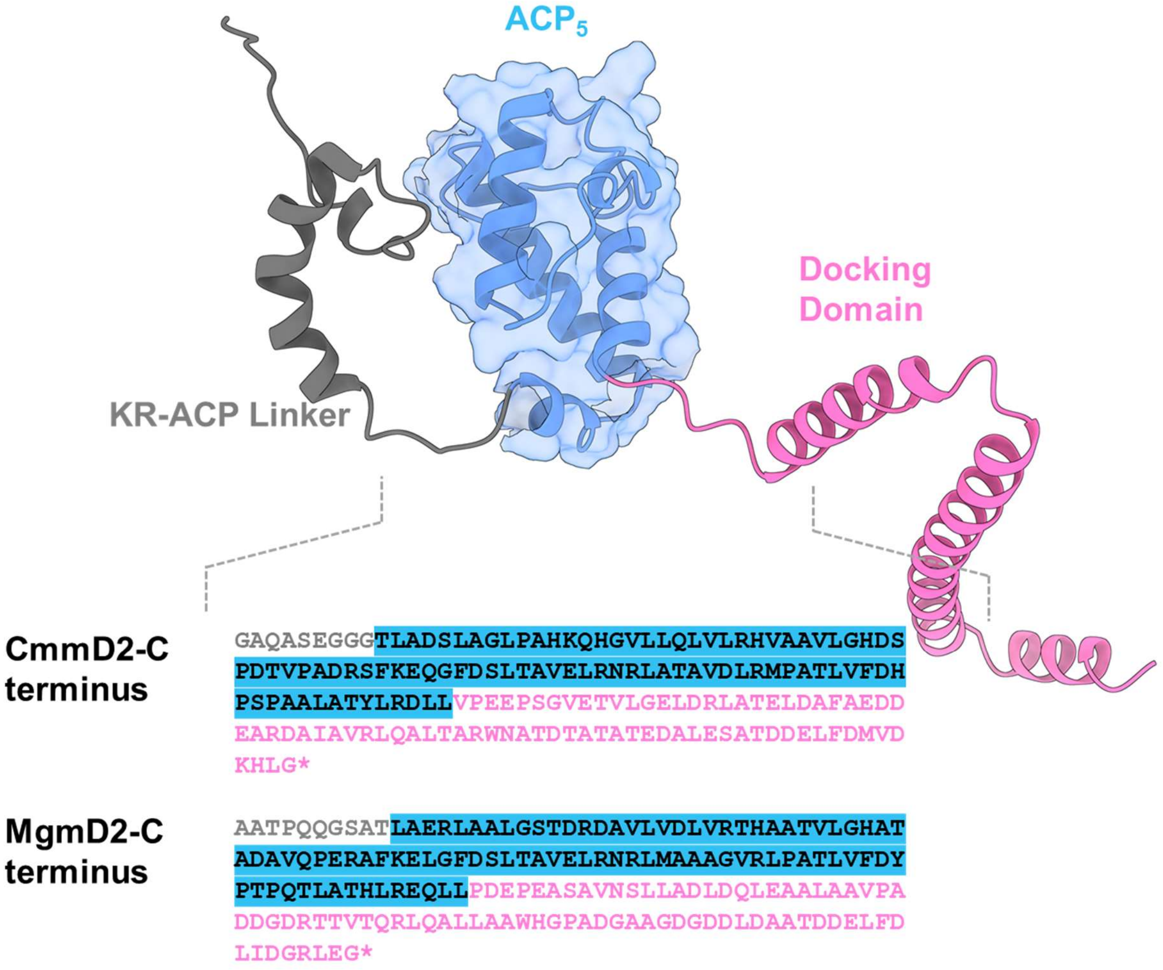

Supplementary Fig. 144. Structural modeling of CmmD2-$ACP_5$ region.

Model was performed using AlphaFold 2.0 and visualized using Chimera X software. CmmD2-$ACP_5$ domain and MgmD2-$ACP_5$ domain are highlighted in blue. C-terminal docking domains of CmmD2 and MgmD2 are highlighted in pink, and the upstream linkers of ACP domains are colored in grey.

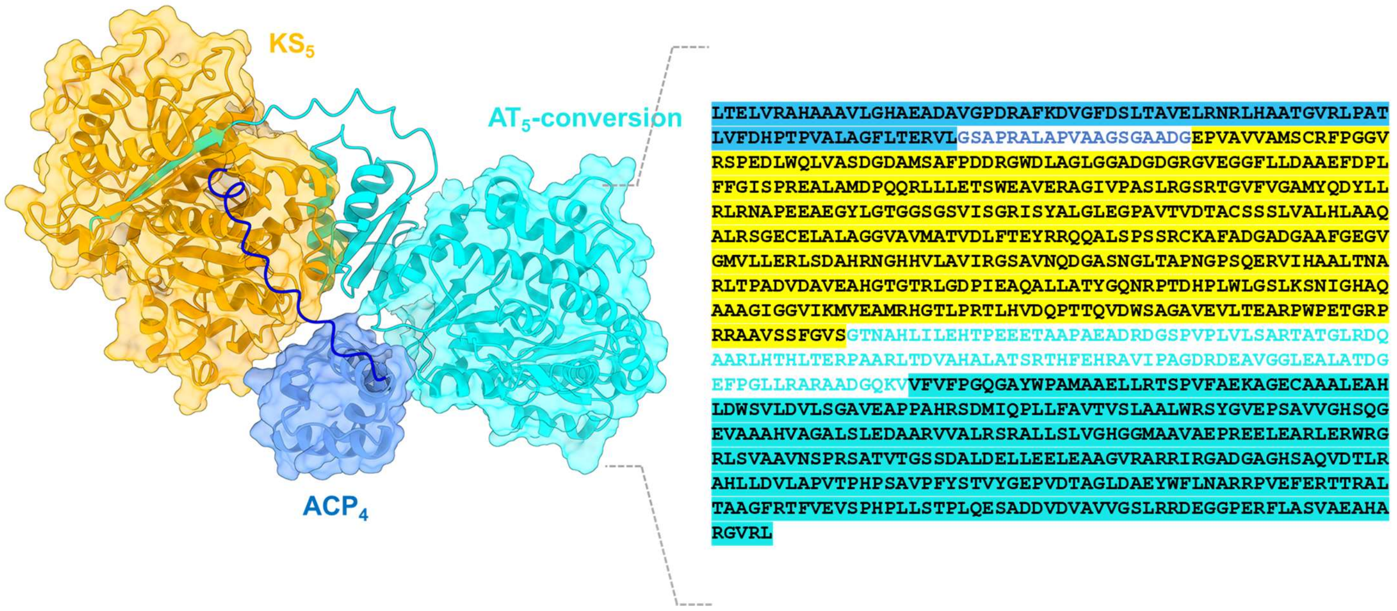

Supplementary Fig. 145. Structural modeling of CmmD2-($ACP_4$-$KS_5$-$AT_5$) subunit.

Model was performed using AlphaFold 2.0 and visualized using Chimera X software. The CmmD2-$KS_5$ domain is highlighted in yellow. CmmD2-$ACP_4$ domain and $ACP_4$-$KS_5$ linker are highlighted in blue, and the downstream CmmD2-$AT_5$ domain is highlighted in cyan.

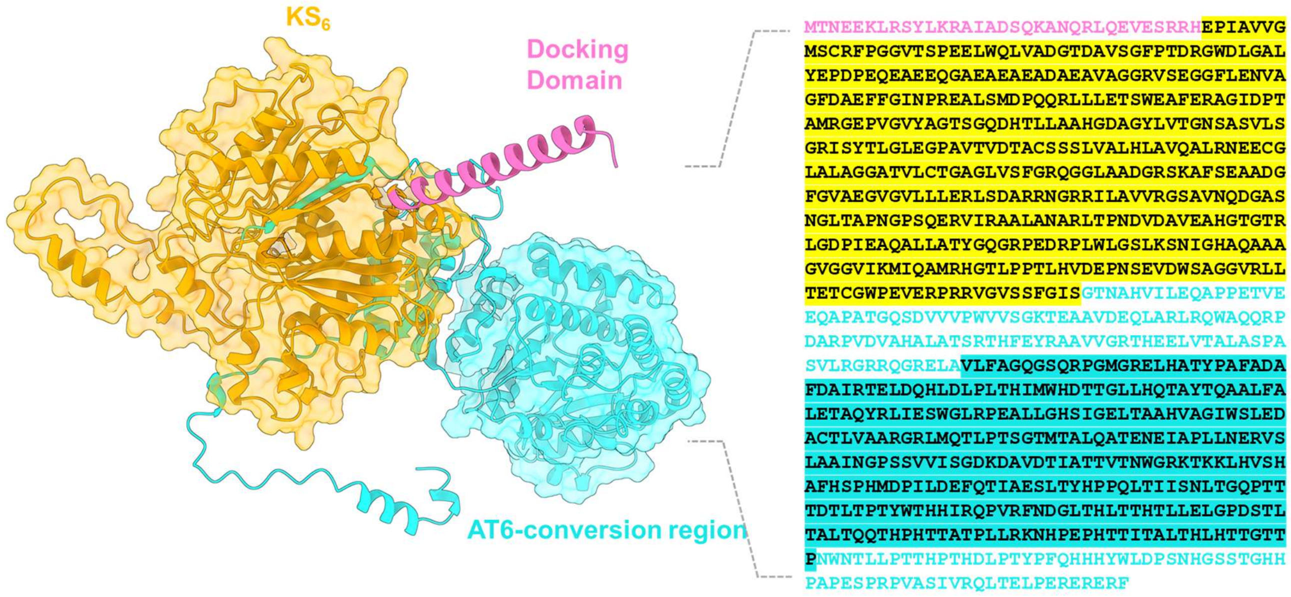

Supplementary Fig. 146. Structural modeling of CmmD3-(Docking domain-$KS_6$-$AT_6$) subunit.

Model was performed using alphafold 2.0 and visualized using Chimera X software. The CmmD3-$KS_6$ domain is highlighted in yellow, indicating the replacement boundaries of the KS domain for mutant S5-mgm$KS_6$. The N-terminal docking domain of CmmD3 is highlighted in pink, and the downstream $AT_6$ domain is highlighted in cyan.

### CmmD2-module 4/5

**a**

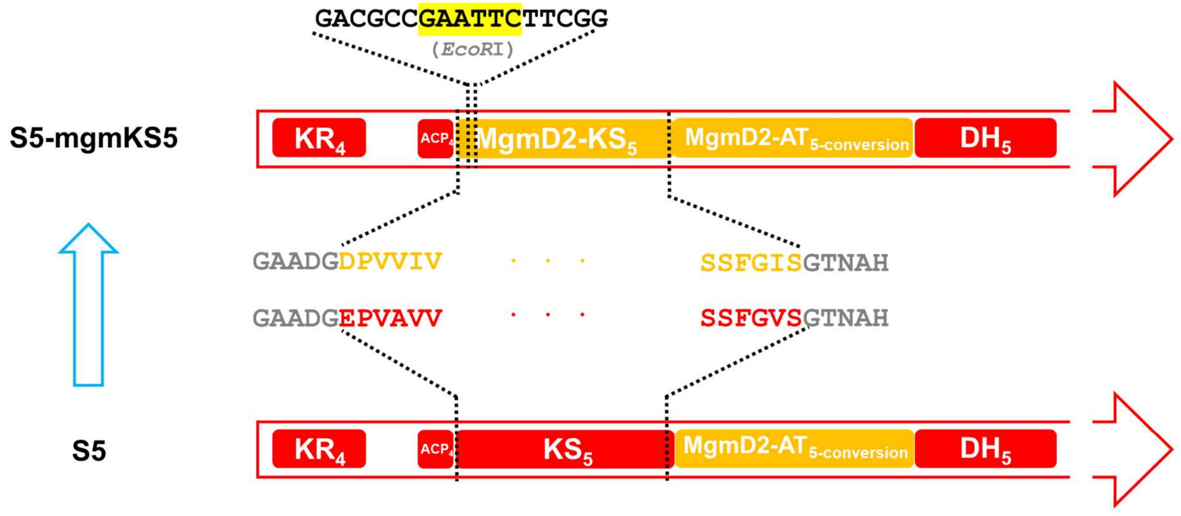

**b**

| Mutant | Primer Name | Primer Sequence (5'→3') | Screening Method |
|---|---|---|---|
| S5-mgmKS5 | sKS5-S1 | CTCTCCGACGAGCCCCAGAC | PCR products were digested using *EcoR*I |
| | sKS5-S2 | ACCAGTCCAGGAGTGGATCG | |

**c**

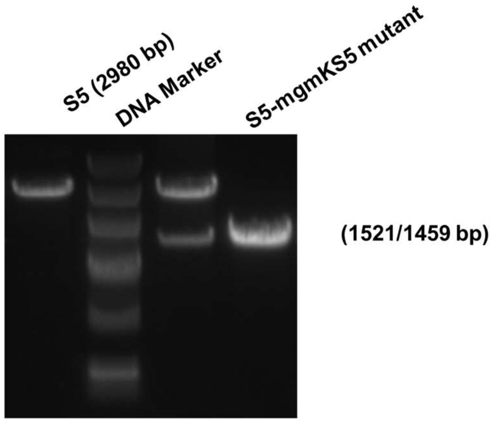

Supplementary Fig. 147. Construction and confirmation of mutant S5-mgmKS5 strain.

(**a**): Schematic diagram of construction of mutant S5-mgmKS5. *EcoR*I restriction sites used for screening is highlighted in yellow; (**b**) Primers and methods for mutant strain screening; (**c**) Agarose gel screening of mutant S5-mgmKS5 by PCR and restriction enzyme digesting. The red squares represent domains from CmmD2.

a

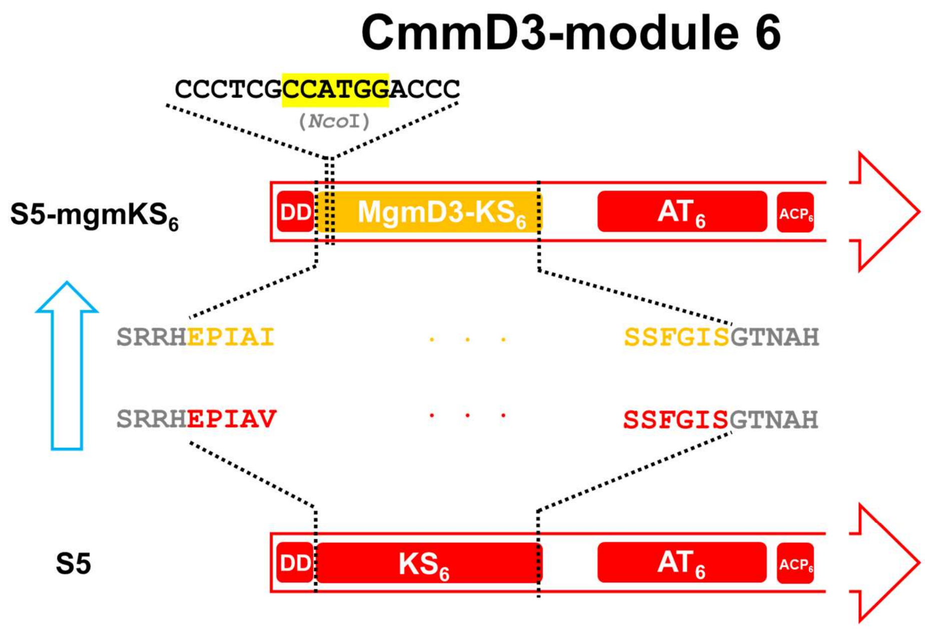

b

| Mutant | Primer Name | Primer Sequence (5'→3') | Screening Method |
|---|---|---|---|
| S5-mgmKS$_6$ | sKS6-S1 | AACGCGACGGAC ACGGCTAC | PCR products were digested using *Nco*I |
| | sKS6-S2 | TGGCGACGGGAC GAGGAGAC | |

c

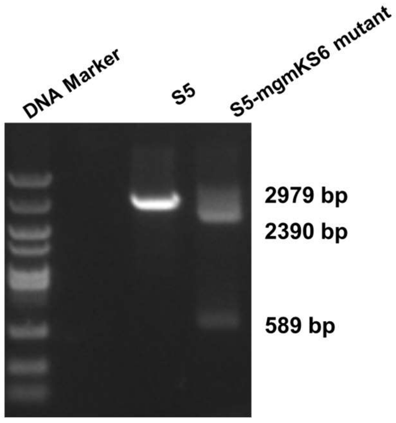

Supplementary Fig. 148. Construction and confirmation of mutant S5-mgmKS$_6$ strain.

(**a**): Schematic diagram of construction of mutant S5-mgmKS$_6$, *Nco*I restriction sites used for screening is highlighted in yellow; (**b**) Primers and methods for mutant strain screening; (**c**) Agarose gel screening of mutant S5-mgmKS$_6$ by PCR and restriction enzyme digesting. The red squares represent domains from CmmD3, and DD is the abbreviation of docking domain.

**a**

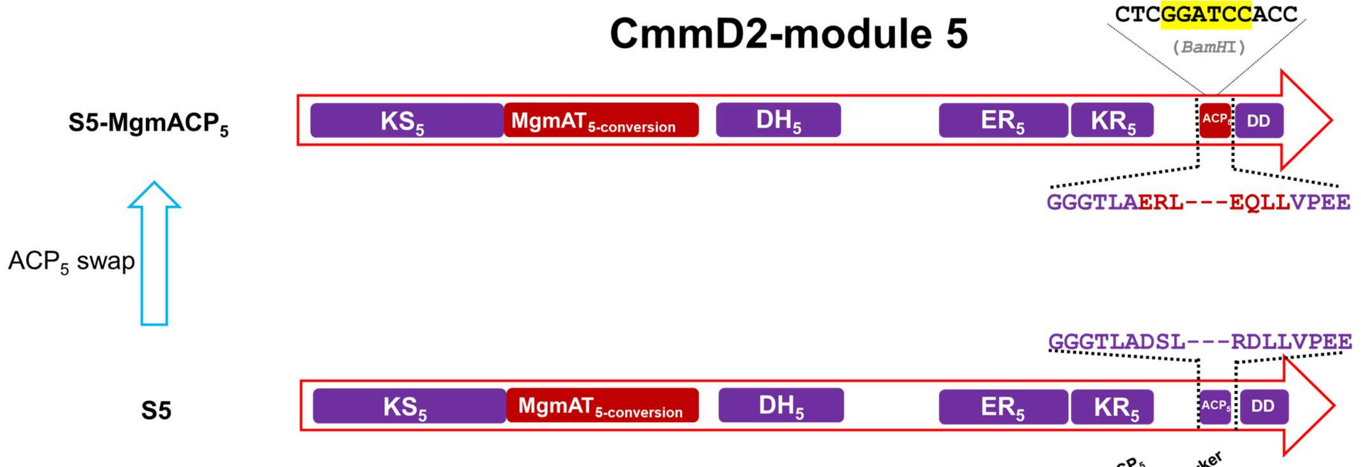

**b**

| Mutant | Primer Name | Primer Sequence (5'→3') | Method |
|---|---|---|---|
| **S5-mgmACP5** | ACP5-S1 | GTCGGCAAGATGG TCCTCAC | PCR products were digested using *BamH*I |
| | ACP5-S2 | GGTTGGCCTTTTG GGAGTCG | |

**c**

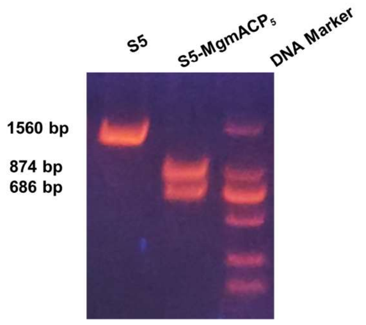

Supplementary Fig. 149. Construction and confirmation of mutant S5-mgmACP5 strain.

(**a**): Schematic diagram of construction of mutant S5-mgmACP5, *Bam*HI restriction sites used for screening is highlighted in yellow; (**b**) Primers and methods for mutant strain screening; (**c**) Agarose gel screening of mutant S5-mgmACP5 by PCR and restriction enzyme digesting. The purple squares represent domains from CmmD2, and DD is the abbreviation of docking domain.

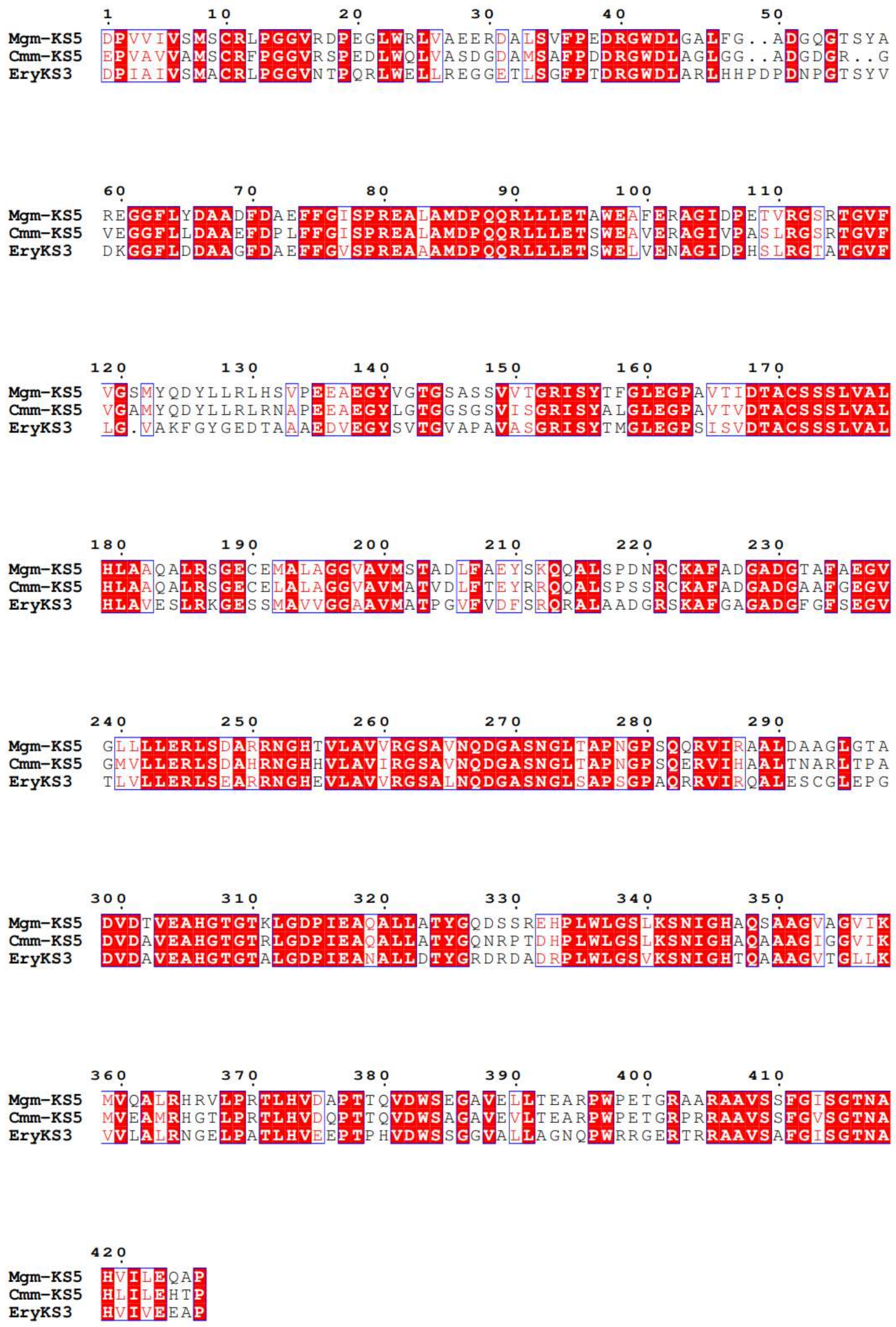

Supplementary Fig. 150. Amino acid sequence alignments of Mgm-KS5, Cmm-KS5 and EryKS3[6].

**a**

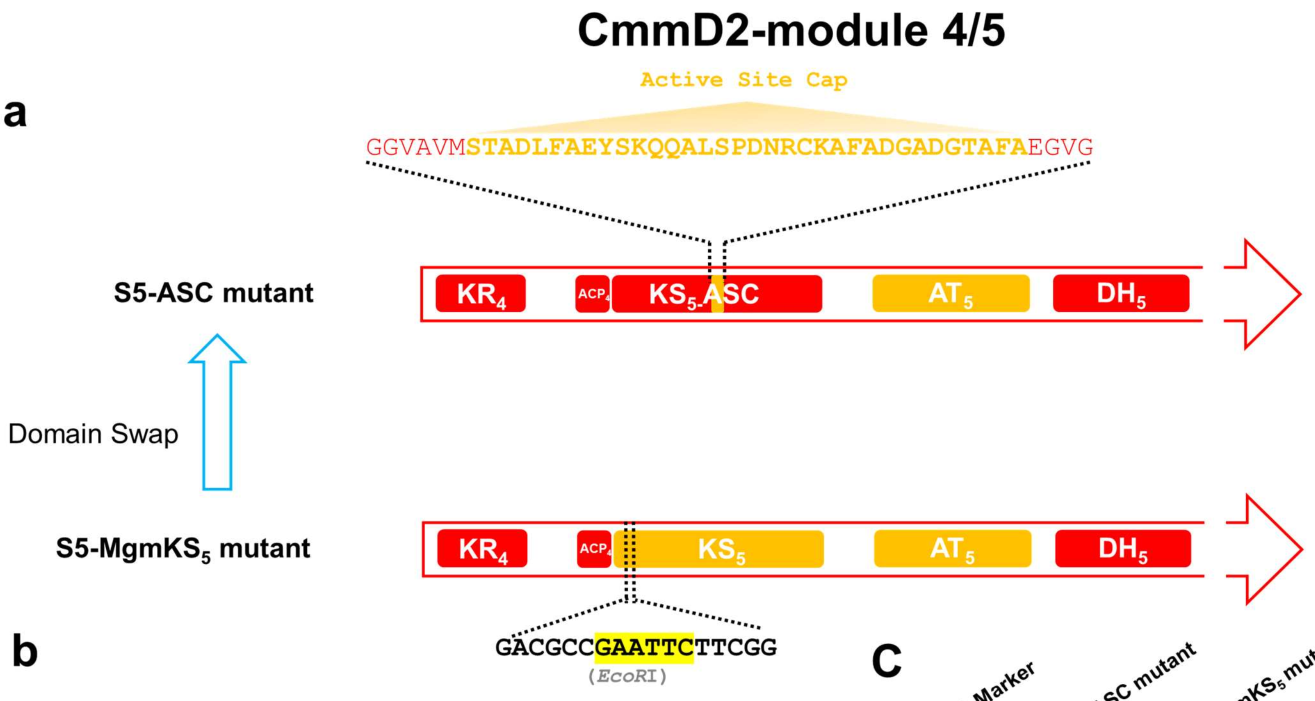

**b**

| Mutant | Primer Name | Primer Sequence (5'→3') | Template |
|---|---|---|---|
| S5-ASC | sKS5-S1 | CTCTCCGACGAG CCCCAGAC | PCR products were digested using *EcoR*I |
| | sKS5-S1 | ACCAGTCCAGGA GTGGATCG | |

**C**

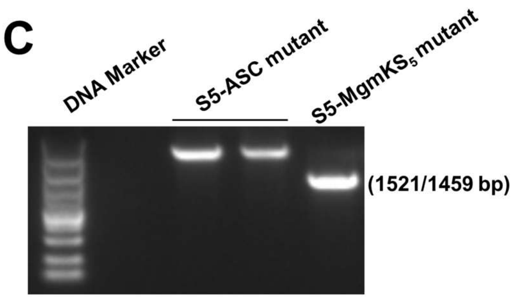

Supplementary Fig. 151. Construction and confirmation of mutant S5-ASC.

(**a**): Schematic diagram of construction of mutant S5-mgmKS5. *EcoR*I restriction sites used for screening is highlighted in yellow; (**b**) Primers and methods for mutant strain screening; (**c**) Agarose gel screening of mutant S5-ASC by PCR and restriction enzyme digesting. The red squares represent domains from CmmD2, and yellow squares represent domains from MgmD2.

a

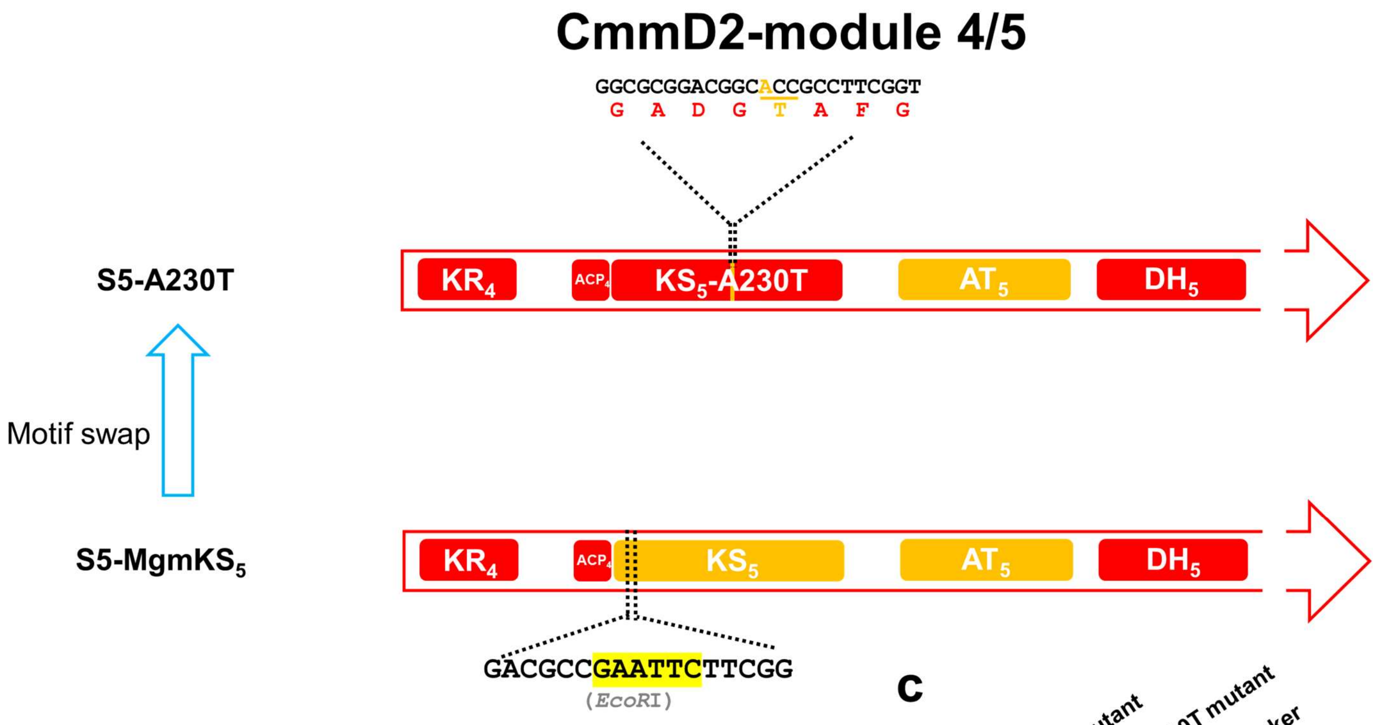

b

| Mutant | Primer Name | Primer Sequence (5'→3') | Template |
|---|---|---|---|
| S5-A230T | sKS5-S1 | CTCTCCGACGAGCCCCAGAC | PCR products were digested using *Eco*RI |
| | sKS5-S1 | ACCAGTCCAGGAGTGGATCG | |

c

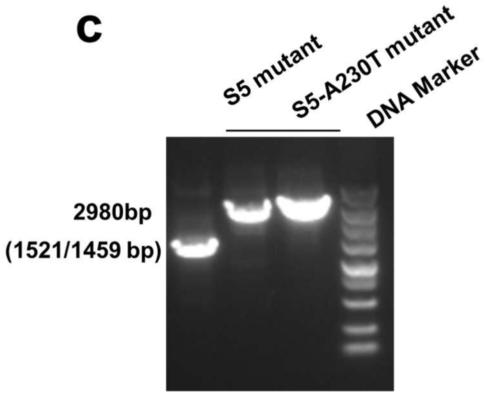

Supplementary Fig. 152. Construction and confirmation of S5-A230T mutant strain.

(**a**): Schematic diagram of construction of mutant S5-A230T. *EcoR*I restriction sites used for screening is highlighted in yellow; (**b**) Primers and methods for mutant strain screening; (**c**) Agarose gel screening of mutant S5-A230T by PCR and restriction enzyme digesting. The red squares represent domains from CmmD2, and yellow squares represent domains from MgmD2.

**a**

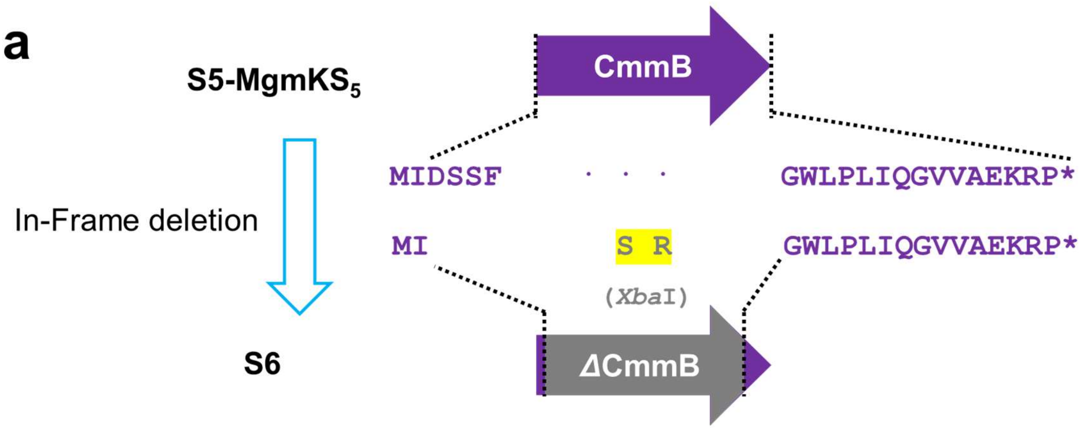

**b**

| Mutant | Primer Name | Primer Sequence (5'→3') | Screening Method |
|---|---|---|---|
| S6 | cmmB-S1 | AGTGCTGCACAGCTGGGACGAGTTC | PCR |
| | cmmB-S2 | TTCGCGGTTTTCATGACGTCACCTGTTCTC | |

**c**

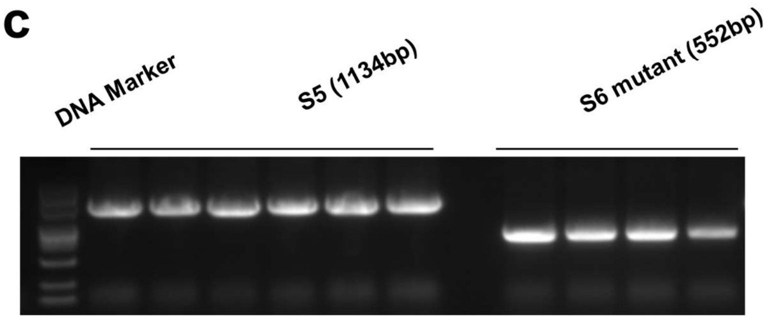

Supplementary Fig. 153. Construction and confirmation of mutant S6 strain.

(**a**): Schematic diagram of construction of mutant S6, *Xba***I** restriction sites is highlighted in yellow; (**b**) Primers and methods for mutant strain screening; (**c**) Agarose gel screening of mutant S6.

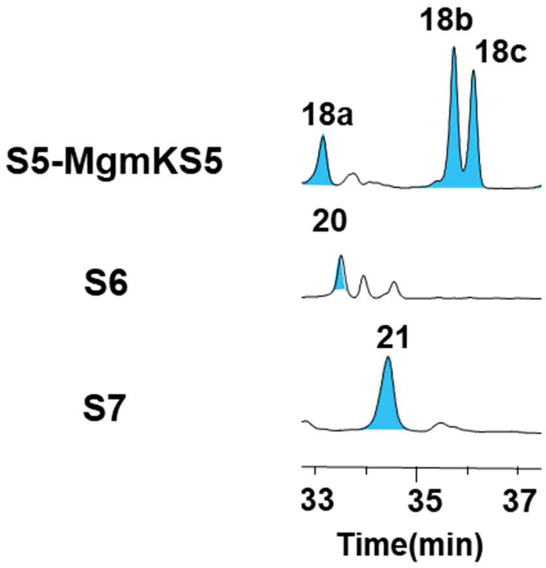

Supplementary Fig. 154. HPLC chromatograms of the fermentation broth from engineered strains of S6 and S7 at λ = 280 nm.

The desired products are shown in blue peaks. Three independent experiments were repeated.

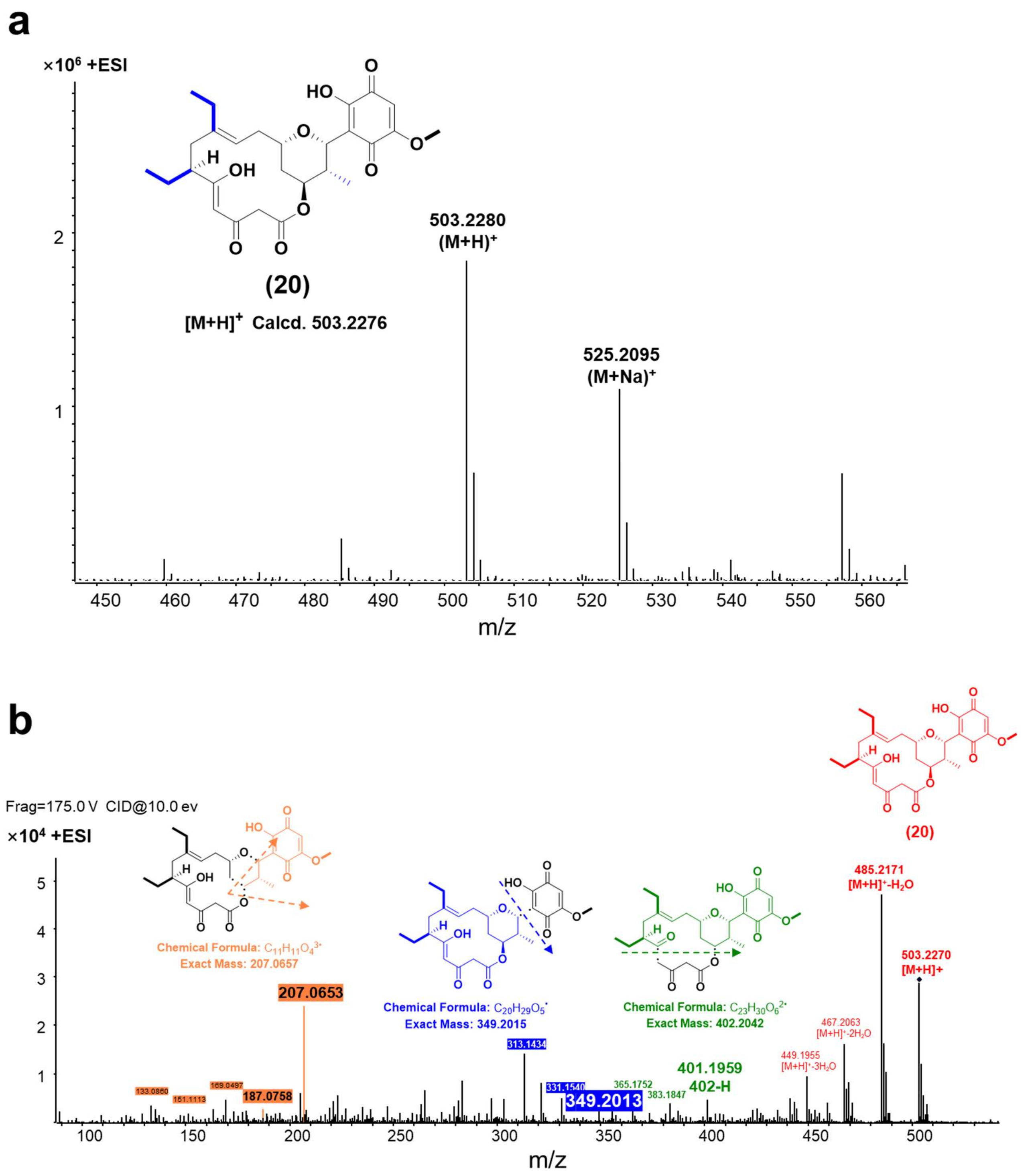

Supplementary Fig. 155. LC-ESI-HRMS (**a**)and MS/MS (**b**) analysis of compound **20**.

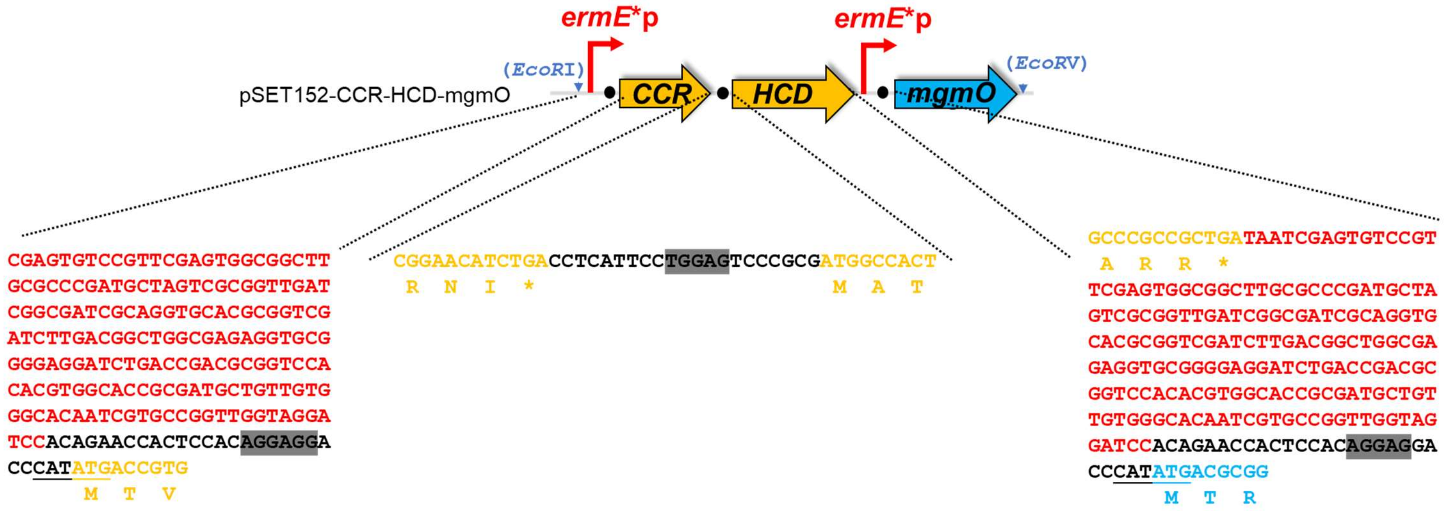

Supplementary Fig. 156. Nucleotide sequence of plasmid pSET1252-CCR-HCD-mgmO.

The "*ermE**p" nucleotide sequence is displayed in red bold font. The nucleotide sequences of CCR and HCD are displayed in yellow bold font. The nucleotide sequences of *mgmO* are displayed in blue bold font.

**a**

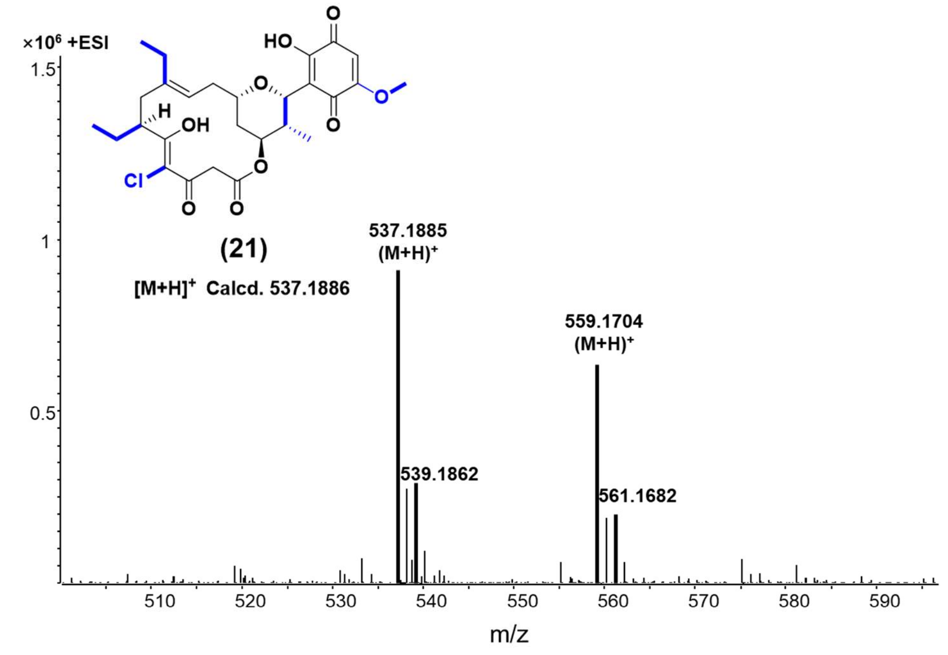

**b**

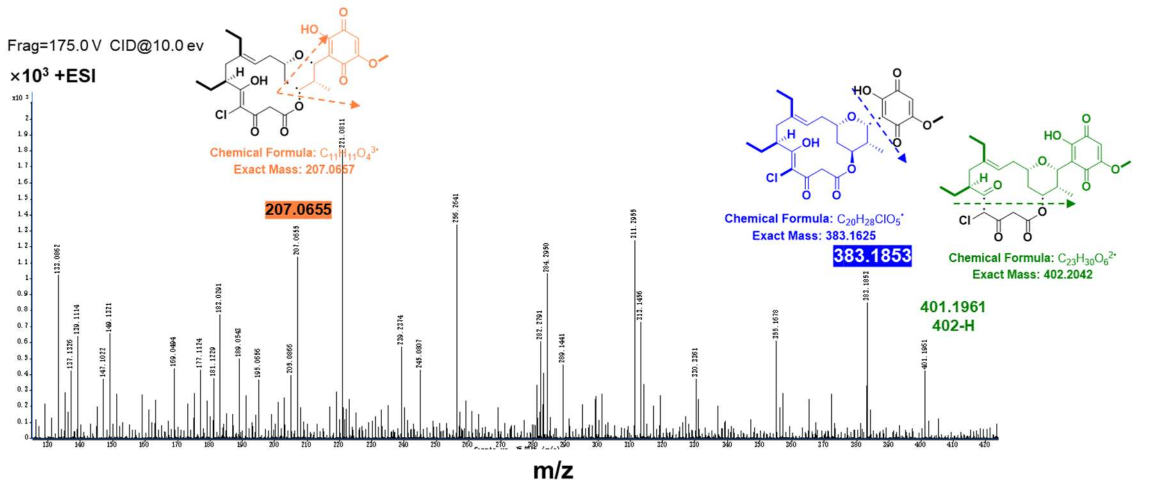

Supplementary Fig. 157. LC-ESI-HRMS spectrum (**a**) and MS/MS analysis (**b**) of compound **21**.

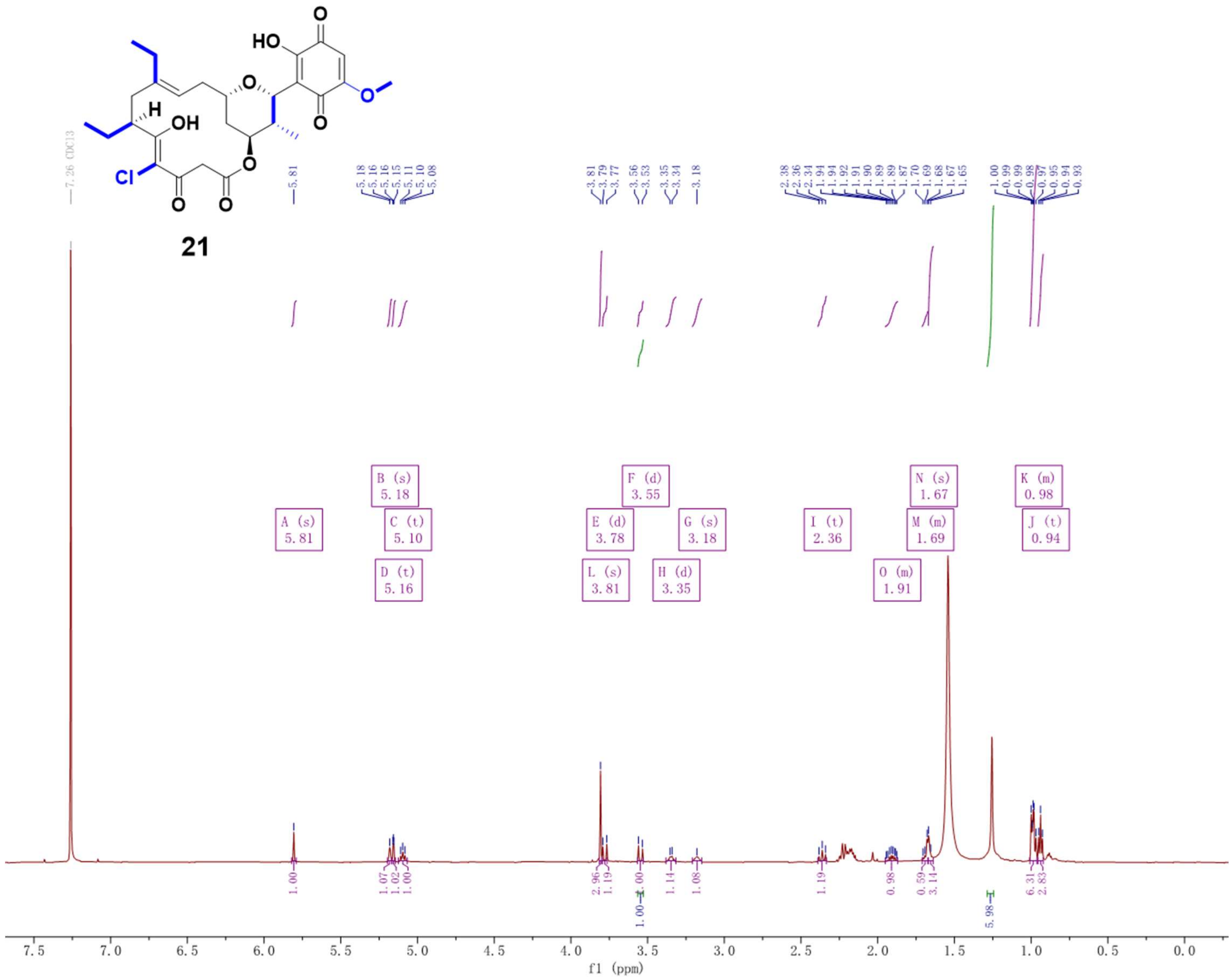

Supplementary Fig. 158. $^{1}H$ NMR spectrum (600 MHz, $CDCl_3$) of compound **21**.

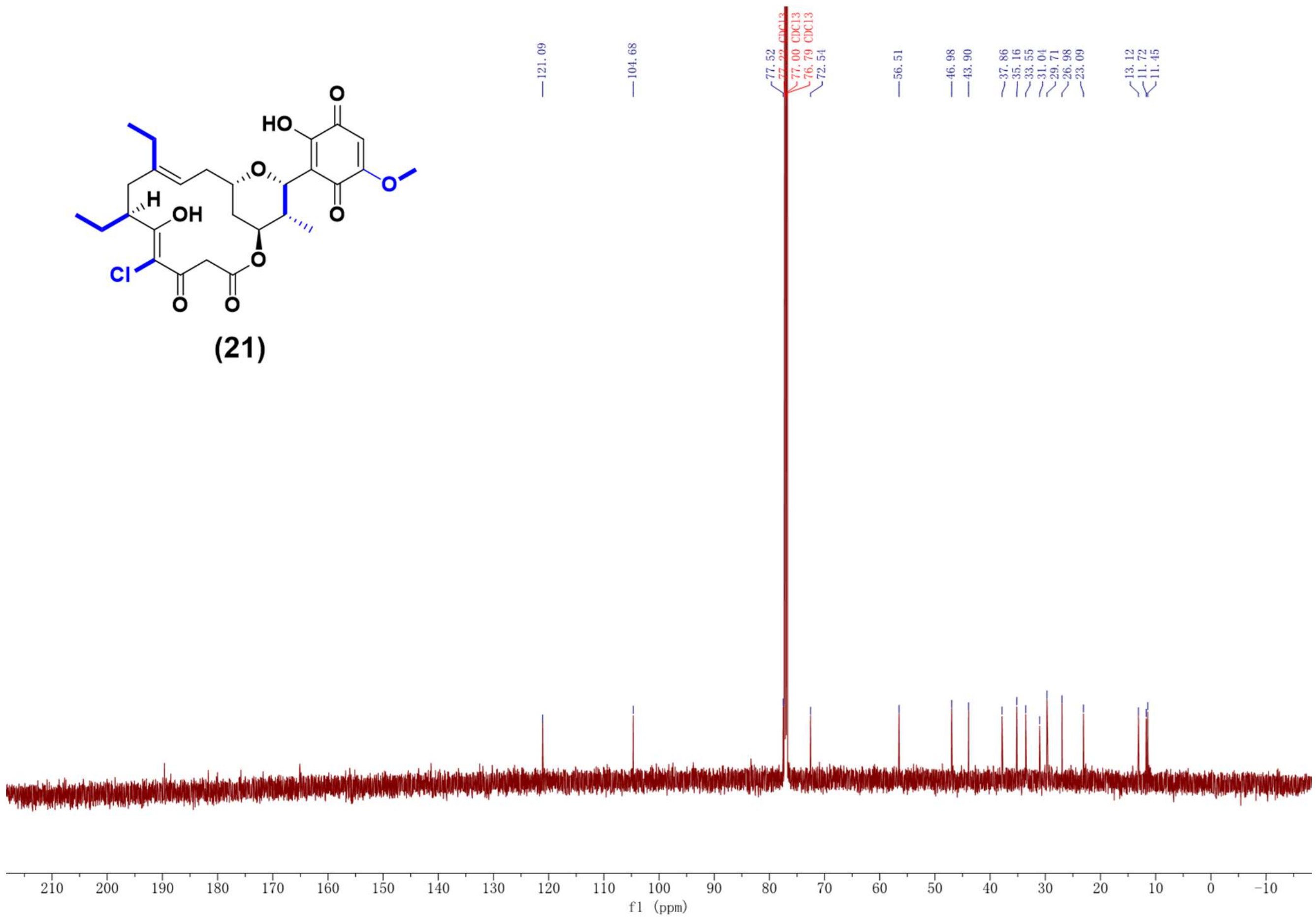

Supplementary Fig. 159. $^{13}C$ NMR spectrum (150 MHz, $CDCl_3$) of compound **21**.

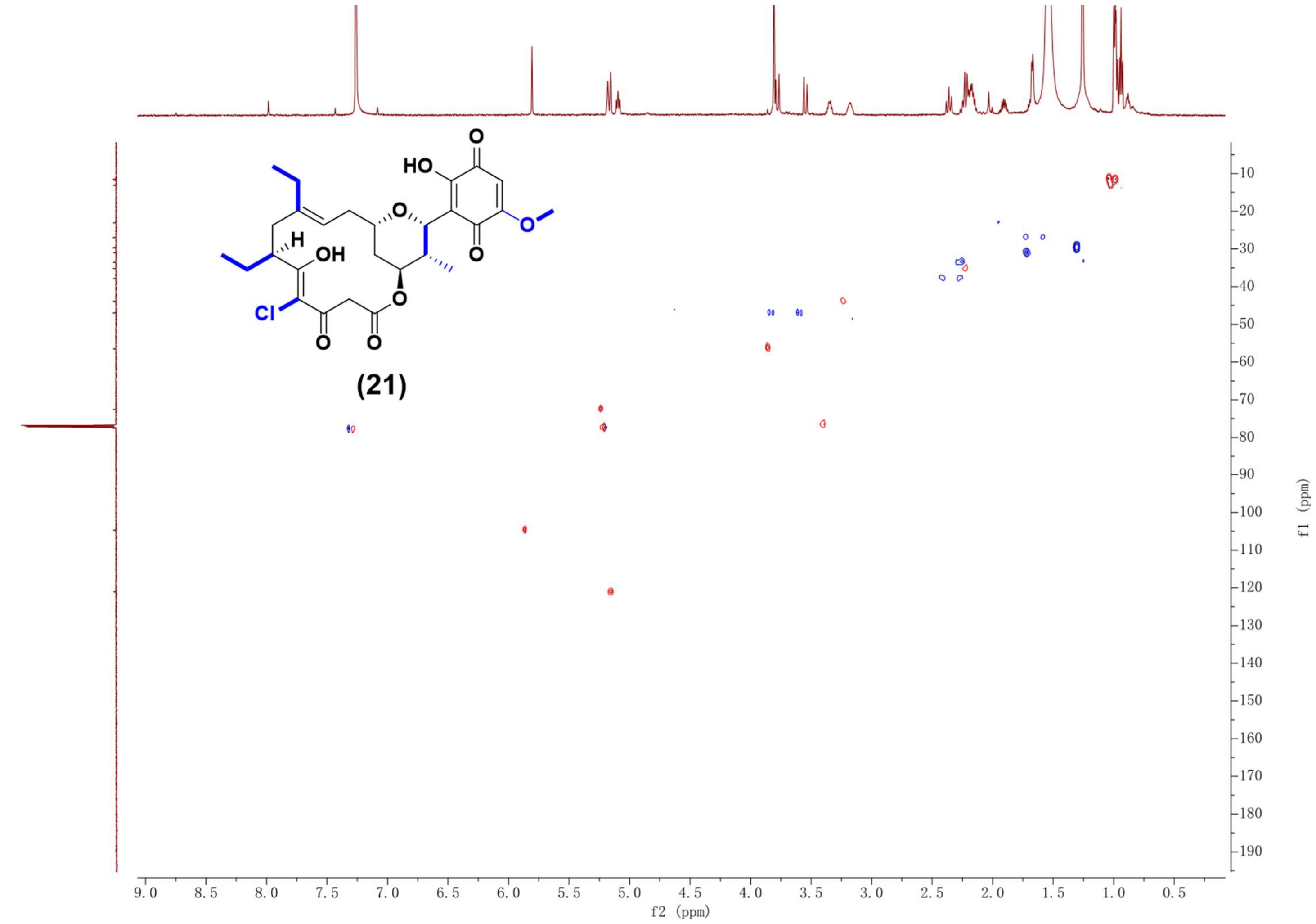

Supplementary Fig. 160. HSQC NMR spectrum (600 MHz, $CDCl_3$) of compound **21**.

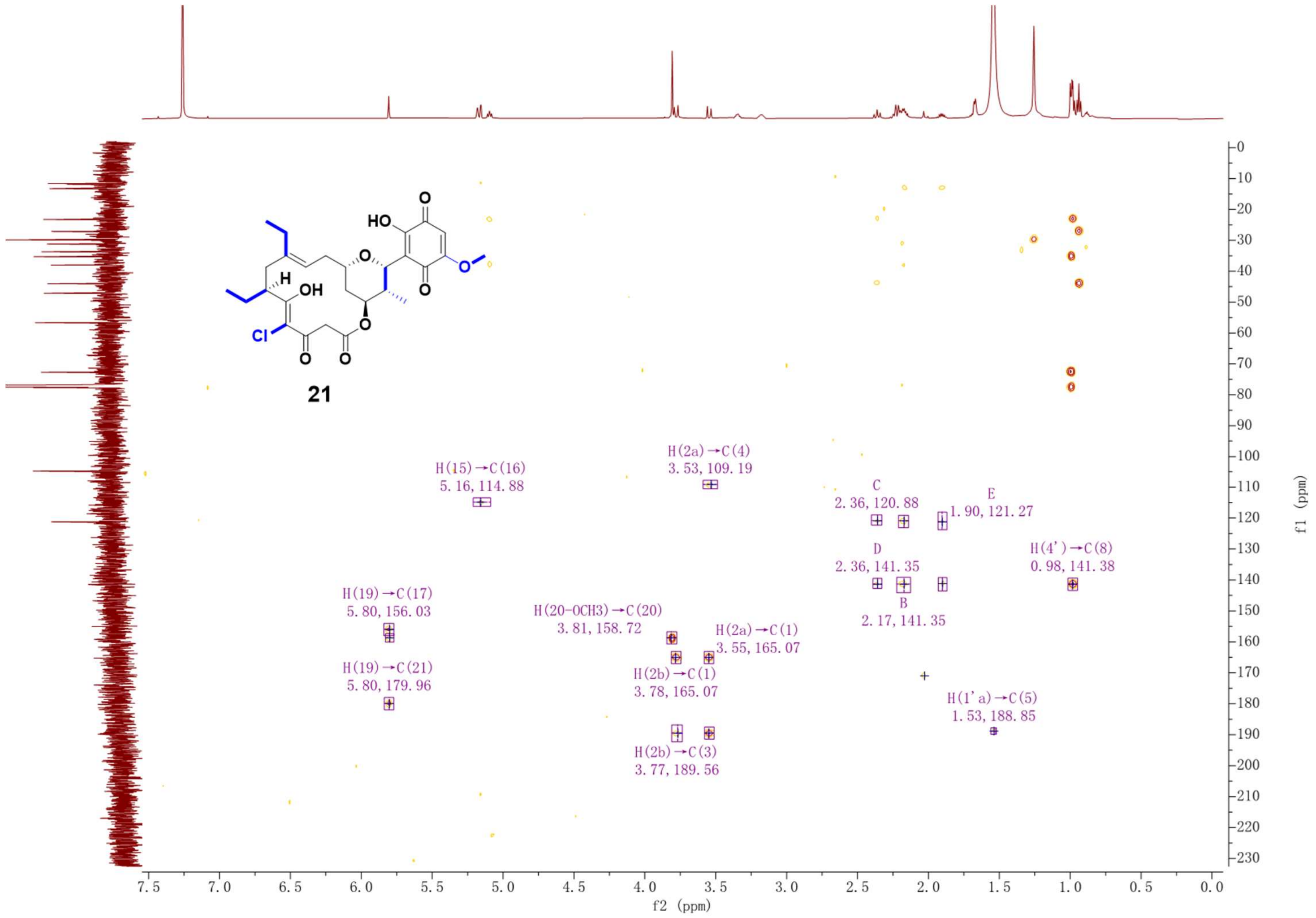

Supplementary Fig. 161. HMBC NMR spectrum (600 MHz, $CDCl_3$) of compound **21**.

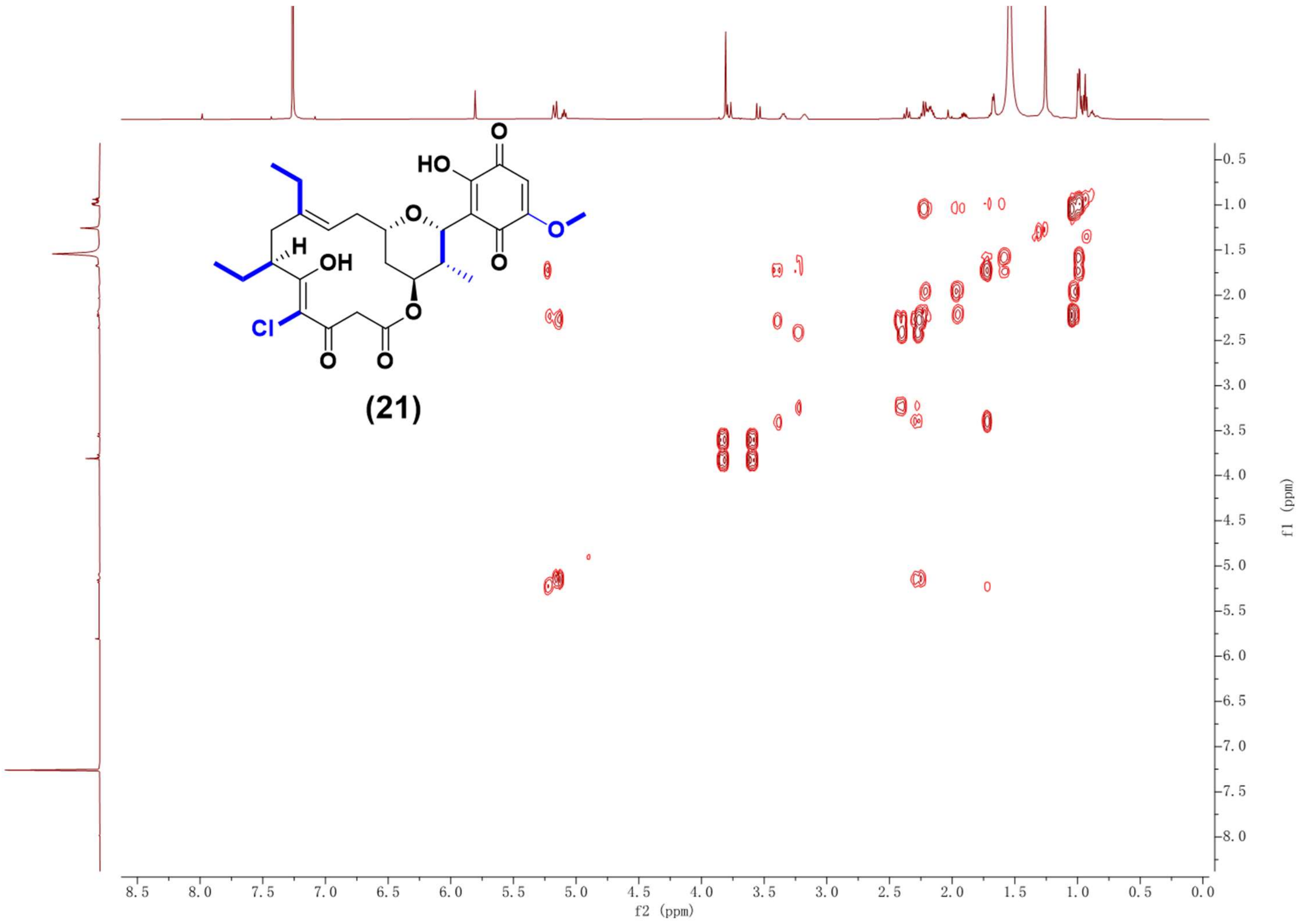

Supplementary Fig. 162. $^1$H-$^1$H COSY NMR spectrum (600 MHz, $CDCl_3$) of compound **21**.

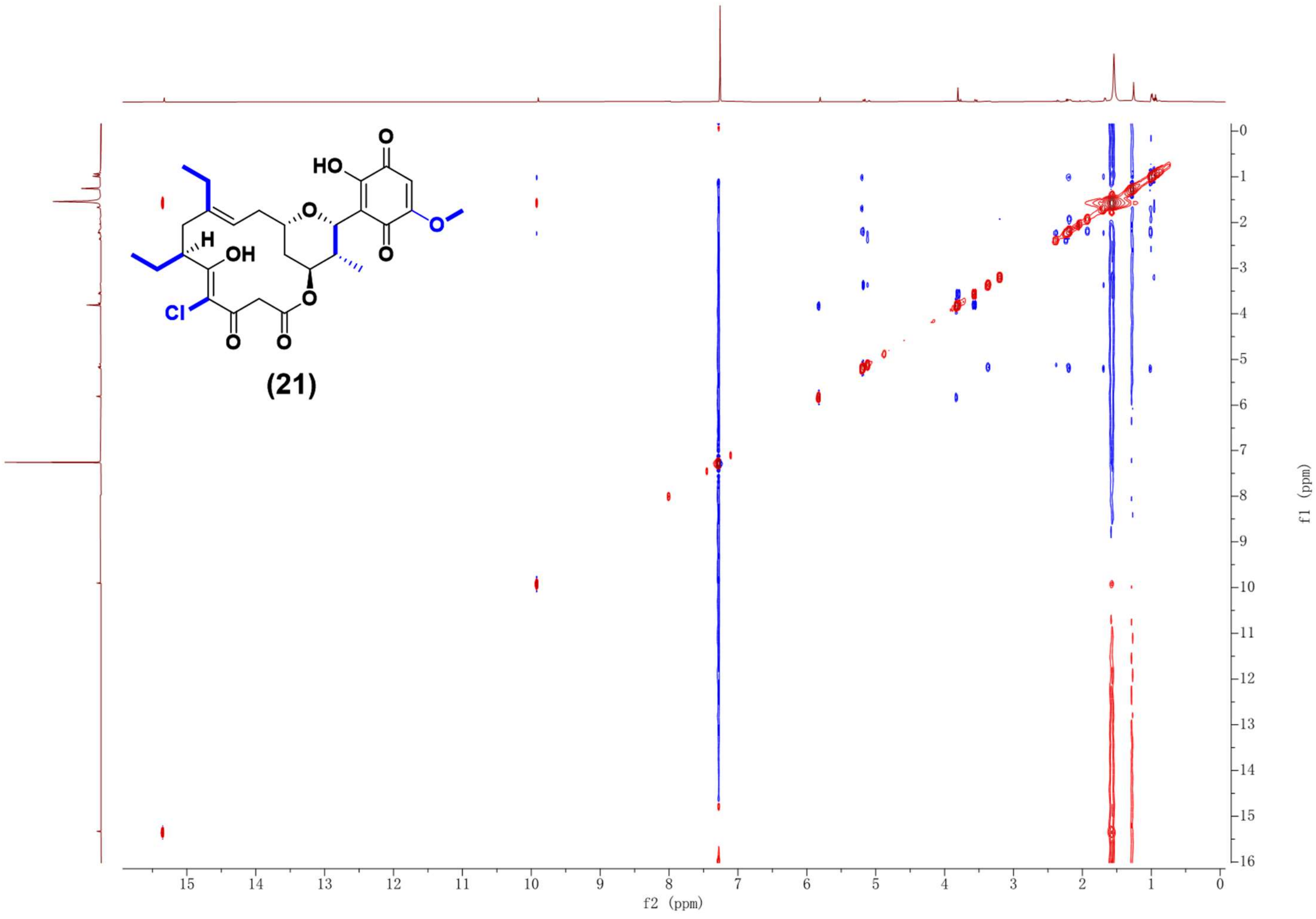

Supplementary Fig. 163. NOESY NMR spectrum (600 MHz, $CDCl_3$) of compound **21**.
